# Screening and identification of key biomarkers in clear cell renal cell carcinoma based on bioinformatics analysis

**DOI:** 10.1101/2020.12.21.423889

**Authors:** Basavaraj Vastrad, Chanabasayya Vastrad, Iranna Kotturshetti

## Abstract

Clear cell renal cell carcinoma (ccRCC) is one of the most common types of malignancy of the urinary system. The pathogenesis and effective diagnosis of ccRCC have become popular topics for research in the previous decade. In the current study, an integrated bioinformatics analysis was performed to identify core genes associated in ccRCC. An expression dataset (GSE105261) was downloaded from the Gene Expression Omnibus database, and included 26 ccRCC and 9 normal kideny samples. Assessment of the microarray dataset led to the recognition of differentially expressed genes (DEGs), which was subsequently used for pathway and gene ontology (GO) enrichment analysis. This data was utilized in the construction of the protein-protein interaction network and module analysis was conducted using Human Integrated Protein-Protein Interaction rEference (HIPPIE) and Cytoscape software. In addation, target gene - miRNA regulatory network and target gene - TF regulatory network were constructed and analysed. Finally, hub genes were validated by survival analysis, expression analysis, stage analysis, mutation analysis, immune histochemical analysis, receiver operating characteristic (ROC) curve analysis, RT-PCR and immune infiltration analysis. The results of these analyses led to the identification of a total of 930 DEGs, including 469 up regulated and 461 down regulated genes. The pathwayes and GO found to be enriched in the DEGs (up and down regulated genes) were dTMP de novo biosynthesis, glycolysis, 4-hydroxyproline degradation, fatty acid beta-oxidation (peroxisome), cytokine, defense response, renal system development and organic acid metabolic process. Hub genes were identified from PPI network according to the node degree, betweenness centrality, stress centrality, closeness centrality and clustering coefficient. Similarly, targate genes were identified from target gene - miRNA regulatory network and target gene - TF regulatory network according to the node degree. Furthermore, survival analysis, expression analysis, stage analysis, mutation analysis, immune histochemical analysis, ROC curve analysis, RT-PCR and immune infiltration analysis revealed that CANX, SHMT2, IFI16, P4HB, CALU, CDH1, ERBB2, NEDD4L, TFAP2A and SORT1 may be associated in the tumorigenesis, advancement or prognosis of ccRCC. In conclusion, the 10 hub genes diagonised in the current study may help researchers in exemplify the molecular mechanisms linked with the tumorigenesis and advancement of ccRCC, and may be powerful and favorable candidate biomarkers for the prognosis, diagnosis and treatment of ccRCC.

## Introduction

Clear cell renal cell carcinoma (ccRCC) is the most common malignancy of the kidney [Heng et al. 2009]. ccRCC ranks 14^th^ in cancer deaths worldwide, responsible for about 403,262 new cases and 175,098 deaths last year [Bray et al. 2018]. Although we have made great development on the early prognosis, diagnosis and new therapy, ccRCC still is one of challengeable diseases in this world [Jiang et al. 2006; Pan et al. 2004; Wood and Margulis, 2009]. However, the molecular mechanisms of ccRCC are not well understood. And due to the absence of specific biomarkers, most ccRCC patients are diagnosed at a late stage, leading to particularly poor outcomes of patients [Patel et al. 2016]. Even worse, some of ccRCC patients suffer from tumor recurrence due to the cancer drug resistance [Mickisch et al. 1990] and nephrectomy [Eggener et al. 2006]. Therefore, it is of outstanding importance to find novel biomarkers, pathways and effective targets for ccRCC patients.

Similar to that of other cancers, the process of ccRCC initiation, progression and invasion of cancer cells associates’ genetic aberrations, and changes in the cancer microenvironment [Slaton et al. 2001]. The literature has also reported that the development and occurrence of ccRCC are closely associated with a variety of genes such as von Hippel-Lindau (VHL) [Nickerson et al. 2008], AL-1/4.1B [Yamada et al. 2006], HIF1A [Wiesener et al. 2001], FOXM1 [Xue et al. 2012] and KISS1R [Chen et al. 2011], and cellular pathways such as IL-6/STAT-3 pathway [Cuadros et al. 2014], notch and TGF-β signaling pathways [Sjölund et al. 2011], MYC pathway [Tang et al. 2009], EGFR/MMP-9 signaling pathway [Liang et al. 2012] and phosphoinositide 3-kinase/Akt pathway [Sourbier et al. 2006]. Thus, it is vital to identify effective early diagnostic methods progression development, in order to intervene in the advancement of the disease at the early stages.

Microarray technology is a high-throughput and powerful tool to make large quantities of data such as gene expression and DNA methylation [Giltnane and Rimm, 2004]. To explore the differential expressed genes (DEGs) in metastatic ccCRCC, diagnose new high-specificity and high-sensitivity tumor biomarkers, and identify effective therapeutic targets, in silico methods were used to study ccRCC. In the current study, GSE105261 was downloaded and analyzed from the Gene Expression Omnibus (GEO) database to obtain DEGs between metastatic ccRCC tissues and normal renal tissues. Subsequently, pathway enrichment analysis, gene ontology (GO) enrichment analysis, protein-protein interaction (PPI) network analysis, module analysis, target gene - miRNA regulatory network analysis and target gene - TF regulatory network analysis were performed to characterize the molecular mechanisms underlying carcinogenesis and progression of ccRCC. Validation of hub genes was performed by using survival analysis, expression analysis, stage analysis, mutation analysis, immune histochemical analysis (IHC) receiver operating characteristic (ROC) curve, RT-PCR and immune infiltration analysis. In conclusion, the present study identified a number of important genes that are associated with the molecular mechanism of ccRCC by integrated bioinformatical analysis, and these genes and regulatory networks may assist with identifying potential gene therapy targets for ccRCC. Our study results should provide novel insights for predicting the risk of ccRCC.

## Materials and methods

### Microarray data source, data preprocessing and identification of DEGs

In the current study, gene microarray datasets comparing the gene expression profiles between metastatic ccRCC tissues and normal renal tissues were downloaded from the GEO (http://www.ncbi.nlm.nih.gov/gds/). The accession number was GSE105261 was submitted by Nam et al (2019). The microarray data of GSE105261 was assessed using the GPL10558 Illumina HumanHT-12 V4.0 expression beadchip. The gene microarray data comprised 26 metastatic ccRCC tissue samples and 9 normal renal tissue samples. IDAT files were converted to expression measures and normalized via the beadarray package in R bioconductor [Dunning et al. 2006]. The aberrantly expressed mRNAs were subsequently determined using the Limma package in R bioconductor [Ritchie et al. 2015], based on the Benjamini and Hochberg method [Kvam et al. 2012]. DEGs between metastatic ccRCC tissues and normal renal tissues were defined by the cut-off criterion of fold change >1.11 for up regulated genes, fold change >1.00 for down regulated genes and a P-value of <0.05.

### Pathway enrichment analysis of DEGs

Pathway enrichment analyses of DEGs were performed according to the enrichment analysis tool of ToppGene (ToppFun) (https://toppgene.cchmc.org/enrichment.jsp) [Chen et al 2009] which integrates various pathway databases such as Kyoto Encyclopedia of Genes and Genomes (KEGG; http://www.genome.jp/kegg/) [Aoki-Kinoshita and Kanehisa, 2007], Pathway Interaction Database (PID, http://pid.nci.nih.gov/) [Schaefer et al 2009], Reactome (https://reactome.org/PathwayBrowser/) [Croft et al 2011], Molecular signatures database (MSigDB, http://software.broadinstitute.org/gsea/msigdb/) [Liberzon et al 2011], GenMAPP (http://www.genmapp.org/) [Dahlquist et al 2002], Pathway Ontology (https://bioportal.bioontology.org/ontologies/PW) [Petri et al 2014], PantherDB (http://www.pantherdb.org/) [Mi et al 2013] and Small Molecule Pathway Database (SMPDB, http://smpdb.ca/) with the cutoff criterion of p value <0.05.

### GO enrichment analysis of DEGs

ToppGene (ToppFun) (https://toppgene.cchmc.org/enrichment.jsp) [Chen et al 2009] is an online biological information database that integrates biological data and analysis tools. GO (http://www.geneontology.org/) [Harris et al 2004] was used to perform bioinformatics analysis and annotate function enrichment of genes. ToppGene was applied to perform the function enrichment and biological analyses of these DEGs with categories of biological processes (BP), cellular component (CC) and molecular function (MF). P<0.05 was considered statistically significant.

### PPI network construction and module analysis

Human Integrated Protein-Protein Interaction rEference (HIPPIE) (http://cbdm.uni-mainz.de/hippie/) [Alanis-Lobato et al. 2017] is a online tool that studies genetic interactions and can assess PPI network and this online tool integrates various PPI databases such as IntAct (https://www.ebi.ac.uk/intact/) [Orchard et al. 2014], BioGRID (https://thebiogrid.org/) [Chatr-Aryamontri et al. 2017], HPRD (http://www.hprd.org/) [Keshava Prasad et al. 2009], MINT (https://mint.bio.uniroma2.it/) [Licata et al. 2012], BIND (http://download.baderlab.org/BINDTranslation/) [Isserlin et al. 2011], MIPS (http://mips.helmholtz-muenchen.de/proj/ppi/) [Pagel et al. 2005] and DIP (http://dip.doe-mbi.ucla.edu/dip/Main.cgi) [Salwinski et al. 2004]. Therefore, the HIPPIE database in Cytoscape version 3.7.2 (http://www.cytoscape.org/) [Shannon et al 2003] was used to identify and map potential associations between the DEGs (up and down regulated genes). Network Analyzer is an application of Cytoscape, and was used for calculating the topological properties such as node degree [Wang et al 2014], betweenness centrality [Peng et al 2015], stress centrality [Carson and Lu, 2015], closeness centrality [Jalili et al 2016] and clustering coefficient [Wiles et al 2010] of hub genes in PPI network. Modules in the PPI network were extracted from Cytoscape using the PEWCC1 [Zaki et al 2013], with the following criteria: Degree cut-off, 2; node score cut-off, 0.2; k-core, 2; and maximum depth, 100.

### Construction of target genes - miRNA regulatory network

The micro RNA (miRNAs) interacts with targeted genes were predicted by two established miRNA target prediction databases such as DIANA-TarBase (http://diana.imis.athena-innovation.gr/DianaTools/index.php?r=tarbase/index) [Vlachos et al 2015] and miRTarBase (http://mirtarbase.mbc.nctu.edu.tw/php/download.php) [Chou et al 2018]. The miRNAs predicted by online tool NetworkAnalyst (https://www.networkanalyst.ca/) [Zhou et al 2019] selected as the targeted genes and converted the results visualized by using Cytoscape version 3.7.2 (http://www.cytoscape.org/) [Shannon et al 2003]. In the network, a blue diamond node represented the miRNA and a green and red circular nodes represented the up and down regulated target gene, their interaction was represented by an line. The numbers of lines in the networks indicated the contribution of one miRNA to the surrounding target genes, and the higher the degree, the more central the target gene was within the network.

### Construction of target genes - TF regulatory network

The transcription factors (TFs) interacts with targeted genes were predicted by established TF target prediction database JASPAR (http://jaspar.genereg.net/) [Khan et al 2018]. The TFs predicted by online tool NetworkAnalyst (https://www.networkanalyst.ca/) [Zhou et al 2019] selected as the targeted genes and converted the results visualized by using Cytoscape version 3.7.2 (http://www.cytoscape.org/) [Shannon et al 2003]. In the network, a blue and yellow triangle node represented the TF and a green and red circular nodes represented the up and down regulated target gene, their interaction was represented by an line. Te numbers of lines in the networks indicated the contribution of one TF to the surrounding target genes, and the higher the degree, the more central the target gene was within the network.

### Validation of hub genes

After hub genes identified from microarray expression profile dataset, UALCAN (http://ualcan.path.uab.edu/analysis.html) [Chandrashekar et al 2017], cBio Cancer Genomics Portal (http://www.cbioportal.org/) [Gao et al 2013] and The Human Protein Atlas (HPA) (https://www.proteinatlas.org/) [Uhlen et al 2010] were used to validate the selected up regulated and down regulated hub genes. UALCAN is an online tool for gene expression analysis between cancer patients and normal person’s data from The Cancer Genome Atlas (TCGA). UALCAN provides data such as survival period, gene expression and tumor staging. cBio Cancer Genomics Portal is an online tool for genetic alterations in cancer patients from TCGA database. cBio Cancer Genomics Portal provides data such as DNA mutations, methylations, gene amplifications, homozygous deletions, protein alterations and phosphorous abundance. The Human Protein Atlas (HPA) is an online tool for map all the human proteins in tissues, cells and organs. HPA provides data about expressions of hub genes in cancer tissue and its normal tissue. Receiver operating characteristic (ROC) curve was performed to evaluate the diagnosis value of hub genes in ccRCC using R package“pROC” [Robin et al 2011]. Total RNA of hub genes were segregated from cultured ccRCC cells using TRIzol reagent (Invitrogen) and converted into complementary DNA (EY001; Shanghai Iyunbio Co). LightCycler 96 Real-Time PCR Systems (Roche) was used to performed RT-PCR. The operations for the real-time polymerase chain reaction (RT-PCR) were as follows: 50C for 3 minutes, 95C for 3 minutes, 95C for 10 seconds, followed by 40 cycles of 95C for 10 seconds and 60C for 30 seconds. The sequences of the primers are shown in Table 1. The relative expression levels of genes were normalized to those of human β-actin, estimated using the 2^-ΔΔCT^ method and were log-transformed [Livak and Schmittgen, 2001]. The hub genes were further analyzed using online tool the TIMER (https://cistrome.shinyapps.io/timer/) [Li et al. 2017] for immune infiltration analysis. Immune cell types such as B cells, CD4+ T cells, CD8+ T cells, neutrophils, macrophages, and dendritic cells were assessed by TIMER on renal cancer sample data, and the correlation between 10 hub genes expression and immune infiltration.

**Table 1.**
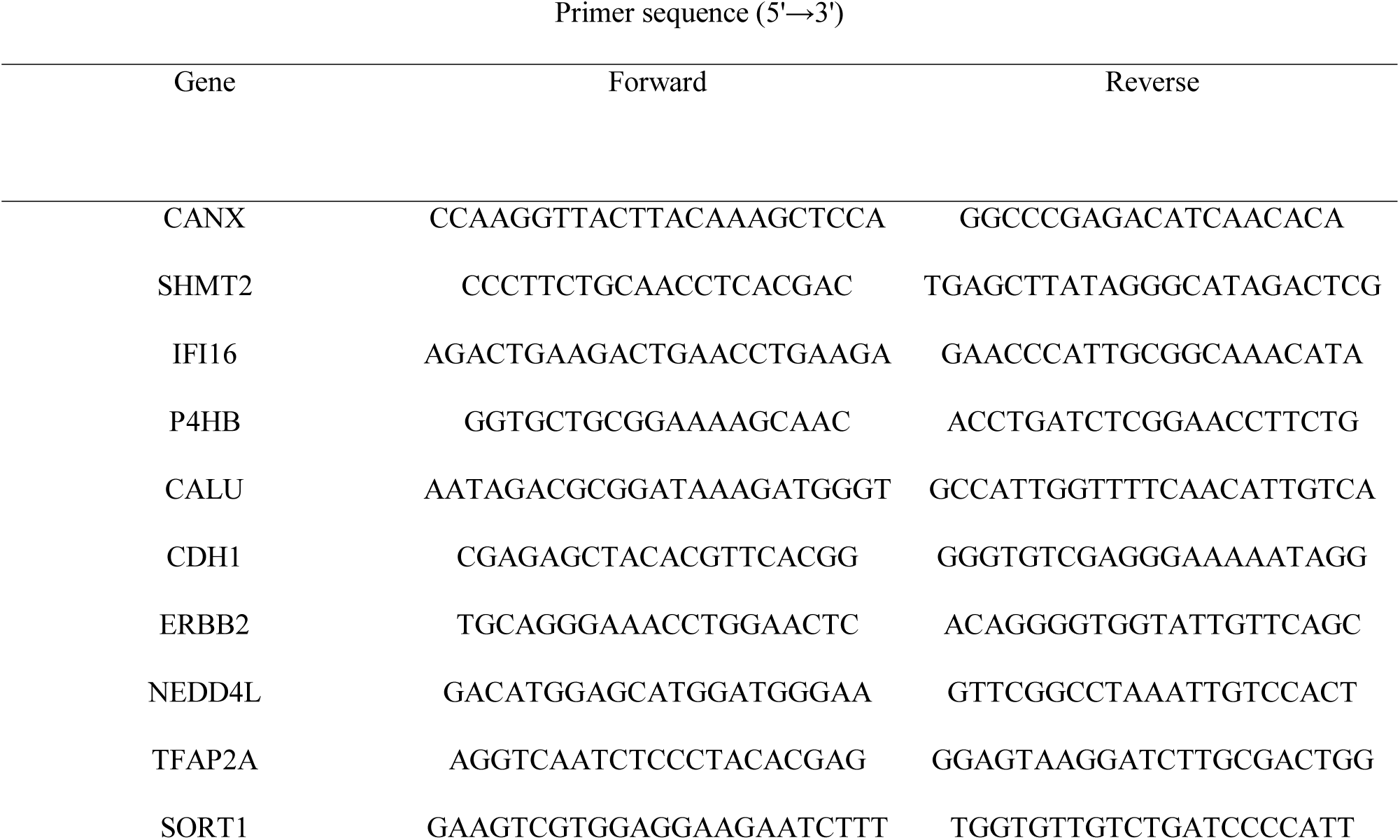
Primers used for quantitative PCR

## Results

### Identification of DEGs

The gene expression dataset GSE33455 was downloaded from the GEO database. After the standardization (normalization) of the microarray data, differentially expressed genes were identified between metastatic ccRCC tissues and normal renal tissues. Box plots were constructed before normalization and after normalization of microarray data (Fig. 1A and Fig. 1B). Upon preprocessing, 930 DEGs (P value <0.05; fold change > 1.11 for up regulated genes; fold change < -1 for down regulated genes) were identified, including 469 up regulated and 461 down regulated genes (Table 1). The volcano plot of gene expression profile data was shown in Fig. 2. The heat maps of the up regulated and down regulated genes are presented in Fig. 3 and Fig. 4.

**Fig. 1.**
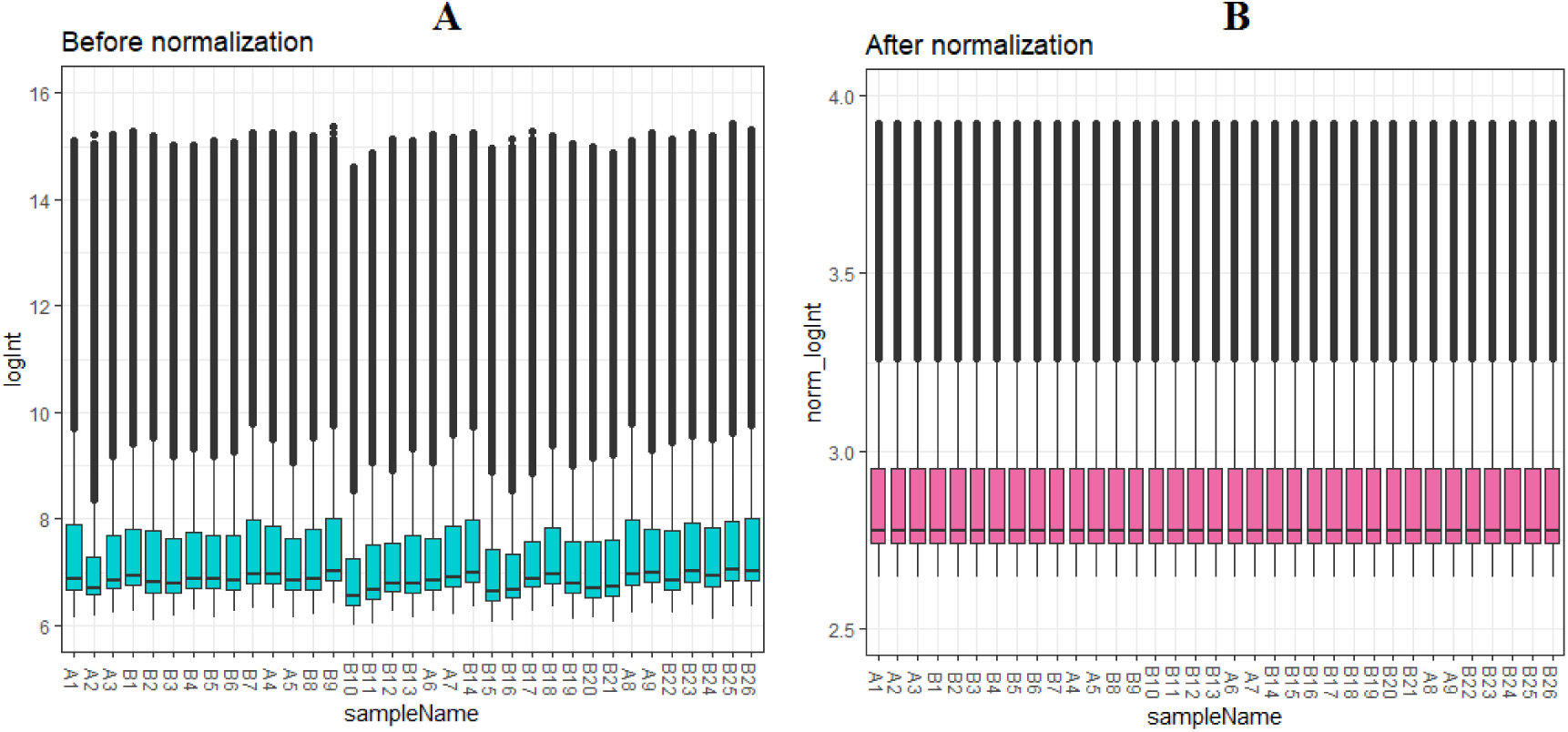
Box plots of the gene expression data before normalization (A) and after normalization (B). Horizontal axis represents the sample symbol and the vertical axis represents the gene expression values. The black line in the box plot represents the median value of gene expression. (A1 – A9 = normal renal tissues samples; B1 – B26 = metastatic ccRCC tissues samples)

**Fig. 2.**
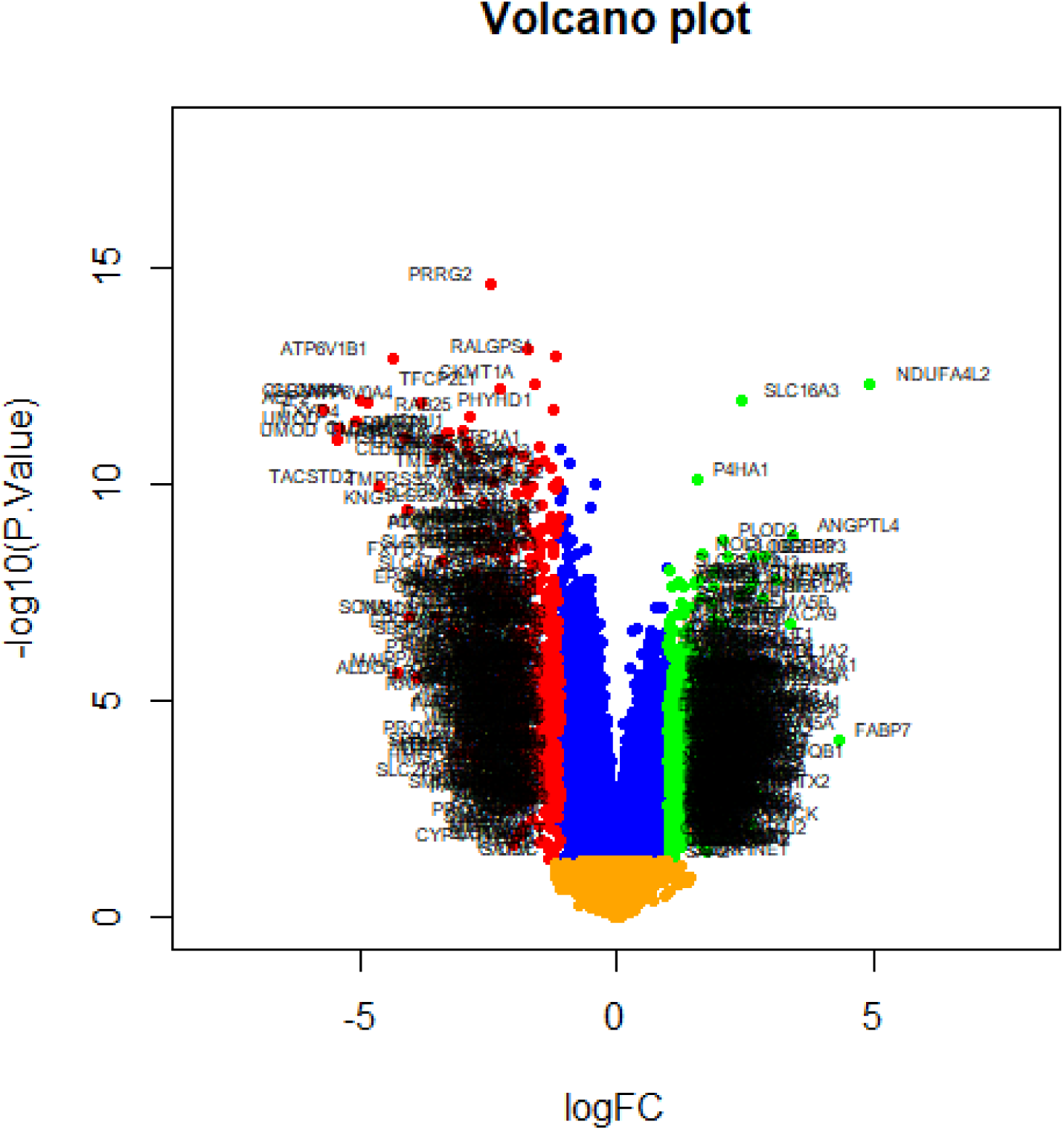
Volcano plot of differentially expressed genes. Genes with a significant change of more than two-fold were selected.

**Fig. 3.**
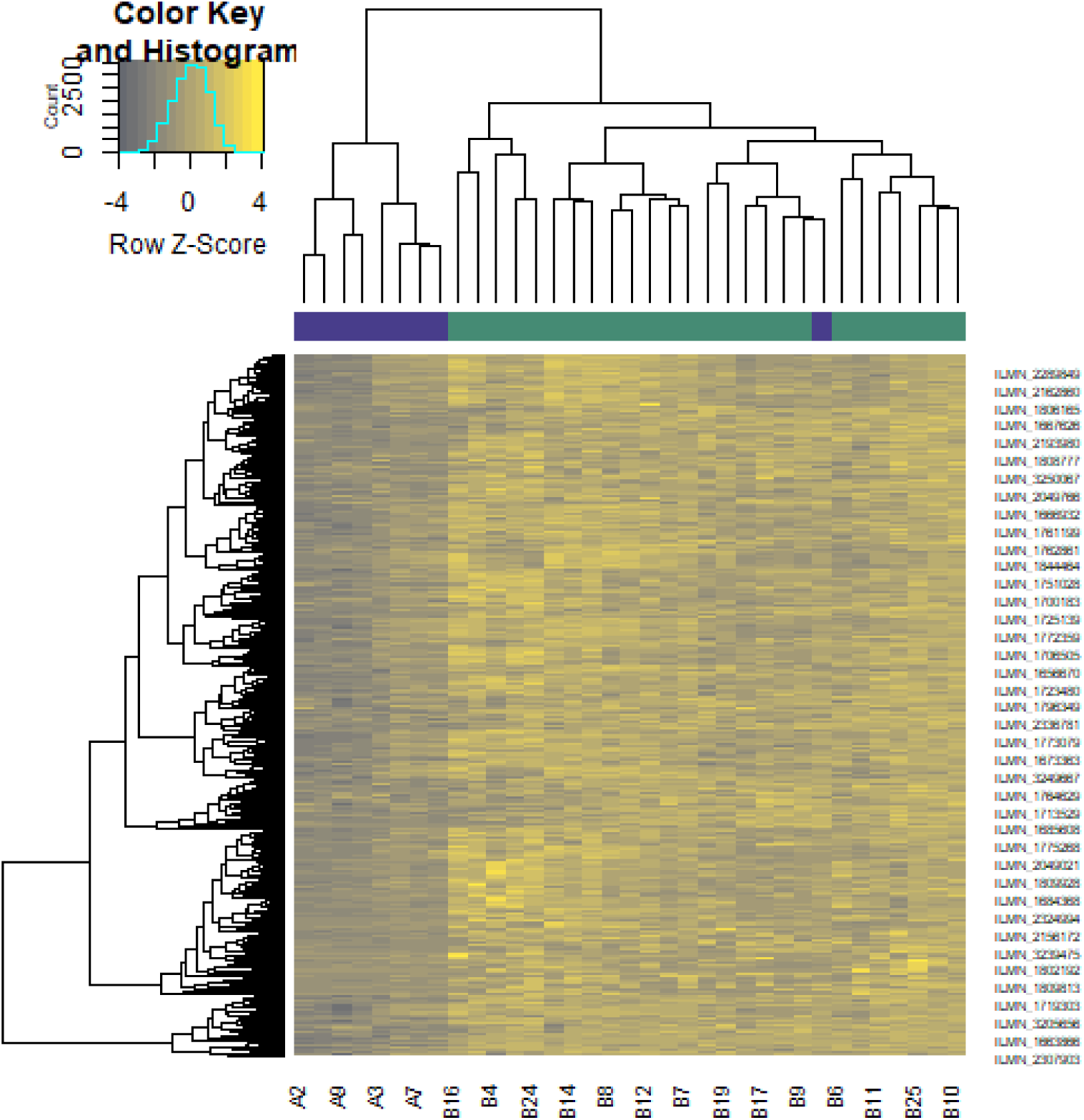
Heat map of up regulated differentially expressed genes. Legend on the top left indicate log fold change of genes. (A1 – A9 = normal renal tissues samples; B1 – B26 = metastatic ccRCC tissues samples)

**Fig. 4.**
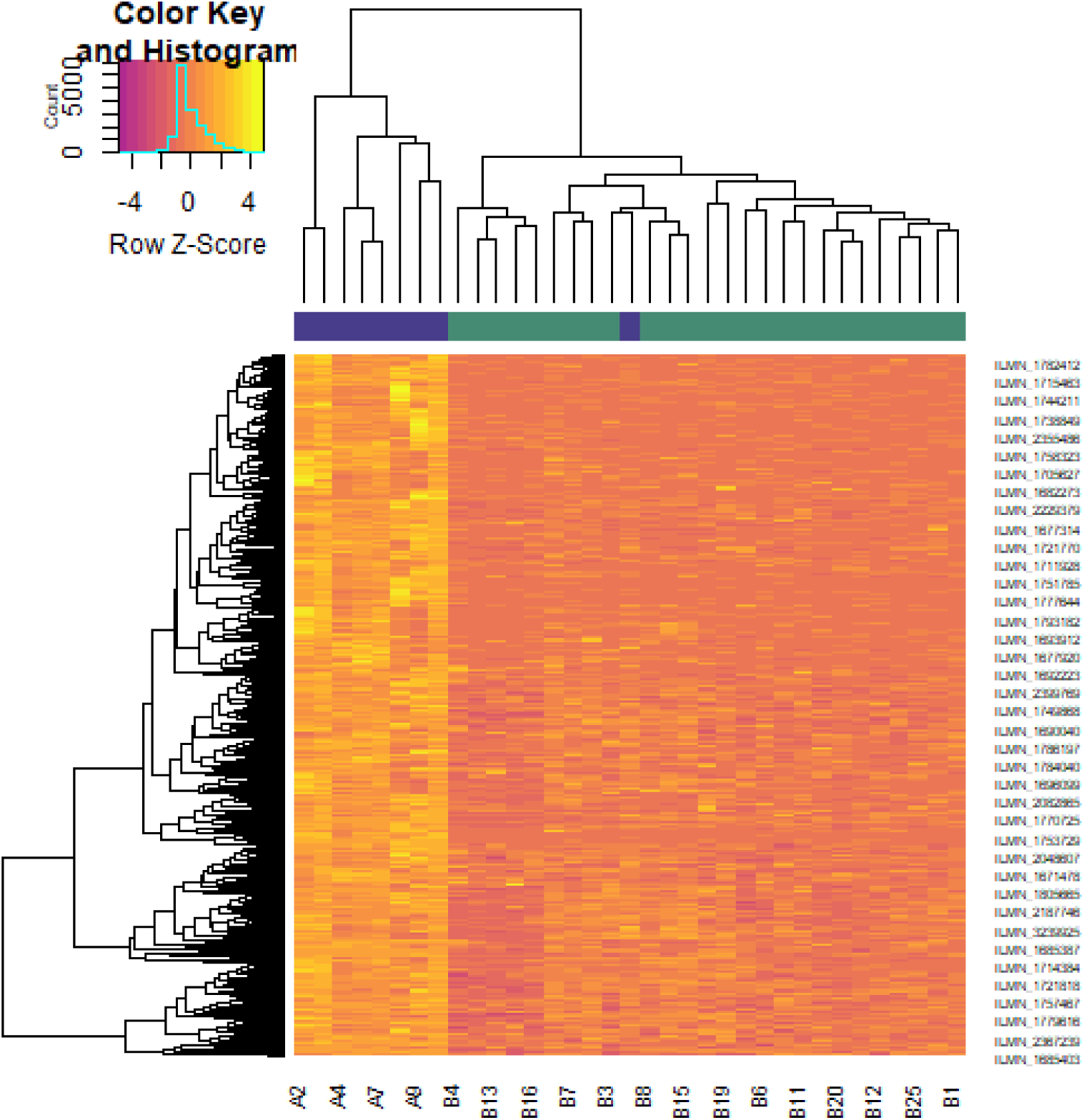
Heat map of down regulated differentially expressed genes. Legend on the top left indicate log fold change of genes. (A1 – A9 = normal renal tissues samples; B1 – B26 = metastatic ccRCC tissues samples)

### Pathway enrichment analysis

Pathway enrichment analyses of DEGs (up and down regulated genes) was performed using ToppGene. As for pathway enrichment, the up regulated genes were enriched in dTMP de novo biosynthesis, glycolysis, phagosome, complement and coagulation cascades, HIF-1-alpha transcription factor network, IL12-mediated signaling events, extracellular matrix organization, cytokine signaling in immune system, gluconeogenesis, one carbon pool by folate, ensemble of genes encoding extracellular matrix and extracellular matrix-associated proteins, ensemble of genes encoding core extracellular matrix including ECM glycoproteins, collagens and proteoglycans, integrin signalling pathway, plasminogen activating cascade, hypertension, reverse cholesterol transport and pyrimidine metabolism are listed in Table 2. Similarly, the down regulated genes were enriched in 4-hydroxyproline degradation, fatty acid beta-oxidation (peroxisome), metabolic pathways, aldosterone-regulated sodium reabsorption, FOXA2 and FOXA3 transcription factor networks, p75(NTR)-mediated signaling, transmembrane transport of small molecules, SLC-mediated transmembrane transport, urea cycle and metabolism of amino groups, valineleucine and isoleucine degradation, platelet amyloid precursor protein pathway, ensemble of genes encoding extracellular matrix and extracellular matrix-associated proteins, blood coagulation, 5-hydroxytryptamine degredation, pathway of urea cycle and metabolism of amino groups, gluconeogenesis pathway, polythiazide pathway and ramipril pathway are listed in Table 3.

**Table 2.**
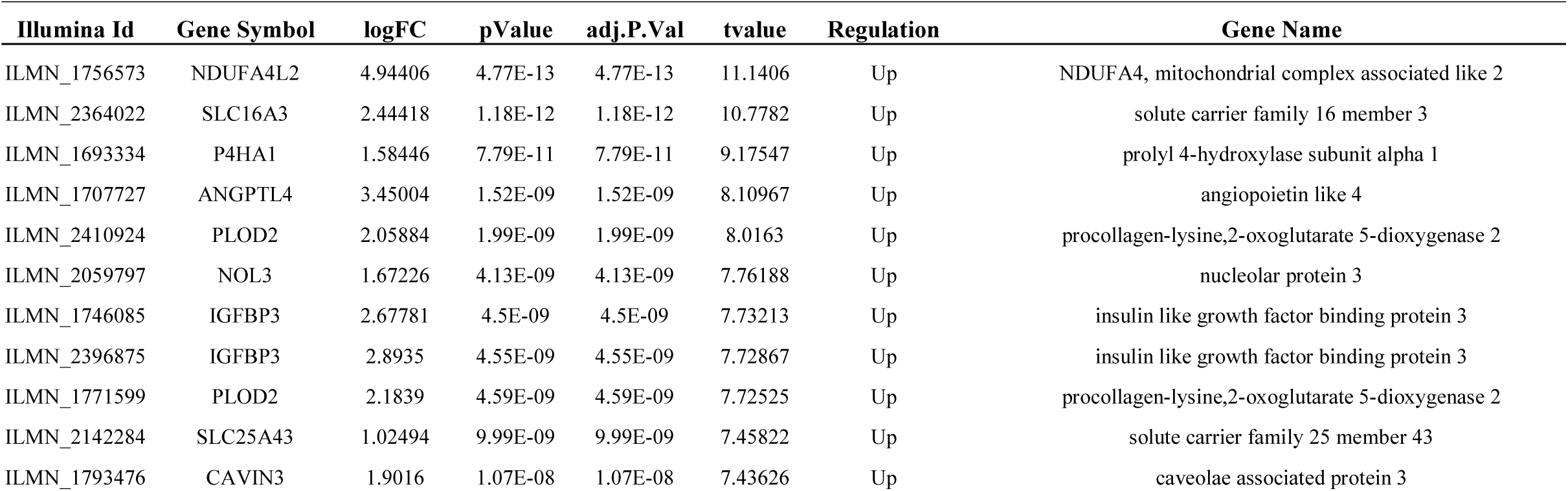
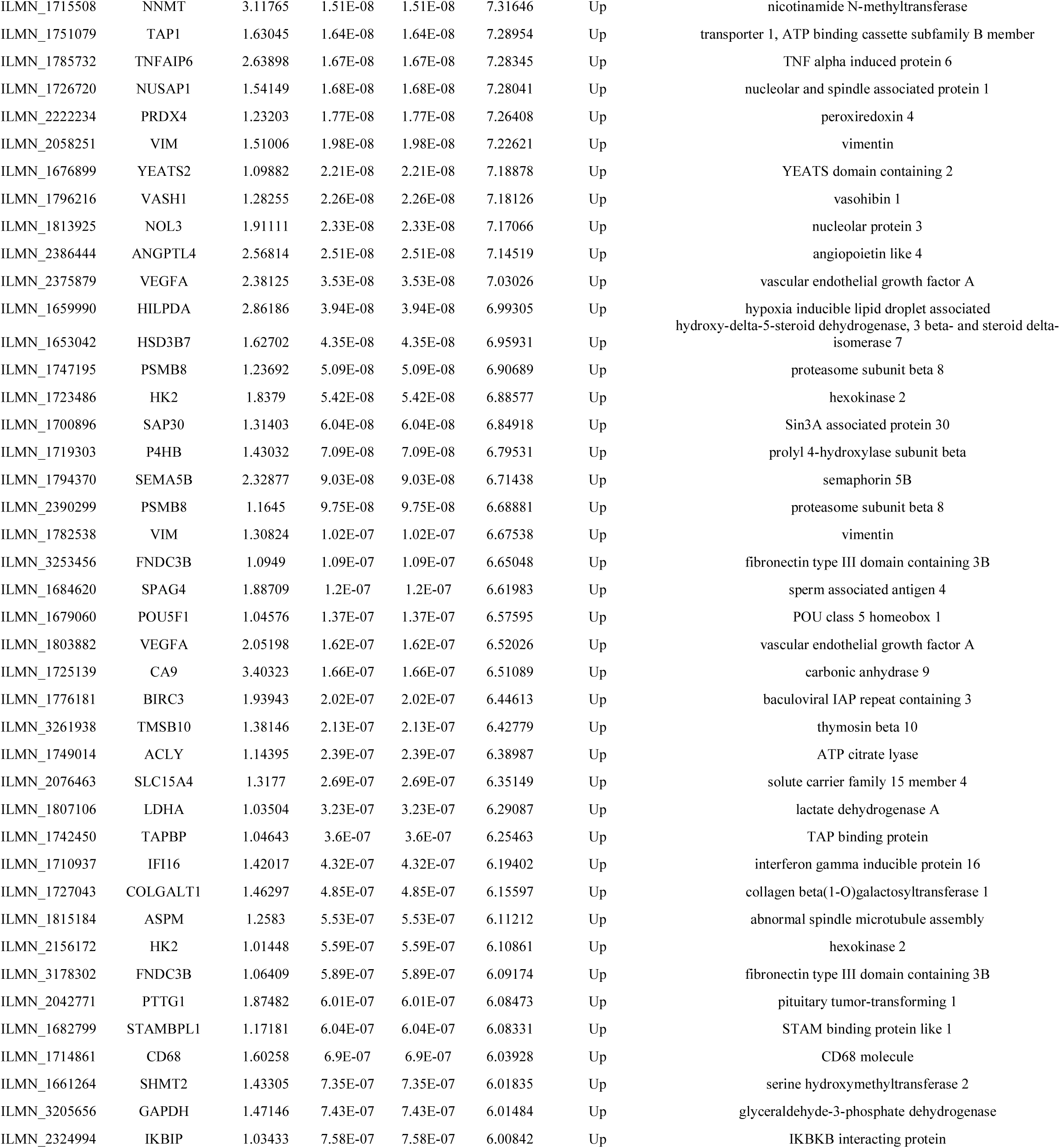
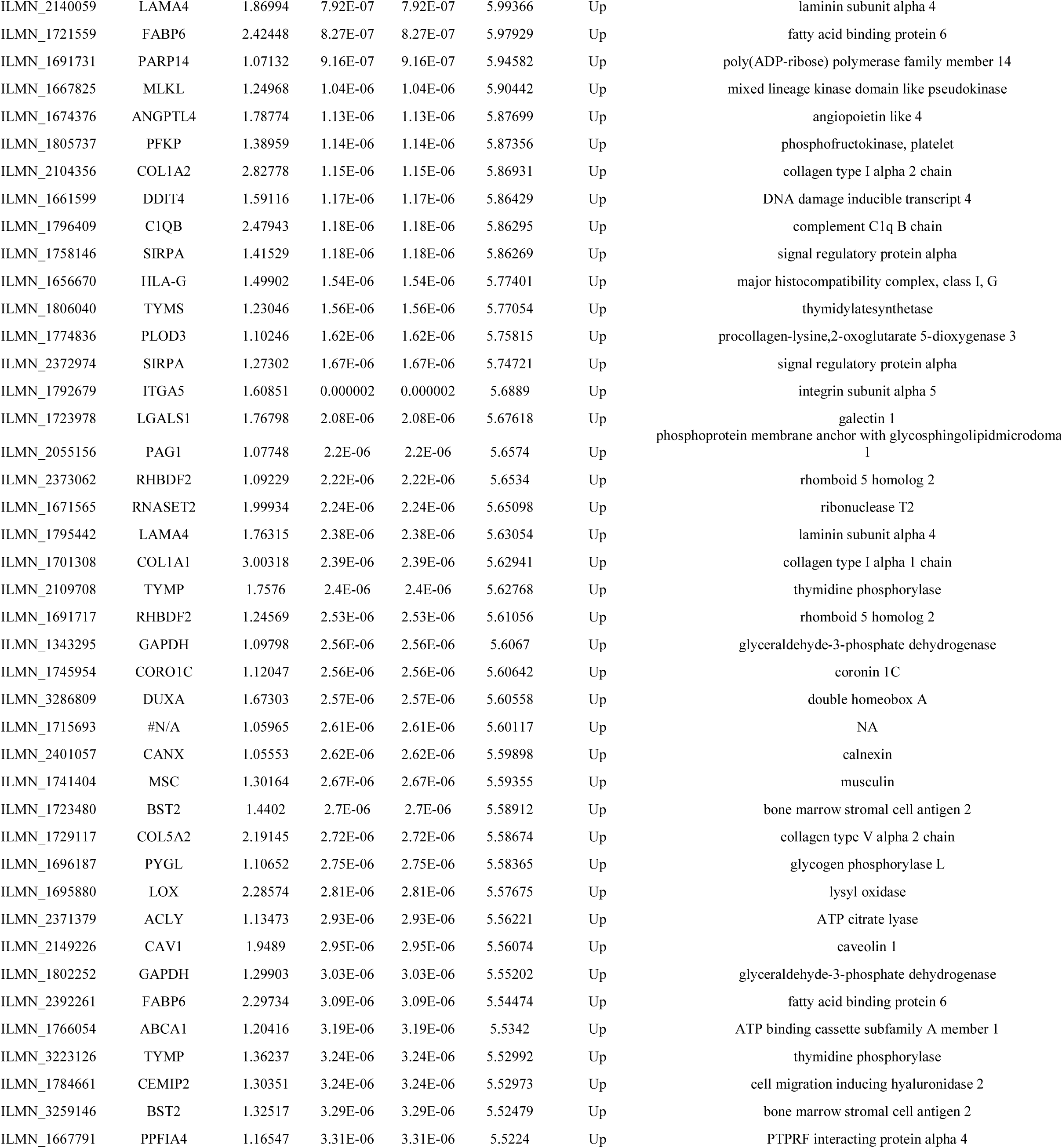
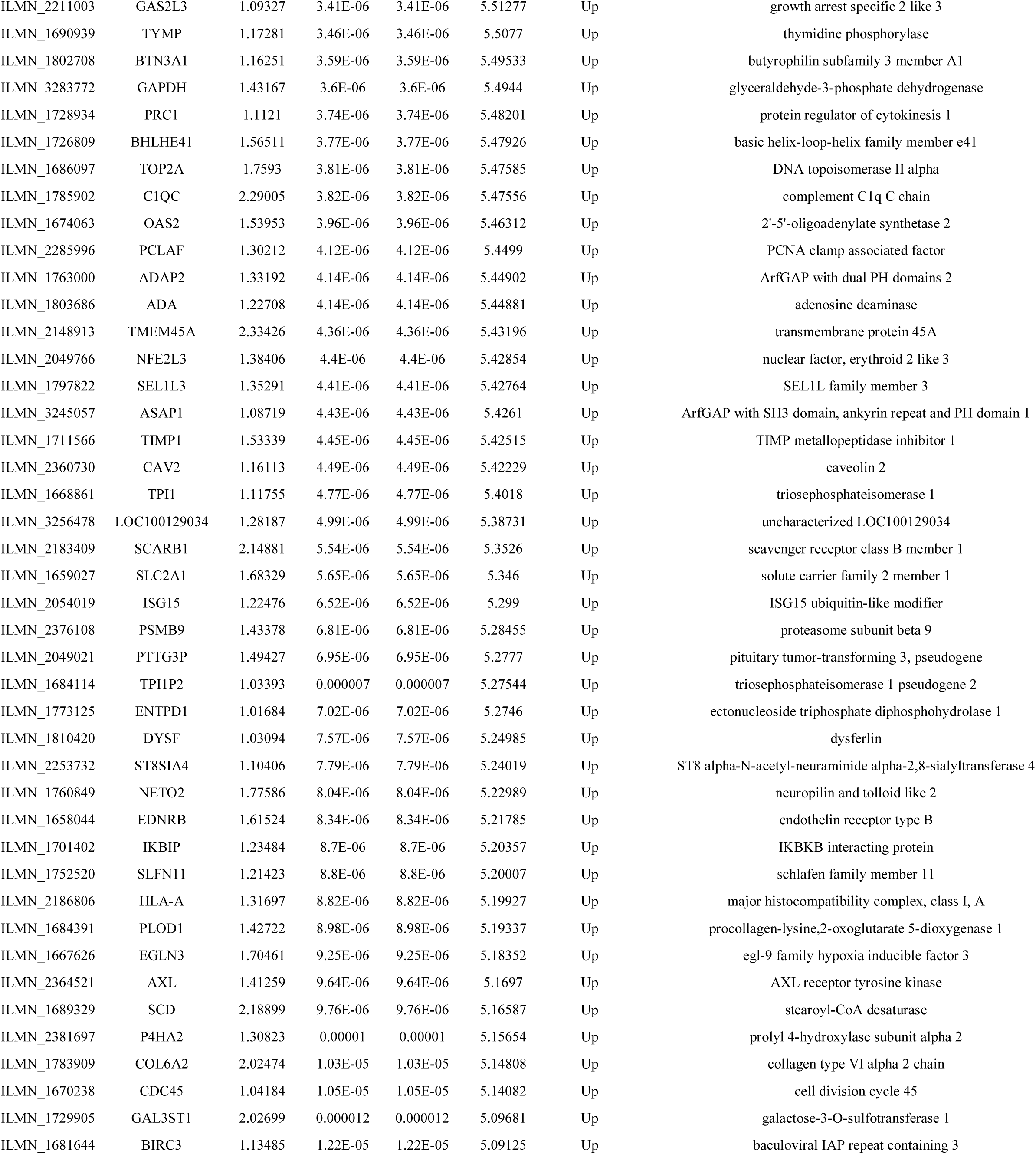
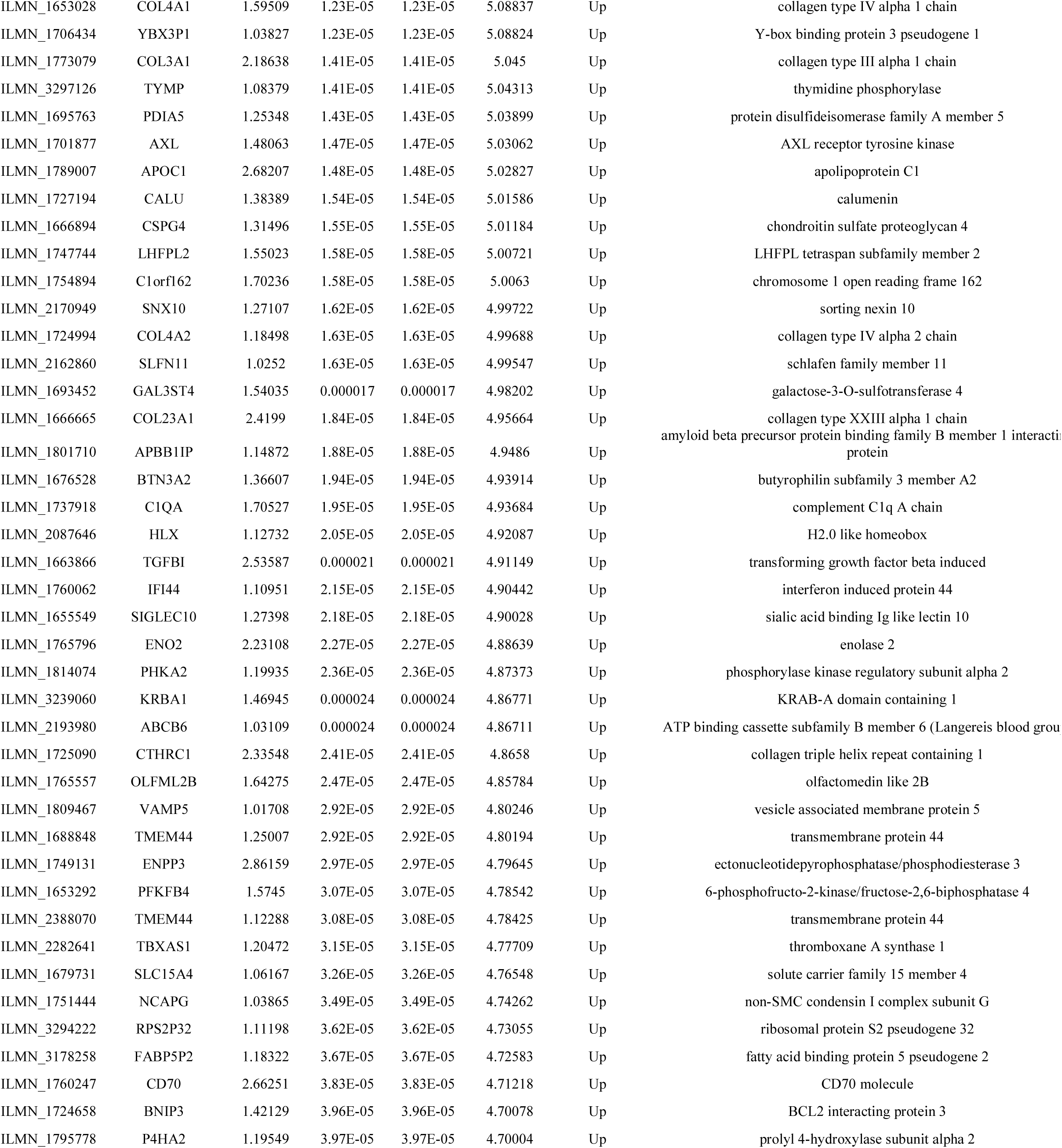
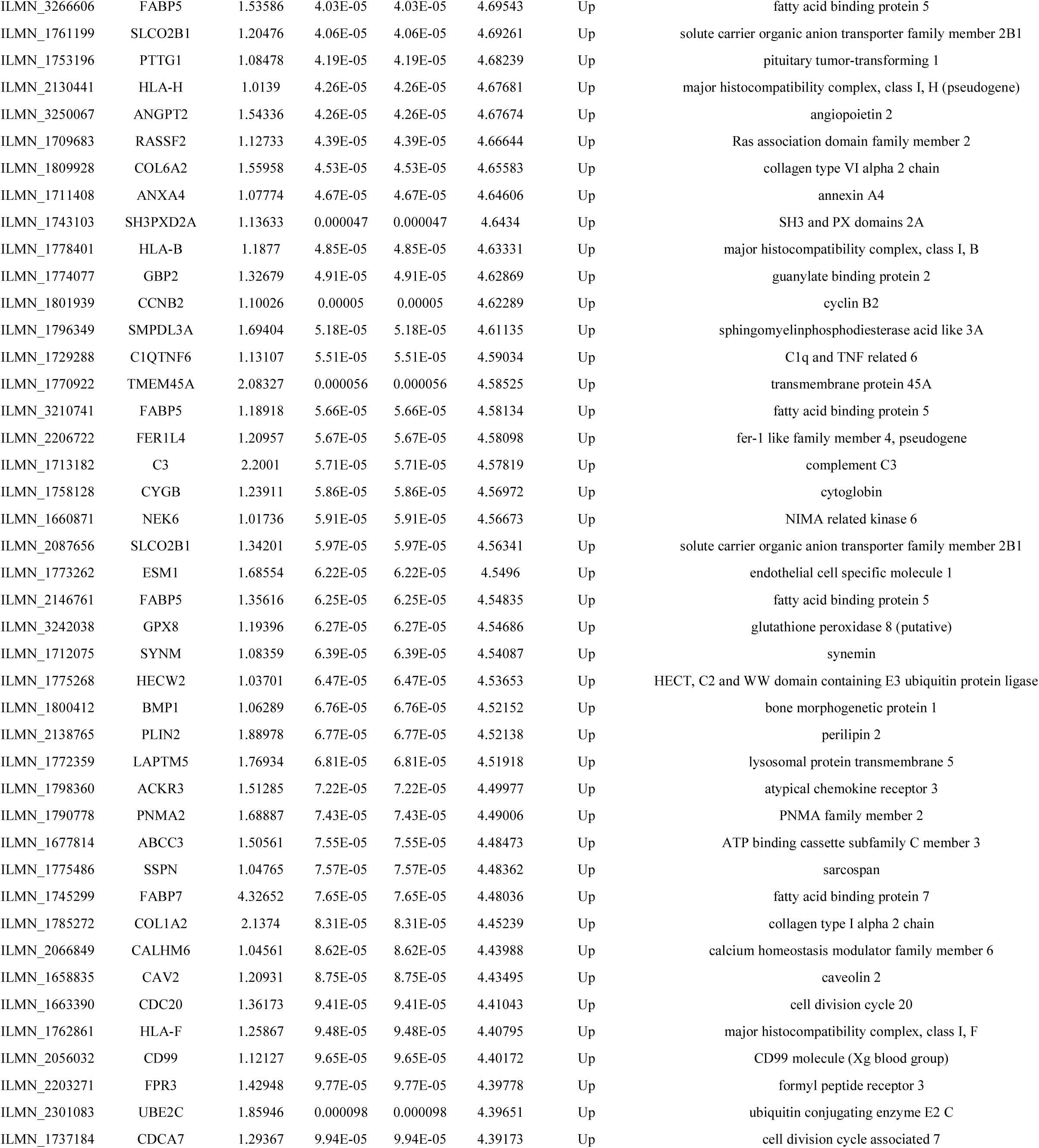
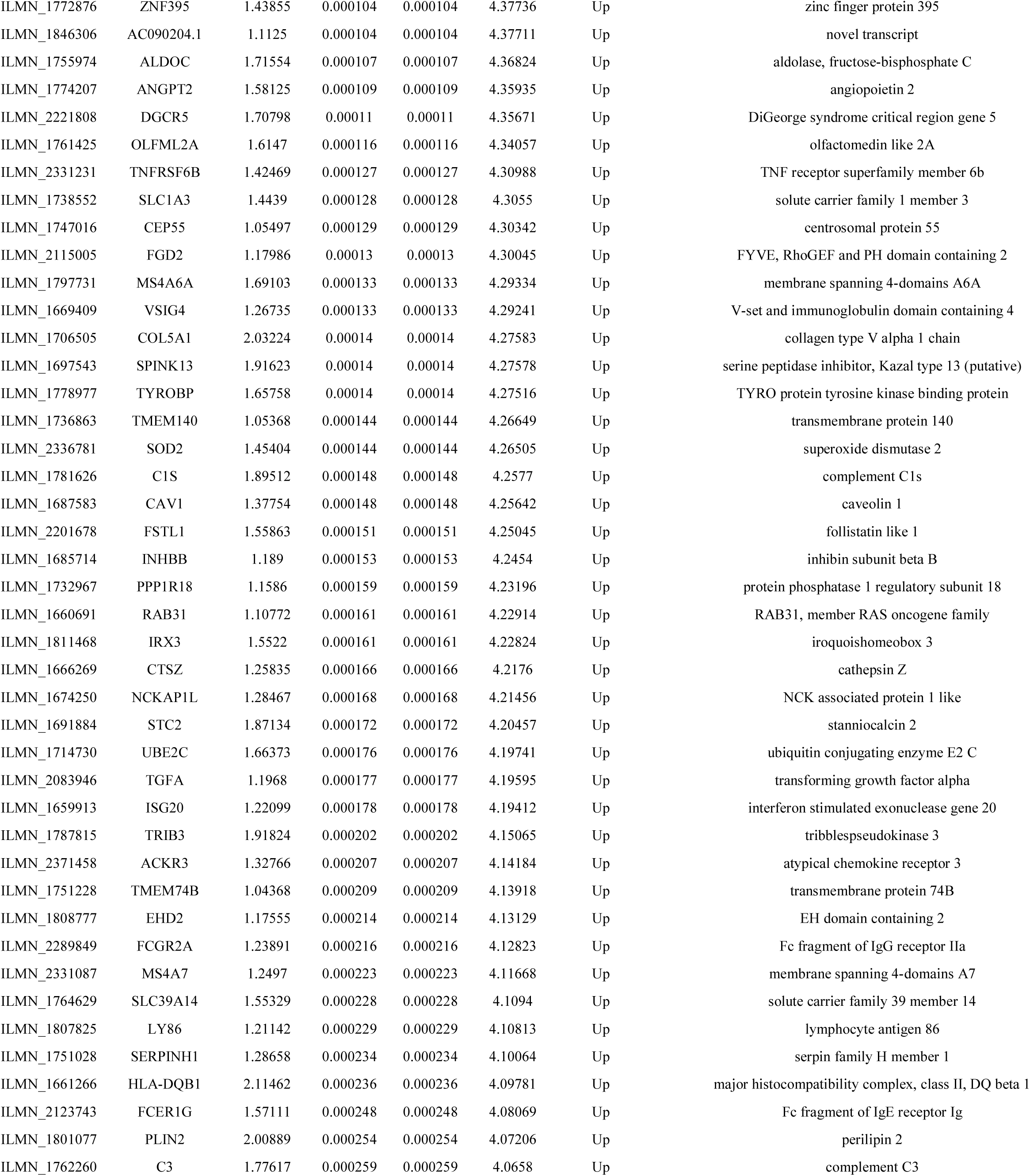
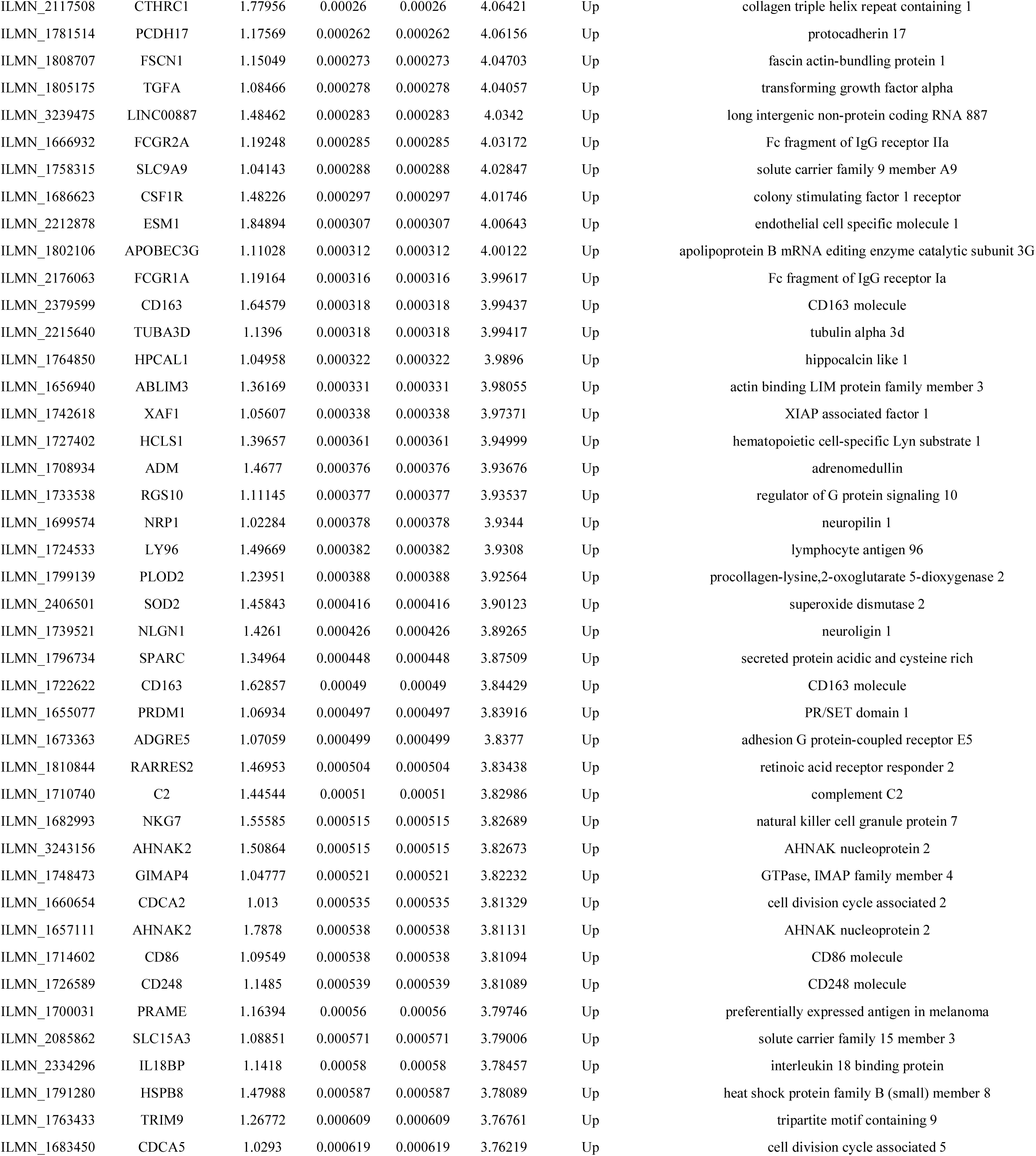
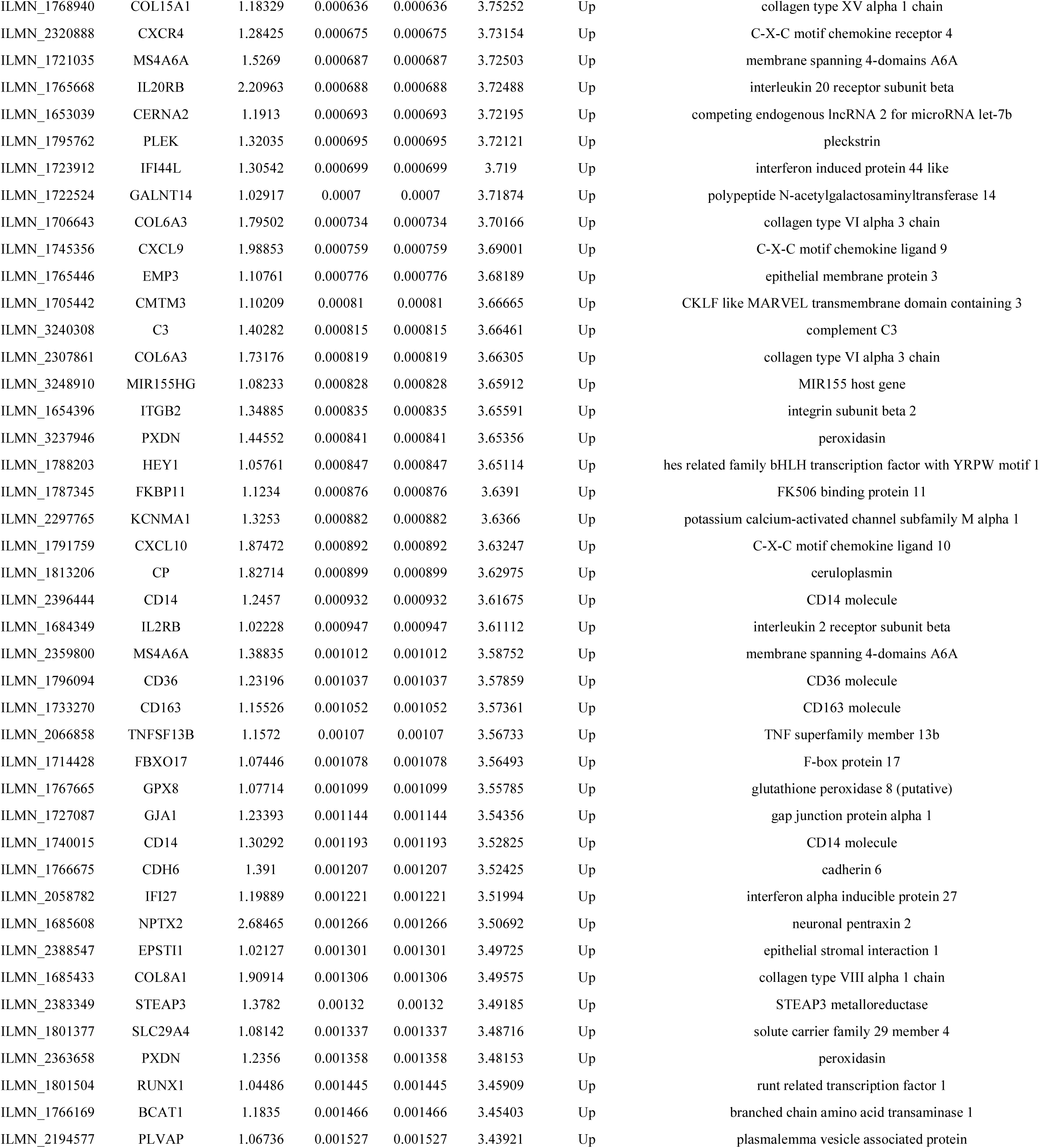
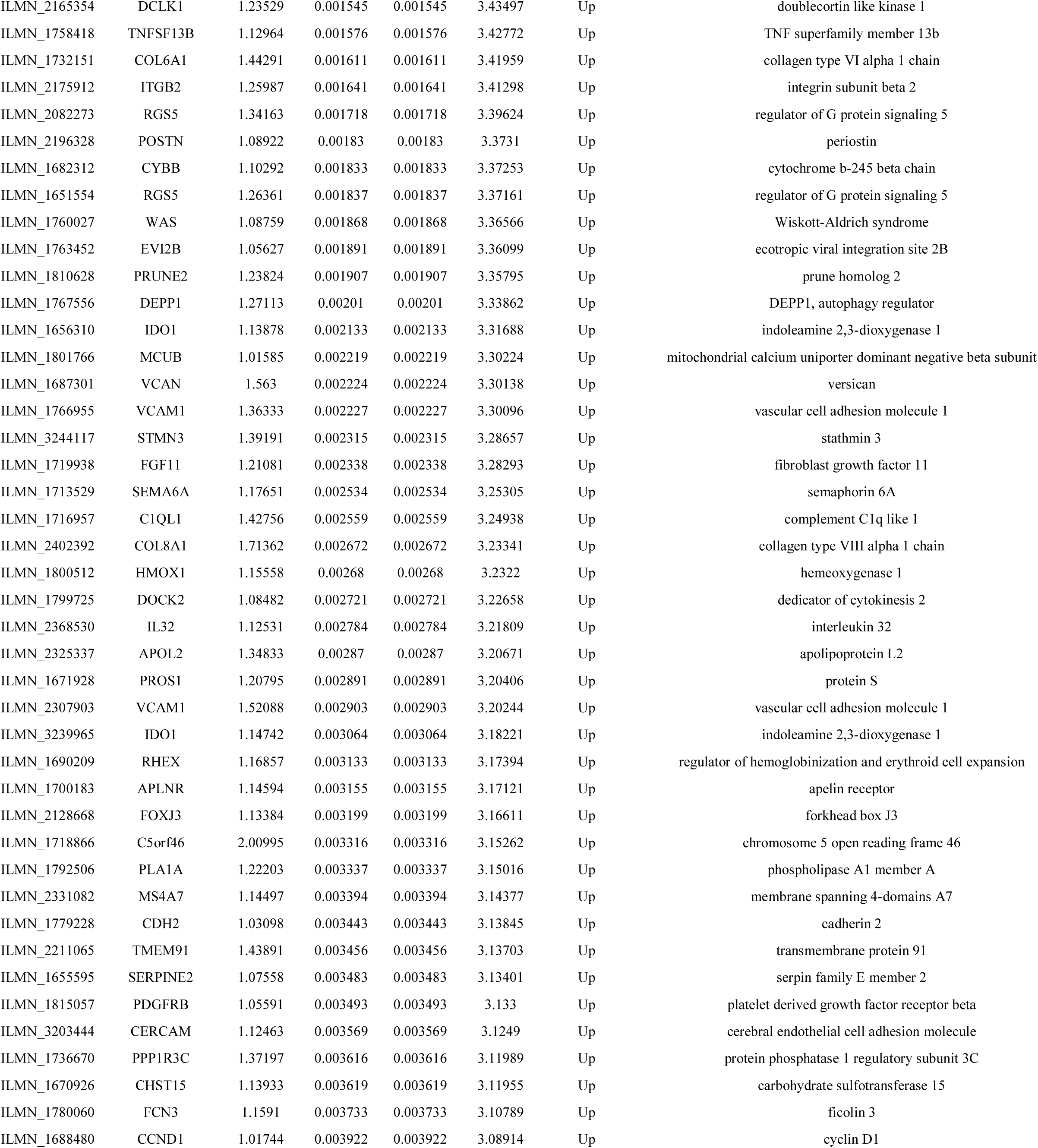
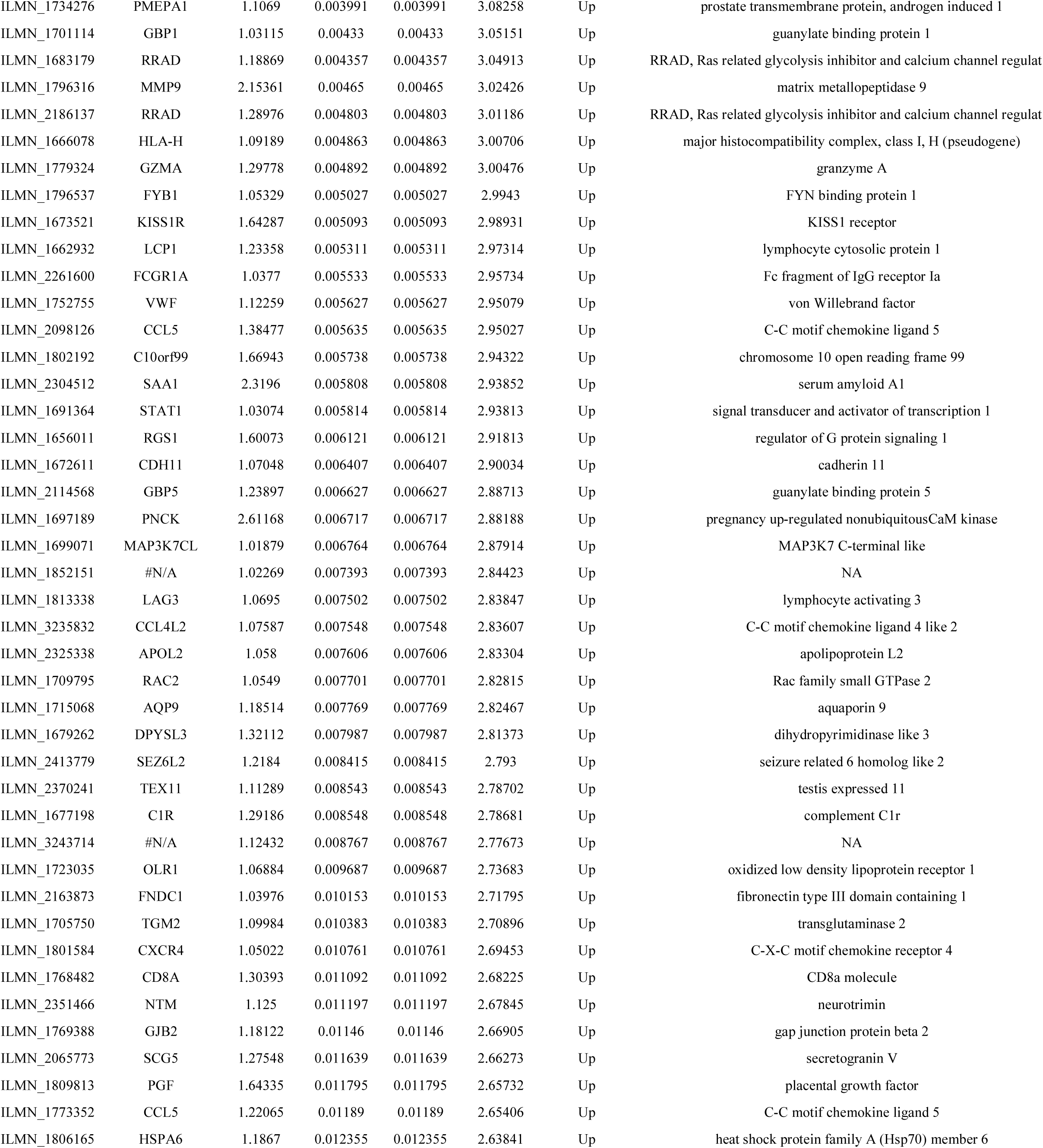
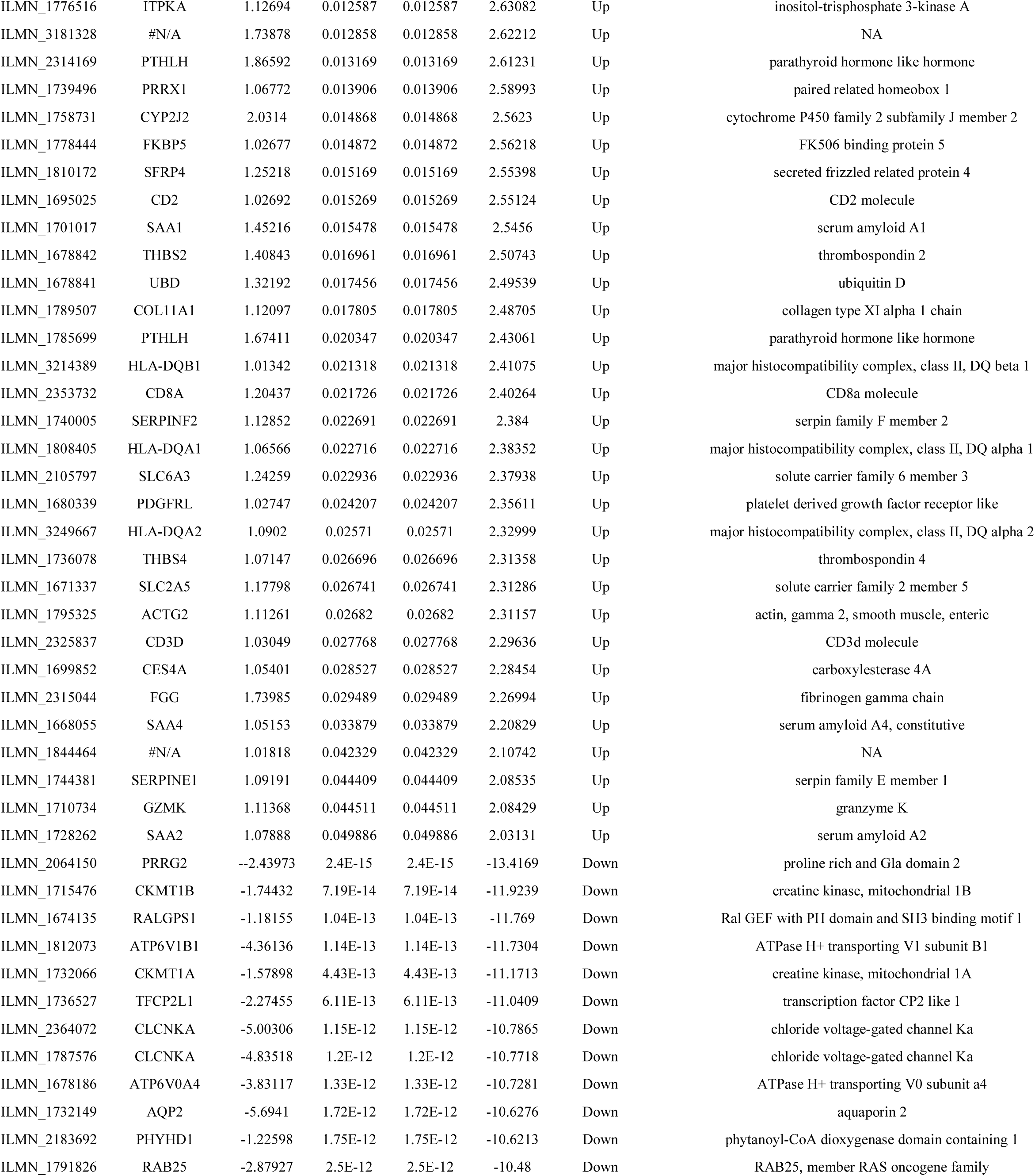
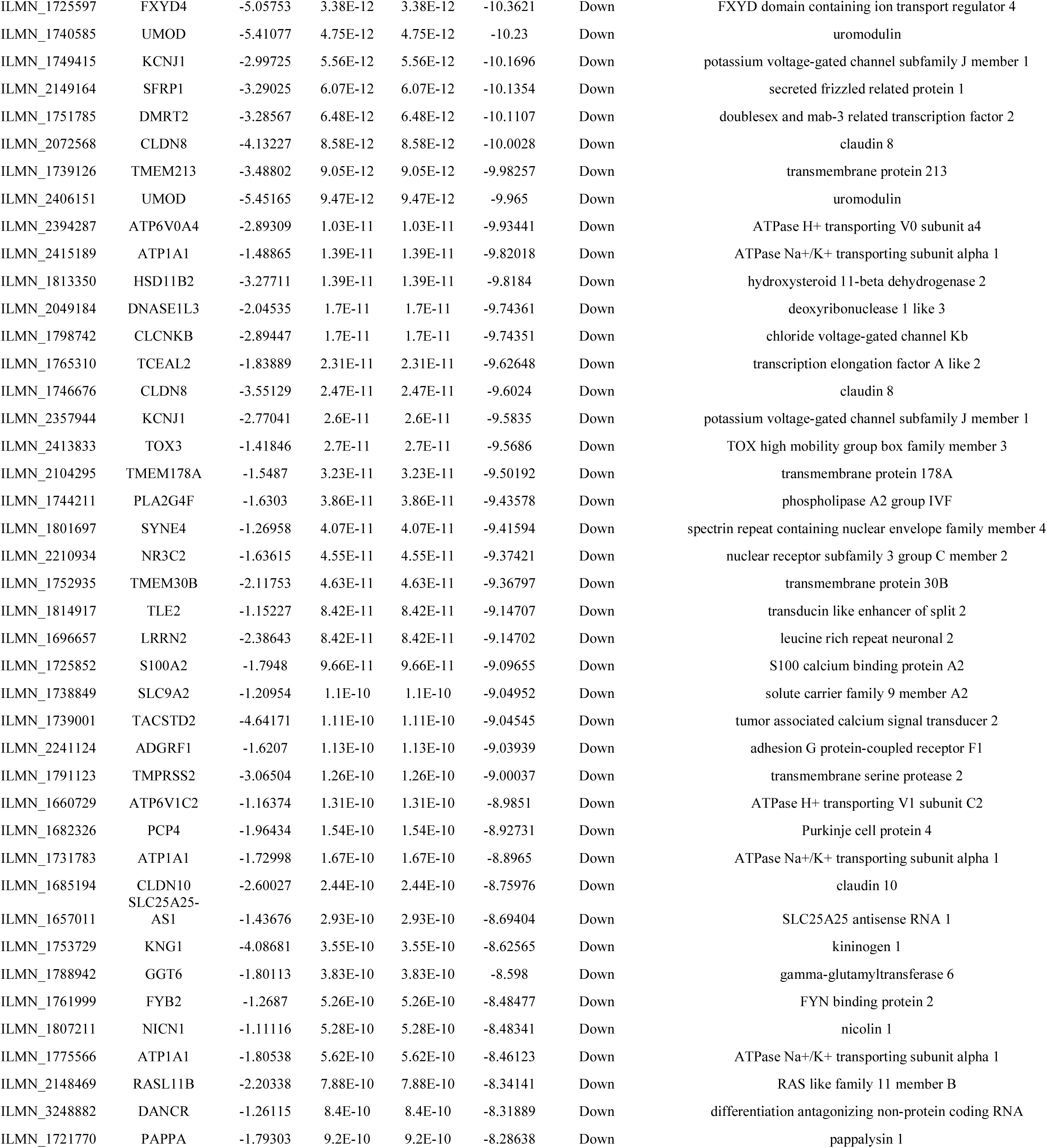
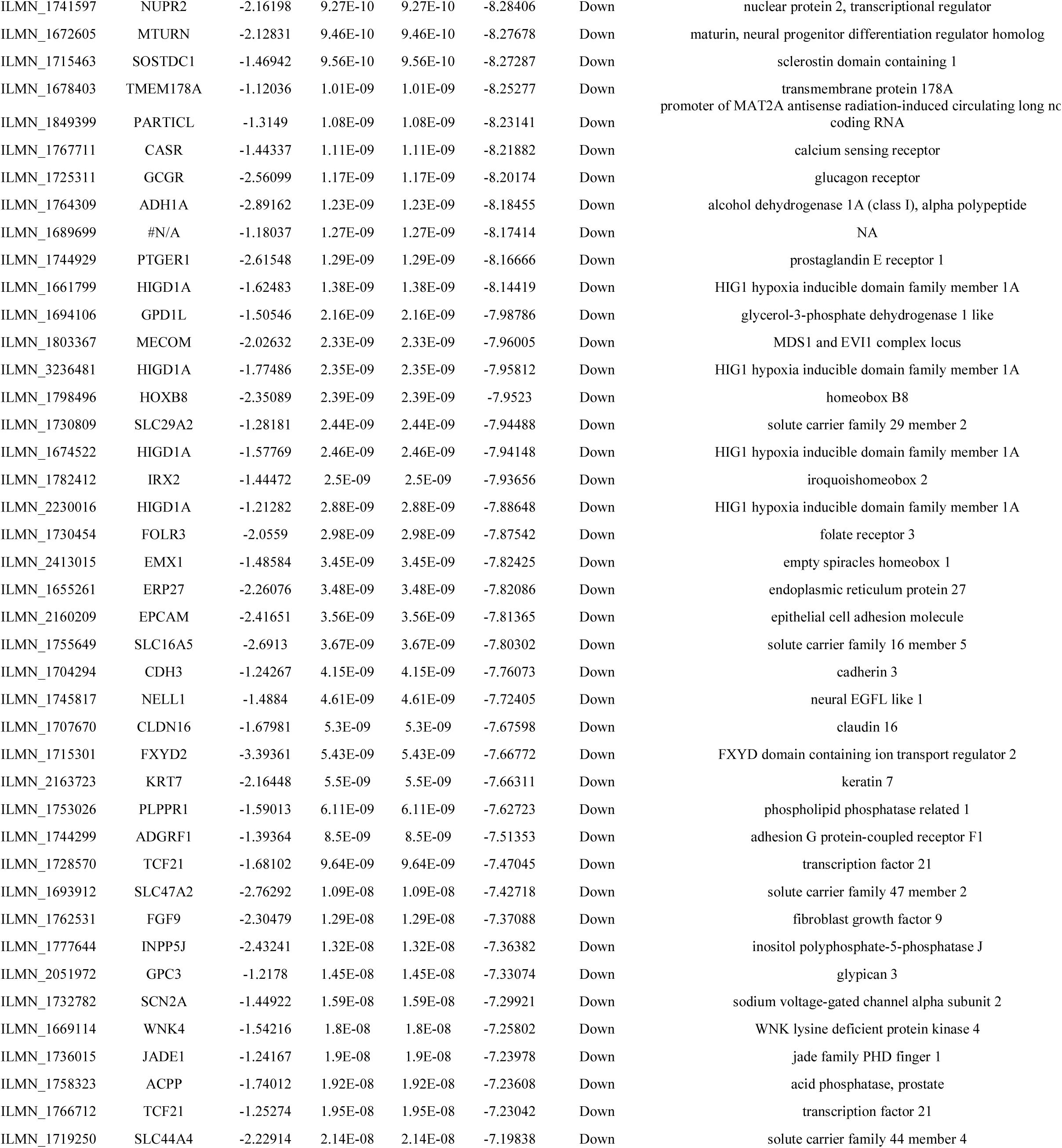
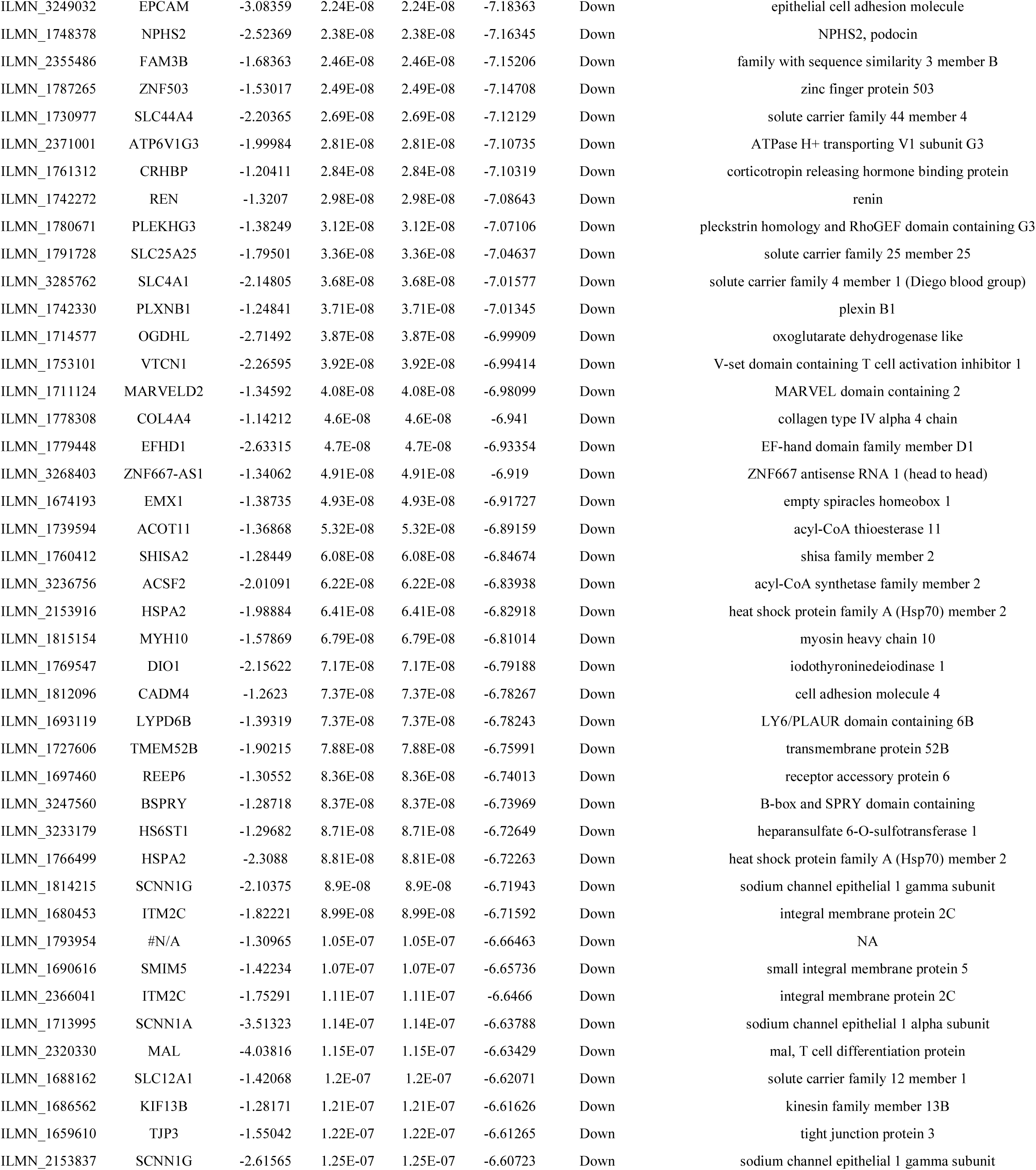
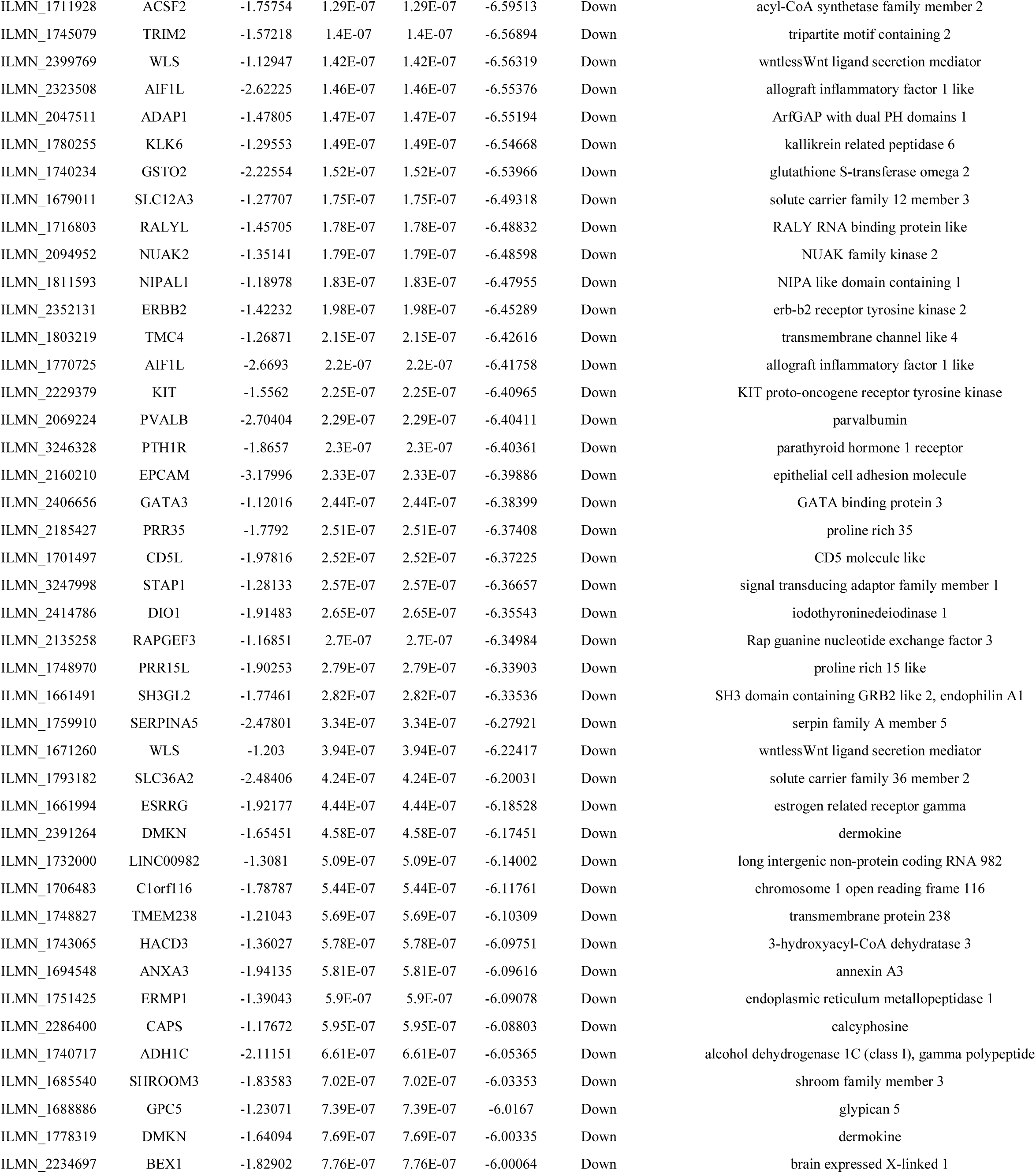
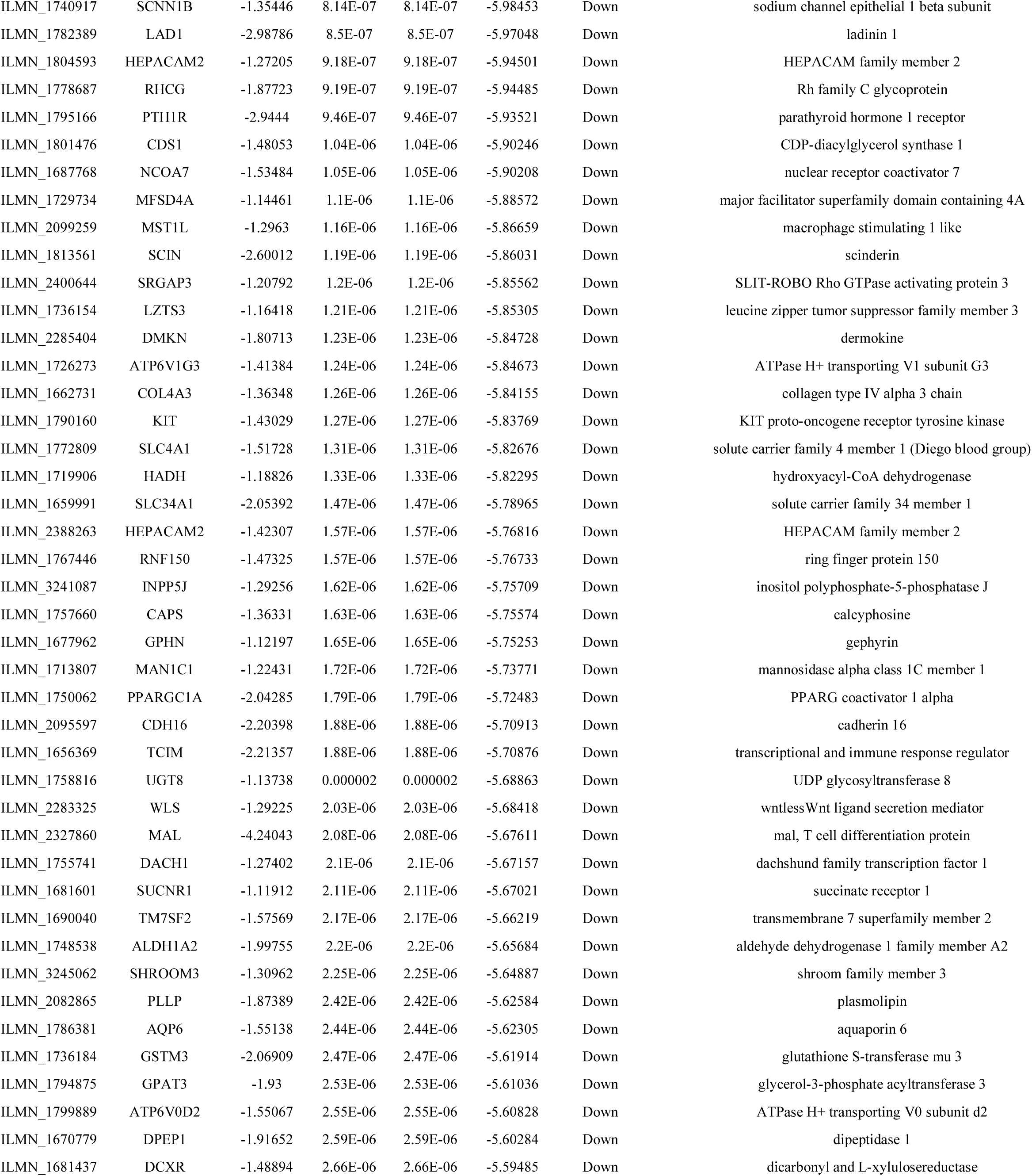
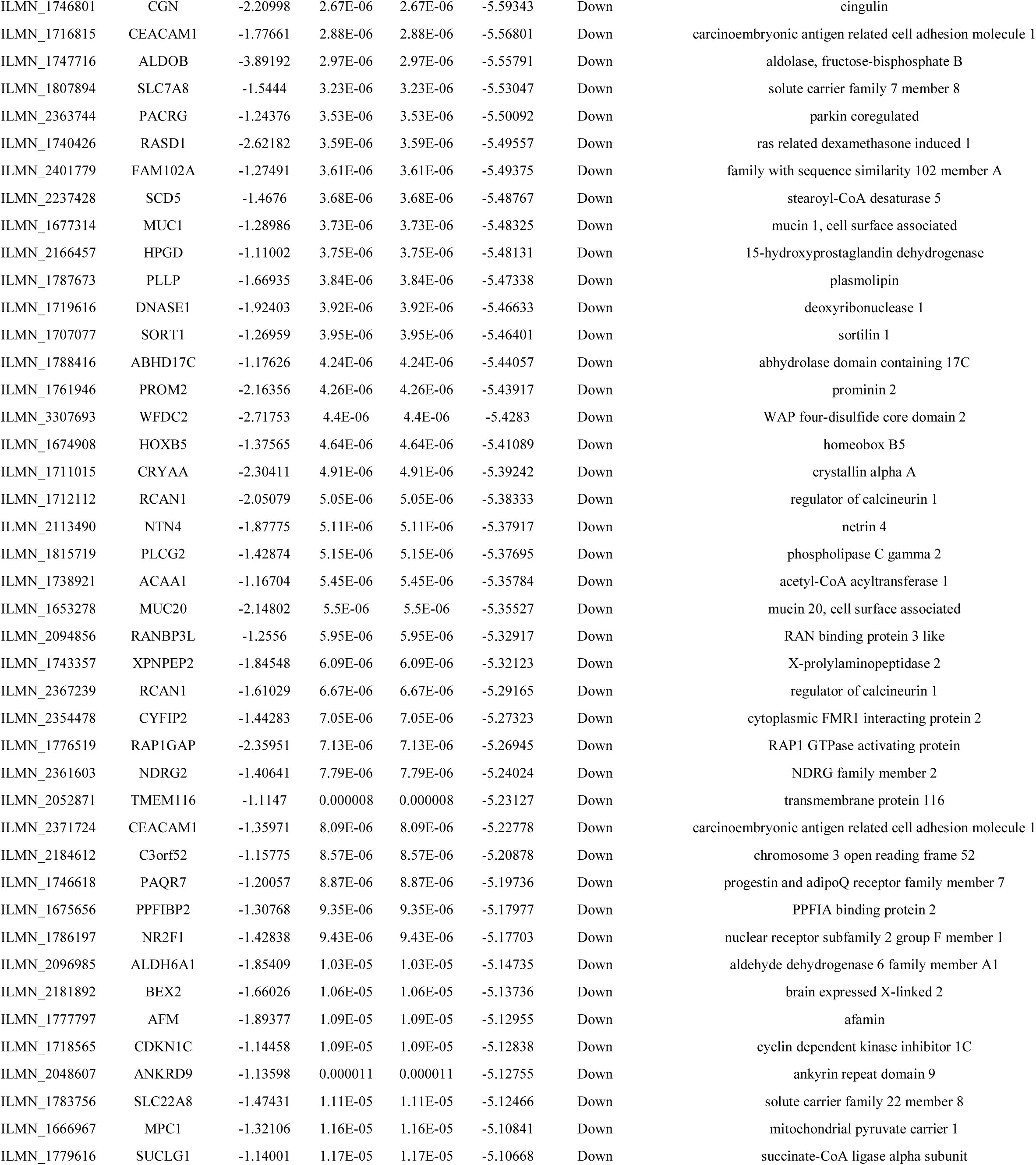
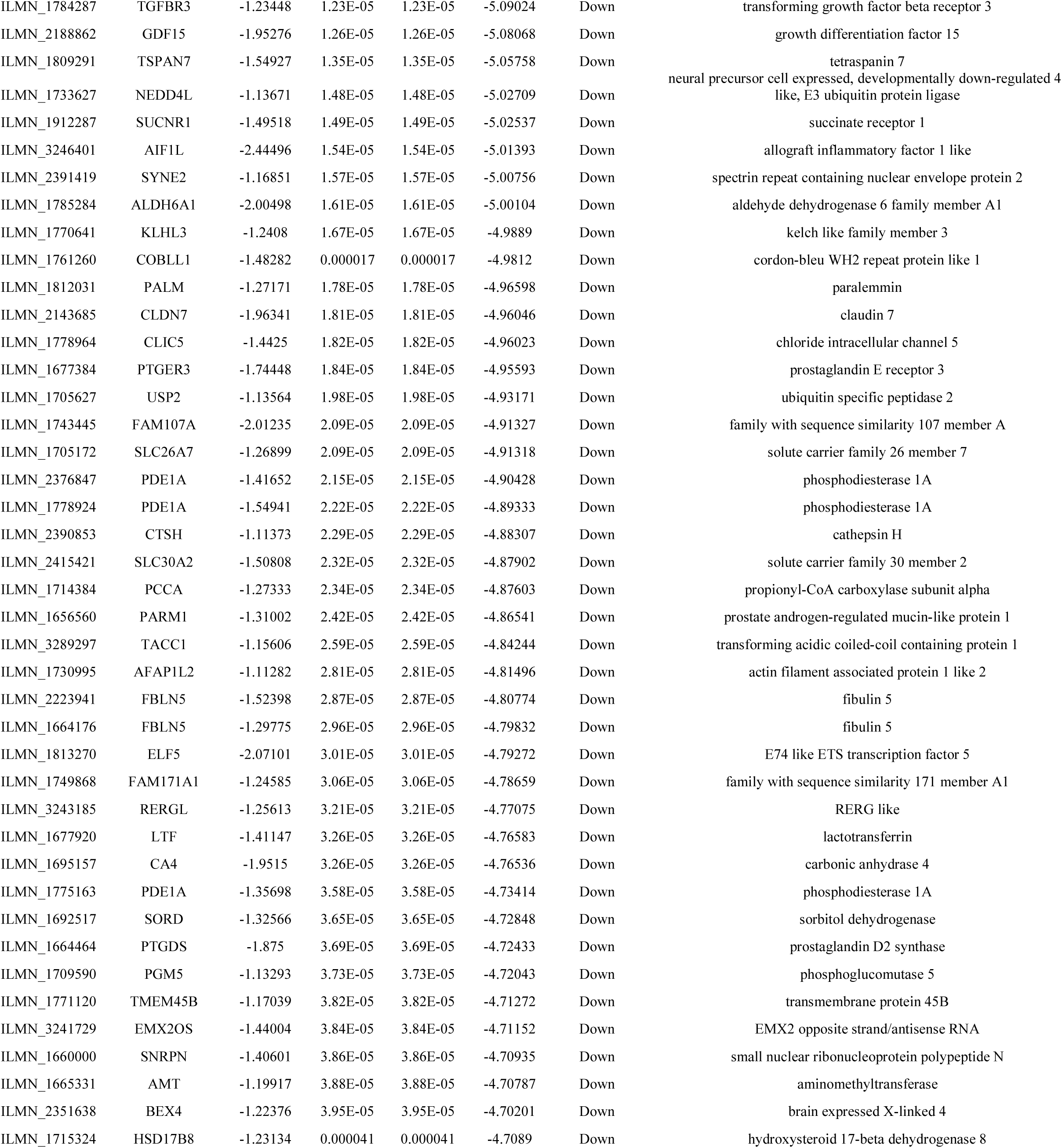
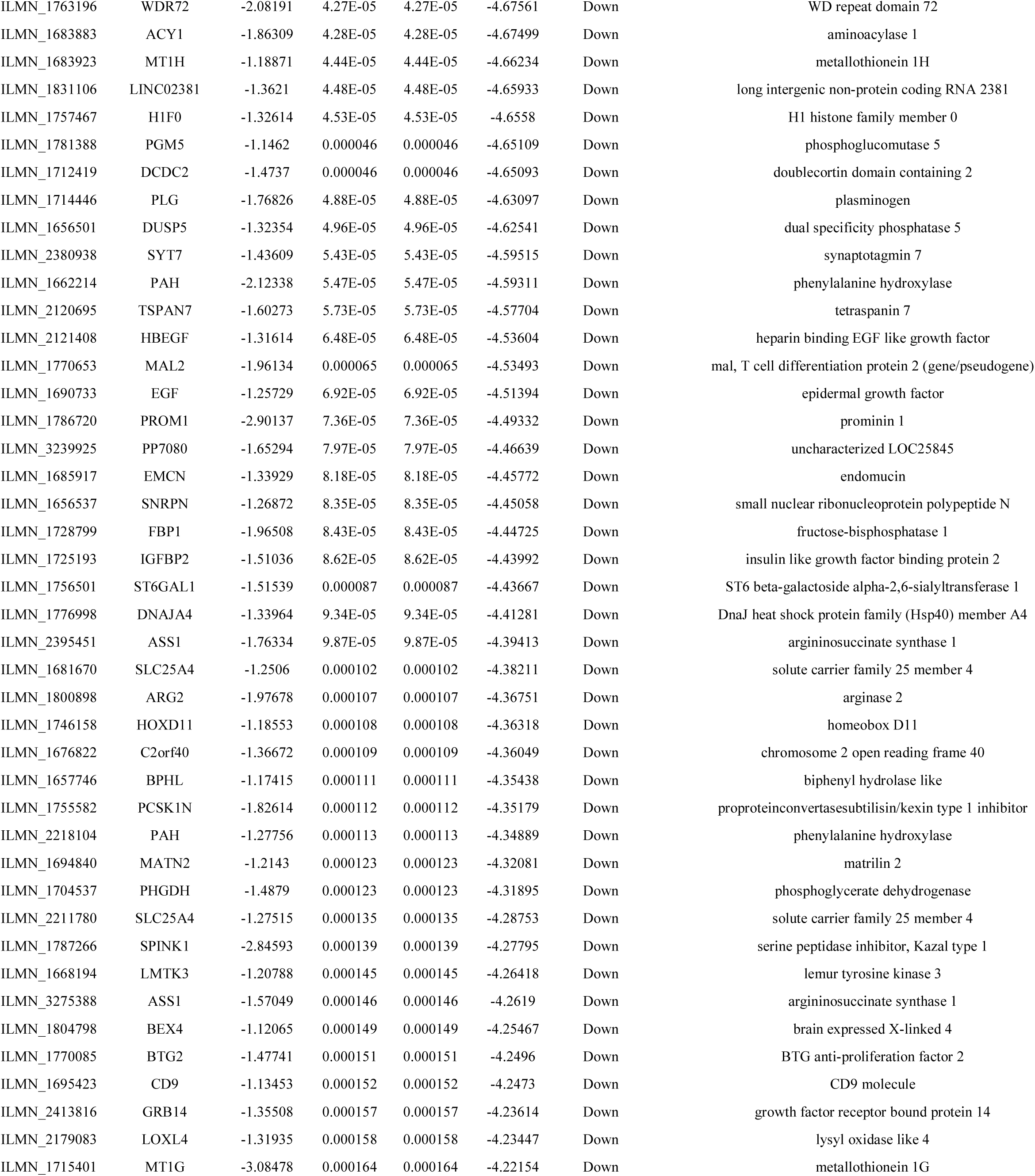
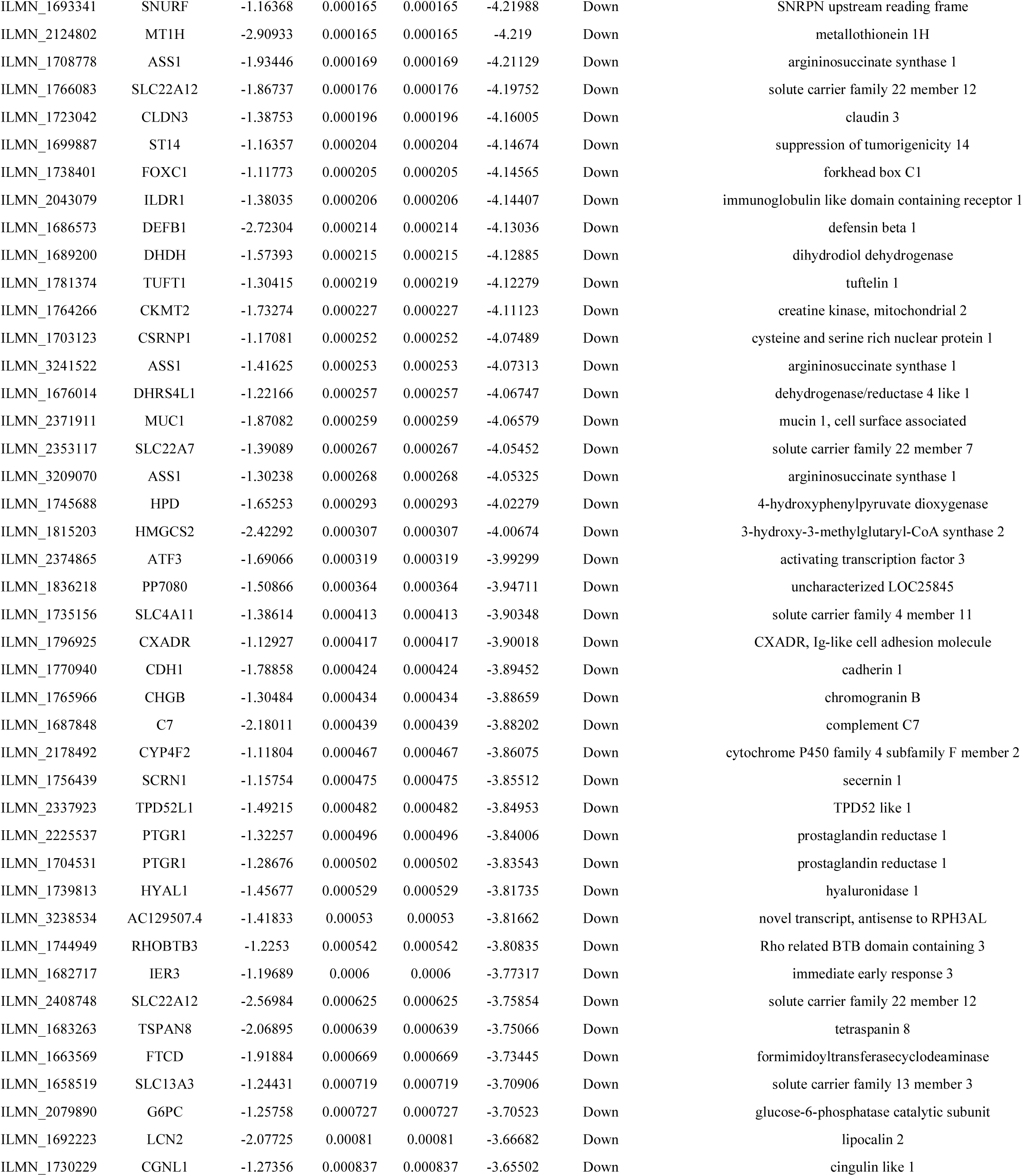
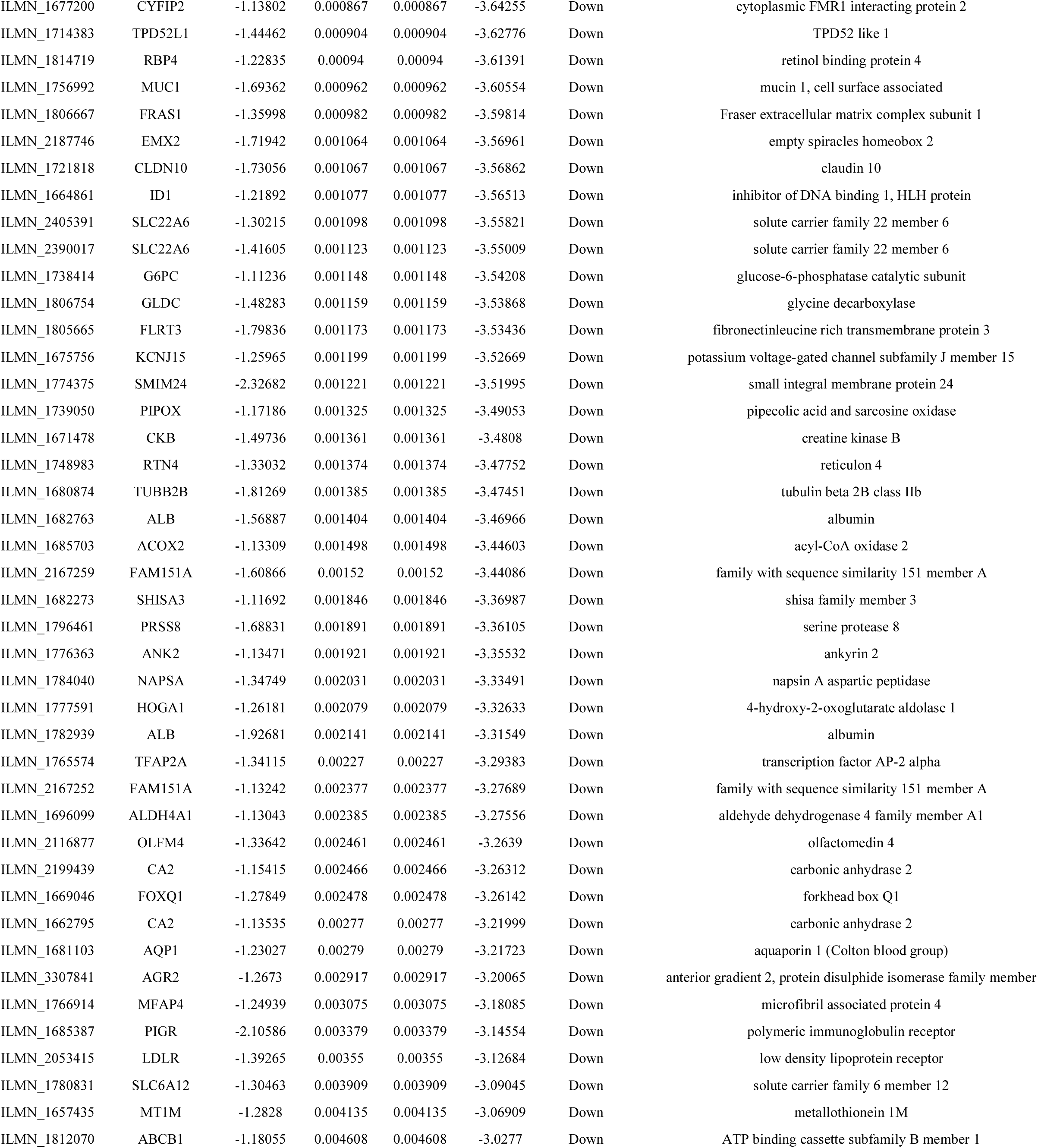
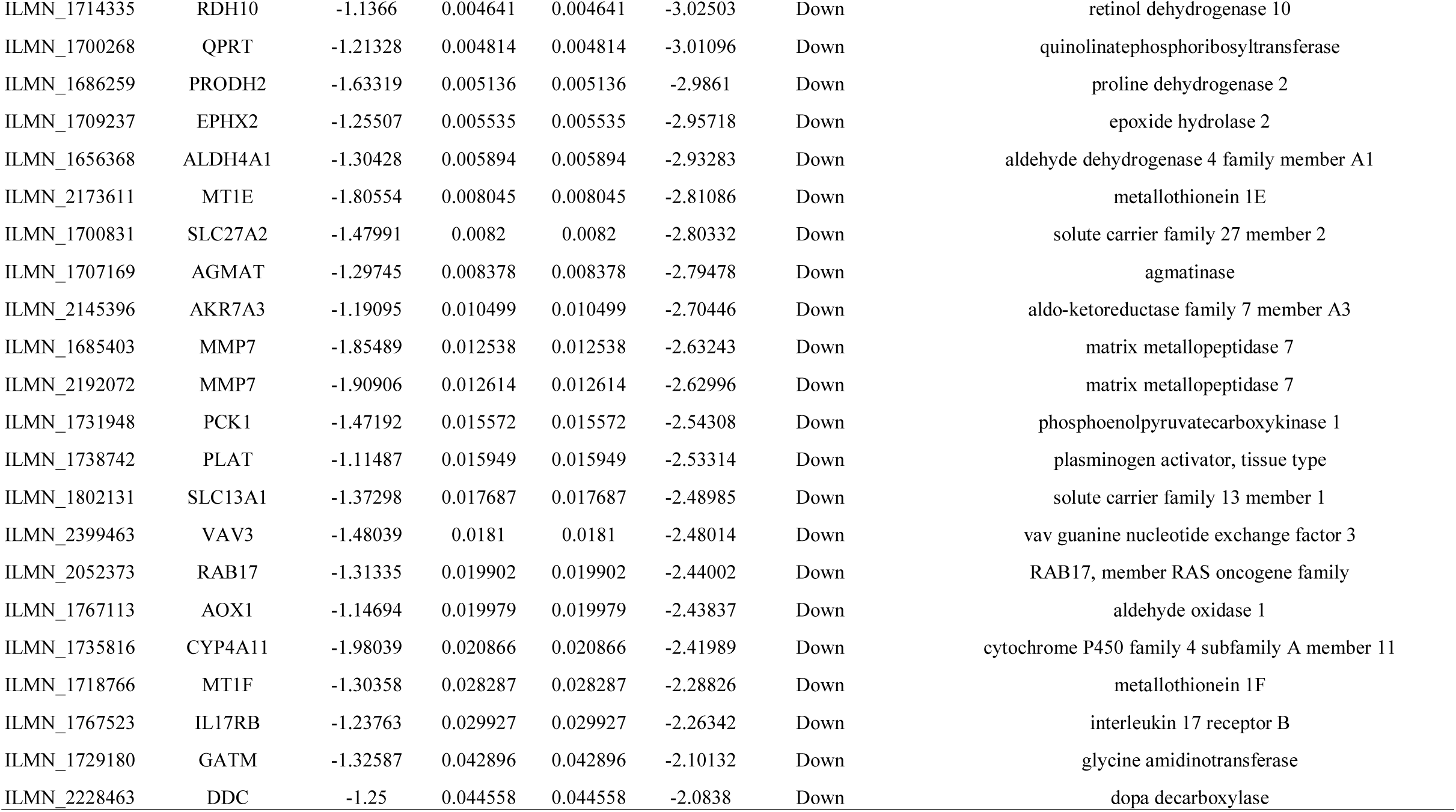
The statistical metrics for key differentially expressed genes (DEGs)

**Table 3.**
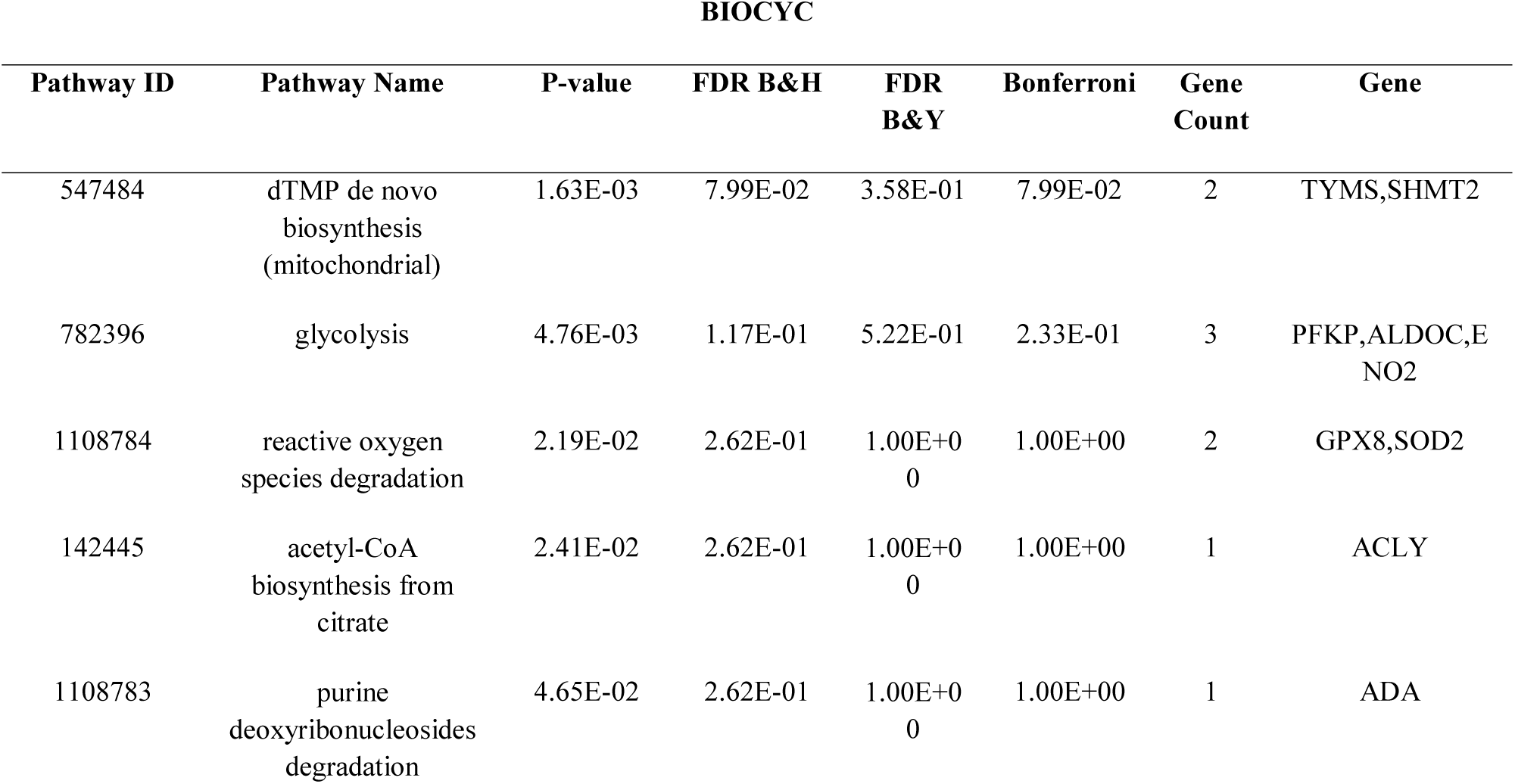
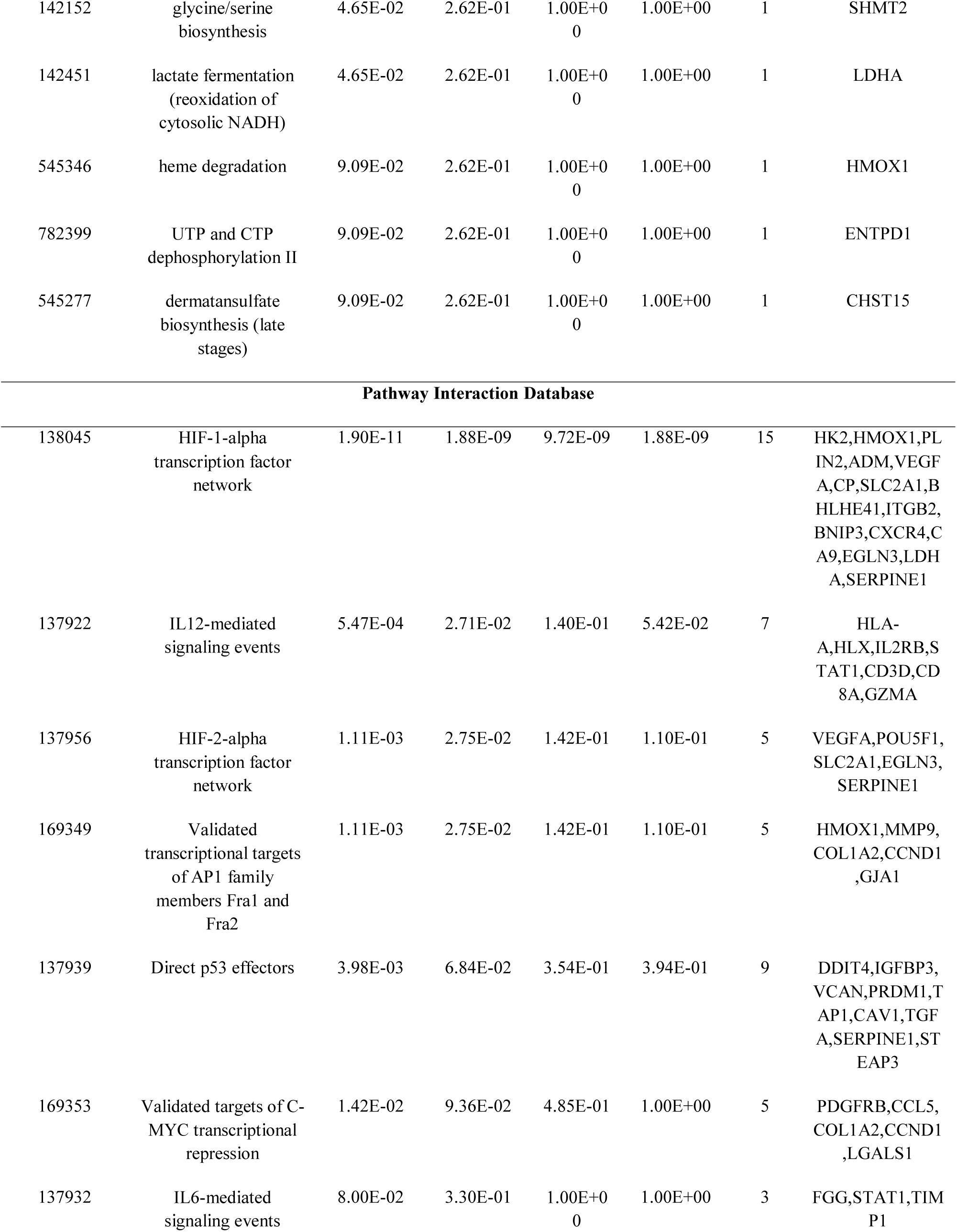
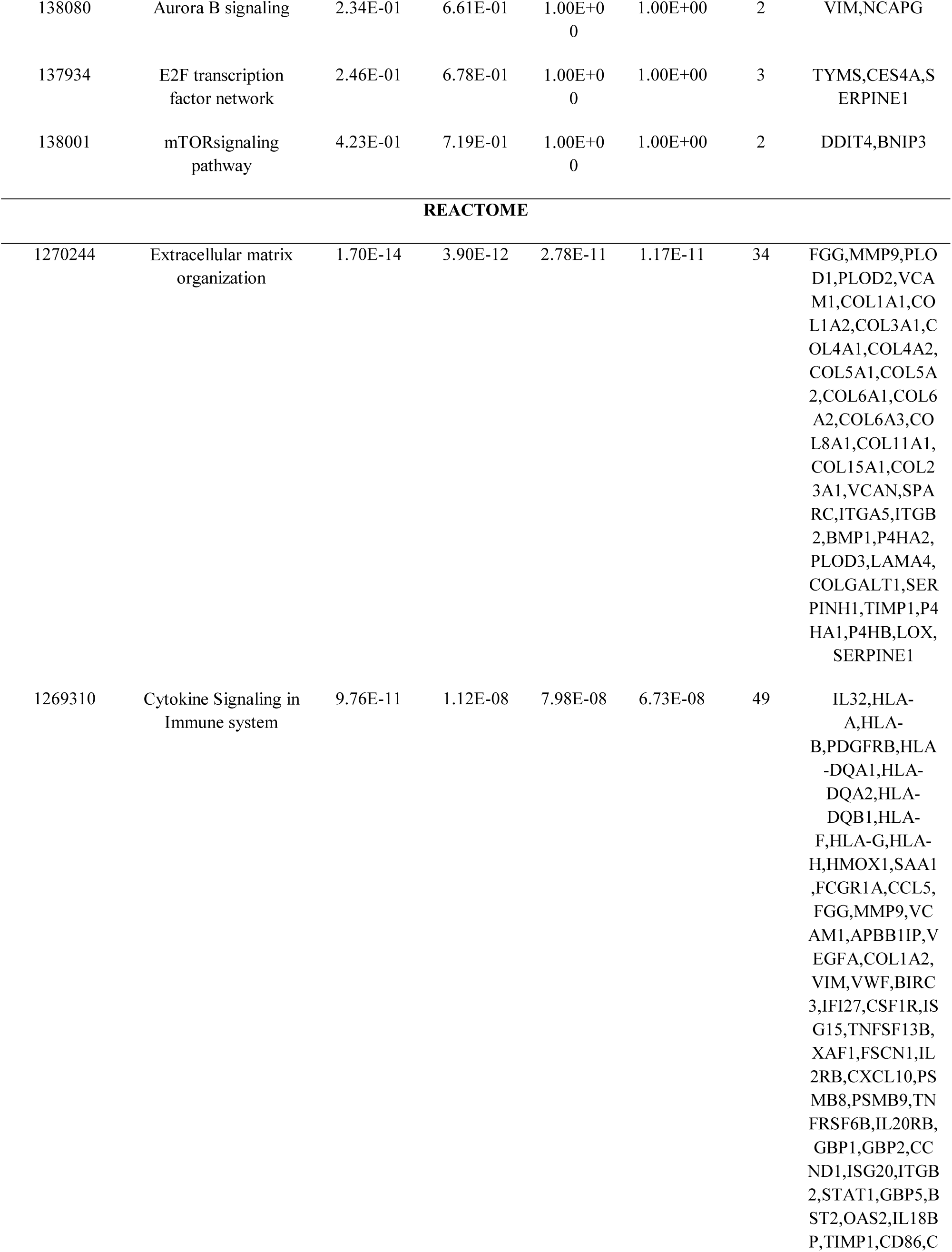
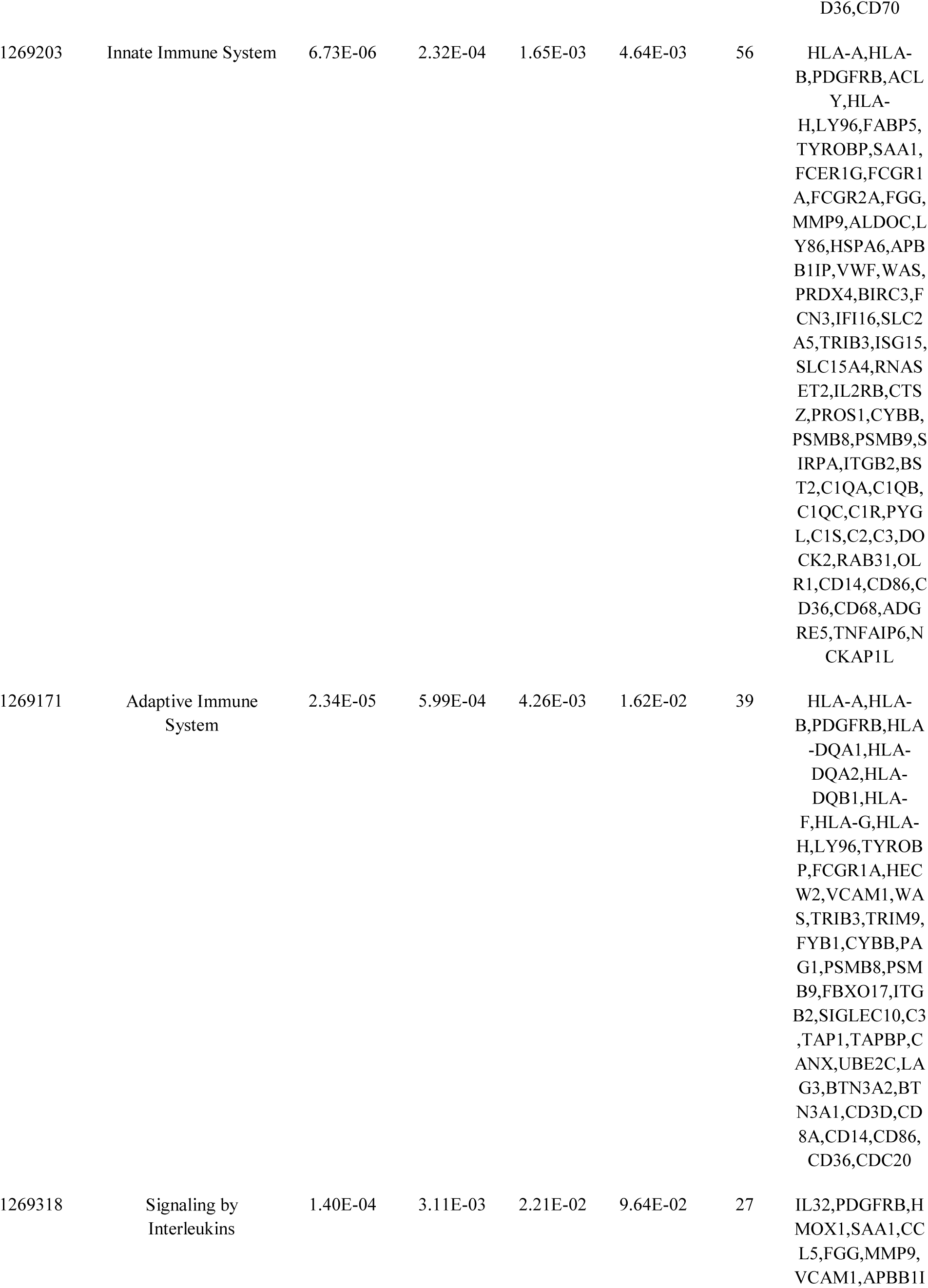
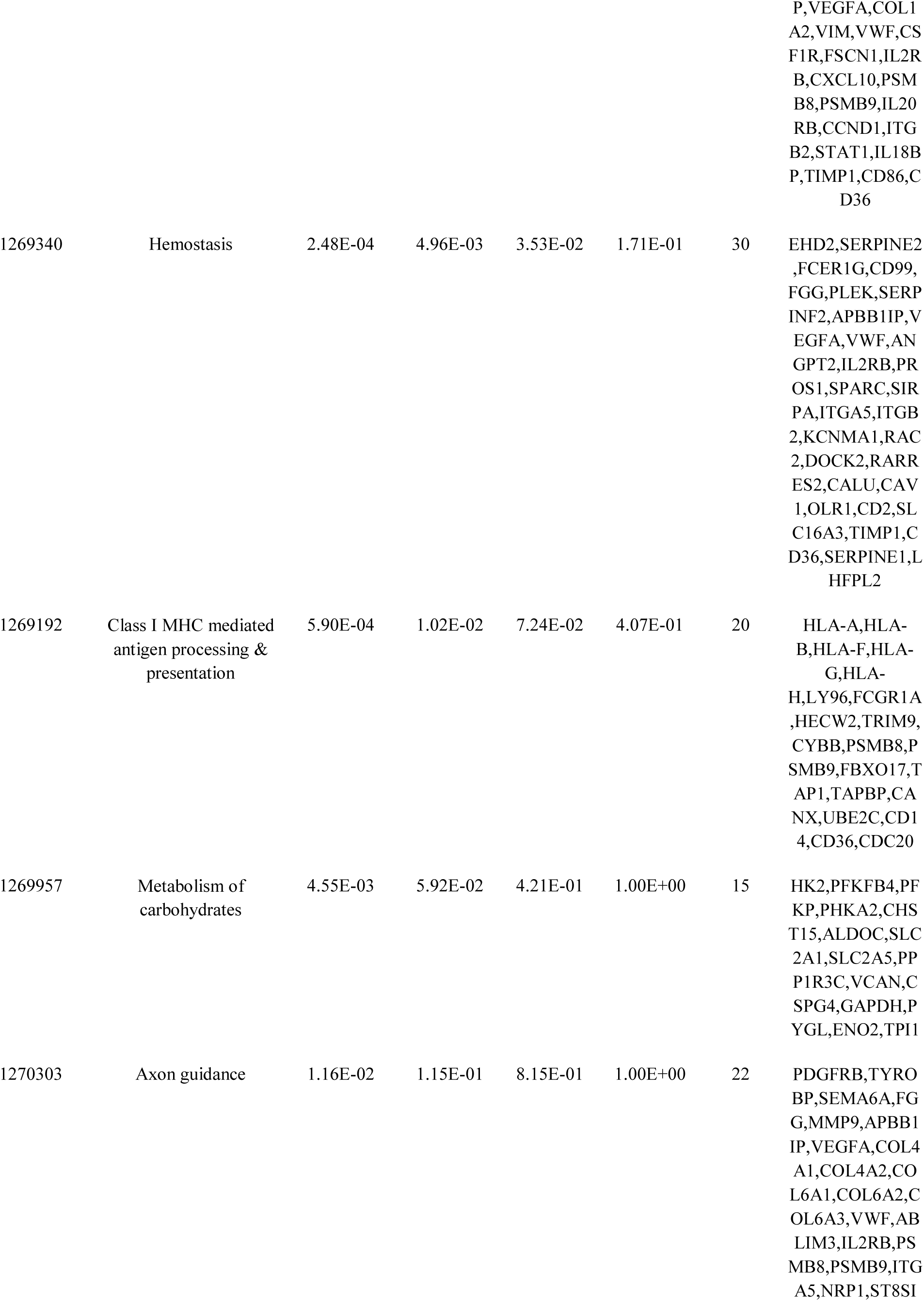
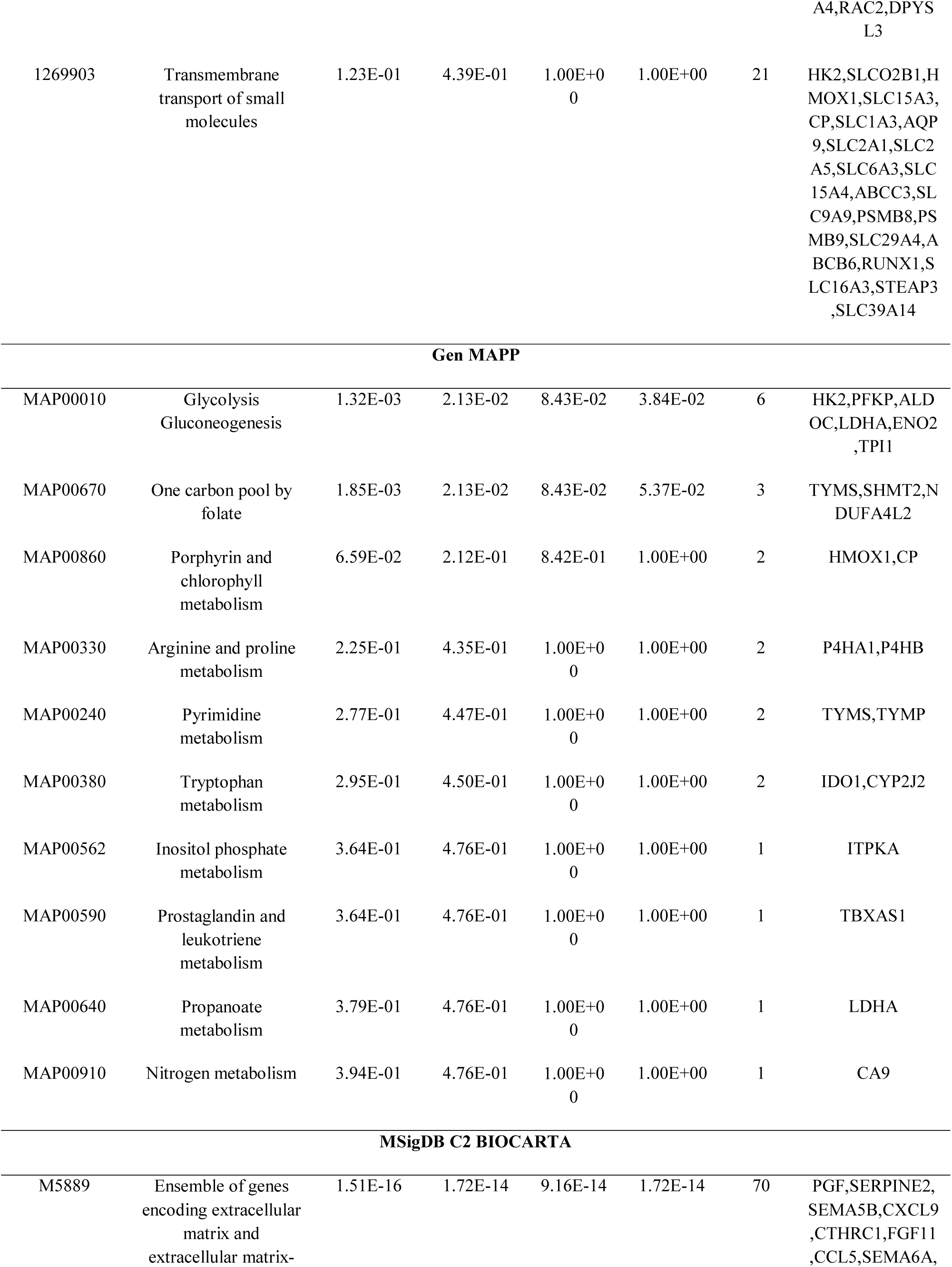
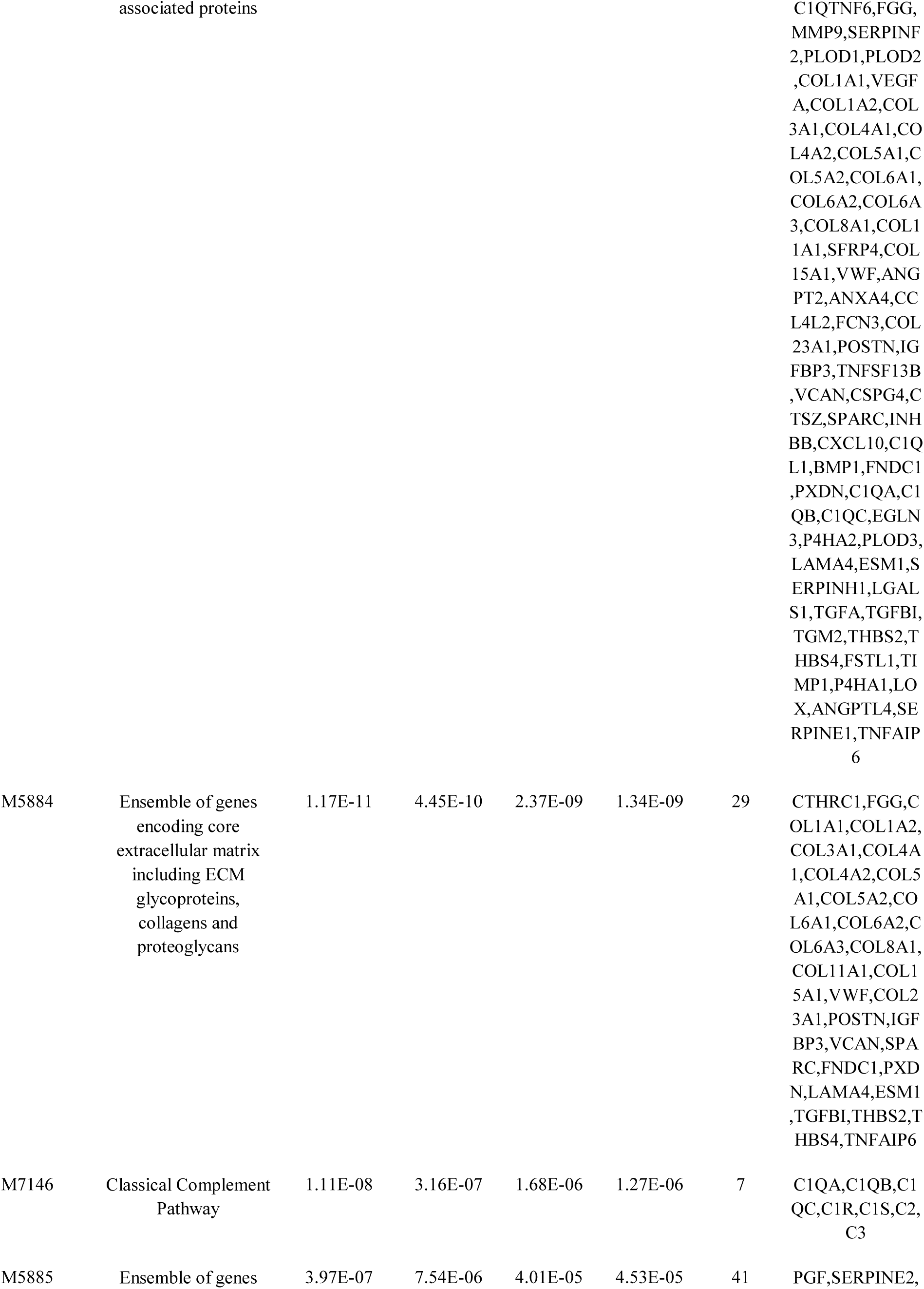
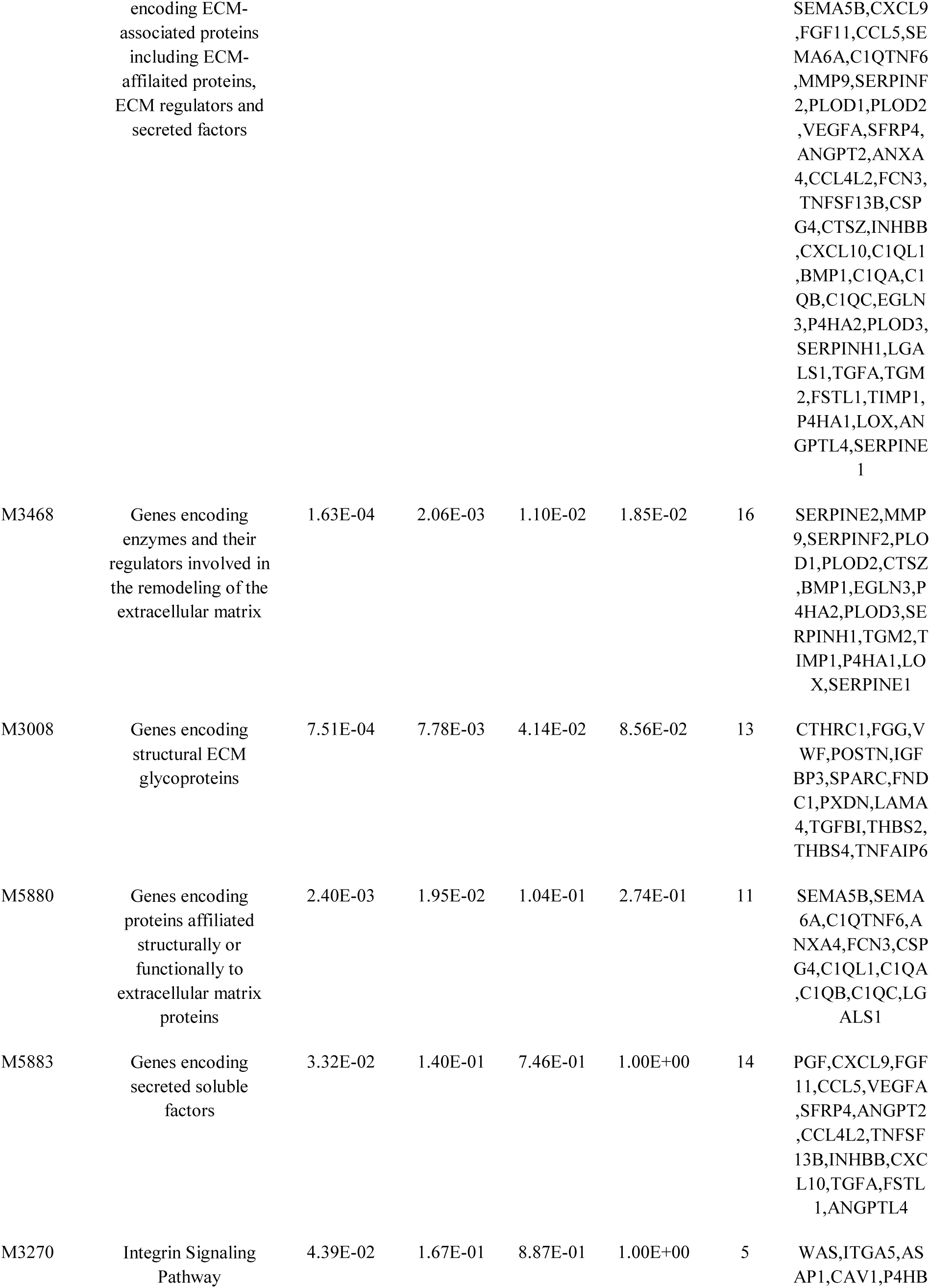
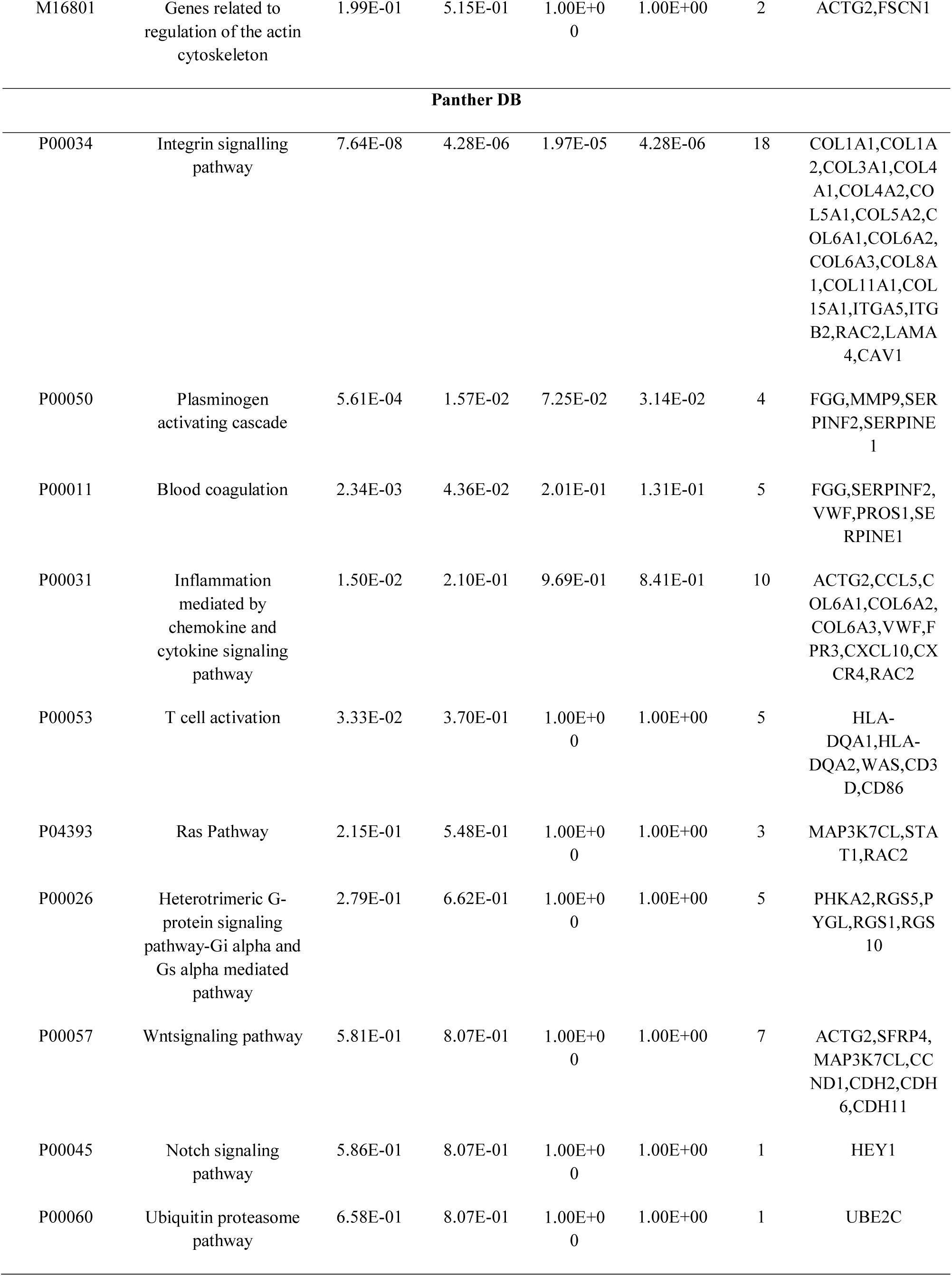
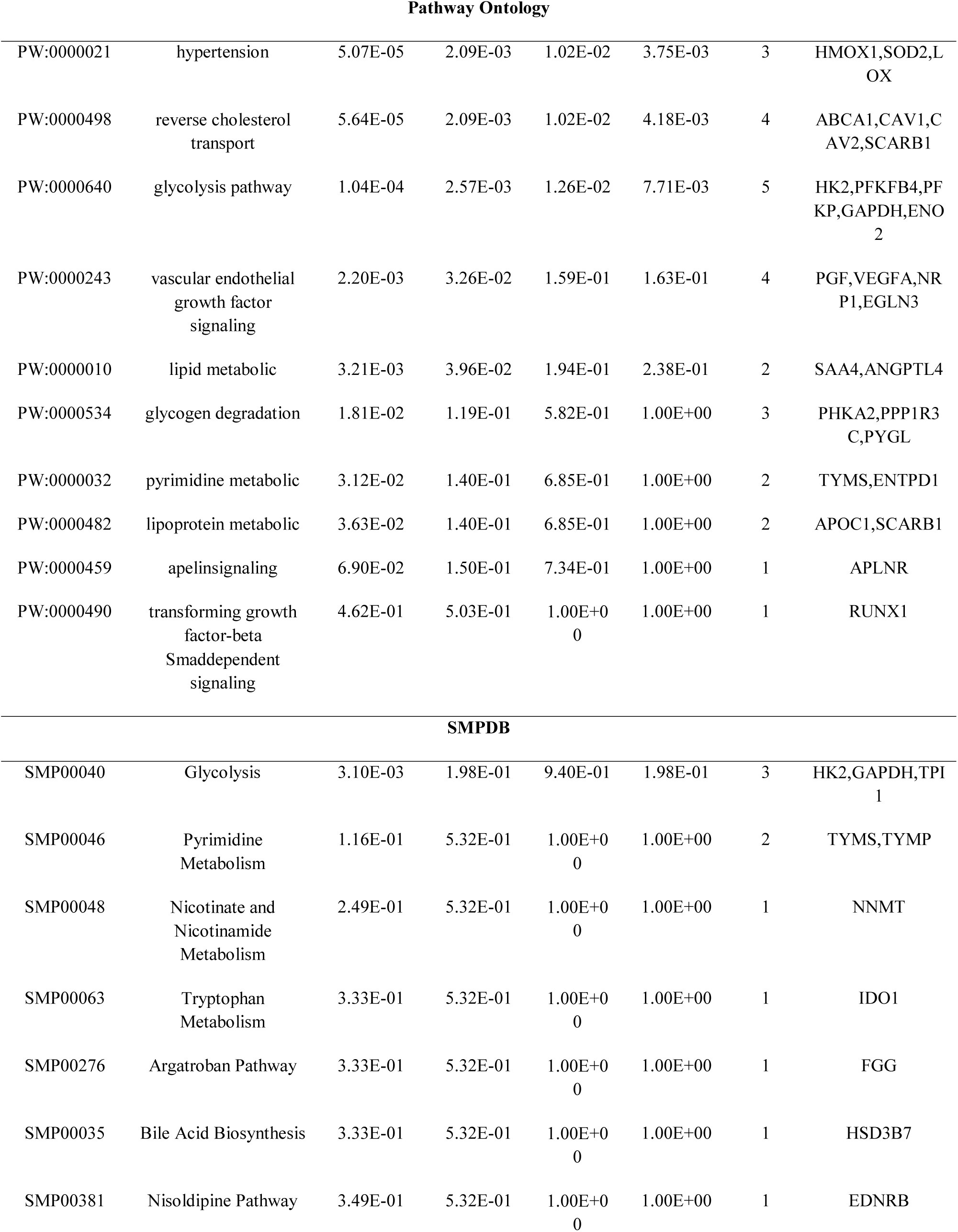
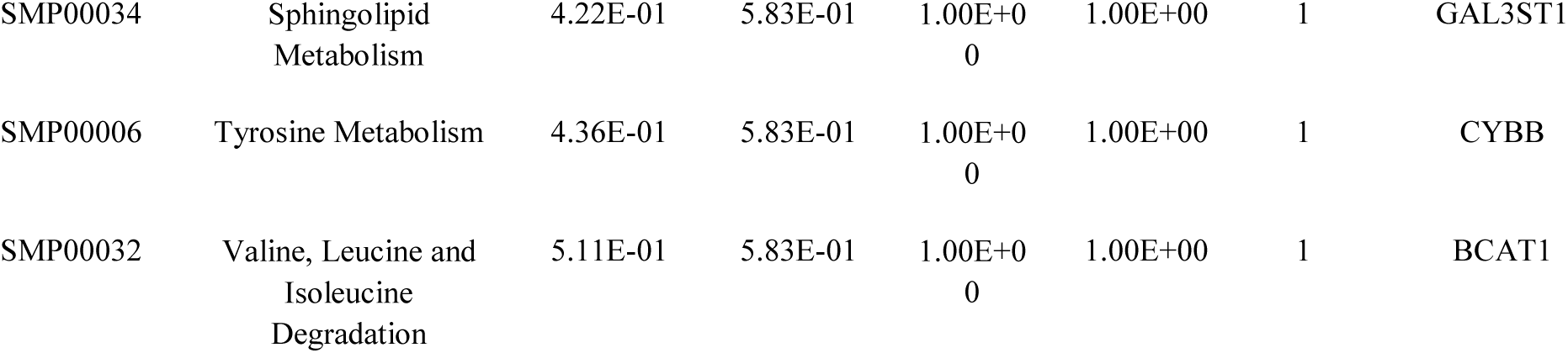
The enriched pathway terms of the up regulated differentially expressed genes

### GO enrichment analysis of DEGs

GO enrichment analysis for the up and down regulated DEGs were performed, respectively. Specifically, GO includes three items, including the BP, CC, and MF. Up regulated genes were mainly enriched in the BP, CC and MF including response to cytokine, defense response, vesicle organization, cell surface, signaling receptor binding and extracellular matrix structural constituent are listed in Table 4. Similarly, down regulated genes were mainly enriched in the BP, CC and MF including renal system development, organic acid metabolic process, apical part of cell, intrinsic component of plasma membrane, ion transmembrane transporter activity and transmembrane transporter activity are listed in Table 5.

**Table 4.**
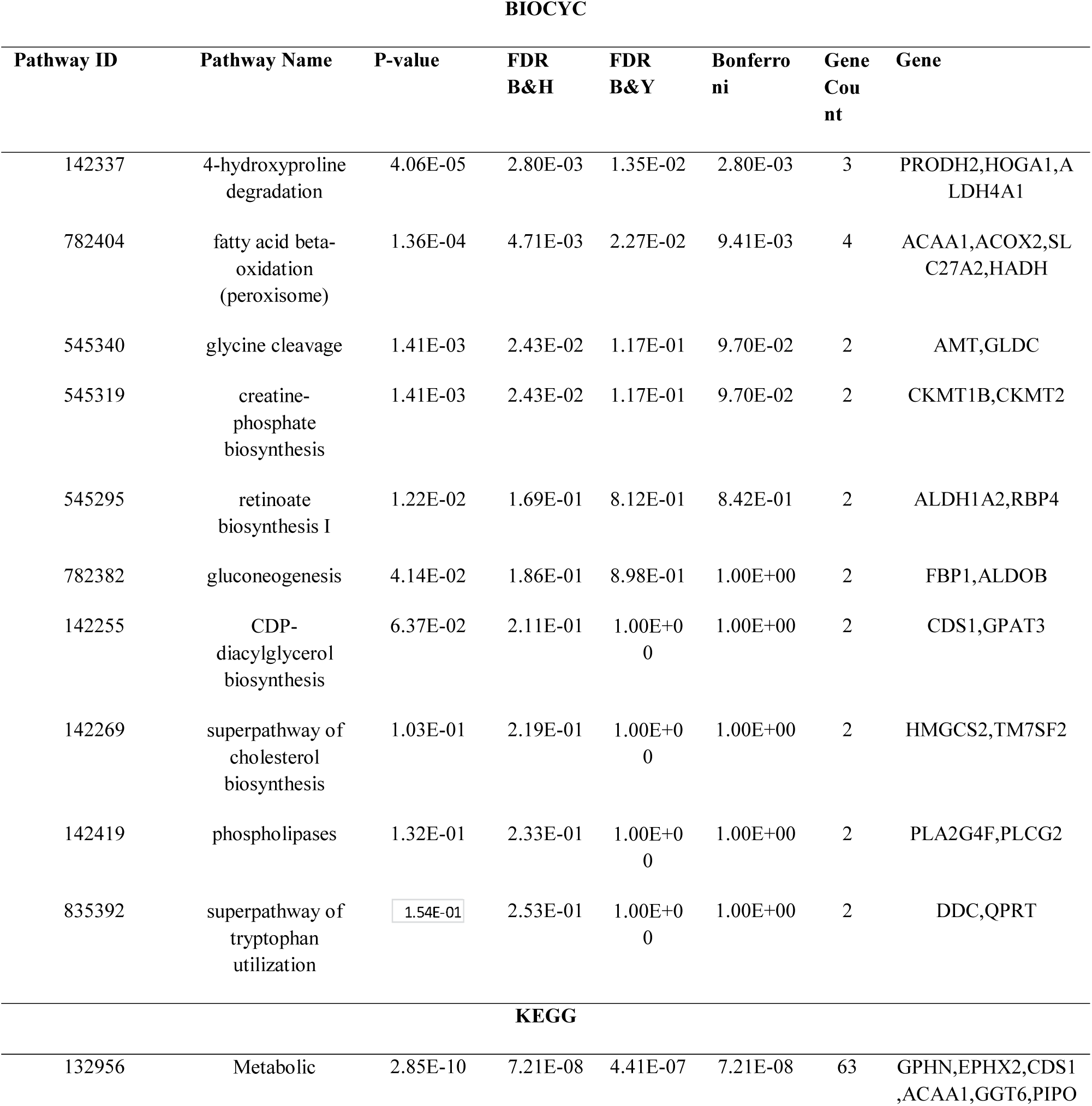
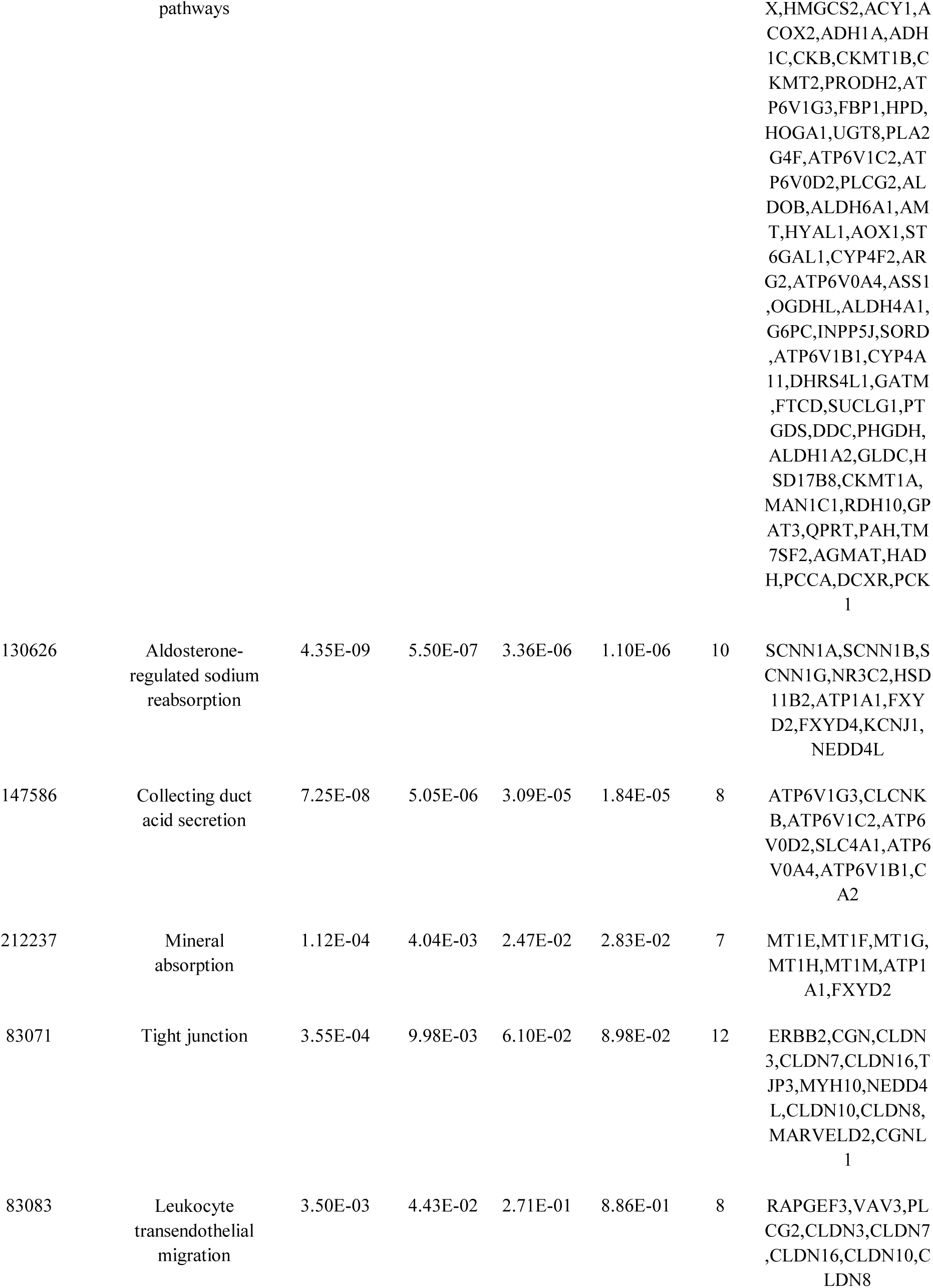
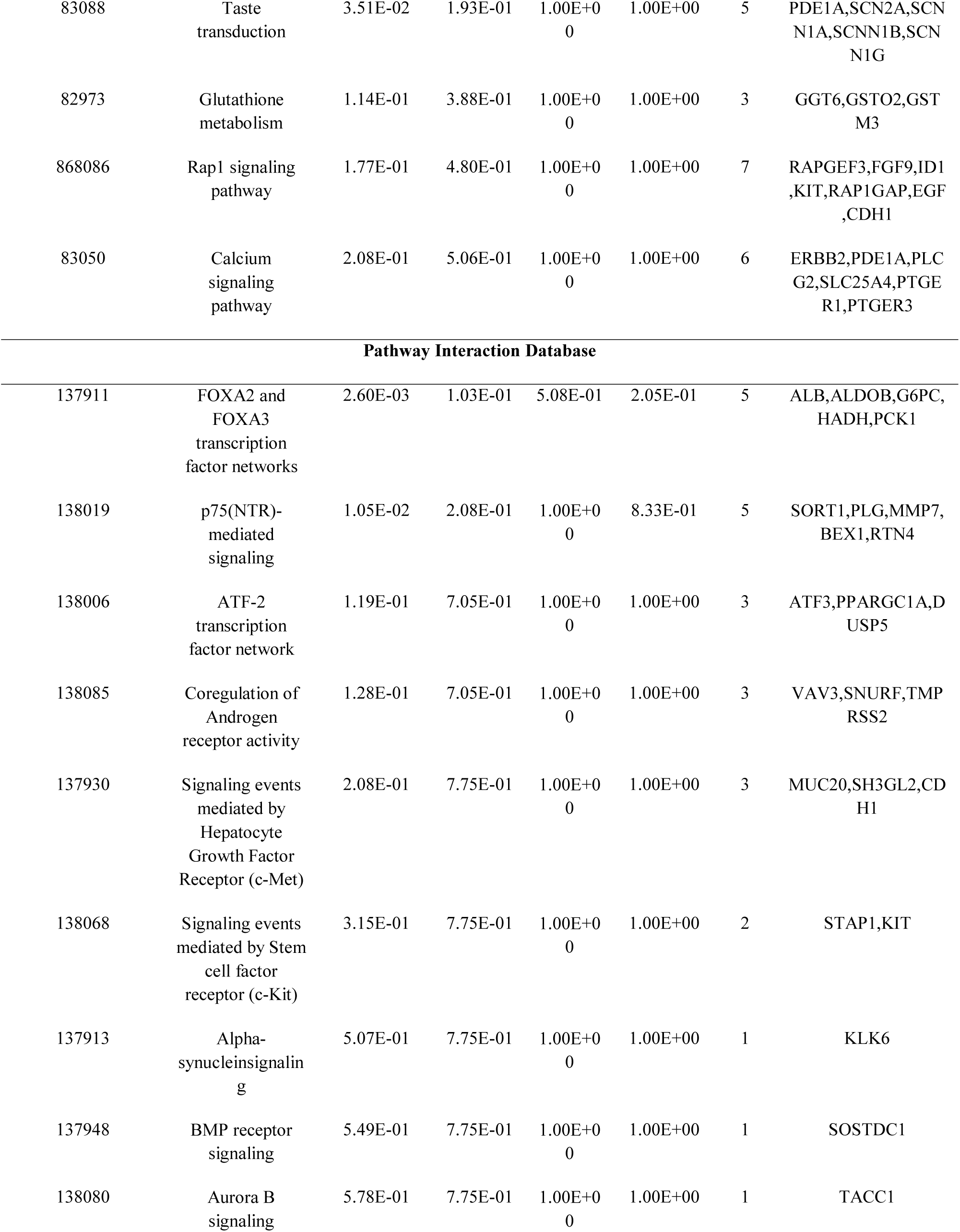
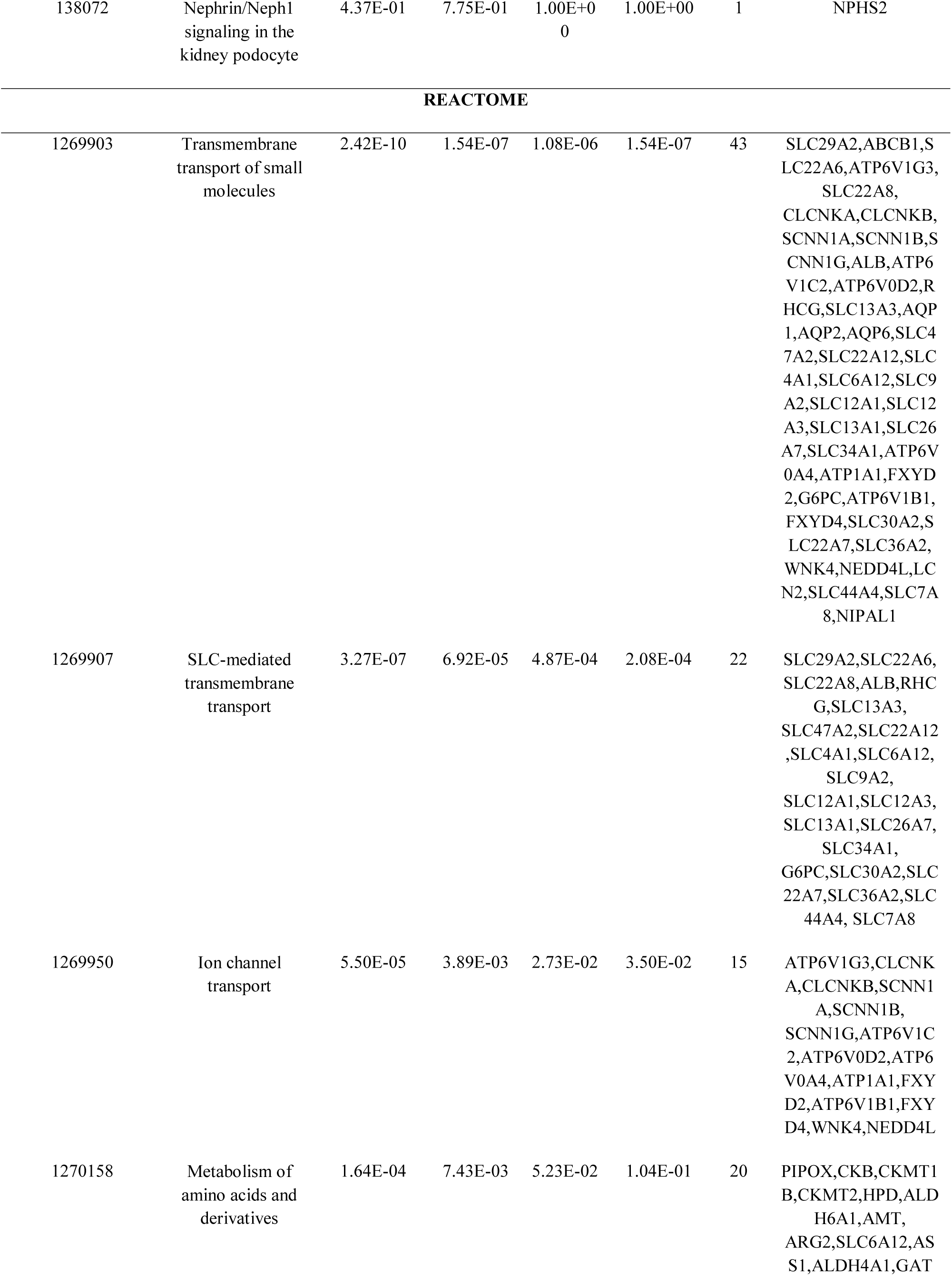
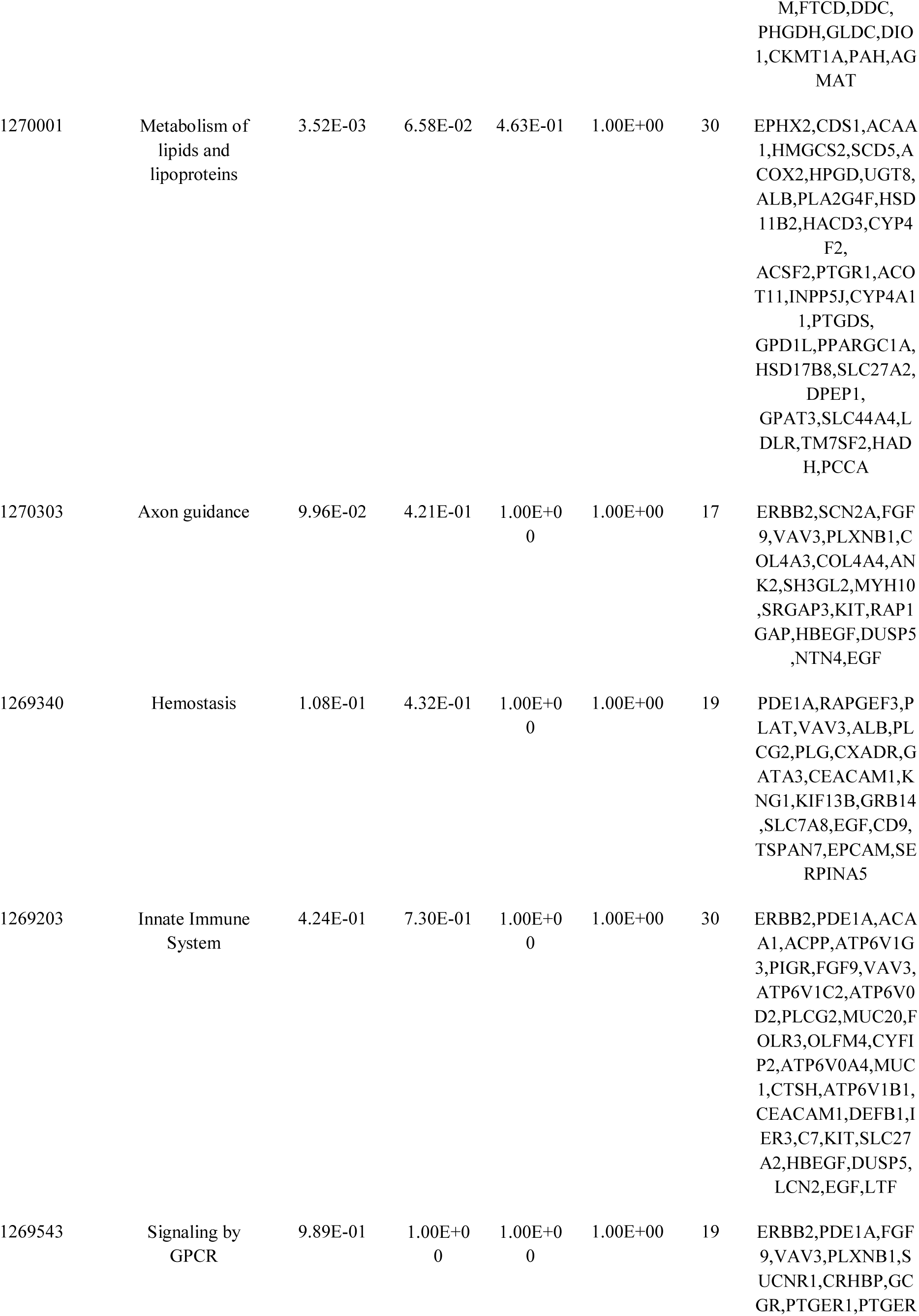
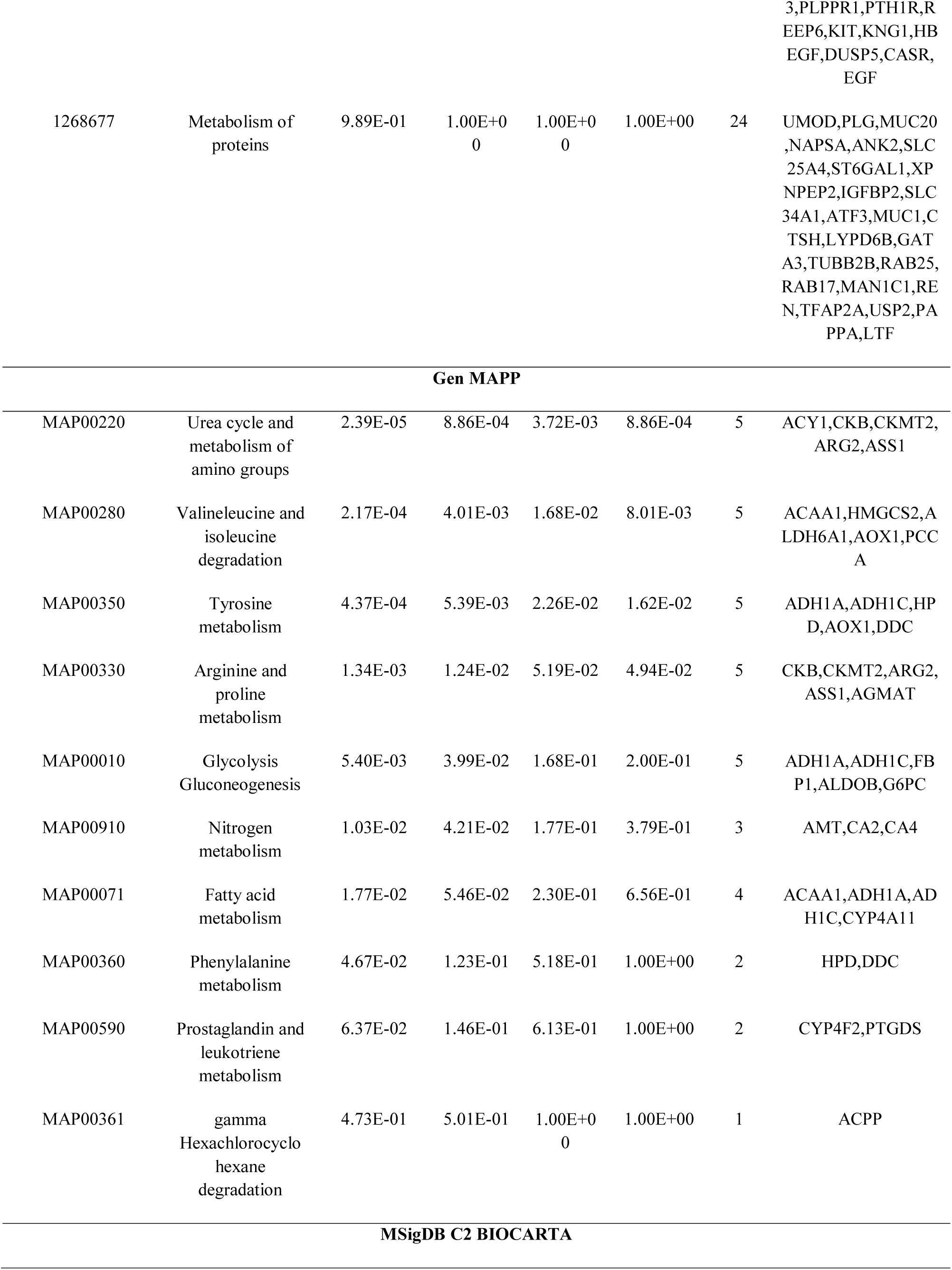
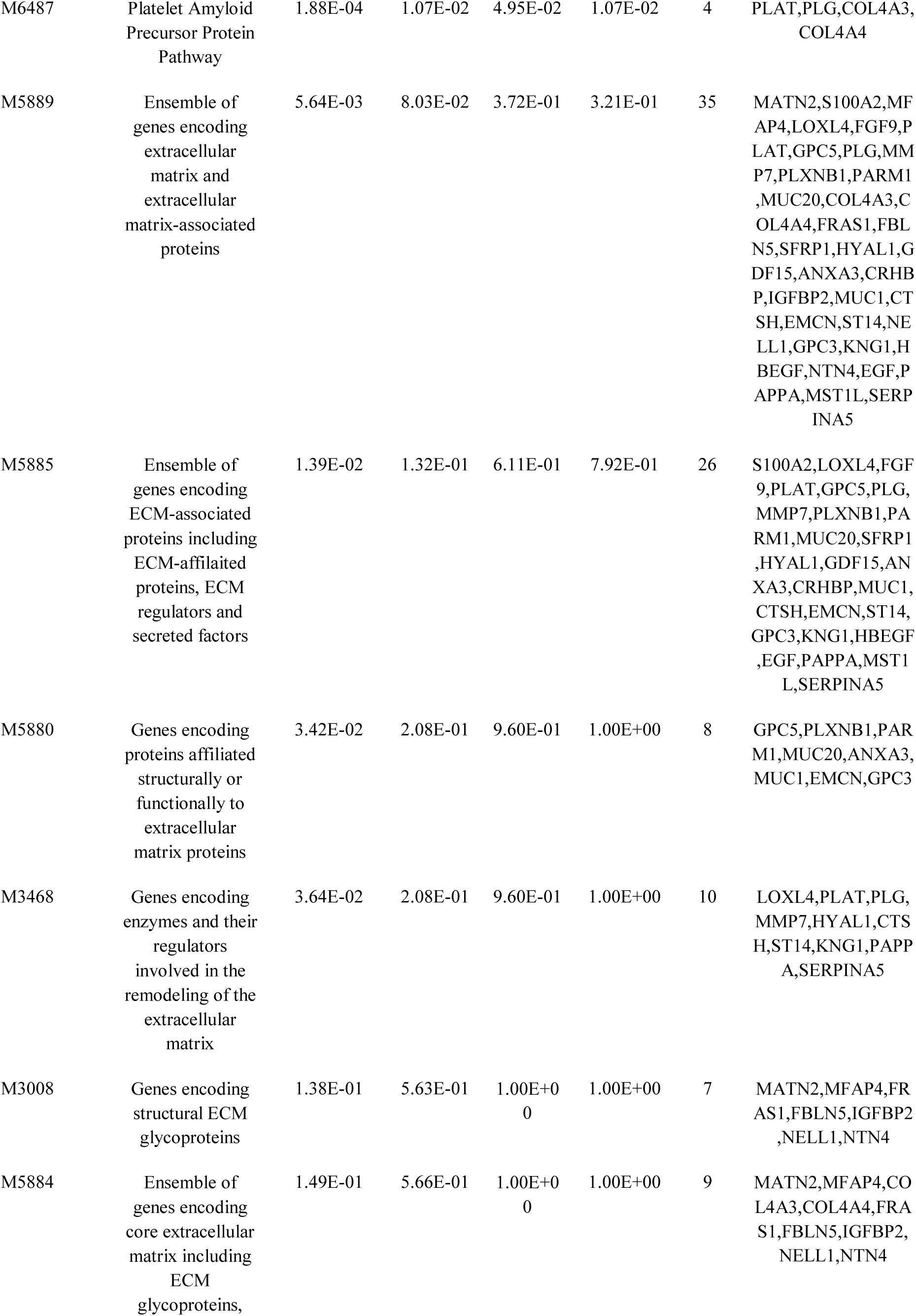
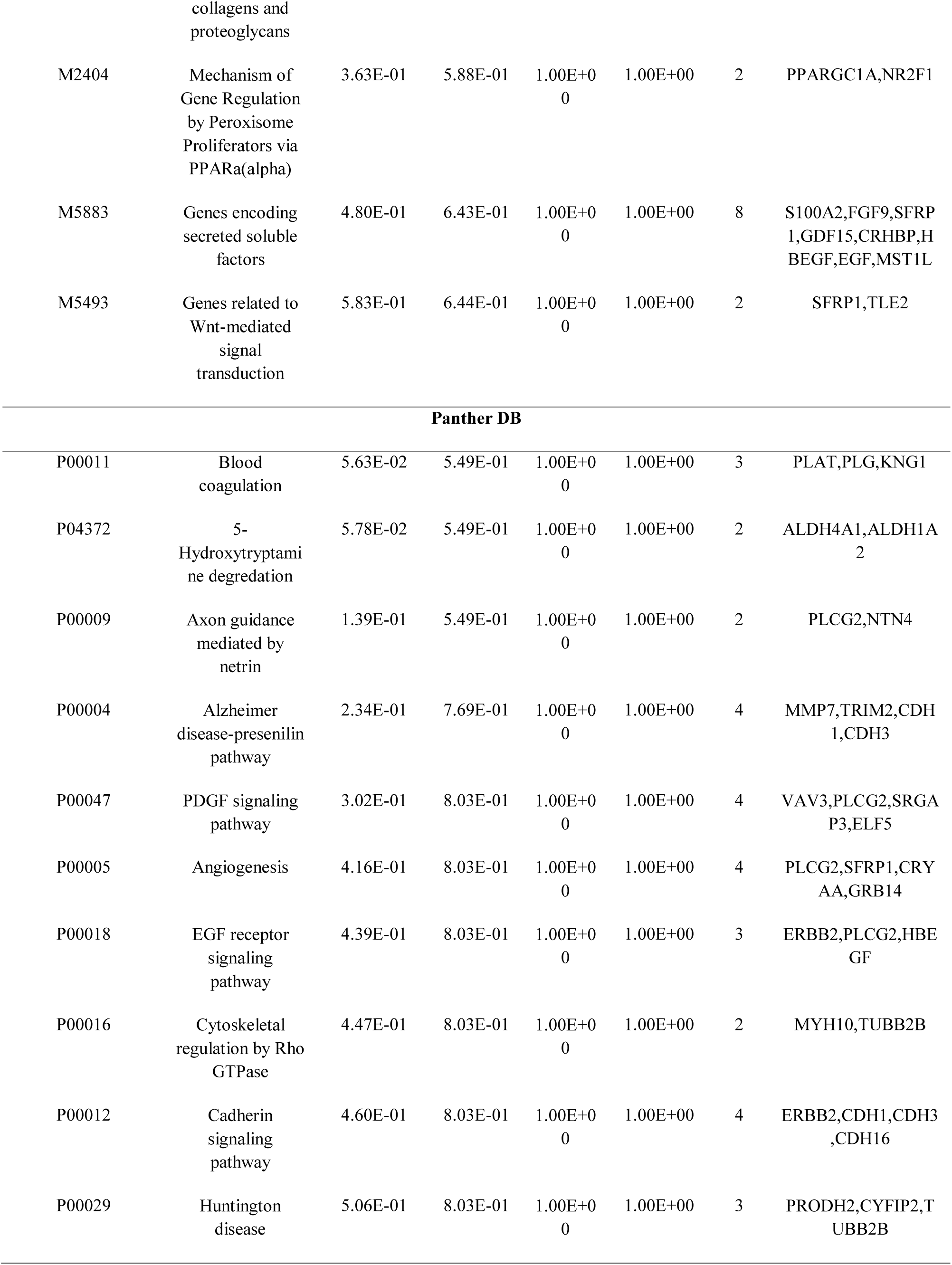
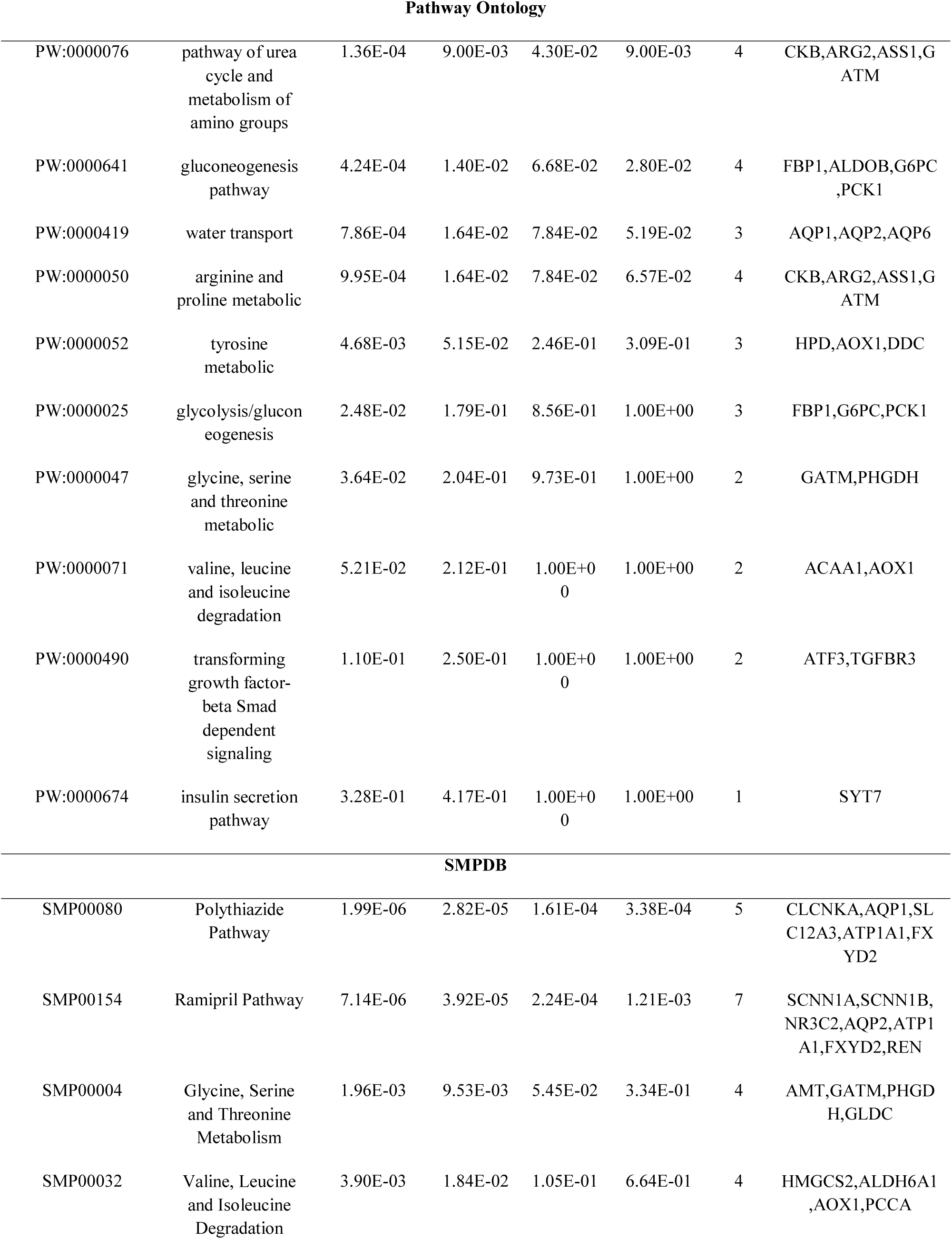
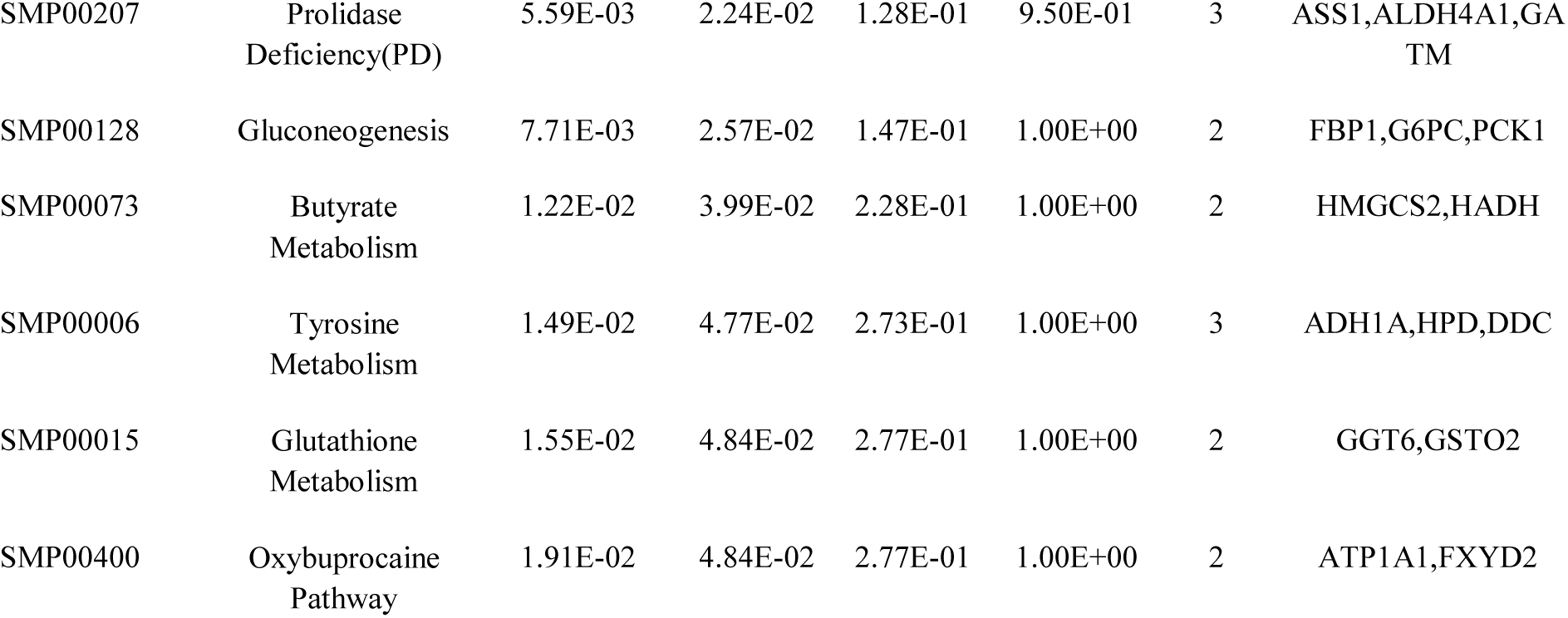
The enriched pathway terms of the down regulated differentially expressed genes

**Table 5.**
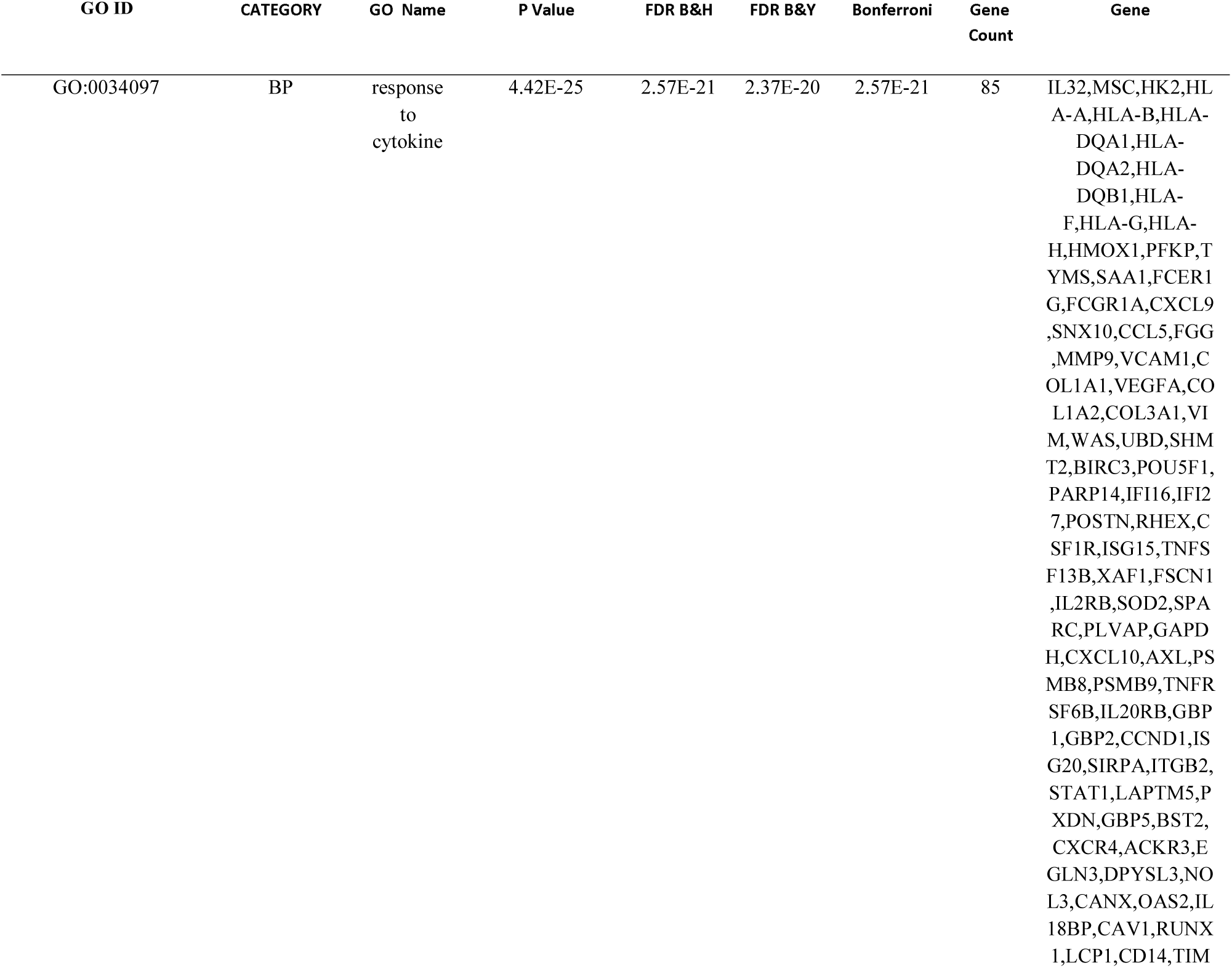
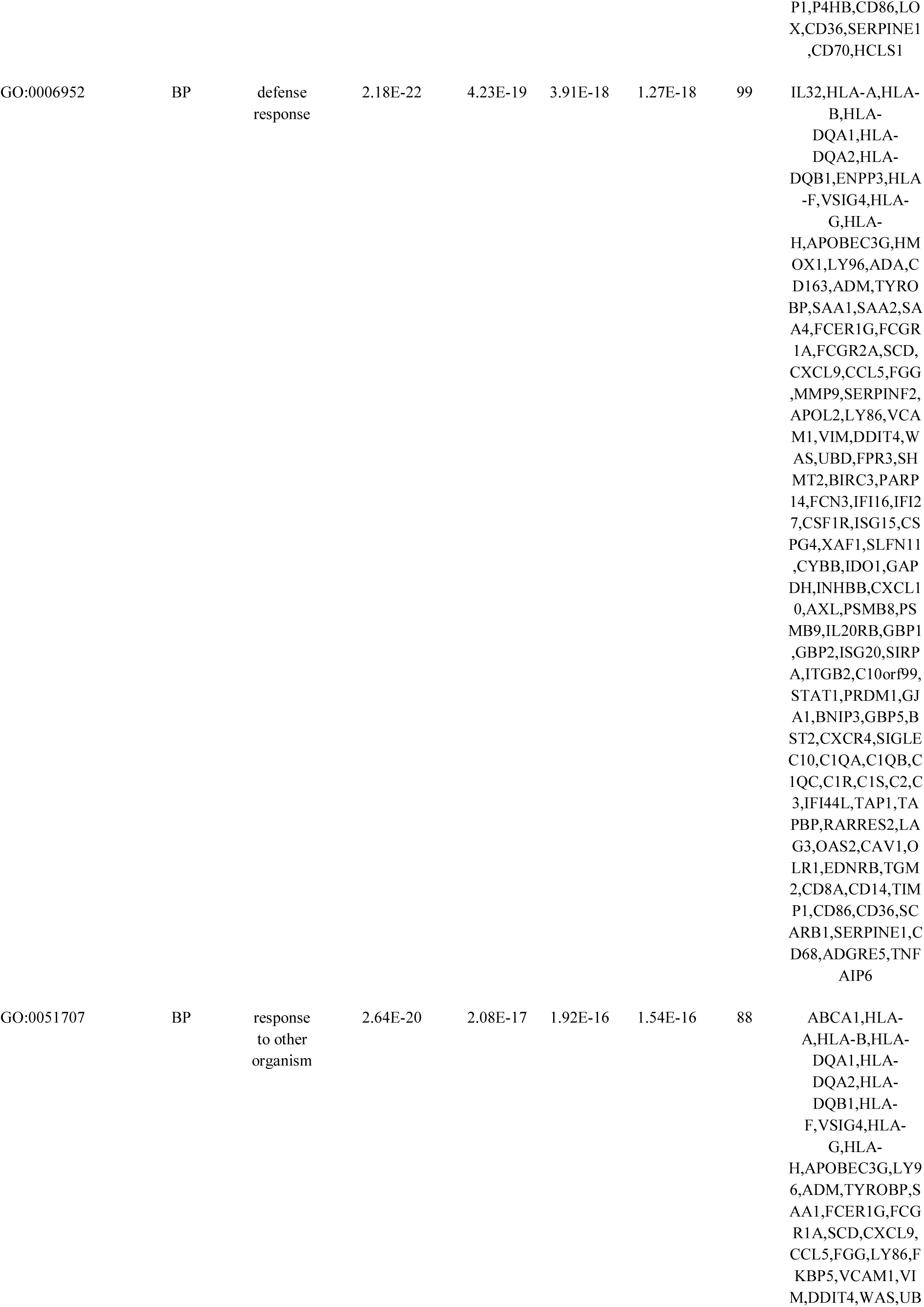
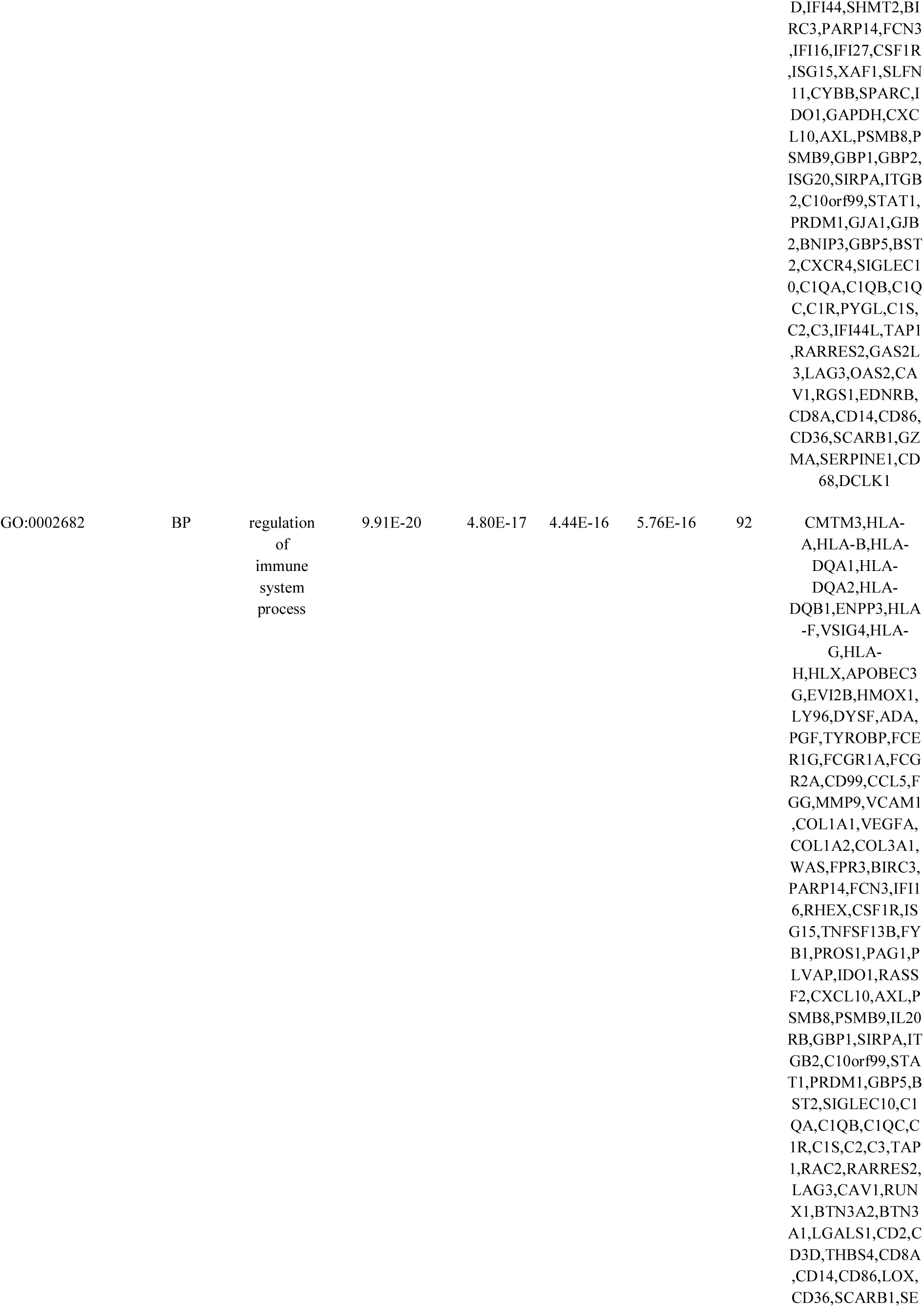
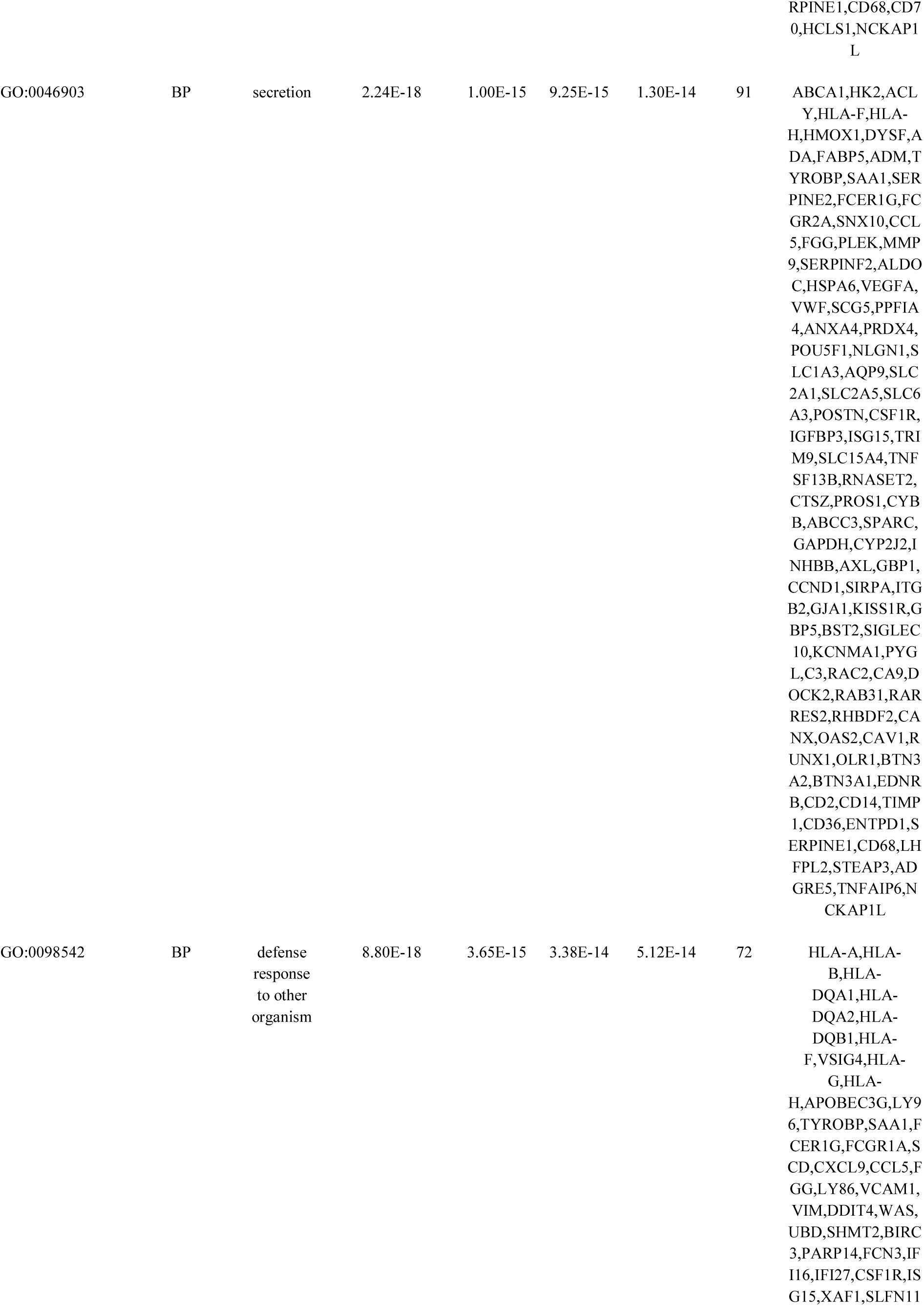
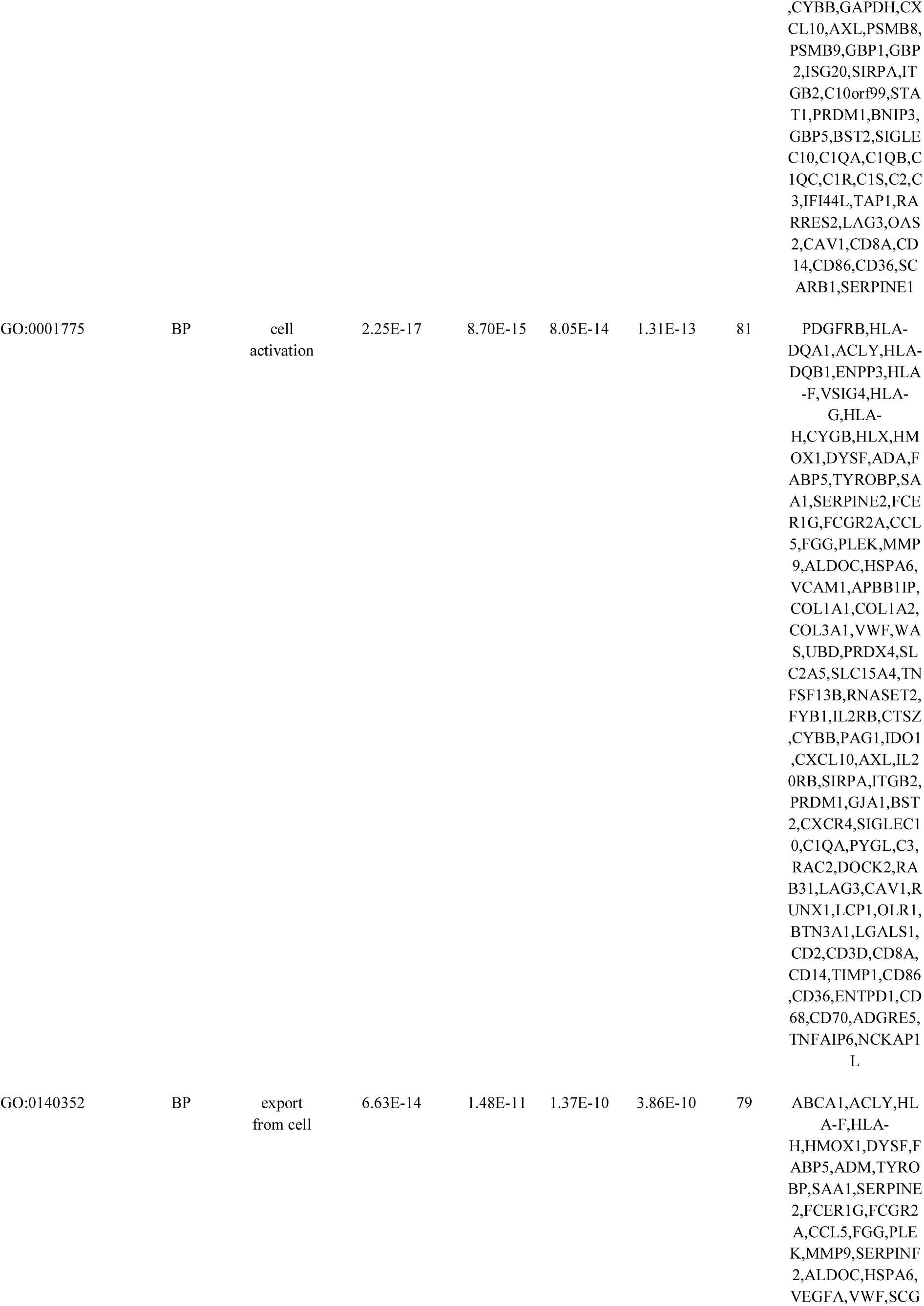
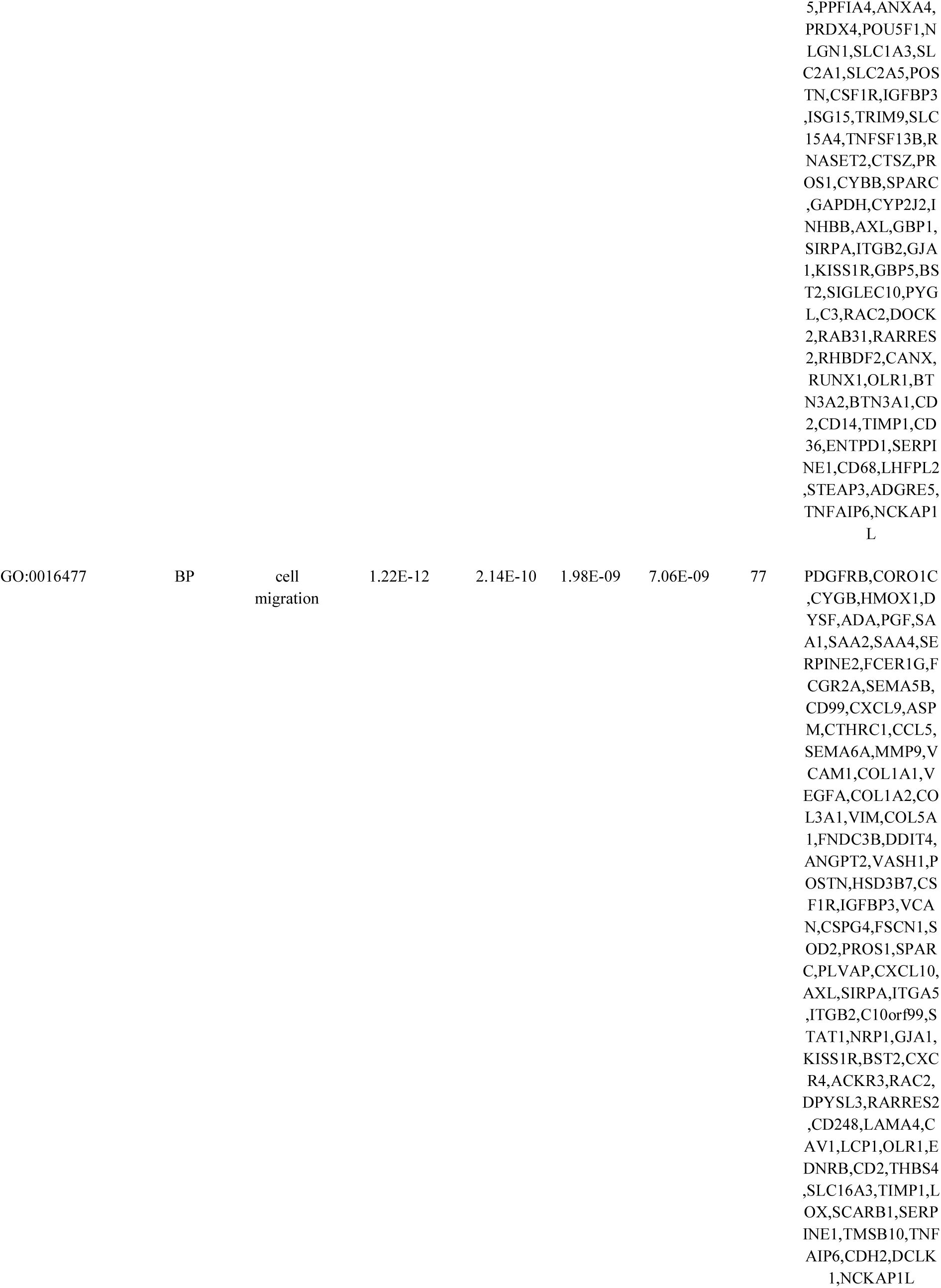
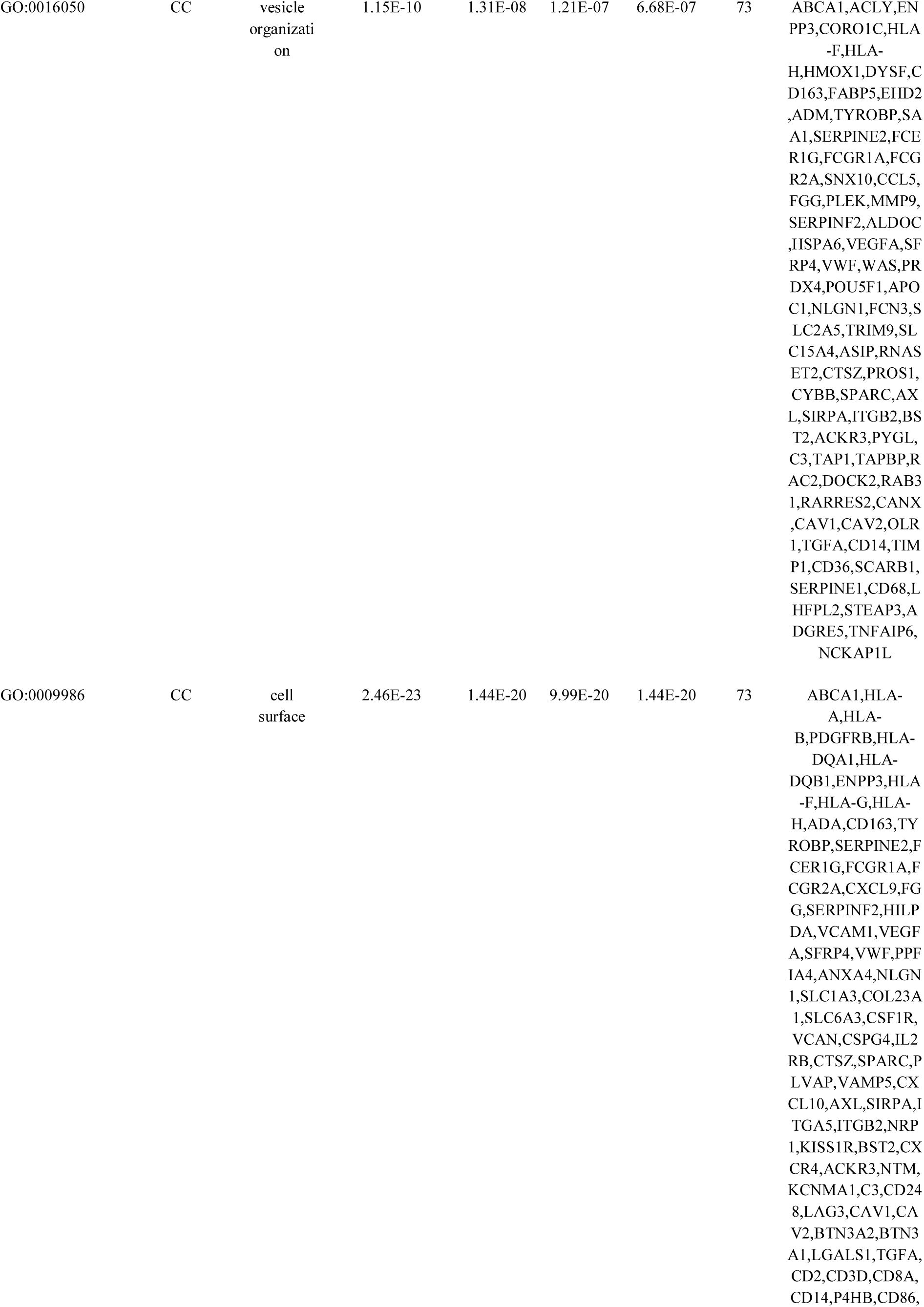
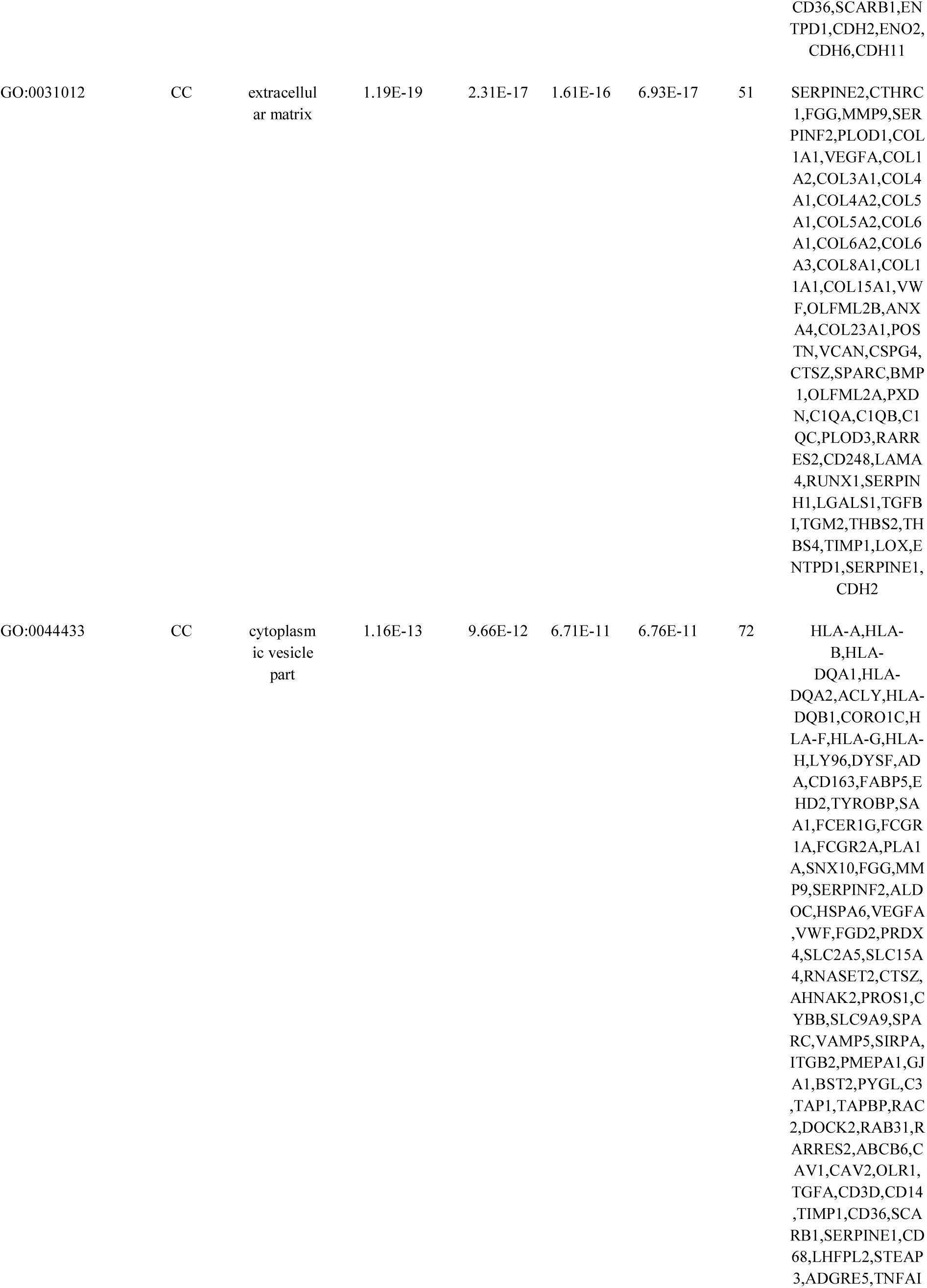
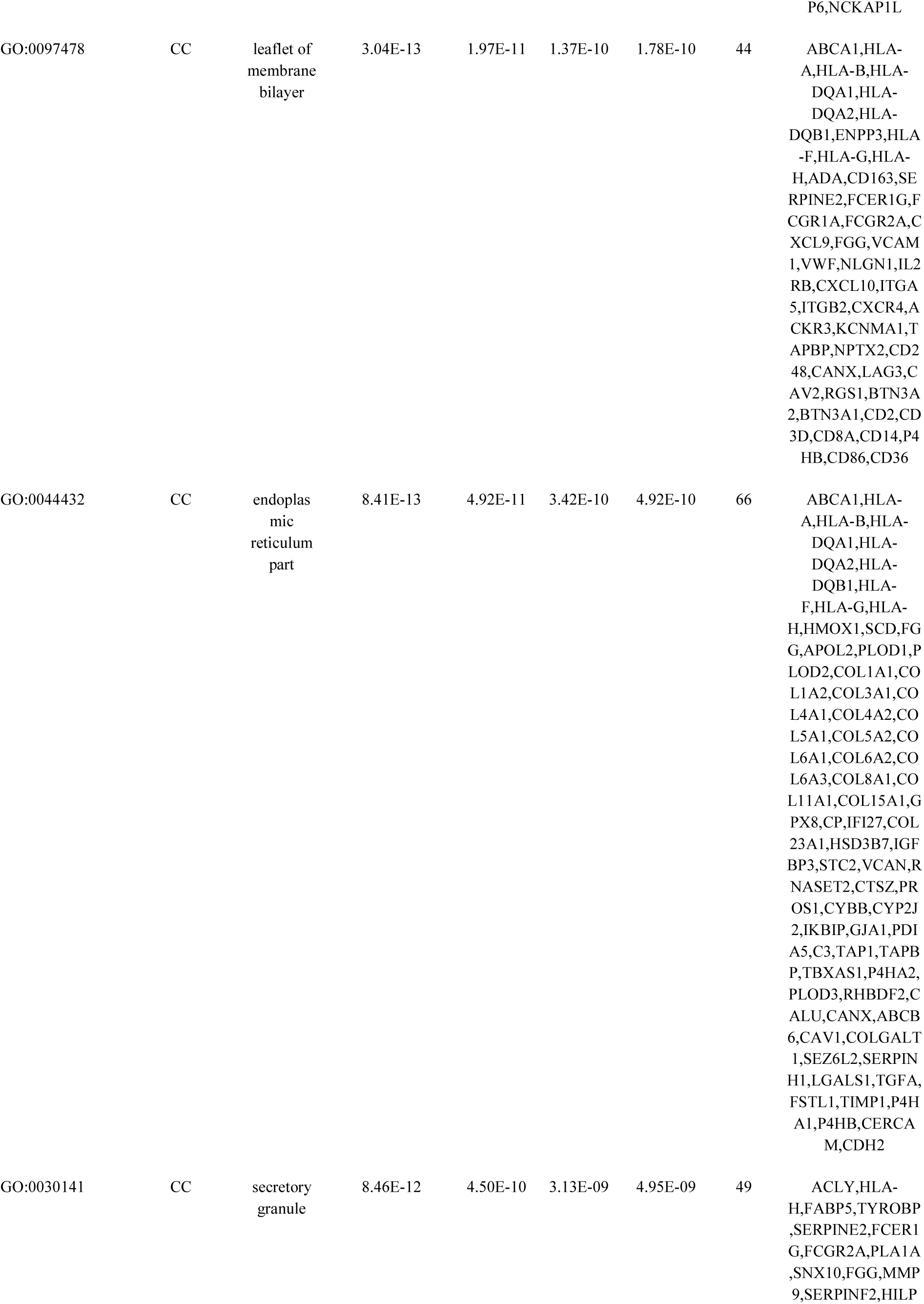
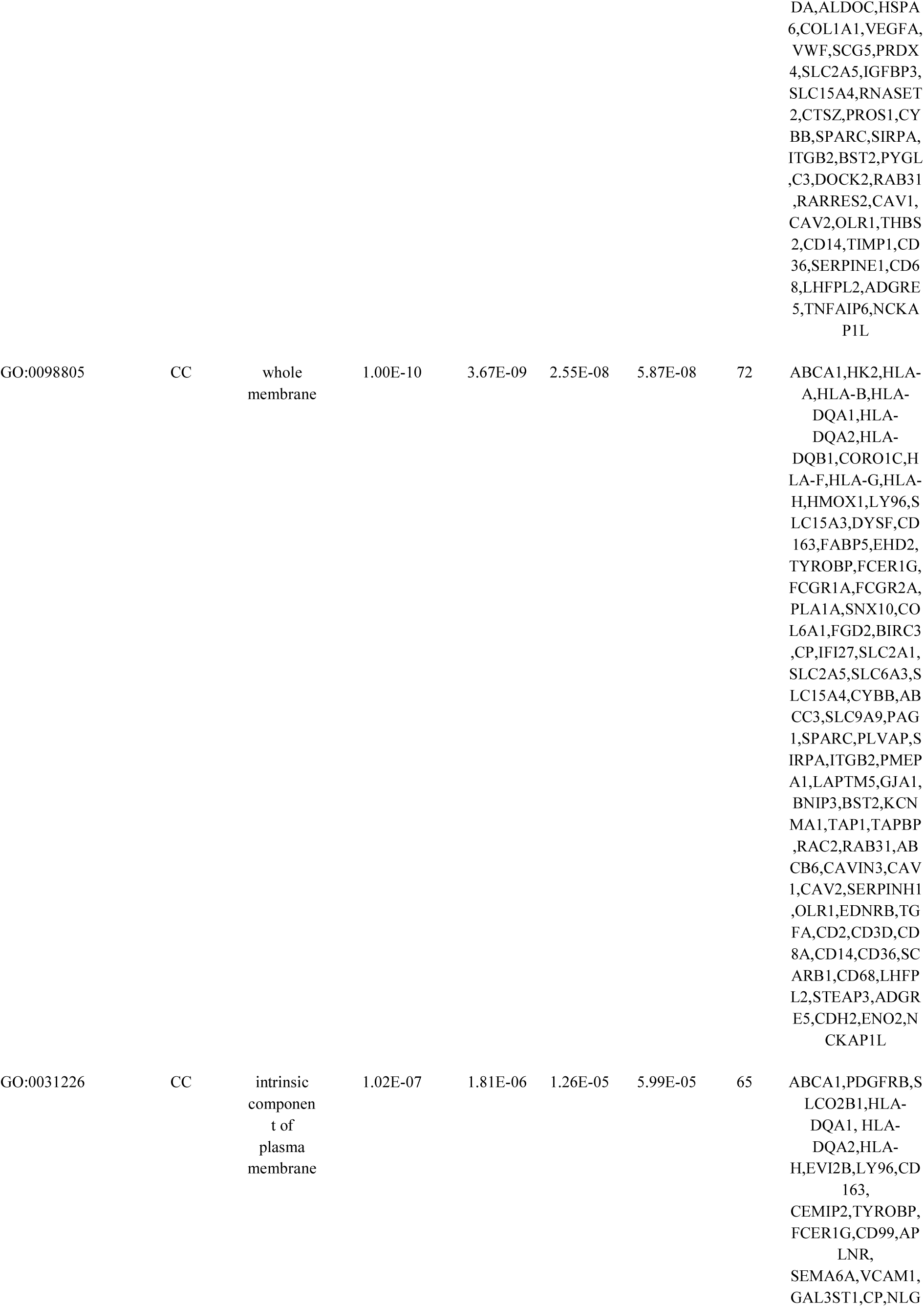
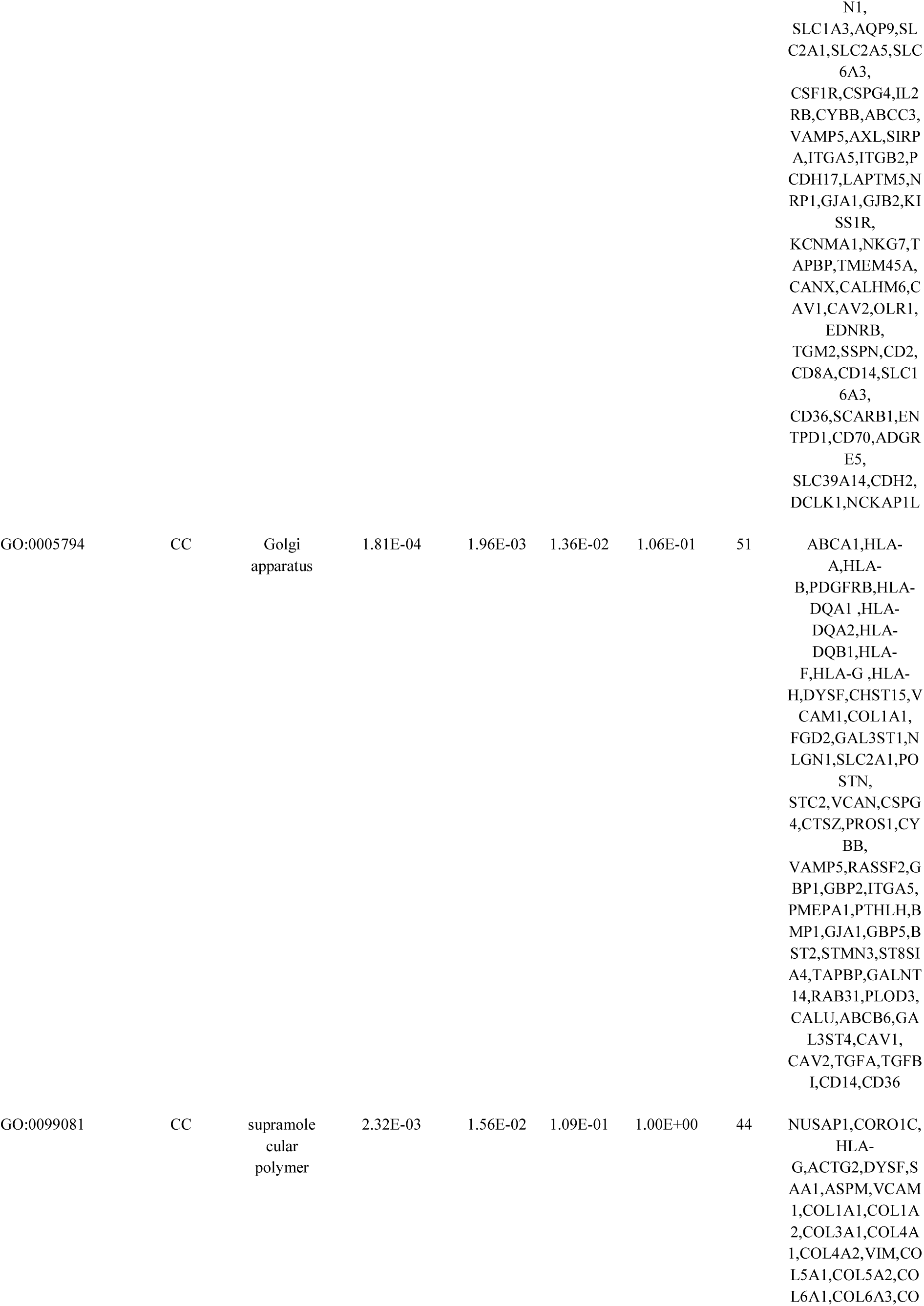
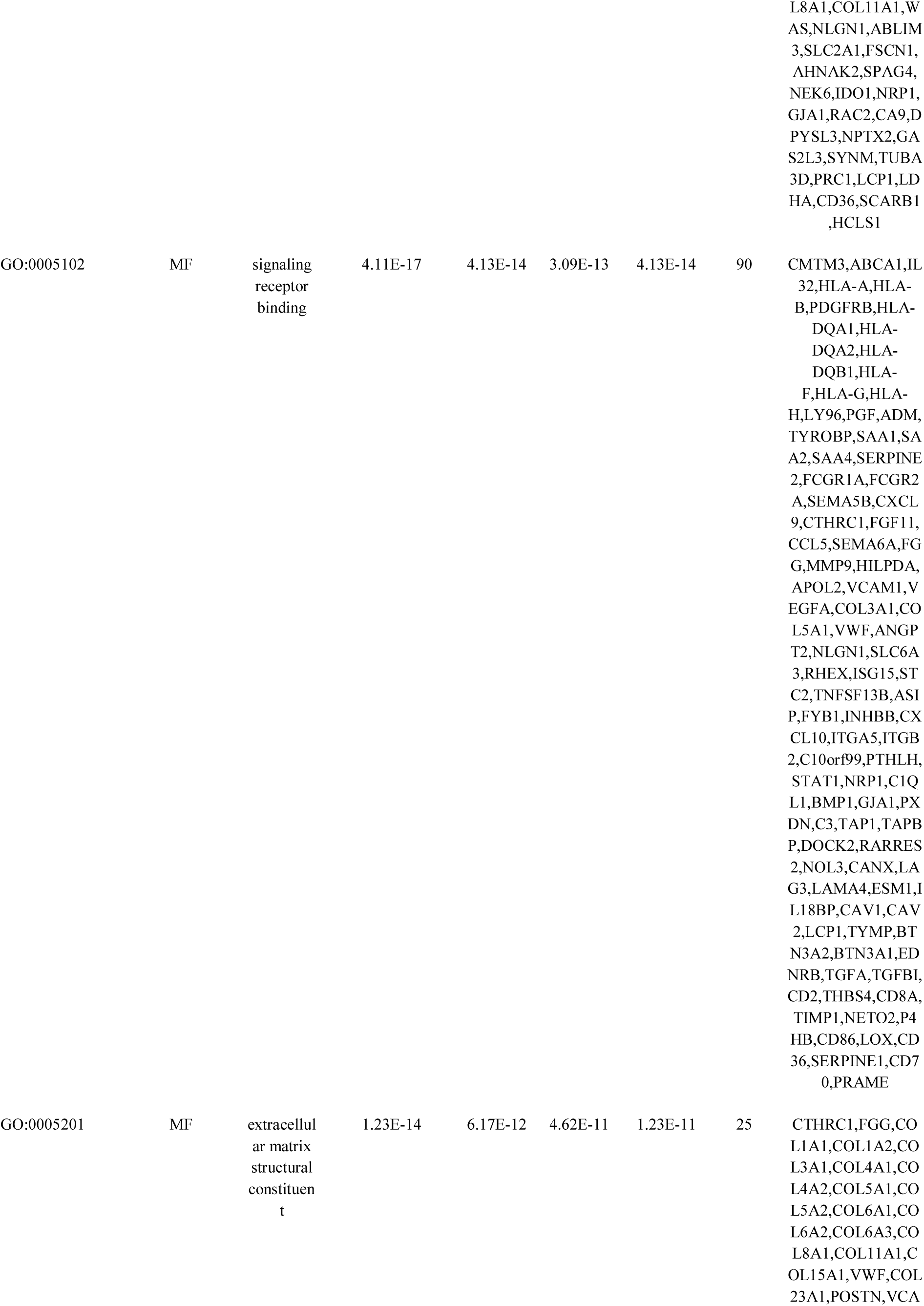
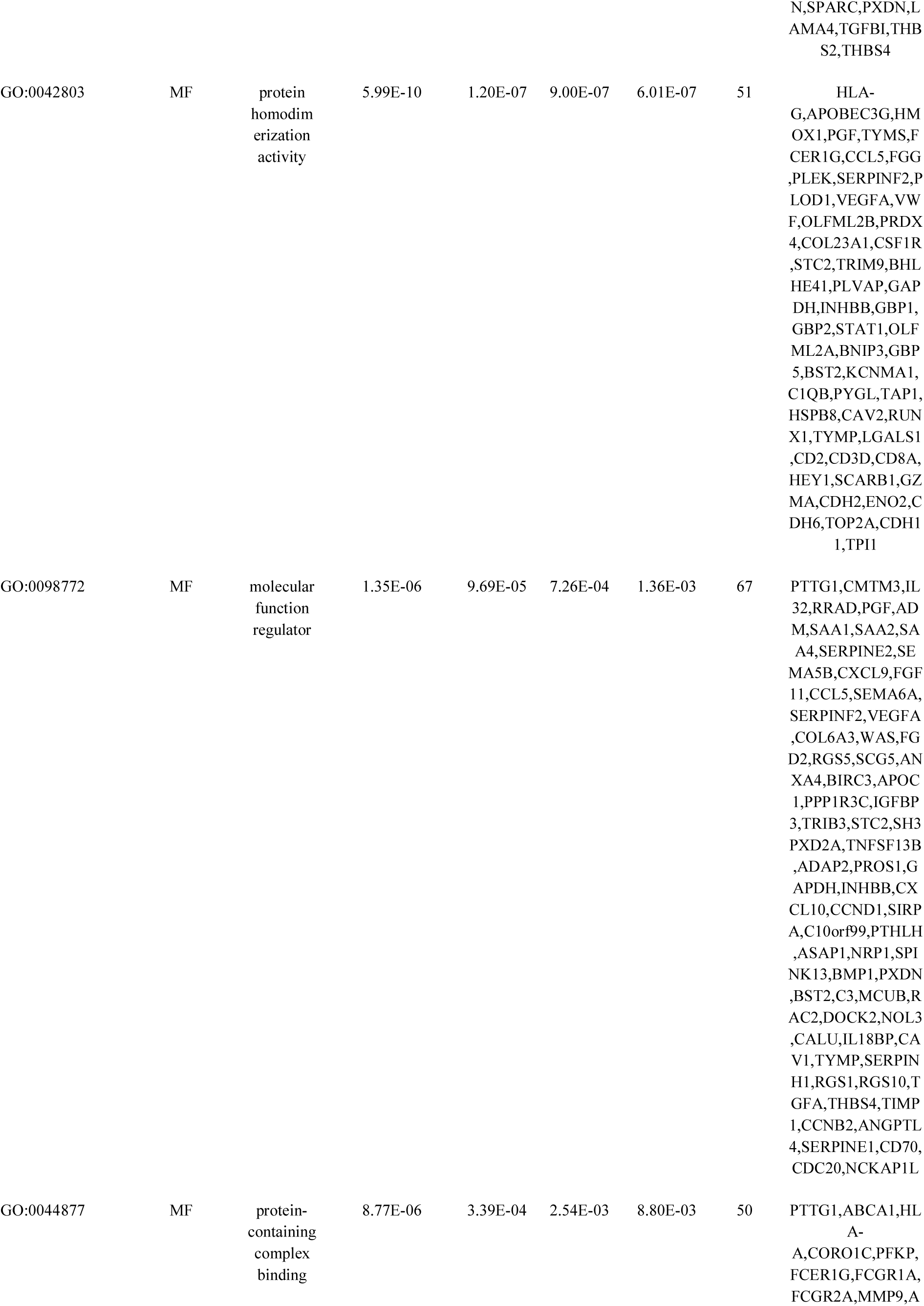
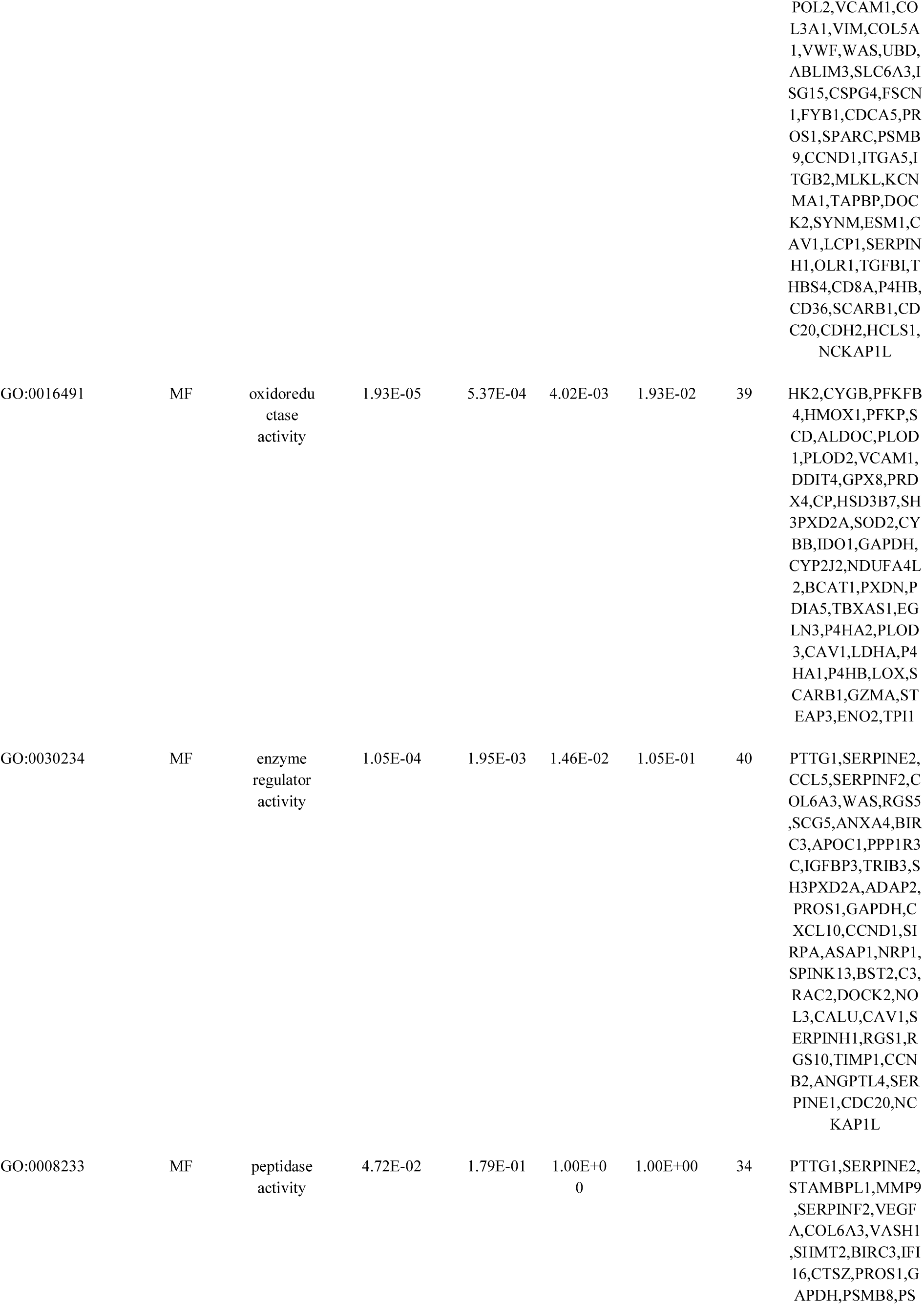
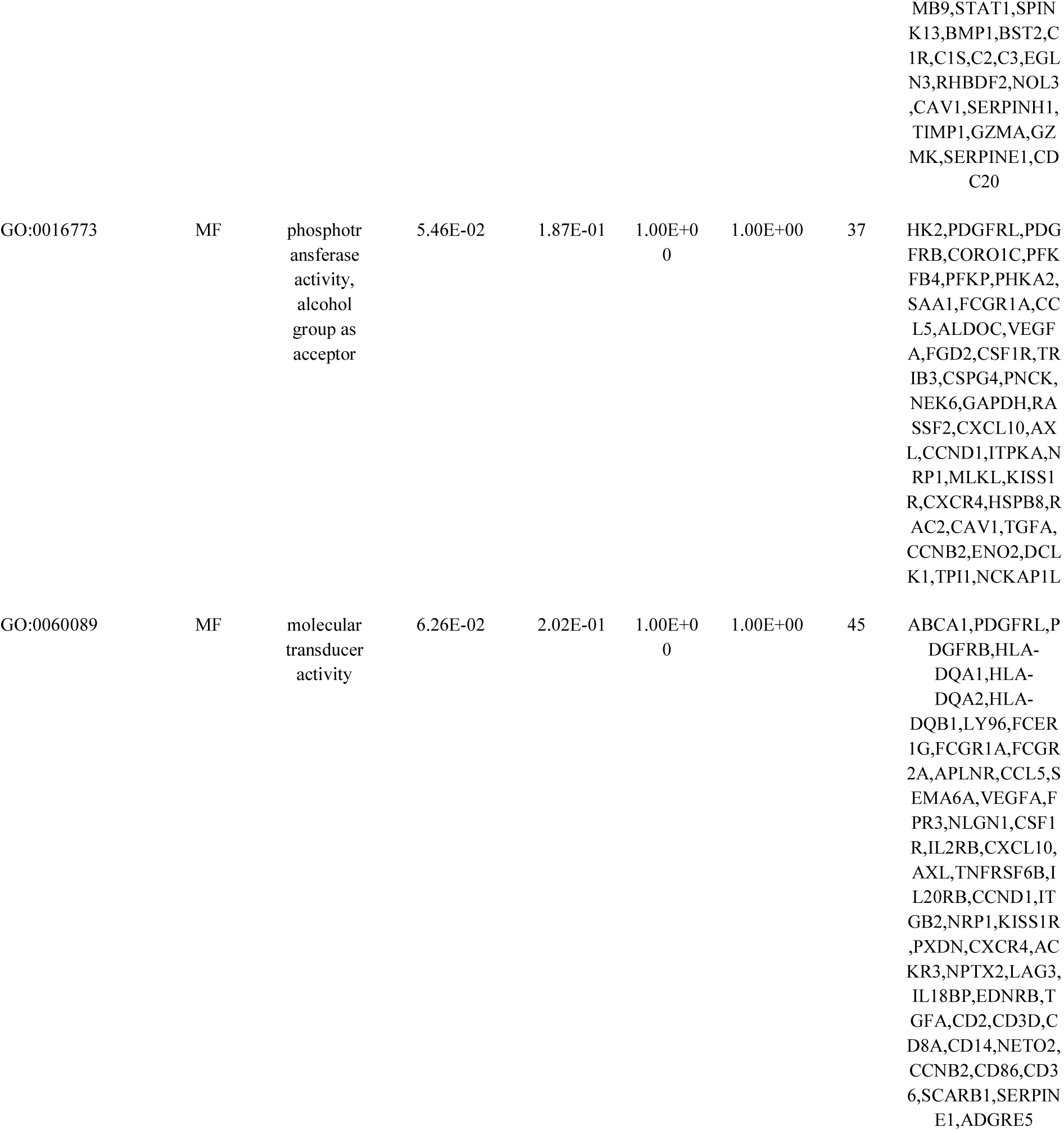
The enriched GO terms of the up regulated differentially expressed genes

| GO ID | CATEGORY | GO Name | P Value | FDR B&H | FDR B&Y | Bonferroni | Gene Count | Gene |
| --- | --- | --- | --- | --- | --- | --- | --- | --- |
| GO:0034097 | BP | response to cytokine | 4.42E-25 | 2.57E-21 | 2.37E-20 | 2.57E-21 | 85 | IL32,MSD,HLA-A,HLA-B,HLA-DQA1,HLA-DQA2,HLA-DQB1,HLA-F,HLA-G,HLA-H,HMOX1,PFKP,TYMS,SAA1,FCER1G,FCGR1A,CXCL9,SNX10,CCL5,FGG,MMP9,VCAM1,COL1A1,VEGFA,COL1A2,COL3A1,VIM,WAS,UBD,SHMT2,BIRC3,POU5F1,PARP14,IFI16,IFI27,POSTN,RHEX,CSF1R,ISG15,TNFSF13B,XAF1,FSCN1,IL2RB,SOD2,SPARC,PLVAP,GAPDH,CXCL10,AXL,PSMB8,PSMB9,TNFRSF6B,IL20RB,GBP1,GBP2,CCND1,ISG20,SIRPA,ITGB2,STAT1,LAPTM5,PXDN,GBP5,BST2,CXCR4,ACKR3,EGLN3,DPYSL3,NOLL3,CANX,OAS2,IL18BP,CAV1,RUNX1,LCP1,CD14,TIM |

|  |  |  |  |  |  |  |  |  |  |
| --- | --- | --- | --- | --- | --- | --- | --- | --- | --- |
|  |  |  |  |  |  |  |  |  | P1,P4HB,CD86,LOX,CD36,SERPINE1,CD70,HCLS1 |
| GO:0006952 | BP | defense response | 2.18E-22 | 4.23E-19 | 3.91E-18 | 1.27E-18 | 99 | IL32,HLA-A,HLA-B,HLA-DQA1,HLA-DQA2,HLA-DQB1,ENPP3,HLA-F,VSIG4,HLA-G,HLA-H,APOBEC3G,HMOX1,LY96,ADA,CD163,ADM,TYROBP,SAA1,SAA2,SAA4,FCER1G,FCGR1A,FCGR2A,SCD,CXCL9,CCL5,FGG,MMP9,SERPINF2,APOL2,LY86,VCAM1,VIM,DDIT4,WAS,UBD,FPR3,SHMT2,BIRC3,PARP14,FCN3,IFI16,IFI27,CSF1R,ISG15,CSPG4,XAF1,SLFN11,CYBB,IDO1,GAPDH,INHBB,CXCL10,AXL,PSMB8,PSMB9,IL20RB,GBP1,GBP2,ISG20,SIRPA,ITGB2,C10orf99,STAT1,PRDM1,GJA1,BNIP3,GBP5,BST2,CXCR4,SIGLEC10,C1QA,C1QB,C1QC,C1R,C1S,C2,C3,IFI44L,TAP1,TAPBP,RARRES2,LAG3,OAS2,CAV1,OLR1,EDNRB,TGM2,CD8A,CD14,TIMP1,CD86,CD36,SCARB1,SERPINE1,CD68,ADGRE5,TNF AIP6 |  |
| GO:0051707 | BP | response to other organism | 2.64E-20 | 2.08E-17 | 1.92E-16 | 1.54E-16 | 88 | ABCA1,HLA-A,HLA-B,HLA-DQA1,HLA-DQA2,HLA-DQB1,HLA-F,VSIG4,HLA-G,HLA-H,APOBEC3G,LY96,ADM,TYROBP,SAA1,FCER1G,FCGR1A,SCD,CXCL9,CCL5,FGG,LY86,FCBP5,VCAM1,VCAM2,DDIT4,WAS,UB |  |
|  |  |  |  |  |  |  |  |  | D,IFI44,SHMT2,BI<br>RC3,PARP14,FCN3<br>,IFI16,IFI27,CSF1R<br>,ISG15,XAF1,SLFN<br>11,CYBB,SPARC,I<br>DO1,GAPDH,CXC<br>L10,AXL,PSMB8,P<br>SMB9,GBP1,GBP2,<br>ISG20,SIRPA,ITGB<br>2,C10orf99,STAT1,<br>PRDM1,GJA1,GJB<br>2,BNIP3,GBP5,BST<br>2,CXCR4,SIGLEC1<br>0,C1QA,C1QB,C1Q<br>C,C1R,PYGL,C1S,<br>C2,C3,IFI44L,TAP1<br>,RARRES2,GAS2L<br>3,LAG3,OAS2,CA<br>V1,RGS1,EDNRB,<br>CD8A,CD14,CD86,<br>CD36,SCARB1,GZ<br>MA,SERPINE1,CD<br>68,DCLK1 |
| GO:0002682 | BP | regulation<br>of<br>immune<br>system<br>process | 9.91E-20 | 4.80E-17 | 4.44E-16 | 5.76E-16 | 92 | CMTM3,HLA-A,HLA-B,HLA-DQA1,HLA-DQA2,HLA-DQB1,ENPP3,HLA<br>-F,VSIG4,HLA-G,HLA-H,HLX,APOBEC3<br>G,EVI2B,HMOX1,<br>LY96,DYSF,ADA,<br>PGF,TYROBP,FCE<br>R1G,FCGR1A,FCG<br>R2A,CD99,CCL5,F<br>GG,MMP9,VCAM1<br>,COL1A1,VEGFA,<br>COL1A2,COL3A1,<br>WAS,FPR3,BIRC3,<br>PARP14,FCN3,IFI1<br>6,RHEX,CSF1R,IS<br>G15,TNFSF13B,FY<br>B1,PROS1,PAG1,P<br>LVAP,IDO1,RASS<br>F2,CXCL10,AXL,P<br>SMB8,PSMB9,IL20<br>RB,GBP1,SIRPA,IT<br>GB2,C10orf99,STA<br>T1,PRDM1,GBP5,B<br>ST2,SIGLEC10,C1<br>QA,C1QB,C1QC,C<br>1R,C1S,C2,C3,TAP<br>1,RAC2,RARRES2,<br>LAG3,CAV1,RUN<br>X1,BTN3A2,BTN3<br>A1,LGALS1,CD2,C<br>D3D,THBS4,CD8A<br>,CD14,CD86,LOX,<br>CD36,SCARB1,SE |  |
|  |  |  |  |  |  |  |  |  | RPINE1,CD68,CD70,HCLS1,NCKAP1L |
| GO:0046903 | BP | secretion | 2.24E-18 | 1.00E-15 | 9.25E-15 | 1.30E-14 | 91 | ABCA1,HK2,ACLY,HLA-F,HLA-H,HMOX1,DYSF,ADA,FABP5,ADM,TYROBP,SAA1,SERPINE2,FCER1G,FCGR2A,SNX10,CCL5,FGG,PLEK,MMP9,SERPINF2,ALDO C,HSPA6,VEGFA,VWF,SCG5,PPFIA4,ANXA4,PRDX4,POU5F1,NLGN1,SLC1A3,AQP9,SLC2A1,SLC2A5,SLC6A3,POSTN,CSF1R,IGFBP3,ISG15,TRIM9,SLC15A4,TNFSF13B,RNASET2,CTSZ,PROS1,CYBB,ABCC3,SPARC,GAPDH,CYP2J2,INHBB,AXL,GBP1,CCND1,SIRPA,ITGB2,GJA1,KISS1R,GBP5,BST2,SIGLEC10,KCNMA1,PYGL,C3,RAC2,CA9,DOCK2,RAB31,RARRES2,RHBDF2,CANX,OAS2,CAV1,RUNX1,OLR1,BTN3A2,BTN3A1,EDNRB,CD2,CD14,TIMP1,CD36,ENTPD1,SERPINE1,CD68,LHFPL2,STEAP3,ADGRE5,TNFAIP6,NCKAP1L |  |
| GO:0098542 | BP | defense response to other organism | 8.80E-18 | 3.65E-15 | 3.38E-14 | 5.12E-14 | 72 | HLA-A,HLA-B,HLA-DQA1,HLA-DQA2,HLA-DQB1,HLA-F,VSIG4,HLA-G,HLA-H,APOBEC3G,LY96,TYROBP,SAA1,FCER1G,FCGR1A,SCD,CXCL9,CCL5,FGG,LY86,VCAM1,VIM,DDIT4,WAS,UBD,SHMT2,BIRC3,PARP14,FCN3,IFI16,IFI27,CSF1R,ISG15,XAF1,SLFN11 |  |
|  |  |  |  |  |  |  |  |  | ,CYBB,GAPDH,CXCL10,AXL,PSMB8,PSMB9,GBP1,GBP2,ISG20,SIRPA,ITGB2,C10orf99,STAT1,PRDM1,BNIP3,GBP5,BST2,SIGLEC10,C1QA,C1QB,C1QC,C1R,C1S,C2,C3,IFI44L,TAP1,RARRRES2,LAG3,OAS2,CAV1,CD8A,CD14,CD86,CD36,SCARB1,SERPINE1 |
| GO:0001775 | BP | cell activation | 2.25E-17 | 8.70E-15 | 8.05E-14 | 1.31E-13 | 81 | PDGFRB,HLA-DQA1,ACLY,HLA-DQB1,ENPP3,HLA-F,VSIG4,HLA-G,HLA-H,CYGB,HLX,HMOX1,DYSF,ADA,FABP5,TYROBP,SAA1,SERPINE2,FCER1G,FCGR2A,CCL5,FGG,PLEK,MMP9,ALDOC,HSPA6,VCAM1,APBB1IP,COL1A1,COL1A2,COL3A1,VWF,WAS,UBD,PRDX4,SLC2A5,SLC15A4,TNFSF13B,RNASET2,FYB1,IL2RB,CTSZ,CYBB,PAG1,IDO1,CXCL10,AXL,IL20RB,SIRPA,ITGB2,PRDM1,GJA1,BST2,CXCR4,SIGLEC10,C1QA,PYGL,C3,RAC2,DOCK2,RAB31,LAG3,CAV1,RUNX1,LCP1,OLR1,BTN3A1,LGALS1,CD2,CD3D,CD8A,CD14,TIMP1,CD86,CD36,ENTPD1,CD68,CD70,ADGRE5,TNFAIP6,NCKAP1L |  |
| GO:0140352 | BP | export from cell | 6.63E-14 | 1.48E-11 | 1.37E-10 | 3.86E-10 | 79 | ABCA1,ACLY,HLA-F,HLA-H,HMOX1,DYSF,FABP5,ADM,TYROBP,SAA1,SERPINE2,FCER1G,FCGR2A,CCL5,FGG,PLEK,MMP9,SERPINF2,ALDOC,HSPA6,VEGFA,VWF,SCG |  |
|  |  |  |  |  |  |  |  | 5,PPFIA4,ANXA4,<br>PRDX4,POU5F1,N<br>LGN1,SLC1A3,SL<br>C2A1,SLC2A5,POS<br>TN,CSF1R,IGFBP3<br>,ISG15,TRIM9,SLC<br>15A4,TNFSF13B,R<br>NASET2,CTSZ,PR<br>OS1,CYBB,SPARC<br>,GAPDH,CYP2J2,I<br>NHBB,AXL,GBP1,<br>SIRPA,ITGB2,GJA<br>1,KISS1R,GBP5,BS<br>T2,SIGLEC10,PYG<br>L,C3,RAC2,DOCK<br>2,RAB31,RARRES<br>2,RHBDF2,CANX,<br>RUNX1,OLR1,BT<br>N3A2,BTN3A1,CD<br>2,CD14,TIMP1,CD<br>36,ENTPD1,SERPI<br>NE1,CD68,LHFPL2<br>,STEAP3,ADGRE5,<br>TNFAIP6,NCKAP1<br>L |
| GO:0016477 | BP | cell<br>migration | 1.22E-12 | 2.14E-10 | 1.98E-09 | 7.06E-09 | 77 | PDGFRB,CORO1C<br>,CYGB,HMOX1,D<br>YSF,ADA,PGF,SA<br>A1,SAA2,SAA4,SE<br>RPINE2,FCER1G,F<br>CGR2A,SEMA5B,<br>CD99,CXCL9,ASP<br>M,CTHRC1,CCL5,<br>SEMA6A,MMP9,V<br>CAM1,COL1A1,V<br>EGFA,COL1A2,CO<br>L3A1,VIM,COL5A<br>1,FNDC3B,DDIT4,<br>ANGPT2,VASH1,P<br>OSTN,HSD3B7,CS<br>F1R,IGFBP3,VCA<br>N,CSPG4,FSCN1,S<br>OD2,PROS1,SPAR<br>C,PLVAP,CXCL10,<br>AXL,SIRPA,ITGA5<br>,ITGB2,C10orf99,S<br>TAT1,NRP1,GJA1,<br>KISS1R,BST2,CXC<br>R4,ACKR3,RAC2,<br>DPYSL3,RARRES2<br>,CD248,LAMA4,C<br>AV1,LCP1,OLR1,E<br>DNRB,CD2,THBS4<br>,SLC16A3,TIMP1,L<br>OX,SCARB1,SERP<br>INE1,TMSB10,TNF<br>AIP6,CDH2,DCLK<br>1,NCKAP1L |
| GO:0016050 | CC | vesicle<br>organizati<br>on | 1.15E-10 | 1.31E-08 | 1.21E-07 | 6.68E-07 | 73 | ABCA1,ACLY,EN<br>PP3,CORO1C,HLA<br>-F,HLA-H,HMOX1,DYSF,C<br>D163,FABP5,EHD2<br>,ADM,TYROBP,SA<br>A1,SERPINE2,FCE<br>R1G,FCGR1A,FCG<br>R2A,SNX10,CCL5,<br>FGG,PLEK,MMP9,<br>SERPINF2,ALDOC<br>,HSPA6,VEGFA,SF<br>RP4,VWF,WAS,PR<br>DX4,POU5F1,APO<br>C1,NLGN1,FCN3,S<br>LC2A5,TRIM9,SL<br>C15A4,ASIP,RNAS<br>ET2,CTSZ,PROS1,<br>CYBB,SPARC,AX<br>L,SIRPA,ITGB2,BS<br>T2,ACKR3,PYGL,<br>C3,TAP1,TAPBP,R<br>AC2,DOCK2,RAB3<br>1,RARRES2,CANX<br>,CAV1,CAV2,OLR<br>1,TGFA,CD14,TIM<br>P1,CD36,SCARB1,<br>SERPINE1,CD68,L<br>HFPL2,STEAP3,A<br>DGRE5,TNFAIP6,<br>NCKAP1L |
| GO:0009986 | CC | cell<br>surface | 2.46E-23 | 1.44E-20 | 9.99E-20 | 1.44E-20 | 73 | ABCA1,HLA-A,HLA-B,PDGFRB,HLA-DQA1,HLA-DQB1,ENPP3,HLA<br>-F,HLA-G,HLA-H,ADA,CD163,TY<br>ROBP,SERPINE2,F<br>CER1G,FCGR1A,F<br>CGR2A,CXCL9,FG<br>G,SERPINF2,HILP<br>DA,VCAM1,VEGF<br>A,SFRP4,VWF,PPF<br>IA4,ANXA4,NLGN<br>1,SLC1A3,COL23A<br>1,SLC6A3,CSF1R,<br>VCAN,CSPG4,IL2<br>RB,CTSZ,SPARC,P<br>LVAP,VAMP5,CX<br>CL10,AXL,SIRPA,I<br>TGA5,ITGB2,NRP<br>1,KISS1R,BST2,CX<br>CR4,ACKR3,NTM,<br>KCNMA1,C3,CD24<br>8,LAG3,CAV1,CA<br>V2,BTN3A2,BTN3<br>A1,LGALS1,TGFA,<br>CD2,CD3D,CD8A,<br>CD14,P4HB,CD86, |
|  |  |  |  |  |  |  |  | CD36,SCARB1,EN<br>TPD1,CDH2,ENO2,<br>CDH6,CDH11 |
| GO:0031012 | CC | extracellul<br>ar matrix | 1.19E-19 | 2.31E-17 | 1.61E-16 | 6.93E-17 | 51 | SERPINE2,CTHRC<br>1,FGG,MMP9,SER<br>PINF2,PLOD1,COL<br>1A1,VEGFA,COL1<br>A2,COL3A1,COL4<br>A1,COL4A2,COL5<br>A1,COL5A2,COL6<br>A1,COL6A2,COL6<br>A3,COL8A1,COL1<br>1A1,COL15A1,VW<br>F,OLFML2B,ANX<br>A4,COL23A1,POS<br>TN,VCAN,CSPG4,<br>CTS2,SPARC,BMP<br>1,OLFML2A,PXD<br>N,C1QA,C1QB,C1<br>QC,PLOD3,RARR<br>ES2,CD248,LAMA<br>4,RUNX1,SERPIN<br>H1,LGALS1,TGFB<br>I,TGM2,THBS2,TH<br>BS4,TIMP1,LOX,E<br>NTPD1,SERPINE1,<br>CDH2 |
| GO:0044433 | CC | cytoplasm<br>ic vesicle<br>part | 1.16E-13 | 9.66E-12 | 6.71E-11 | 6.76E-11 | 72 | HLA-A,HLA-B,HLA-DQA1,HLA-DQA2,ACLY,HLA-DQB1,CORO1C,H<br>LA-F,HLA-G,HLA-H,LY96,DYSF,AD<br>A,CD163,FABP5,E<br>HD2,TYROBP,SA<br>A1,FCER1G,FCGR<br>1A,FCGR2A,PLA1<br>A,SNX10,FGG,MM<br>P9,SERPINF2,ALD<br>OC,HSPA6,VEGFA<br>,VWF,FGD2,PRDX<br>4,SLC2A5,SLC15A<br>4,RNASET2,CTS2,<br>AHNAK2,PROS1,C<br>YBB,SLC9A9,SPA<br>RC,VAMP5,SIRPA,<br>ITGB2,PMEPA1,GJ<br>A1,BST2,PYGL,C3<br>,TAP1,TAPBP,RAC<br>2,DOCK2,RAB31,R<br>ARRES2,ABCB6,C<br>AV1,CAV2,OLR1,<br>TGFA,CD3D,CD14<br>,TIMP1,CD36,SCA<br>RB1,SERPINE1,CD<br>68,LHFPL2,STEAP<br>3,ADGRE5,TNFAI |
|  |  |  |  |  |  |  |  | P6,NCKAP1L |
| GO:0097478 | CC | leaflet of membrane bilayer | 3.04E-13 | 1.97E-11 | 1.37E-10 | 1.78E-10 | 44 | ABCA1,HLA-A,HLA-B,HLA-DQA1,HLA-DQA2,HLA-DQB1,ENPP3,HLA-F,HLA-G,HLA-H,ADA,CD163,SERPINE2,FCER1G,FCGR1A,FCGR2A,CXCL9,FGG,VCAM1,VWF,NLGN1,IL2RB,CXCL10,ITGA5,ITGB2,CXCR4,ACKR3,KCNMA1,TAPBP,NPTX2,CD248,CANX,LAG3,CAV2,RGS1,BTN3A2,BTN3A1,CD2,CD3D,CD8A,CD14,P4HB,CD86,CD36 |
| GO:0044432 | CC | endoplasmic reticulum part | 8.41E-13 | 4.92E-11 | 3.42E-10 | 4.92E-10 | 66 | ABCA1,HLA-A,HLA-B,HLA-DQA1,HLA-DQA2,HLA-DQB1,HLA-F,HLA-G,HLA-H,HMOX1,SCD,FGG,APOL2,PLOD1,PLOD2,COL1A1,COL1A2,COL3A1,COL4A1,COL4A2,COL5A1,COL5A2,COL6A1,COL6A2,COL6A3,COL8A1,COL11A1,COL15A1,GPX8,CP,IFI27,COL23A1,HSD3B7,IGFBP3,STC2,VCAN,RNASET2,CTS2,PRSS1,CYBB,CYP2J2,IKBIP,GJA1,PDI A5,C3,TAP1,TAPBP,TBXAS1,P4HA2,PLOD3,RHBDF2,CALU,CANX,ABCB6,CAV1,COLGALT1,SEZ6L2,SERPINH1,LGALS1,TGFA,FSTL1,TIMP1,P4HA1,P4HB,CERCAM,CDH2 |
| GO:0030141 | CC | secretory granule | 8.46E-12 | 4.50E-10 | 3.13E-09 | 4.95E-09 | 49 | ACLY,HLA-H,FABP5,TYROBP,SERPINE2,FCER1G,FCGR2A,PLA1A,SNX10,FGG,MMP9,SERPINF2,HILP |

|  |  |  |  |  |  |  |  |  |  |
| --- | --- | --- | --- | --- | --- | --- | --- | --- | --- |
|  |  |  |  |  |  |  |  |  | DA,ALDOC,HSPA6,COL1A1,VEGFA,VWF,SCG5,PRDX4,SLC2A5,IGFBP3,SLC15A4,RNASET2,CTSZ,PROS1,CYBB,SPARC,SIRPA,ITGB2,BST2,PYGL,C3,DOCK2,RAB31,RARRES2,CAV1,CAV2,OLR1,THBS2,CD14,TIMP1,CD36,SERPINE1,CD68,LHFPL2,ADGRE5,TNFAIP6,NCKA1P1L |
| GO:0098805 | CC | whole membrane | 1.00E-10 | 3.67E-09 | 2.55E-08 | 5.87E-08 | 72 | ABCA1,HK2,HLA-A,HLA-B,HLA-DQA1,HLA-DQA2,HLA-DQB1,CORO1C,HLA-F,HLA-G,HLA-H,HMOX1,LY96,SLC15A3,DYSF,CD163,FABP5,EHD2,TYROBP,FCER1G,FCGR1A,FCGR2A,PLA1A,SNX10,COL6A1,FGD2,BIRC3,CP,IFI27,SLC2A1,SLC2A5,SLC6A3,SLC15A4,CYBB,ABCC3,SLC9A9,PAG1,SPARC,PLVAP,SIRPA,ITGB2,PMEPA1,LAPTM5,GJA1,BNIP3,BST2,KCNMA1,TAP1,TAPBP,RAC2,RAB31,ABCB6,CAVIN3,CAV1,CAV2,SERPINH1,OLR1,EDNRB,TGFA,CD2,CD3D,CD8A,CD14,CD36,SCARB1,CD68,LHFPL2,STEAP3,ADGRE5,CDH2,ENO2,NCKAP1L |  |
| GO:0031226 | CC | intrinsic component of plasma membrane | 1.02E-07 | 1.81E-06 | 1.26E-05 | 5.99E-05 | 65 | ABCA1,PDGFRB,SLC2B1,HLA-DQA1,HLA-DQA2,HLA-DQB1,EVI2B,LY96,CD163,CEMIP2,TYROBP,FCER1G,CD99,APLNR,SEMA6A,VCAM1,GAL3ST1,CP,NLG |  |
|  |  |  |  |  |  |  |  |  | N1,<br>SLC1A3,AQP9,SLC2A1,SLC2A5,SLC6A3,<br>CSF1R,CSPG4,IL2RB,CYBB,ABCC3,VAMP5,AXL,SIRPA,ITGA5,ITGB2,PCDH17,LAPTM5,NRP1,GJA1,GJB2,KISS1R,<br>KCNMA1,NKG7,TAPBP,TMEM45A,CANX,CALHM6,CAV1,CAV2,OLR1,EDNRB,<br>TGM2,SSPN,CD2,CD8A,CD14,SLC16A3,<br>CD36,SCARB1,ENTPD1,CD70,ADGRE5,<br>SLC39A14,CDH2,DCLK1,NCKAP1L |
| GO:0005794 | CC | Golgi apparatus | 1.81E-04 | 1.96E-03 | 1.36E-02 | 1.06E-01 | 51 | ABCA1,HLA-A,HLA-B,PDGFRB,HLA-DQA1,HLA-DQA2,HLA-DQB1,HLA-F,HLA-G,HLA-H,DYSF,CHST15,VCAM1,COL1A1,FGD2,GAL3ST1,NLGN1,SLC2A1,POSTN,<br>STC2,VCAN,CSPG4,CTSZ,PROS1,CYBB,<br>VAMP5,RASSF2,GBP1,GBP2,ITGA5,PMEPA1,PTHLH,BMP1,GJA1,GBP5,BST2,STMN3,ST8SIA4,TAPBP,GALNT14,RAB31,PLOD3,CALU,ABCB6,GAL3ST4,CAV1,CAV2,TGFA,TGFB1,CD14,CD36 |  |
| GO:0099081 | CC | supramolecular polymer | 2.32E-03 | 1.56E-02 | 1.09E-01 | 1.00E+00 | 44 | NUSAP1,CORO1C,HLA-G,ACTG2,DYSF,SSAA1,ASPM,VCAM1,COL1A1,COL1A2,COL3A1,COL4A1,COL4A2,VIM,COL5A1,COL5A2,COL6A1,COL6A3,CO |  |
|  |  |  |  |  |  |  |  |  | L8A1,COL11A1,W<br>AS,NLGN1,ABLM<br>3,SLC2A1,FSCN1,<br>AHNAK2,SPAG4,<br>NEK6,IDO1,NRP1,<br>GJA1,RAC2,CA9,D<br>PYSL3,NPTX2,GA<br>S2L3,SYNM,TUBA<br>3D,PRC1,LCP1,LD<br>HA,CD36,SCARB1<br>,HCLS1 |
| GO:0005102 | MF | signaling<br>receptor<br>binding | 4.11E-17 | 4.13E-14 | 3.09E-13 | 4.13E-14 | 90 | CMTM3,ABCA1,IL<br>32,HLA-A,HLA-B,PDGFRB,HLA-DQA1,HLA-DQA2,HLA-DQB1,HLA-F,HLA-G,HLA-H,LY96,PGF,ADM,<br>TYROBP,SAA1,SA<br>A2,SAA4,SERPINE<br>2,FCGR1A,FCGR2<br>A,SEMA5B,CXCL<br>9,CTHRC1,FGF11,<br>CCL5,SEMA6A,FG<br>G,MMP9,HILPDA,<br>APOL2,VCAM1,V<br>EGFA,COL3A1,CO<br>L5A1,VWF,ANGP<br>T2,NLGN1,SLC6A<br>3,RHEX,ISG15,ST<br>C2,TNFSF13B,ASI<br>P,FYB1,INHBB,CX<br>CL10,ITGA5,ITGB<br>2,C10orf99,PTHLH,<br>STAT1,NRP1,C1Q<br>L1,BMP1,GJA1,PX<br>DN,C3,TAP1,TAPB<br>P,DOCK2,RARRES<br>2,NOL3,CANX,LA<br>G3,LAMA4,ESM1,I<br>L18BP,CAV1,CAV<br>2,LCP1,TYMP,BT<br>N3A2,BTN3A1,ED<br>NRB,TGFA,TGFBI,<br>CD2,THBS4,CD8A,<br>TIMP1,NETO2,P4<br>HB,CD86,LOX,CD<br>36,SERPINE1,CD7<br>0,PRAME |  |
| GO:0005201 | MF | extracellul<br>ar matrix<br>structural<br>constituen<br>t | 1.23E-14 | 6.17E-12 | 4.62E-11 | 1.23E-11 | 25 | CTHRC1,FGG,CO<br>L1A1,COL1A2,CO<br>L3A1,COL4A1,CO<br>L4A2,COL5A1,CO<br>L5A2,COL6A1,CO<br>L6A2,COL6A3,CO<br>L8A1,COL11A1,C<br>OL15A1,VWF,COL<br>23A1,POSTN,VCA |  |
|  |  |  |  |  |  |  |  | N,SPARC,PXDN,L<br>AMA4,TGFBI,THB<br>S2,THBS4 |
| GO:0042803 | MF | protein<br>homodim<br>erization<br>activity | 5.99E-10 | 1.20E-07 | 9.00E-07 | 6.01E-07 | 51 | HLA-G,APOBEC3G,HM<br>OX1,PGF,TYMS,F<br>CER1G,CCL5,FGG<br>,PLEK,SERPINF2,P<br>LOD1,VEGFA,VW<br>F,OLFML2B,PRDX<br>4,COL23A1,CSF1R<br>,STC2,TRIM9,BHL<br>HE41,PLVAP,GAP<br>DH,INHBB,GBP1,<br>GBP2,STAT1,OLF<br>ML2A,BNIP3,GBP<br>5,BST2,KCNMA1,<br>C1QB,PYGL,TAP1,<br>HSPB8,CAV2,RUN<br>X1,TYMP,LGALS1<br>,CD2,CD3D,CD8A,<br>HEY1,SCARB1,GZ<br>MA,CDH2,ENO2,C<br>DH6,TOP2A,CDH1<br>1,TPII |
| GO:0098772 | MF | molecular<br>function<br>regulator | 1.35E-06 | 9.69E-05 | 7.26E-04 | 1.36E-03 | 67 | PTTG1,CMTM3,IL<br>32,RRAD,PGF,AD<br>M,SAA1,SAA2,SA<br>A4,SERPINE2,SE<br>MA5B,CXCL9,FGF<br>11,CCL5,SEMA6A,<br>SERPINF2,VEGFA<br>,COL6A3,WAS,FG<br>D2,RGS5,SCG5,AN<br>XA4,BIRC3,APOC<br>1,PPP1R3C,IGFBP<br>3,TRIB3,STC2,SH3<br>PXD2A,TNFSF13B<br>,ADAP2,PROS1,G<br>APDH,INHBB,CX<br>CL10,CCND1,SIRP<br>A,C10orf99,PTHLH<br>,ASAP1,NRP1,SPI<br>NK13,BMP1,PXDN<br>,BST2,C3,MCUB,R<br>AC2,DOCK2,NOL3<br>,CALU,IL18BP,CA<br>V1,TYMP,SERPIN<br>H1,RGS1,RGS10,T<br>GFA,THBS4,TIMP<br>1,CCNB2,ANGPTL<br>4,SERPINE1,CD70,<br>CDC20,NCKAP1L |
| GO:0044877 | MF | protein-containing<br>complex<br>binding | 8.77E-06 | 3.39E-04 | 2.54E-03 | 8.80E-03 | 50 | PTTG1,ABCA1,HL<br>A-A,CORO1C,PFKP,<br>FCER1G,FCGR1A,<br>FCGR2A,MMP9,A |

|  |  |  |  |  |  |  |  |  |  |
| --- | --- | --- | --- | --- | --- | --- | --- | --- | --- |
|  |  |  |  |  |  |  |  |  | POL2,VCAM1,COL3A1,VIM,COL5A1,VWF,WAS,UBD,ABLM3,SLC6A3,ISG15,CSPG4,FSCN1,FYB1,CDCA5,PROS1,SPARC,PSMB9,CCND1,ITGA5,ITGB2,MLKL,KCNMA1,TAPBP,DOCK2,SYNM,ESM1,CAV1,LCP1,SERPINH1,OLR1,TGFBI,TBBS4,CD8A,P4HB,CD36,SCARB1,CD20,CDH2,HCLS1,NCKAP1L |
| GO:0016491 | MF | oxidoreductase activity | 1.93E-05 | 5.37E-04 | 4.02E-03 | 1.93E-02 | 39 | HK2,CYGB,PFKFB4,HMOX1,PFKP,SCD,ALDOC,PLOD1,PLOD2,VCAM1,DDIT4,GPX8,PRDX4,CP,HSD3B7,SH3PXD2A,SOD2,CYBB,IDO1,GAPDH,CYP2J2,NDUFA4L2,BCAT1,PXDN,PDIAS,TBXAS1,EGLN3,P4HA2,PLOD3,CAV1,LDHA,P4HA1,P4HB,LOX,SCARB1,GZMA,STEAP3,ENO2,TPI1 |  |
| GO:0030234 | MF | enzyme regulator activity | 1.05E-04 | 1.95E-03 | 1.46E-02 | 1.05E-01 | 40 | PTTG1,SERPINE2,CCL5,SERPINF2,COL6A3,WAS,RGS5,SCG5,ANXA4,BIRC3,APOC1,PPP1R3C,IGFBP3,TRIB3,SH3PXD2A,ADAP2,PROS1,GAPDH,CXCL10,CCND1,SIRPA,ASAP1,NRP1,SPINK13,BST2,C3,RAC2,DOCK2,NOLL3,CALU,CAV1,SERPINH1,RGS1,RGS10,TIMP1,CCNB2,ANGPTL4,SERPINE1,CDC20,NCKAP1L |  |
| GO:0008233 | MF | peptidase activity | 4.72E-02 | 1.79E-01 | 1.00E+00 | 1.00E+00 | 34 | PTTG1,SERPINE2,STAMBPL1,MMP9,SERPINF2,VEGFA,COL6A3,VASH1,SHMT2,BIRC3,IFI16,CTSZ,PROS1,GAPDH,PSMB8,PS |  |
|  |  |  |  |  |  |  |  |  | MB9,STAT1,SPIN<br>K13,BMP1,BST2,C<br>1R,C1S,C2,C3,EGL<br>N3,RHBD2,NOL3<br>,CAV1,SERPINH1,<br>TIMP1,GZMA,GZ<br>MK,SERPINE1,CD<br>C20 |
| GO:0016773 | MF | phosphotr<br>ansferase<br>activity,<br>alcohol<br>group as<br>acceptor | 5.46E-02 | 1.87E-01 | 1.00E+0<br>0 | 1.00E+00 | 37 | HK2,PDGFRL,PDG<br>FRB,CORO1C,PFK<br>FB4,PFKP,PHKA2,<br>SAA1,FCGR1A,CC<br>L5,ALDOC,VEGF<br>A,FGD2,CSF1R,TR<br>IB3,CSPG4,PNCK,<br>NEK6,GAPDH,RA<br>SSF2,CXCL10,AX<br>L,CCND1,ITPKA,N<br>RP1,MLKL,KISS1<br>R,CXCR4,HSPB8,R<br>AC2,CAV1,TGFA,<br>CCNB2,ENO2,DCL<br>K1,TPI1,NCKAP1L |  |
| GO:0060089 | MF | molecular<br>transducer<br>activity | 6.26E-02 | 2.02E-01 | 1.00E+0<br>0 | 1.00E+00 | 45 | ABCA1,PDGFRL,P<br>DGFRB,HLA-DQA1,HLA-DQA2,HLA-DQB1,LY96,FCER<br>1G,FCGR1A,FCGR<br>2A,APLNR,CCL5,S<br>EMA6A,VEGFA,F<br>PR3,NLGN1,CSF1<br>R,IL2RB,CXCL10,<br>AXL,TNFRSF6B,I<br>L20RB,CCND1,IT<br>GB2,NRP1,KISS1R<br>,PXDN,CXCR4,AC<br>KR3,NPTX2,LAG3,<br>IL18BP,EDNRB,T<br>GFA,CD2,CD3D,C<br>D8A,CD14,NETO2,<br>CCNB2,CD86,CD3<br>6,SCARB1,SERPIN<br>E1,ADGRE5 |  |

### PPI network construction

The PPI network of up regulated genes consisted of 6829 nodes and 16639 edges is shown in Fig. 5. When ‘node degree distribution ≥ 150’, ‘betweenness centrality ≥ 0.00567913’, ‘stress centrality ≥ 16321080’, ‘closeness centrality ≥ 0.33036363’ and ‘clustering coefficient = 0’ were set as the cut-off criterion, a total of 12 genes (EGLN3, PCLAF, VCAM1, CANX, SHMT2, IFI16, GAPDH, MS4A7, FGF11, IKBIP, SCD and TRIM9) were selected as hub genes from DEGs and detailed listed of hub genes are given in Table 6. The statistical results and scatter plot for node degree distribution, betweenness centrality, stress centrality, closeness centrality and clustering coefficient are presented in Fig. 6A - 6E. It was found that the pathway and GO enrichment anlysis of 12 hub genes are mainly referred to HIF-1 signaling pathway, extracellular matrix organization, phagosome, dTMP de novo biosynthesis, innate immune system, response to cytokine, pathways in cancer, endoplasmic reticulum part, defense response and adaptive immune system. The PPI network of down regulated genes consisted of 5108 nodes and 9299 edges is shown in Fig. 7. When ‘node degree distribution ≥ 150’, ‘betweenness centrality ≥ 0.01687101’, ‘stress centrality ≥ 8184666’, ‘closeness centrality ≥ 0.30386246’ and ‘clustering coefficient = 0’ were set as the cut-off criterion, a total of 12 genes (CDH1, ERBB2, ATP1A1, ALB, NEDD4L, PHGDH, RTN4, ACOT11, ANKRD9, VTCN1, SLC22A7 and MST1L) were selected as hub genes from DEGs and detailed listed of hub genes are given in Table 6. The statistical results and scatter plot for node degree distribution, betweenness centrality, stress centrality, closeness centrality and clustering coefficient are presented in Fig. 8A - 8E. It was found that the pathway and GO enrichment anlysis of 12 hub genes are mainly referred to Rap1 signaling pathway, tight junction, aldosterone-regulated sodium reabsorption, FOXA2 and FOXA3 transcription factor networks, ion transport, metabolic pathways, p75(NTR)-mediated signaling, metabolism of lipids and lipoproteins, plasma membrane bounded cell projection part, signaling receptor binding, transmembrane transport of small molecules and ensemble of genes encoding extracellular matrix and extracellular matrix-associated proteins.

**Fig. 5.**
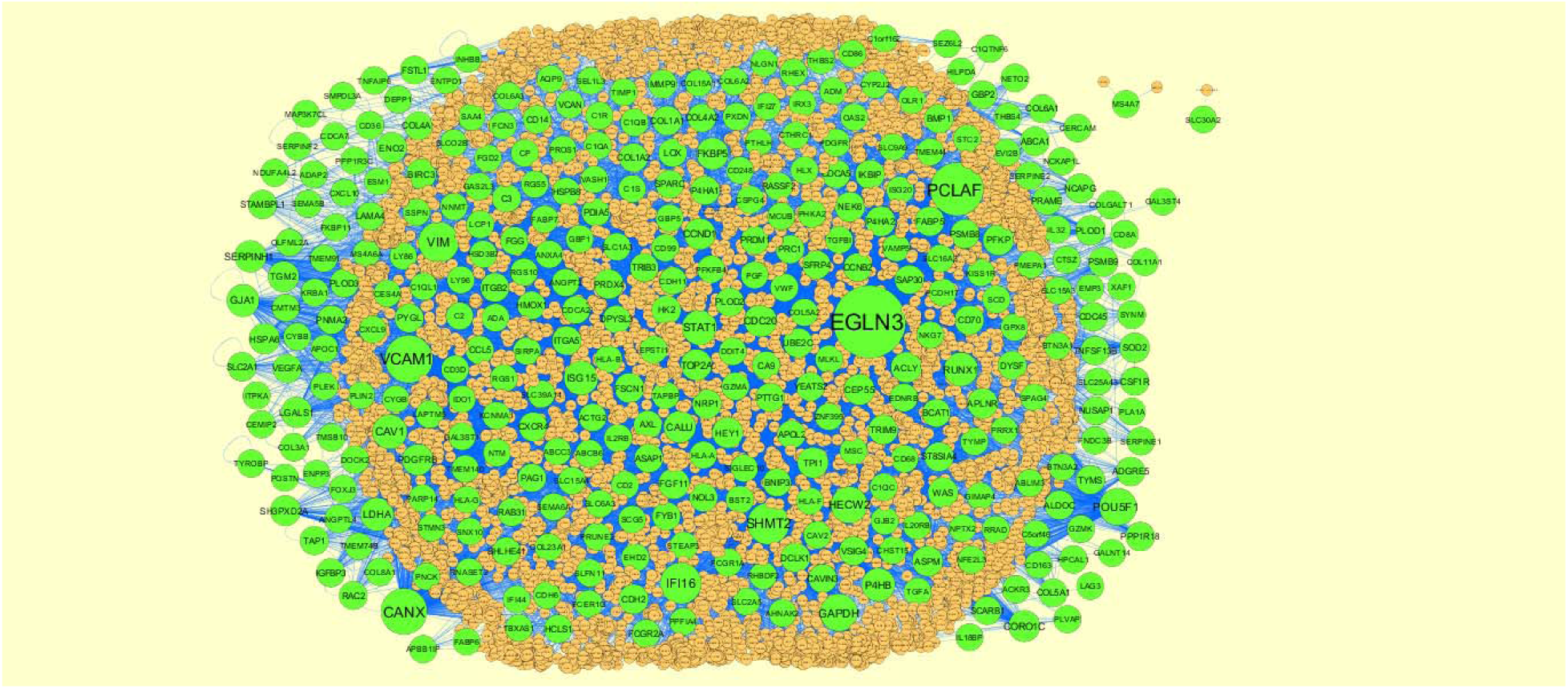
Protein–protein interaction network of up regulated genes. Green nodes denotes up regulated genes.

**Fig. 6.**
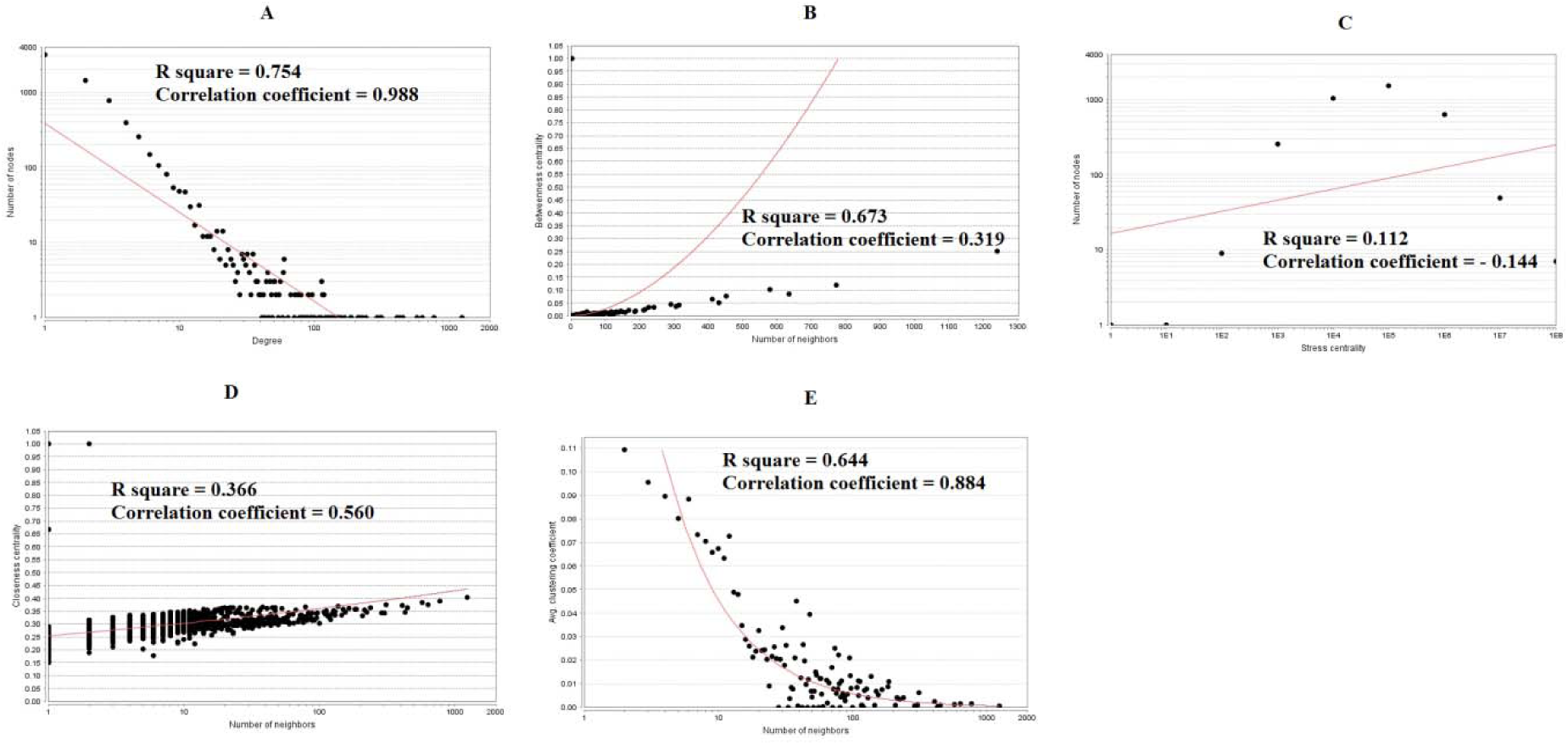
Scatter plot for up regulated genes. (A- Node degree; B- Betweenness centrality; C- Stress centrality; D- Closeness centrality; E- Clustering coefficient)

**Fig. 7.**
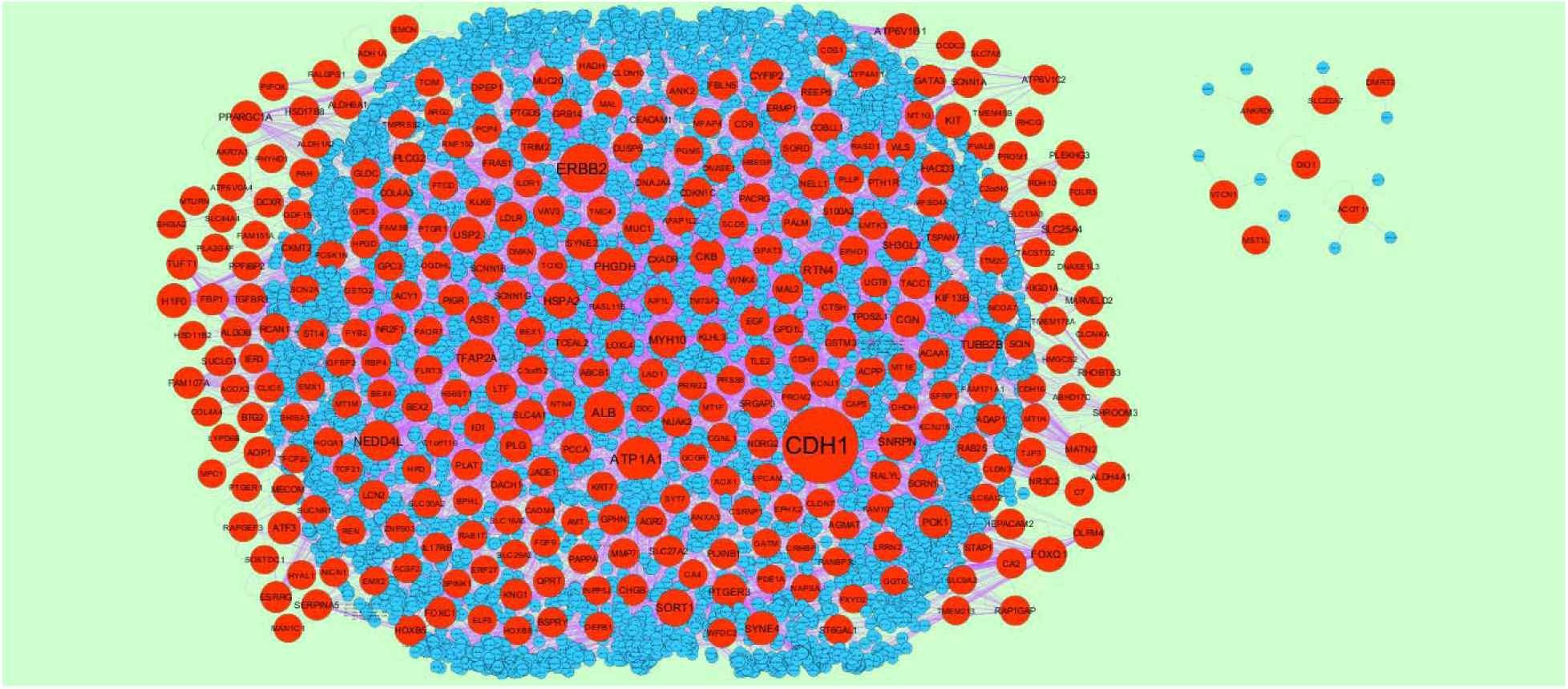
Protein–protein interaction network of down regulated genes. Red nodes denotes down regulated genes.

**Fig. 8.**
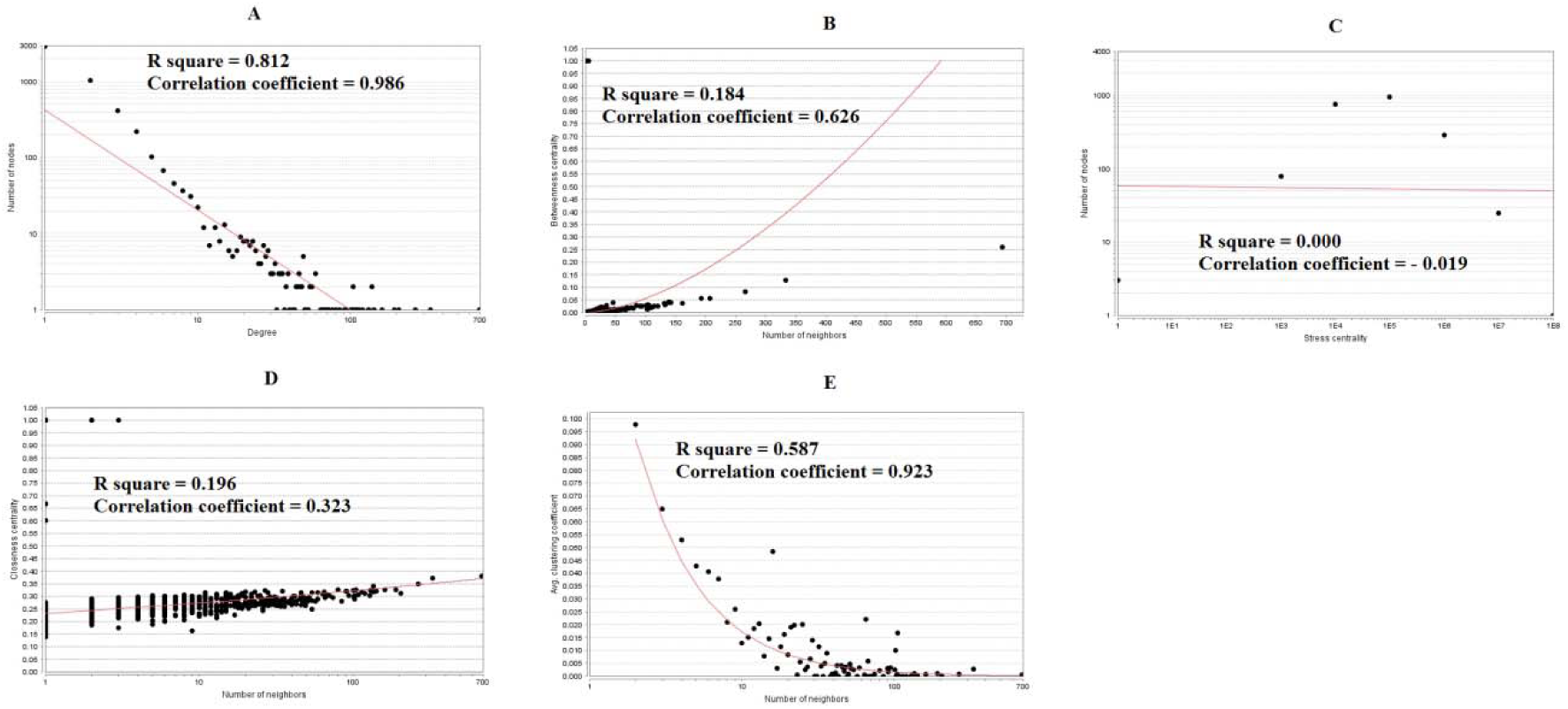
Scatter plot for down regulated genes. (A- Node degree; B- Betweenness centrality; C- Stress centrality; D- Closeness centrality; E- Clustering coefficient)

**Table 6.**
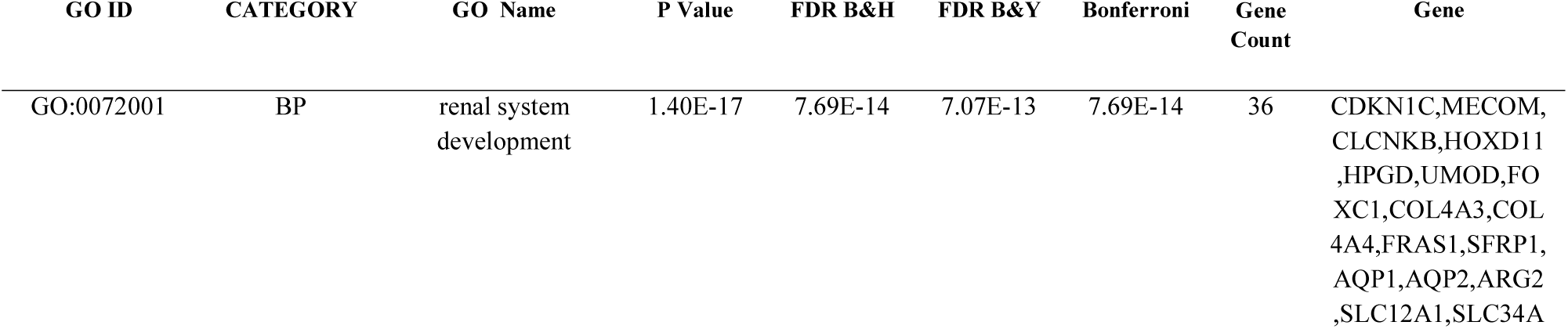
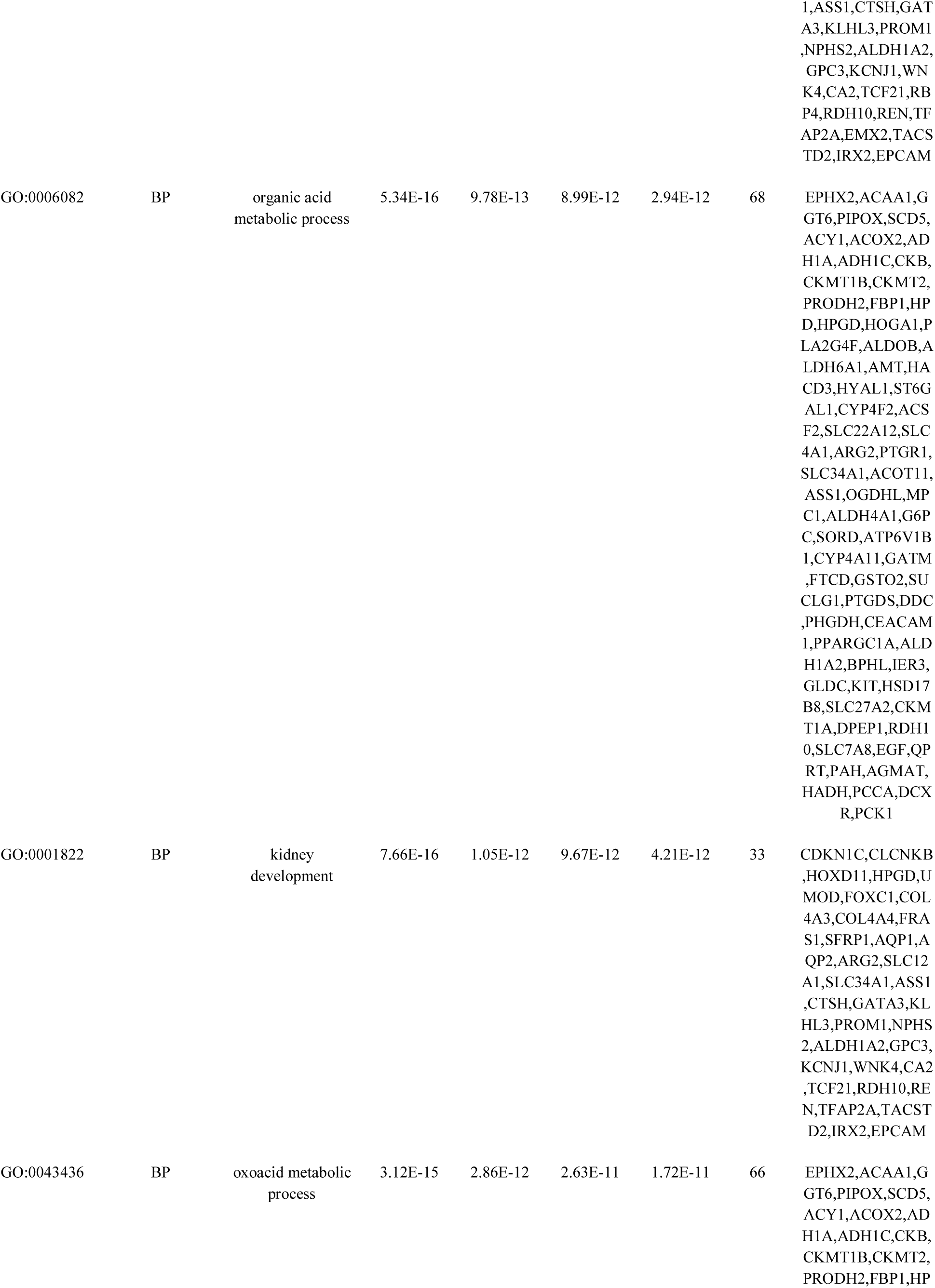
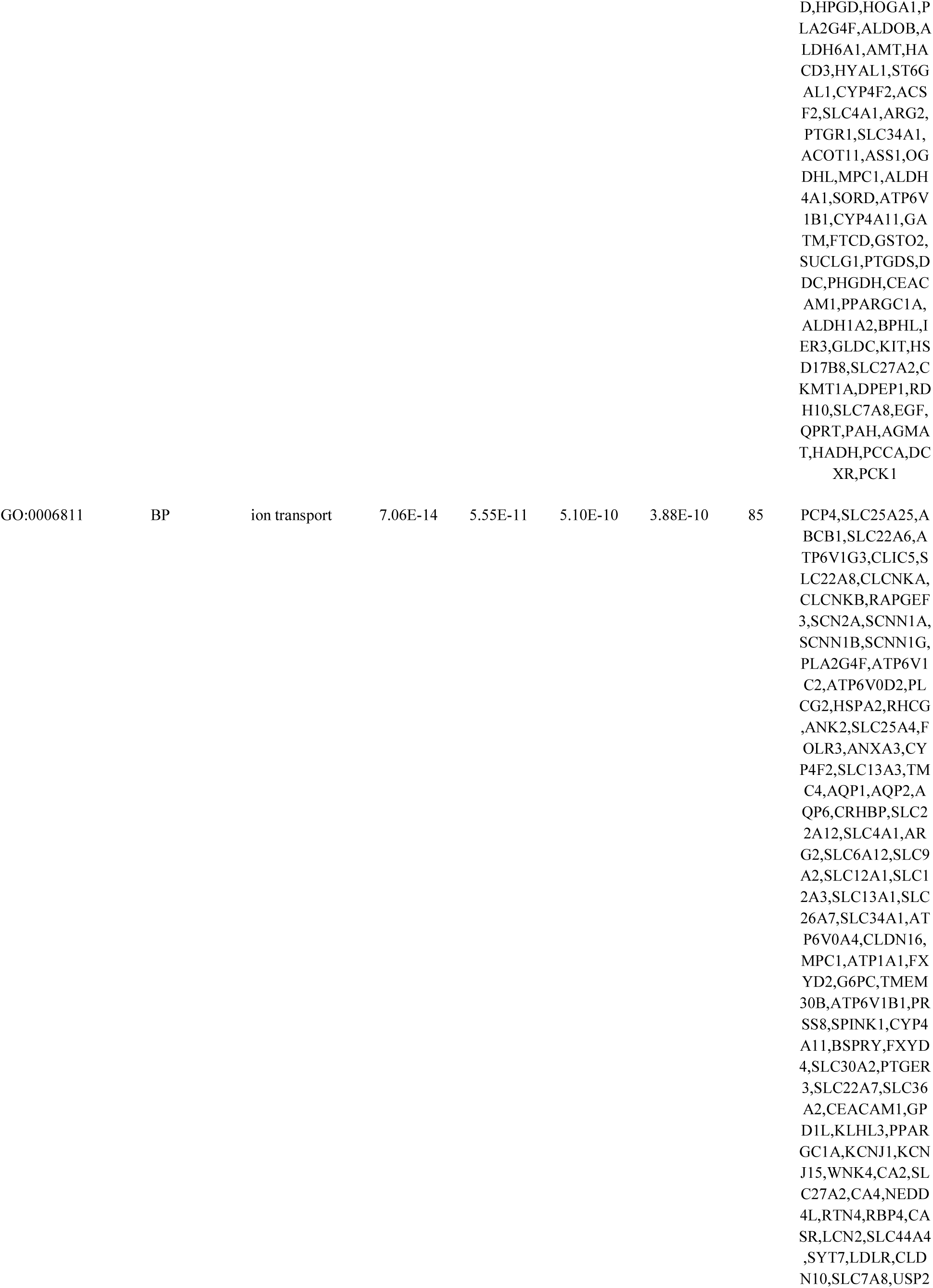
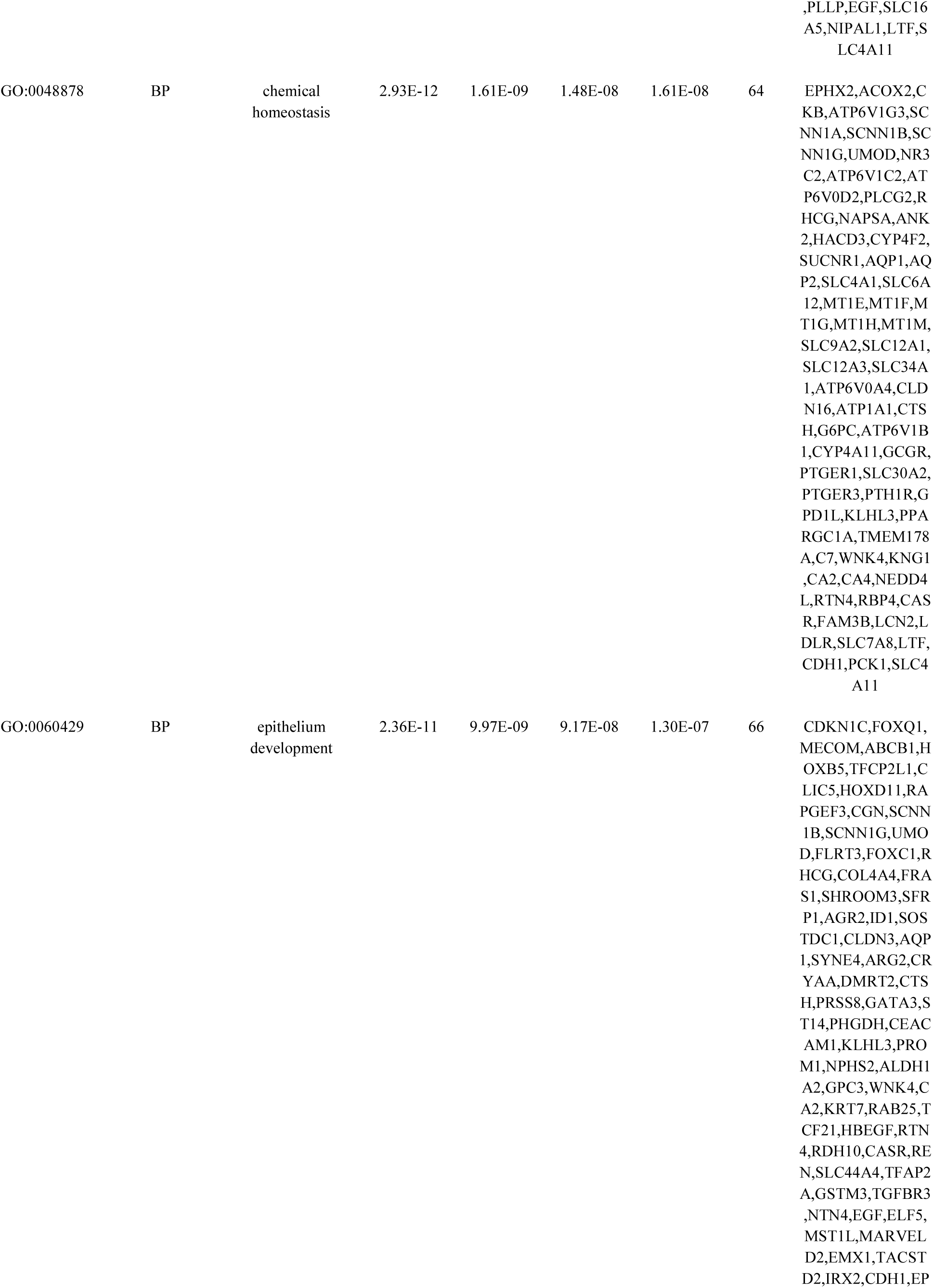
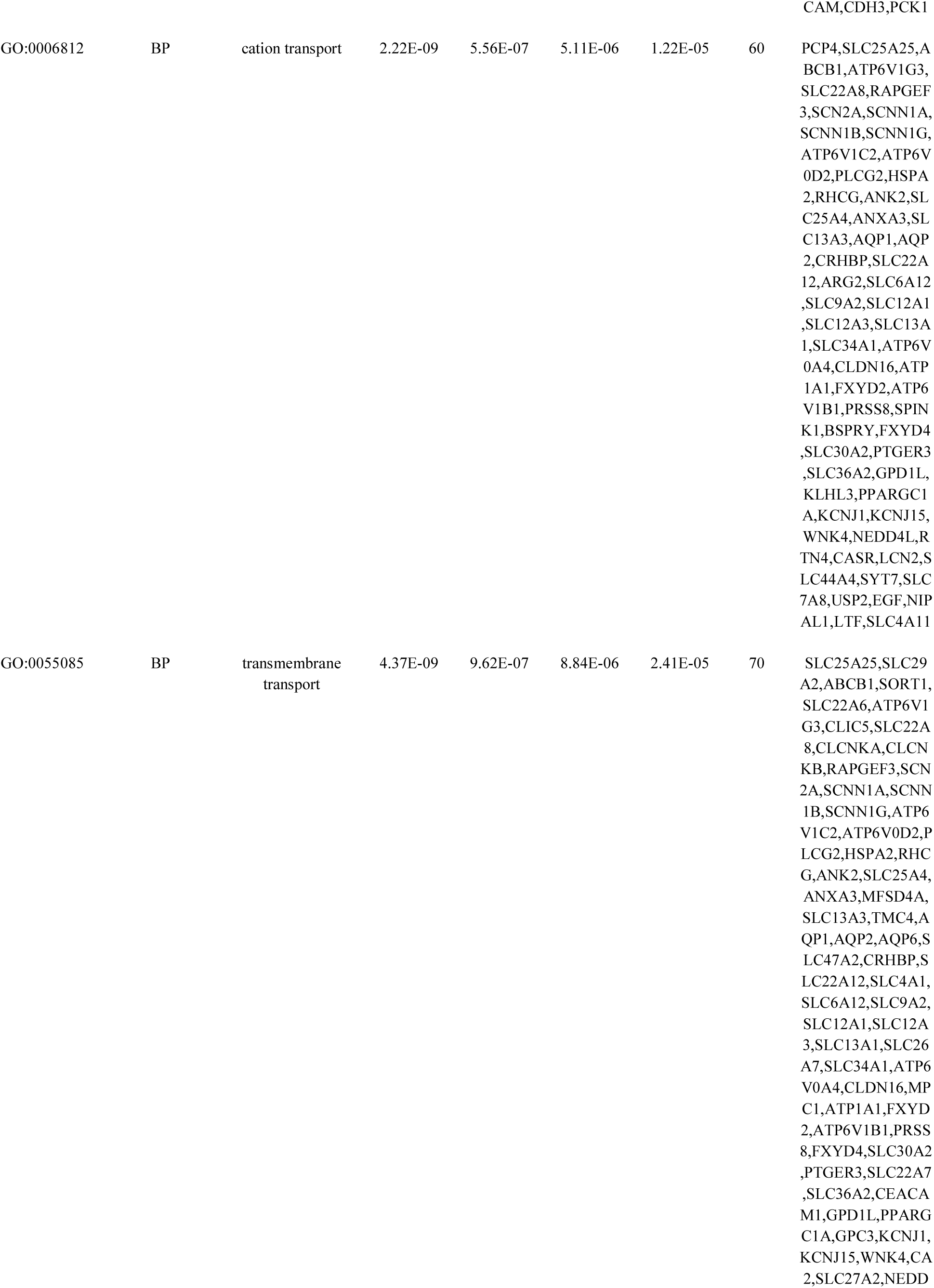
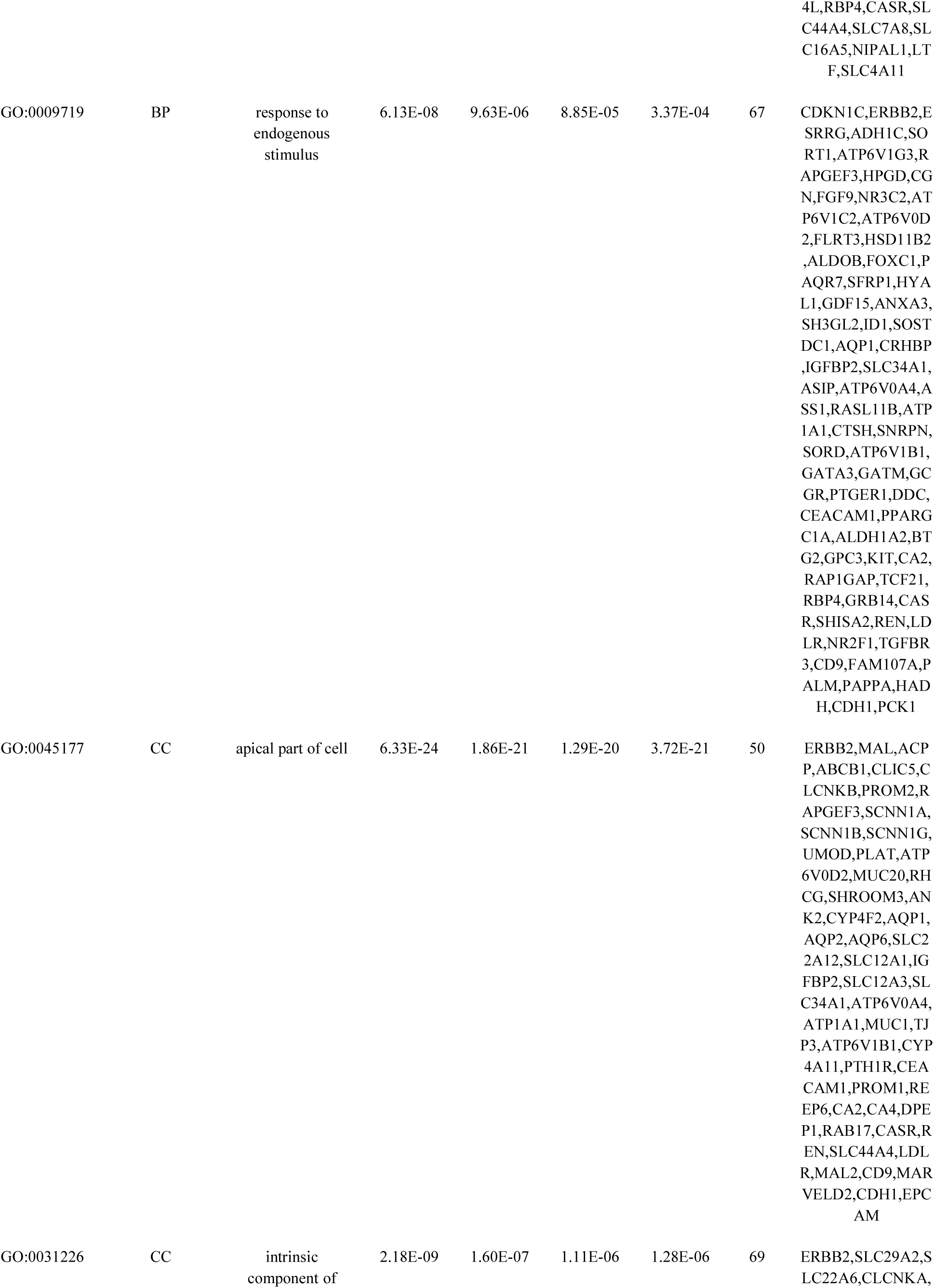
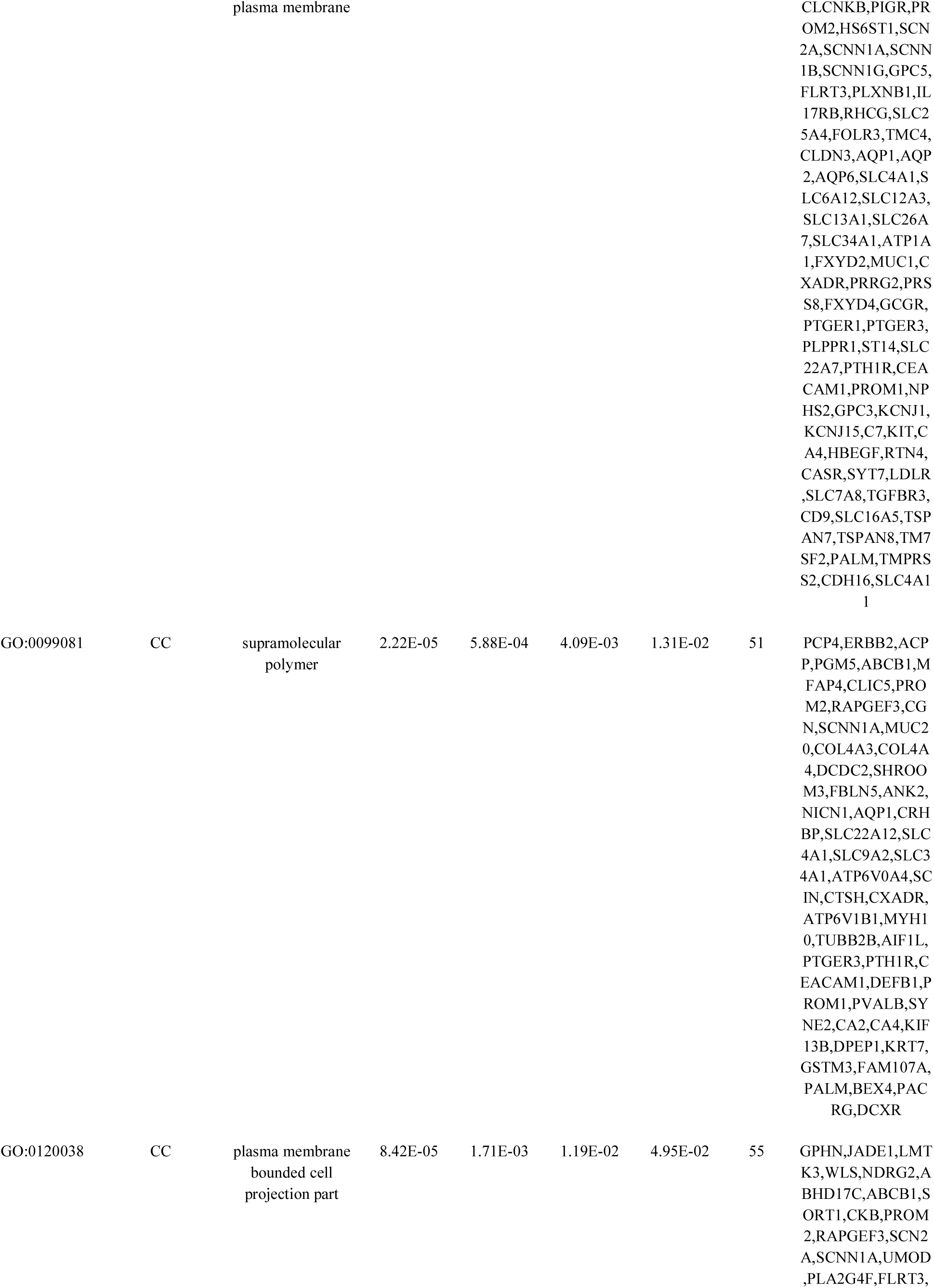
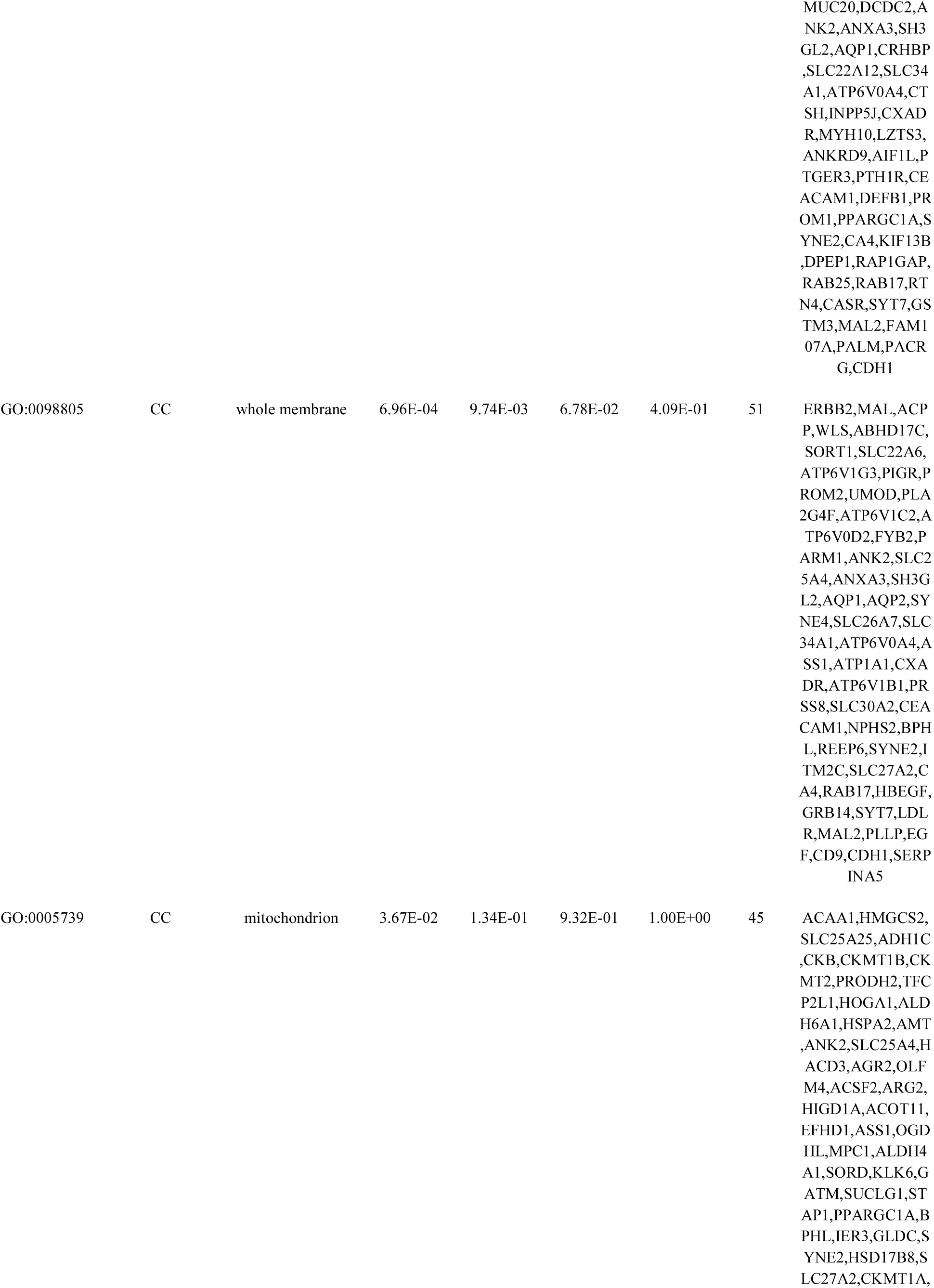
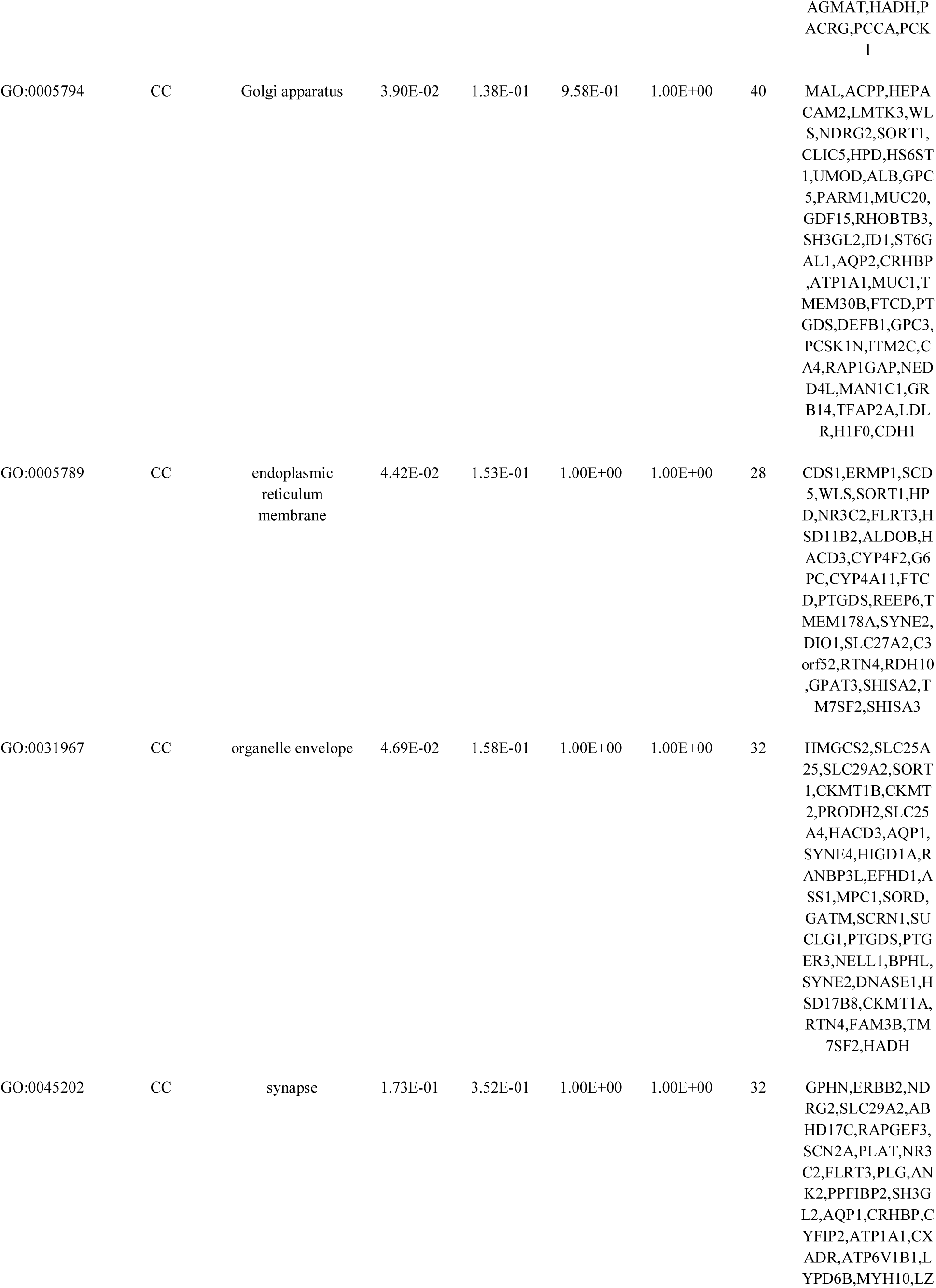
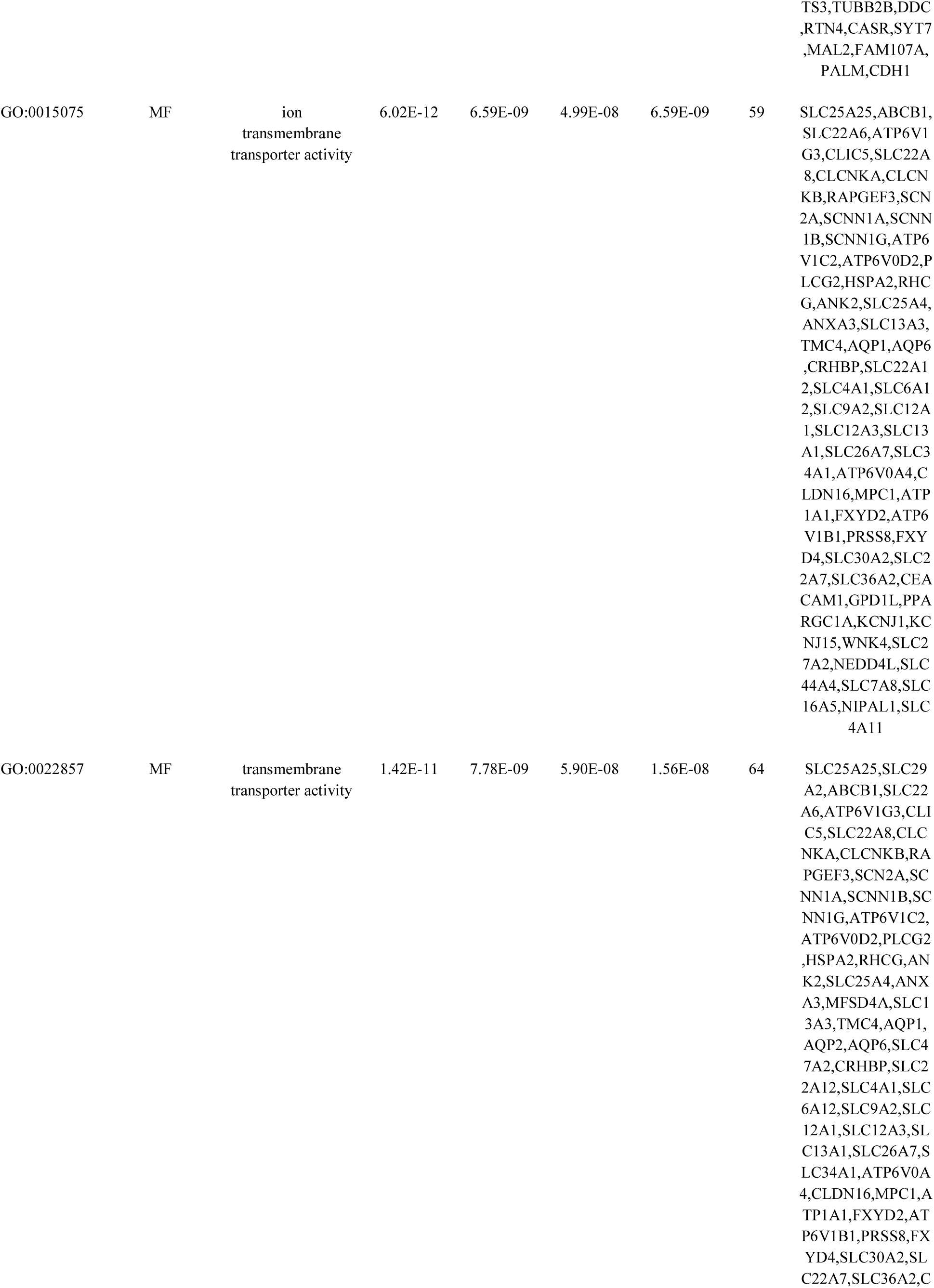
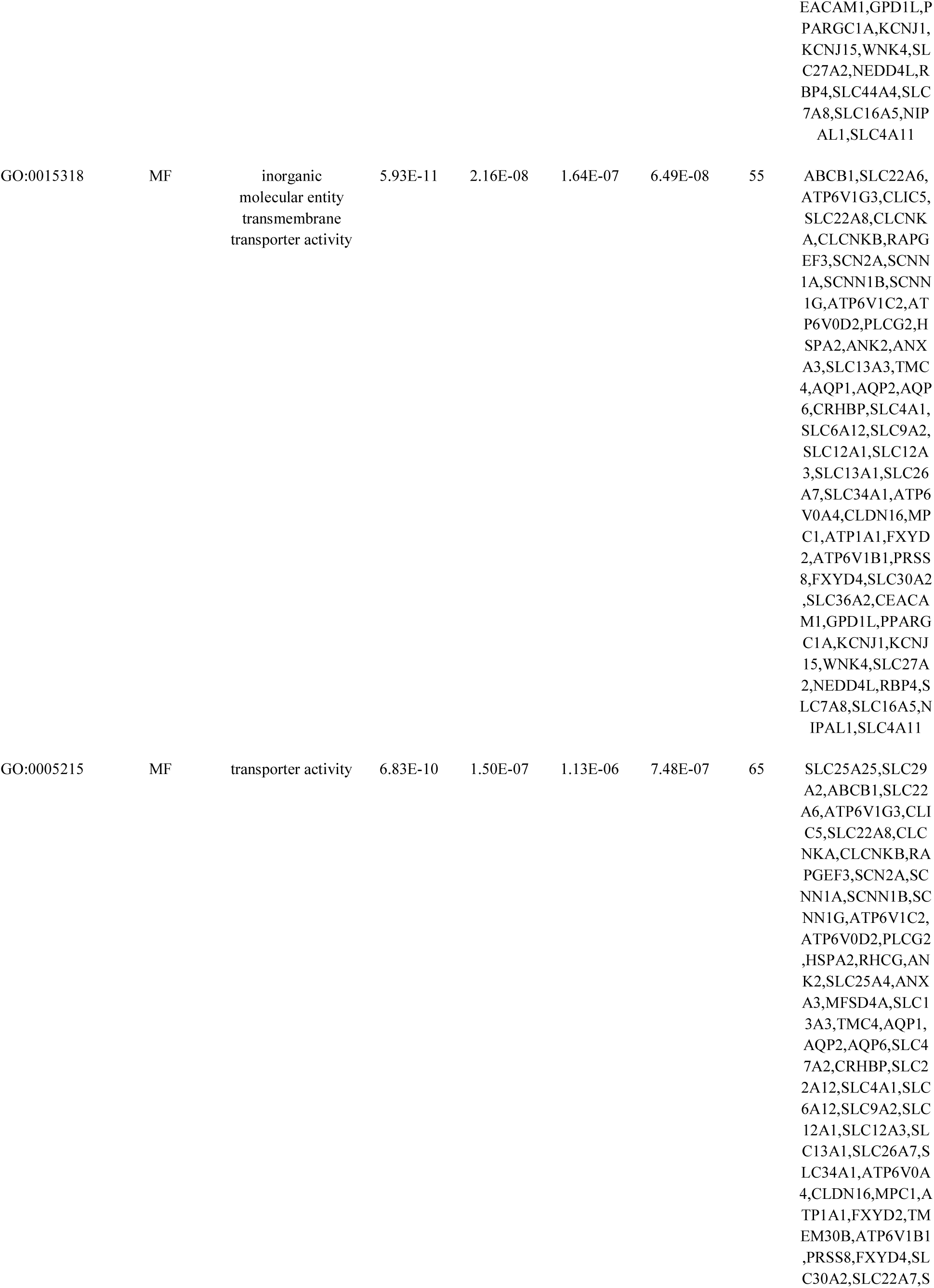
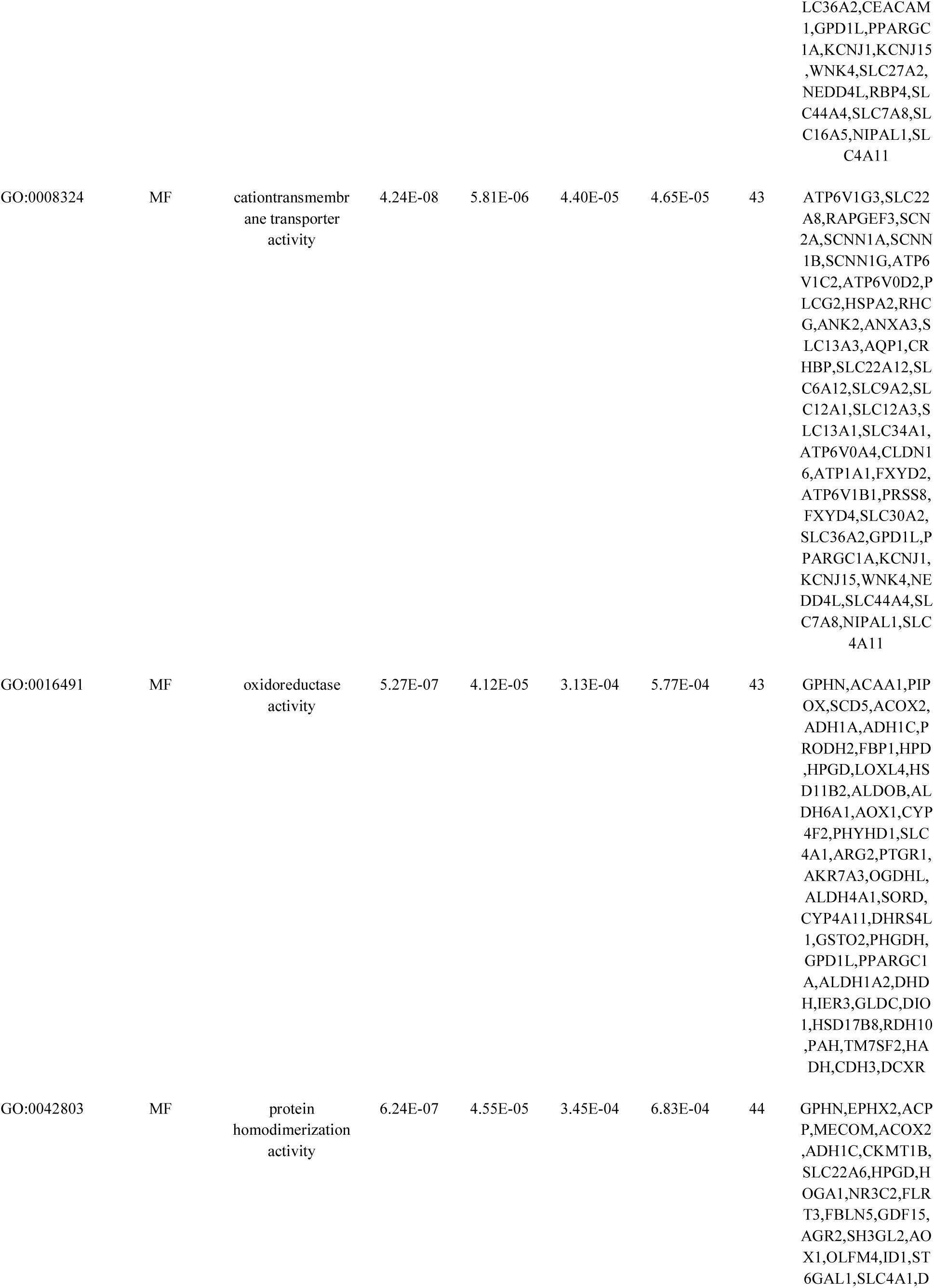
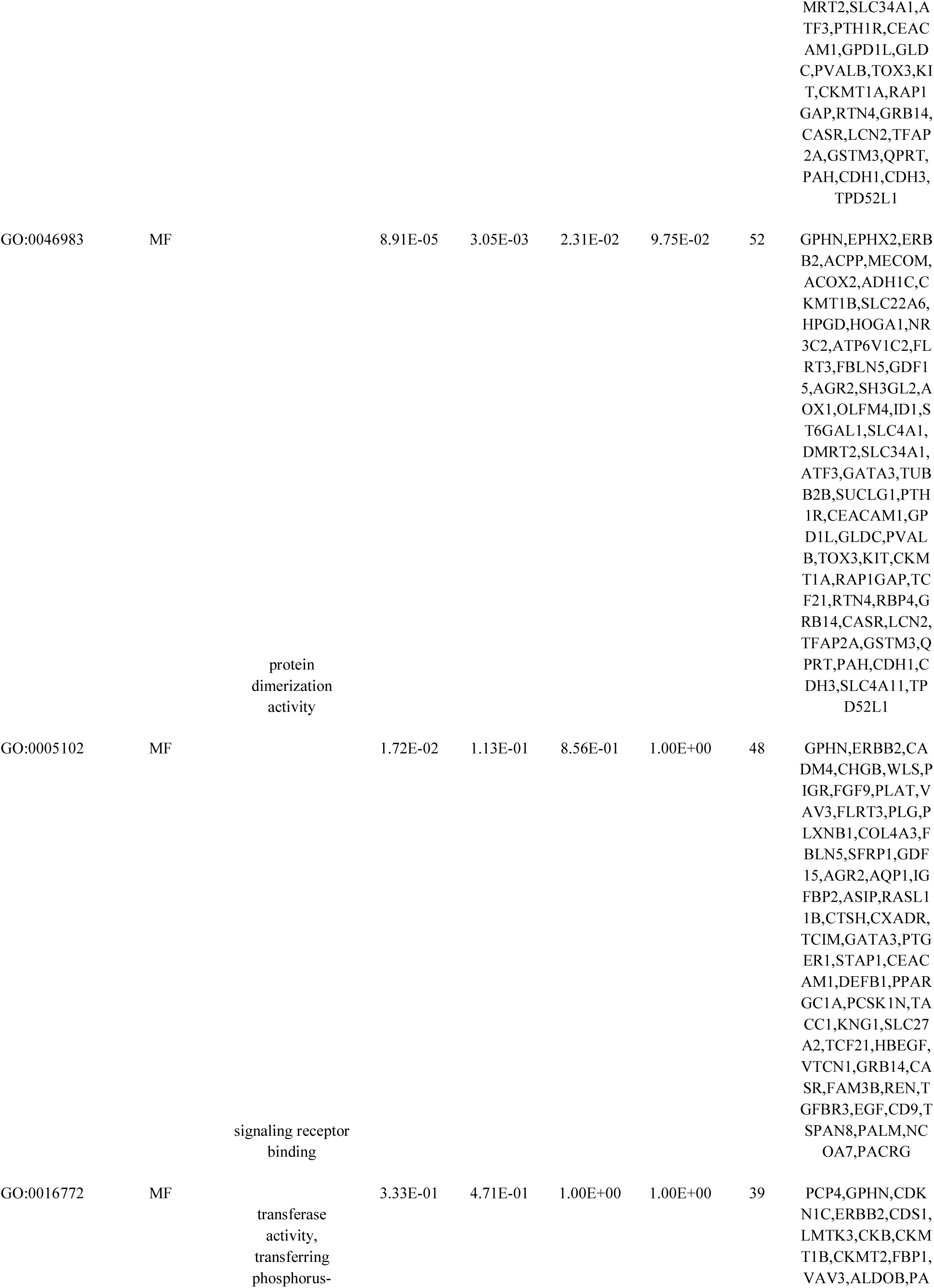
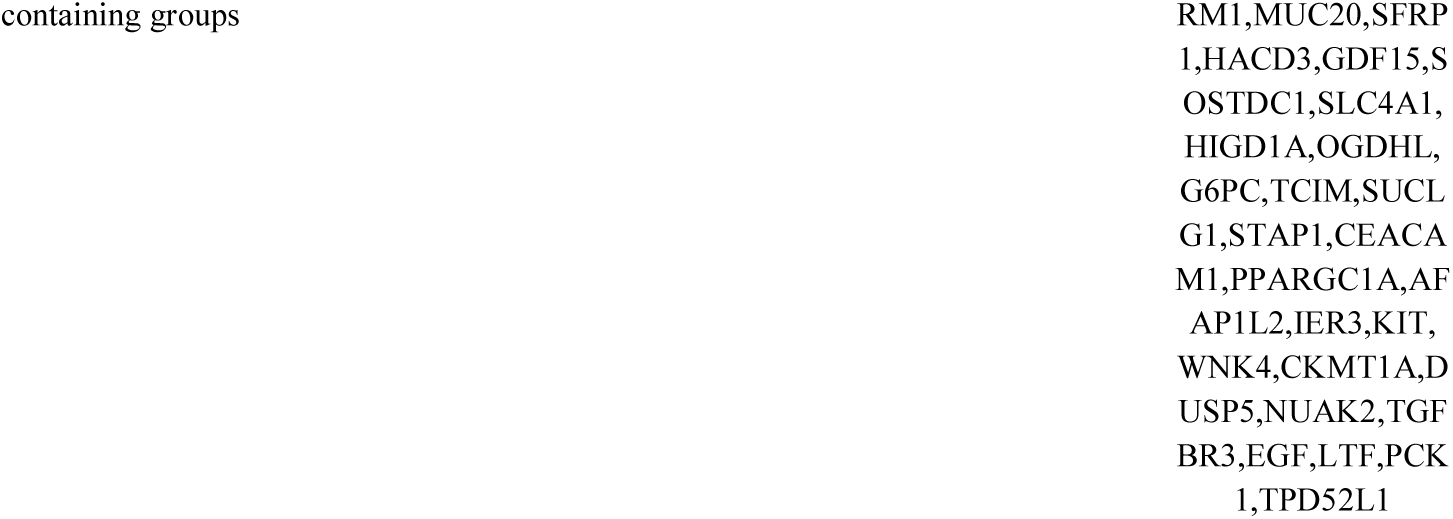
The enriched GO terms of the down-regulated differentially expressed genes

### Module analysis

Total 2083 modules were obtained from the PPI network for up regulated genes based on the PEWCC1 analysis in Cytoscape. We chose the foure most significant modules for further analysis (Fig.9). Module 8 consisted of 41 nodes and 108 edges, module 11 consisted of 34 nodes and 90 edges, module 21 consisted of 26 nodes and 62 edges, and module 32 consisted of 18 nodes and 64 edges. Pathway and GO enrichment analysis showed that GAPDH, VIM, STAT1, COL6A1, ENO2, SERPINH1, CAV1 and SPARC were included in module 8, module 11, module 21 and module 32, and participated in HIF-1 signaling pathway, aurora B signaling, pathways in cancer, focal adhesion, cell surface, extracellular matrix, response to cytokine and response to other organism. Total 932 modules were obtained from the PPI network for down regulated genes based on the PEWCC1 analysis in Cytoscape. We chose the foure most significant modules for further analysis (Fig.10). Module 3 consisted of 25 nodes and 65 edges, module 5 consisted of 22 nodes and 47 edges, module 6 consisted of 17 nodes and 46 edges, and module 8 consisted of 15 nodes and 31 edges. Pathway and GO enrichment analysis showed that ERBB2, CDH1, NEDD4L, KLK6, PLCG2, RTN4, SCNN1A and COL4A3 were included in module 3, module 5, module 6 and module 8, and participated in tight junction, rap1 signaling pathway, aldosterone-regulated sodium reabsorption, alpha-synuclein signaling, ion transport, chemical homeostasis, cation transport and renal system development.

**Fig. 9.**
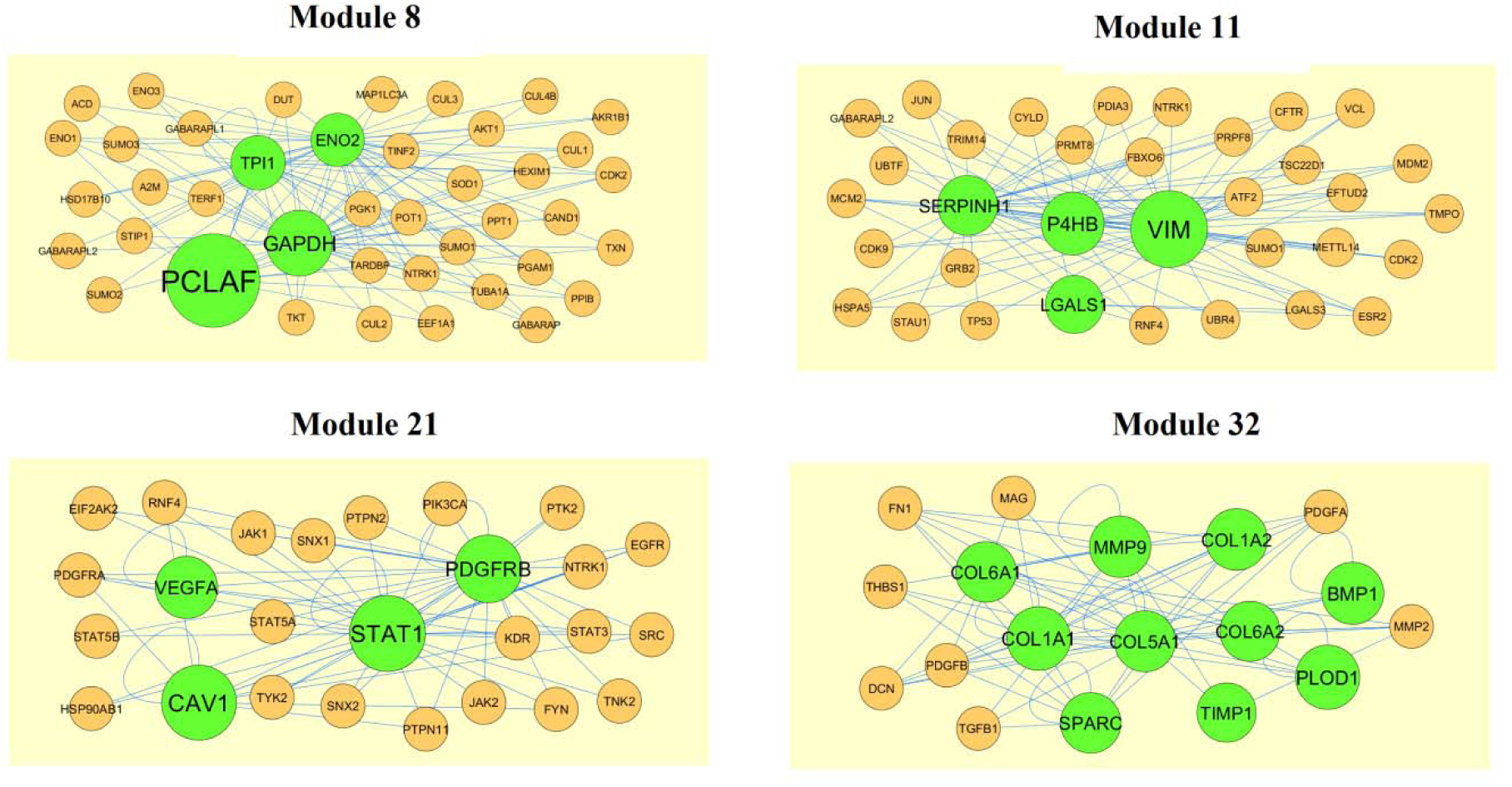
Modules in PPI network. The green nodes denote the up regulated genes

**Fig. 10.**
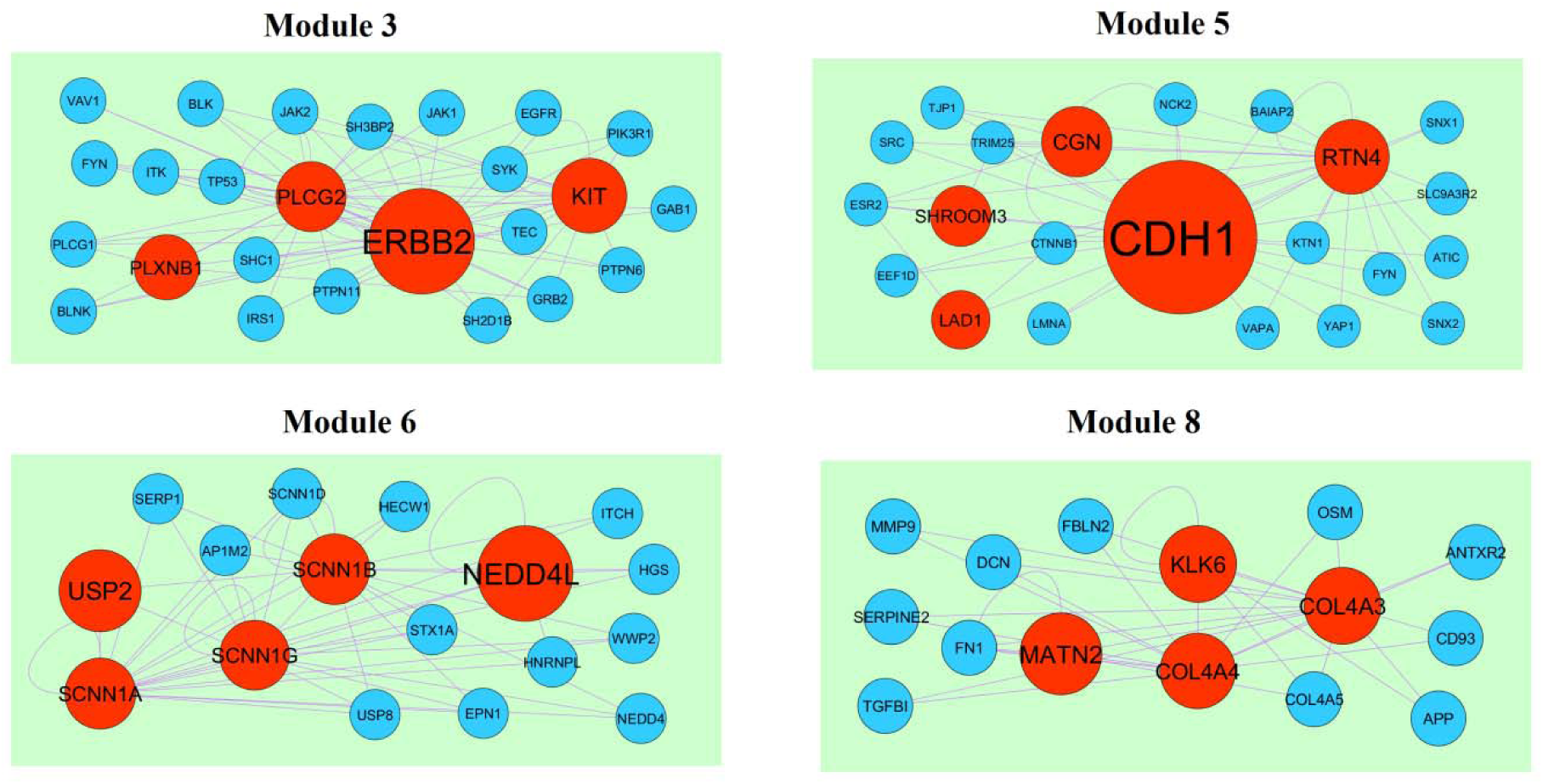
Modules in PPI network. The red nodes denote the down regulated genes

### Construction of target genes - miRNA regulatory network

Most of miRNAs were responsible for progression of CCRCC [Liu et al 2010]. In order to investigate the regulatory network of target genes - miRNA for CCRCC, DIANA-TarBase and miRTarBase were utilized to obtain miRNAs for the top 5 up regulated and down regulated genes. The up regulated genes and miRNAs is shown in Fig. 11. Specifically, that SOD2 regulates 257 miRNAs (ex; hsa-mir-4424), CCND1 regulates197 miRNAs (ex; hsa-mir-5009-3p), SCD regulates 167 miRNAs (ex; hsa-mir-3671), VEGFA regulates 108 miRNAs (ex; hsa-mir-4497) and SH3PXD2A regulates 105 miRNAs (ex; hsa-mir-6818-5p) are listed in Table 7. The pathway and GO enrichment analysis reveled that up regulated targate genes were enriched in reactive oxygen species degradation, Focal adhesion, defense response, regulation of immune system process and molecular function regulator. Similarly, down regulated genes and miRNAs is shown in Fig. 12. Specifically, BTG2 regulates 227 miRNAs (ex; hsa-mir-4319), VAV3 regulates164 miRNAs (ex; hsa-mir-5001-5p), LDLR regulates 123 miRNAs (ex; hsa-mir-3121-3p), FOXC1 regulates 103 miRNAs (ex; hsa-mir-5688) and SYT7 regulates 75 miRNAs (ex; hsa-mir-4270) are listed in Table 7. The pathway and GO enrichment analysis reveled that down regulated targate genes were enriched in response to endogenous stimulus, leukocyte transendothelial migration, metabolism of lipids and lipoproteins, renal system development and insulin secretion pathway.

**Fig. 11.**
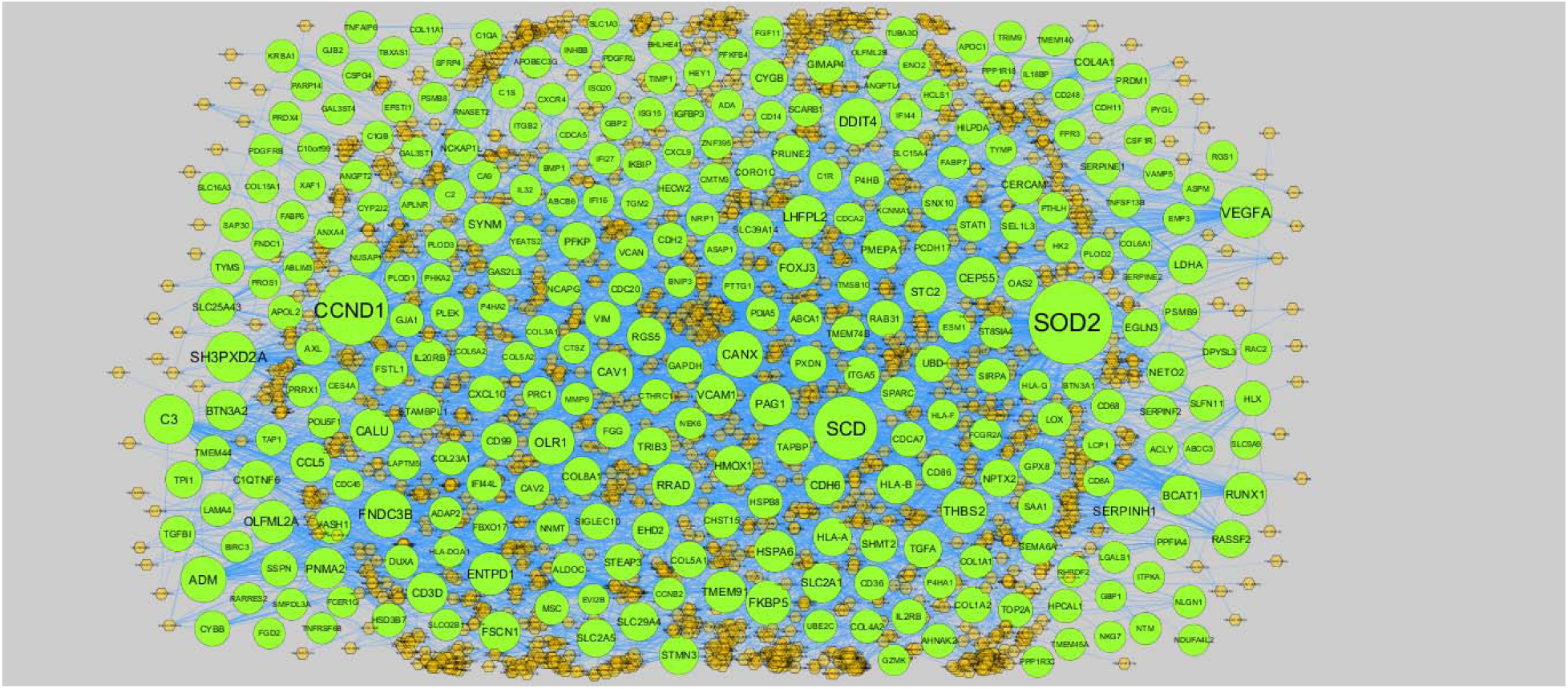
The network of up regulated target gene and their related miRNAs. The green circles nodes are the up regulated target gene, and orange diamond nodes are the miRNAs

**Fig. 12.**
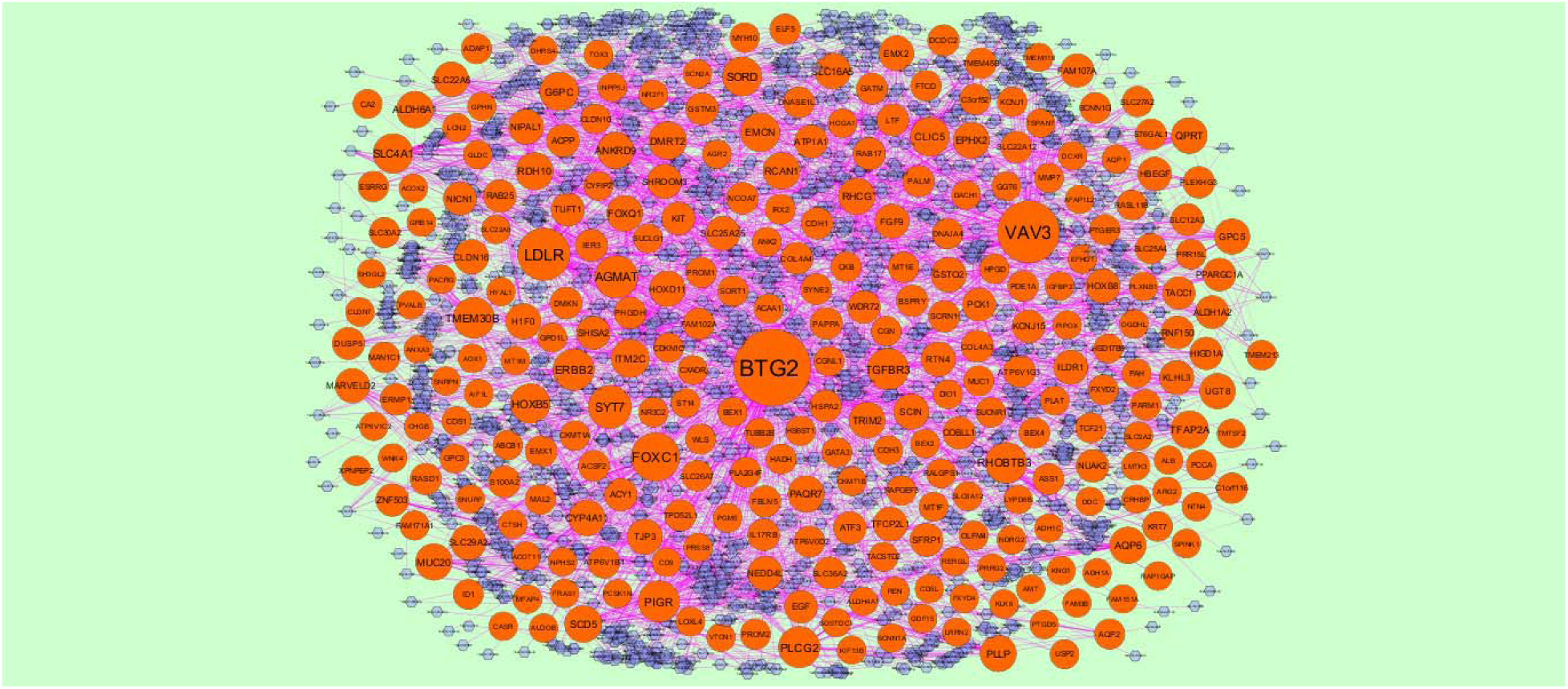
The network of down regulated target gene and their related miRNAs. The red circles nodes are the down regulated target gene, and blue diamond nodes are the miRNAs

**Table 7.**
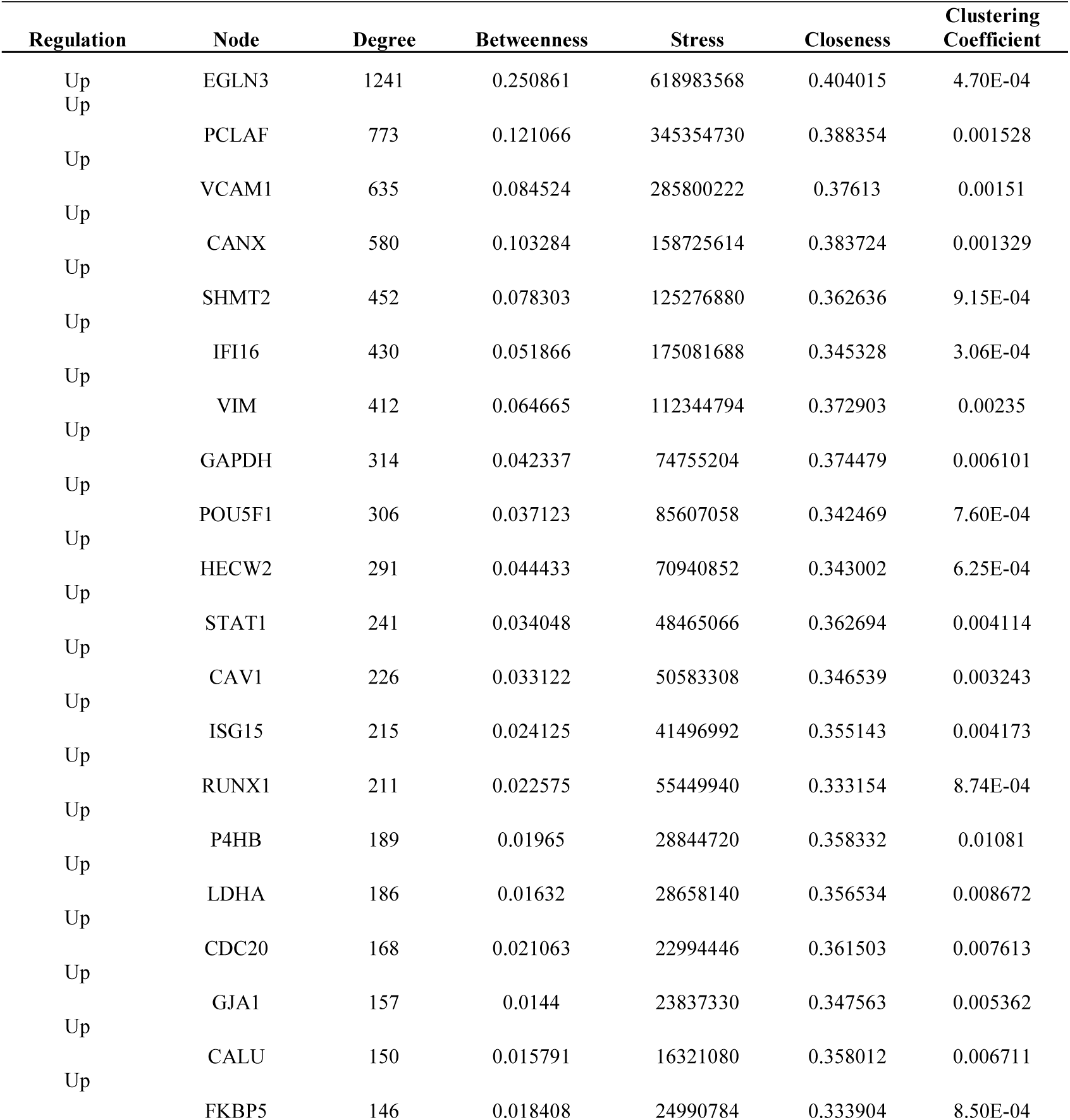
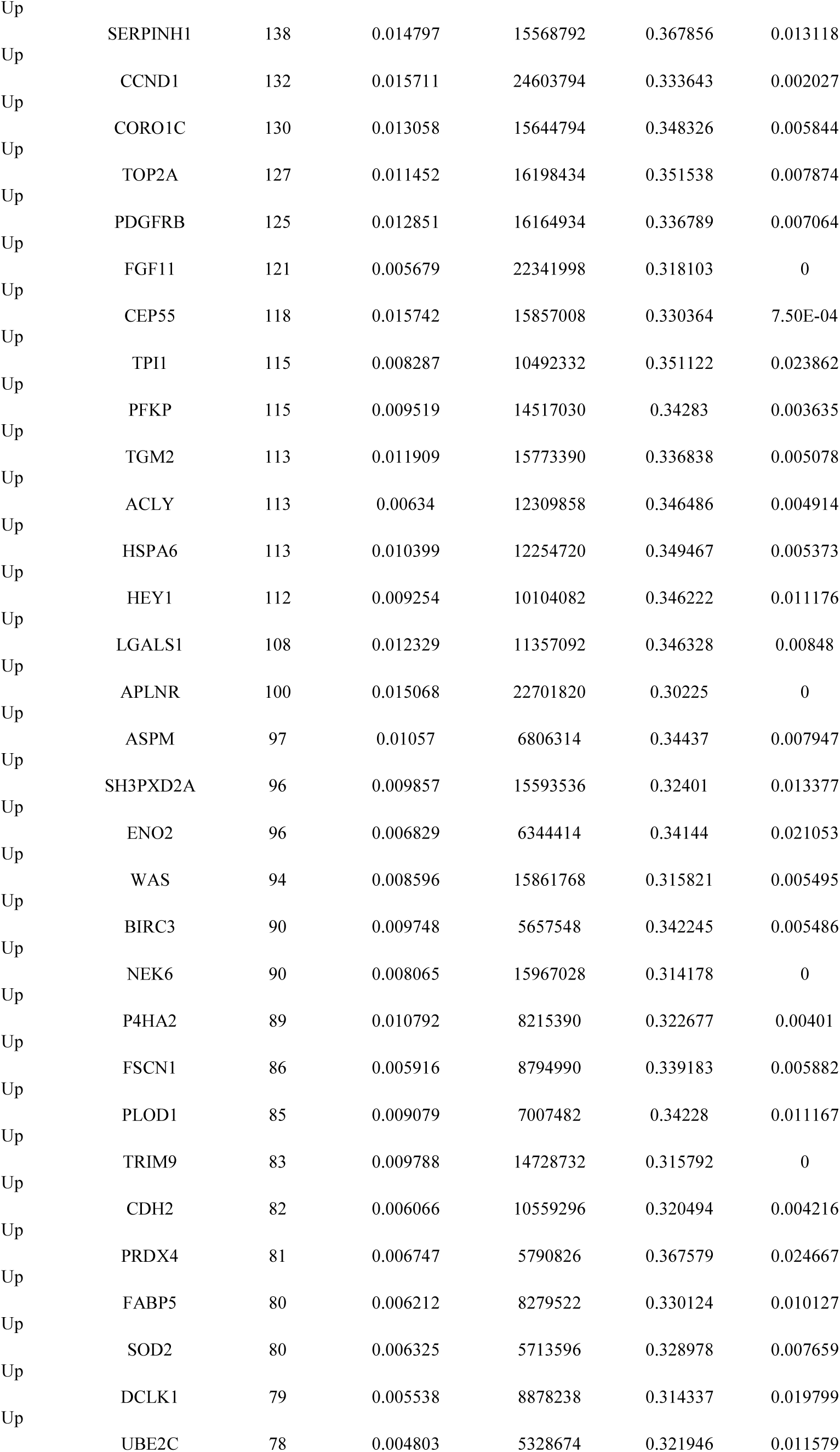
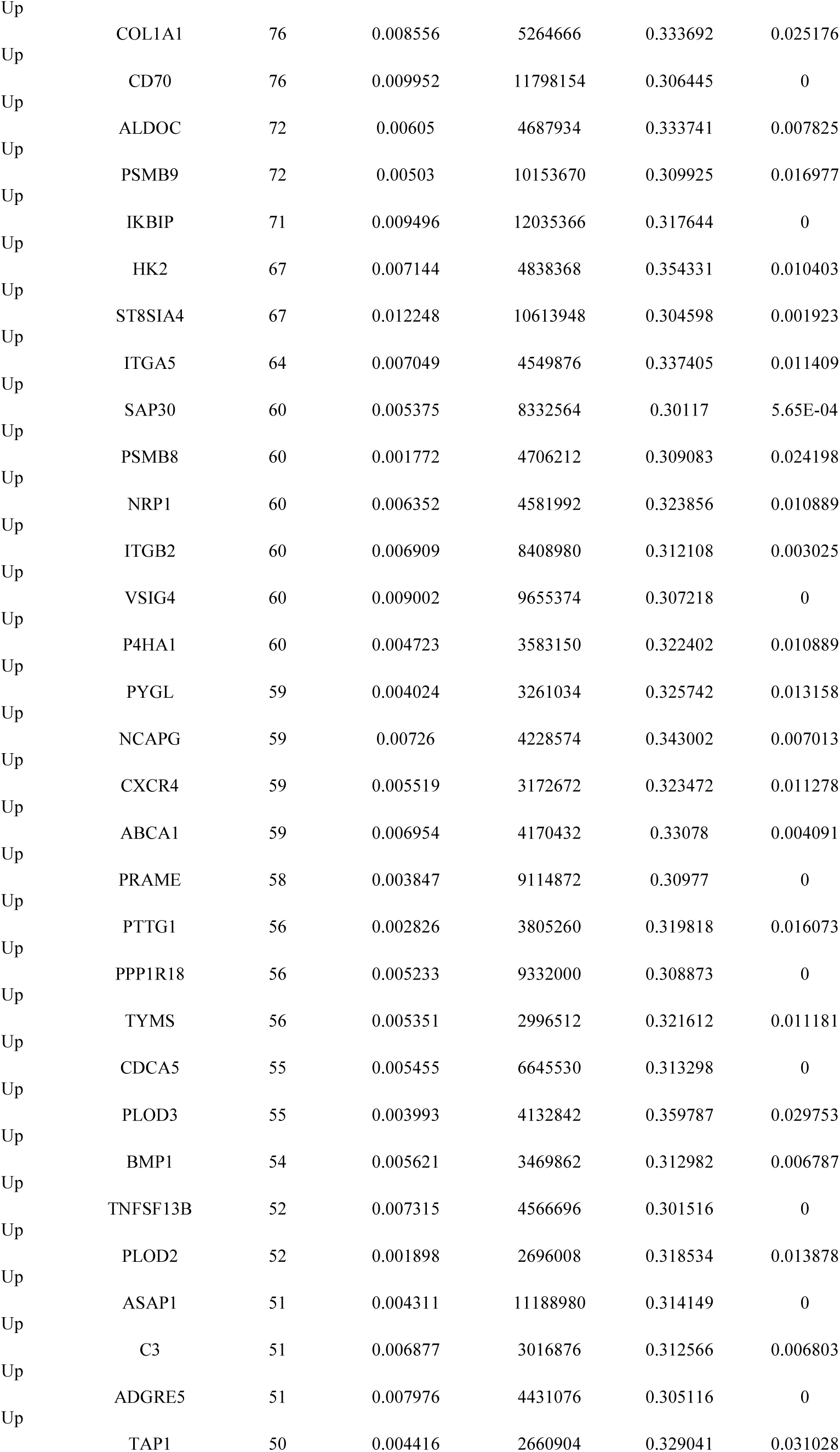
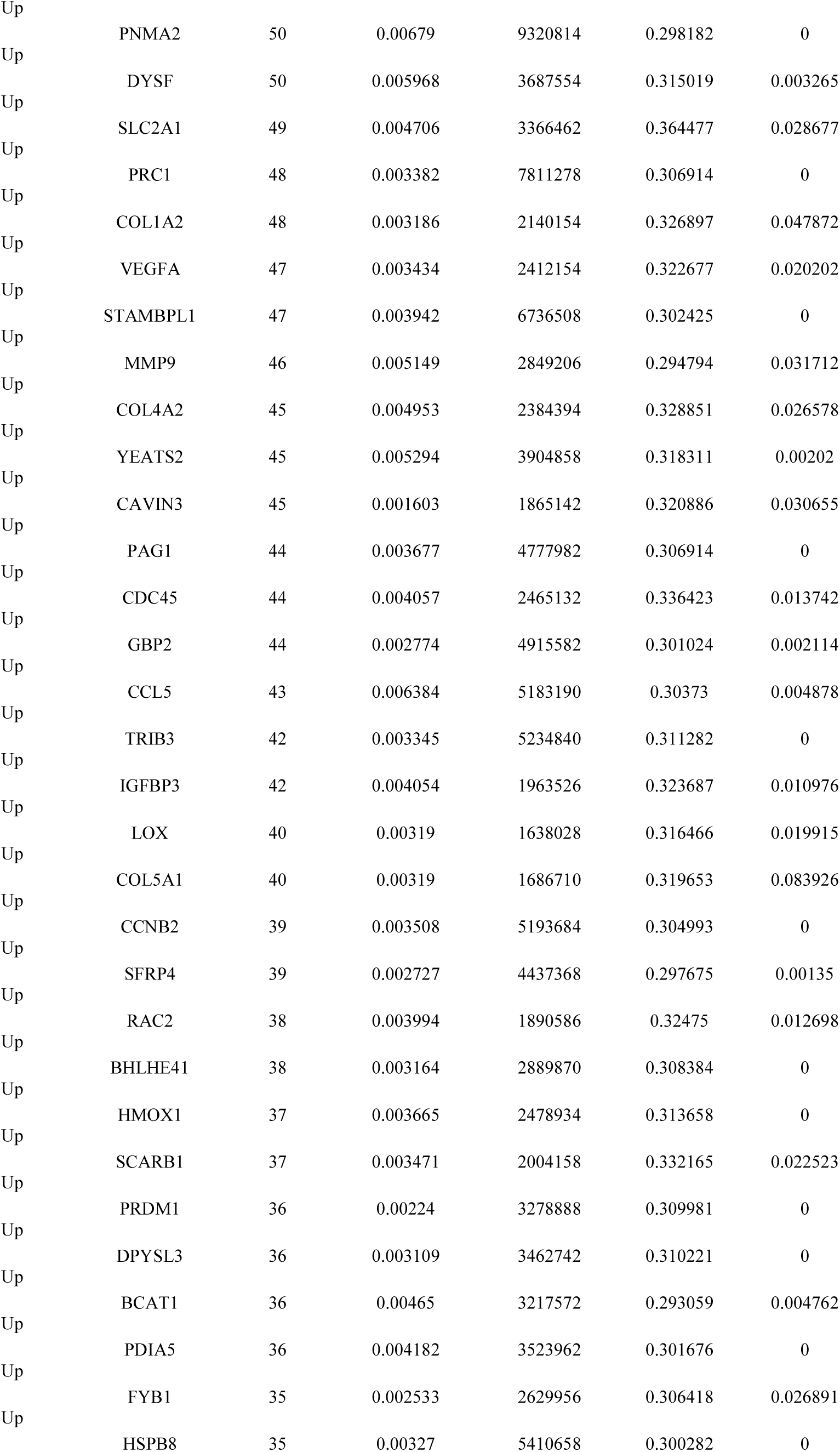
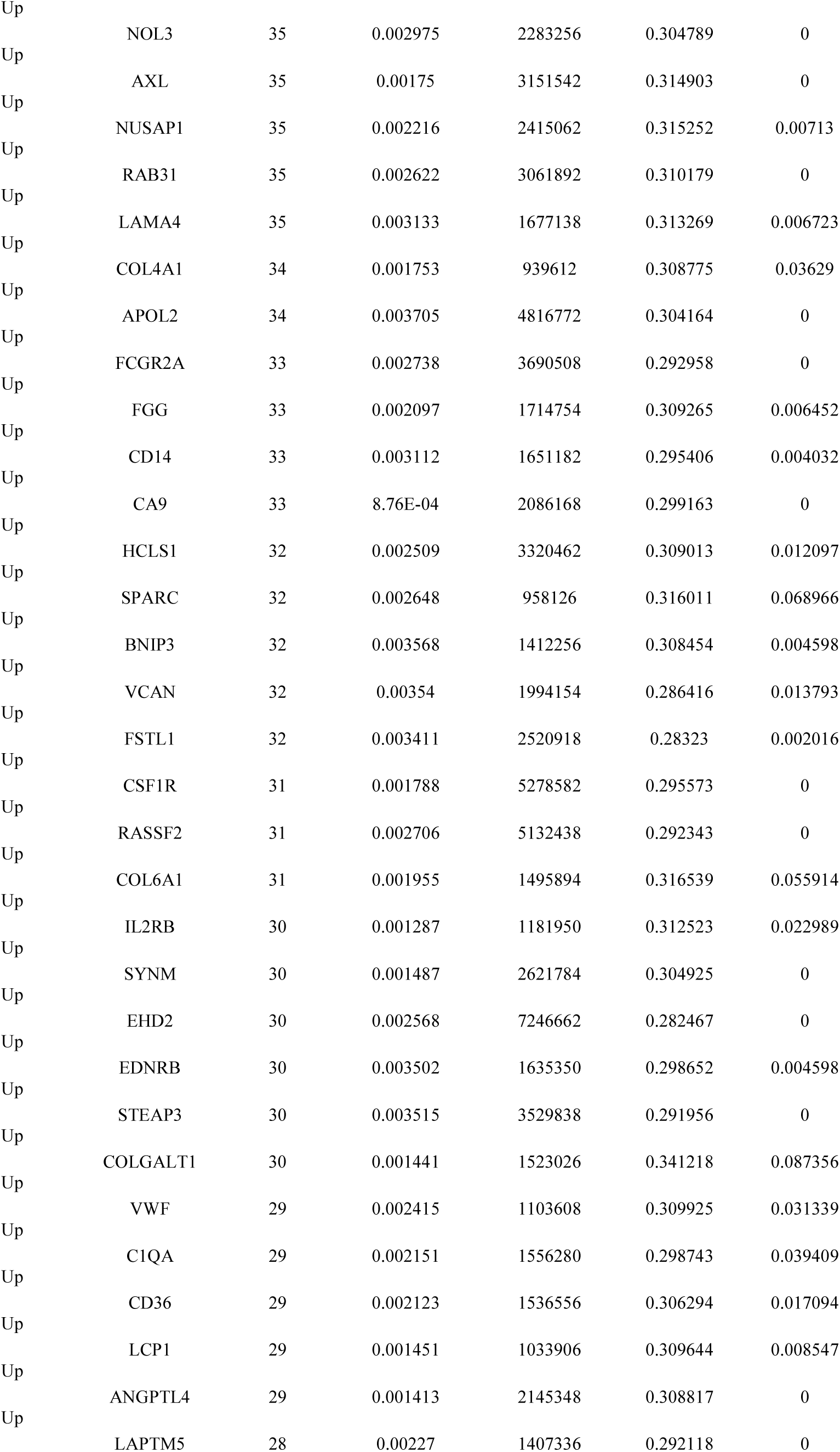
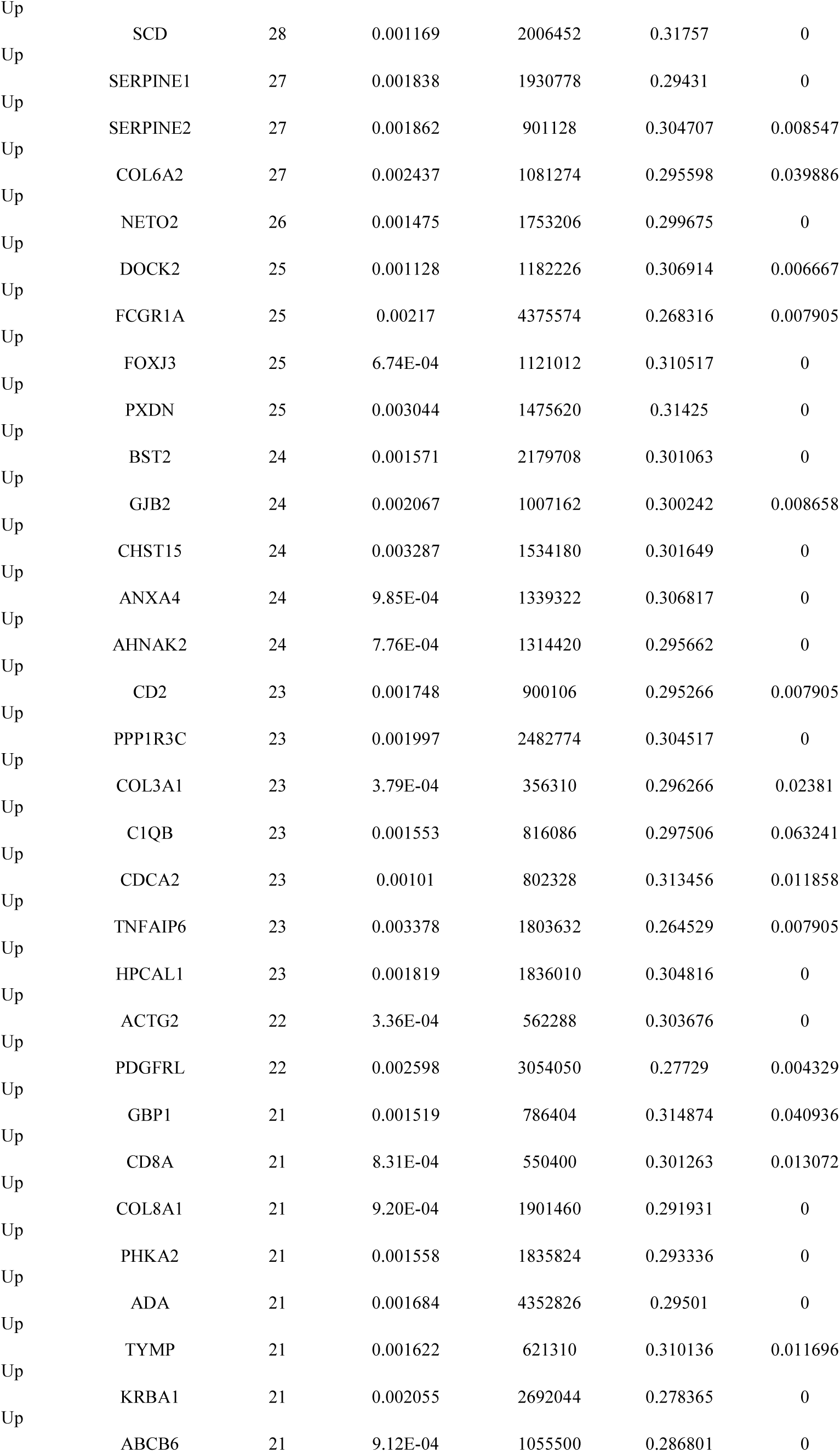
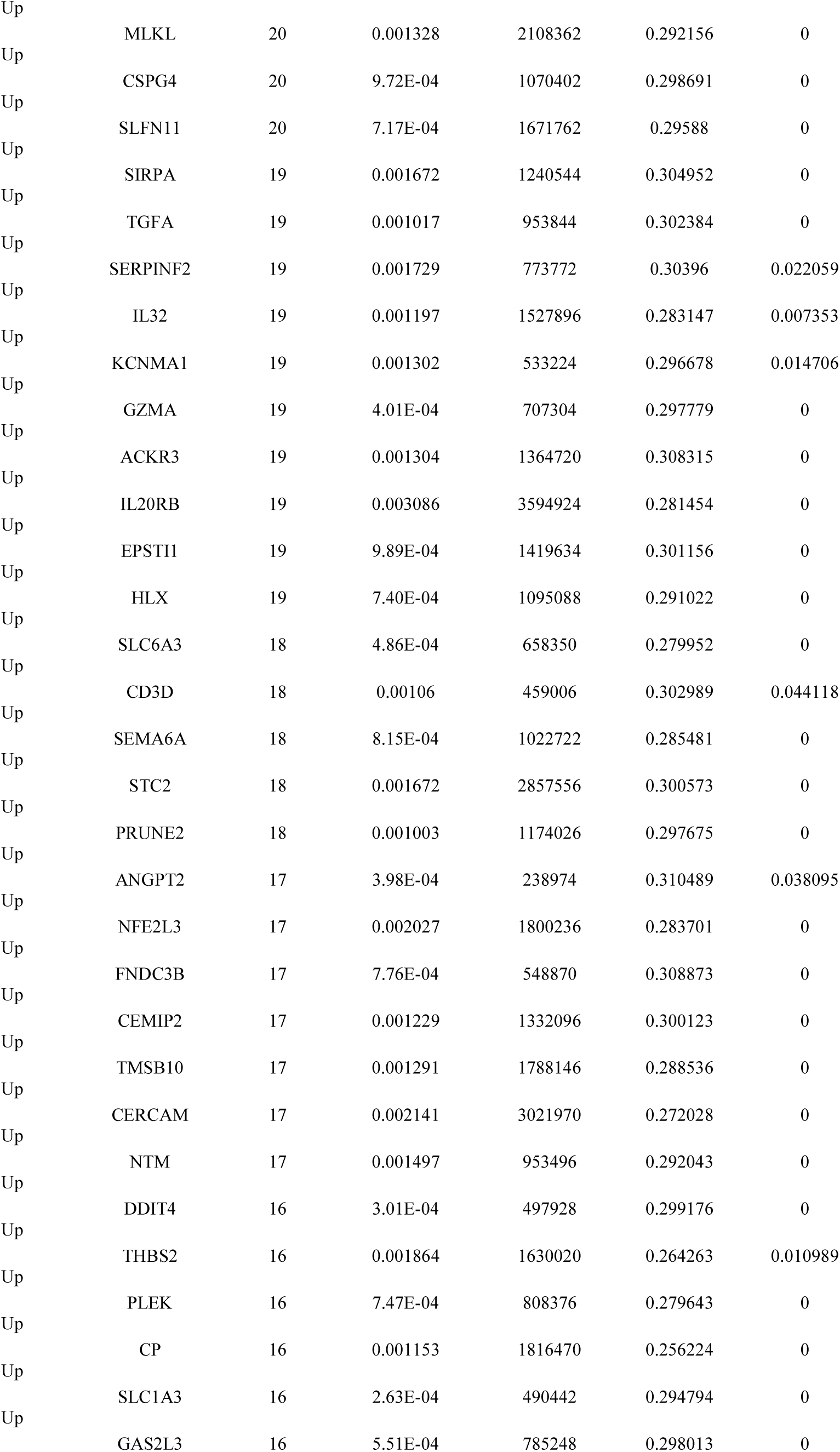
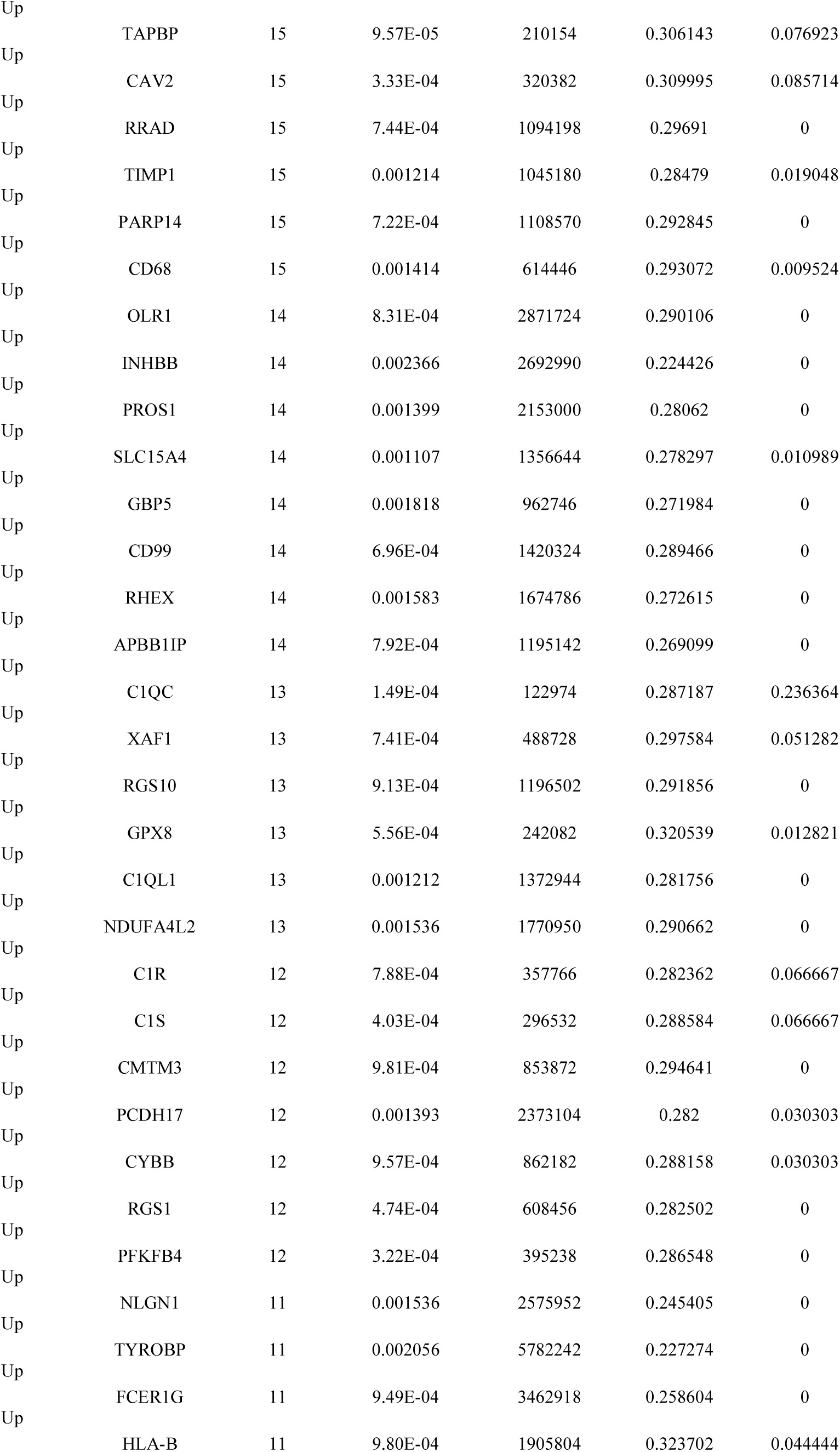
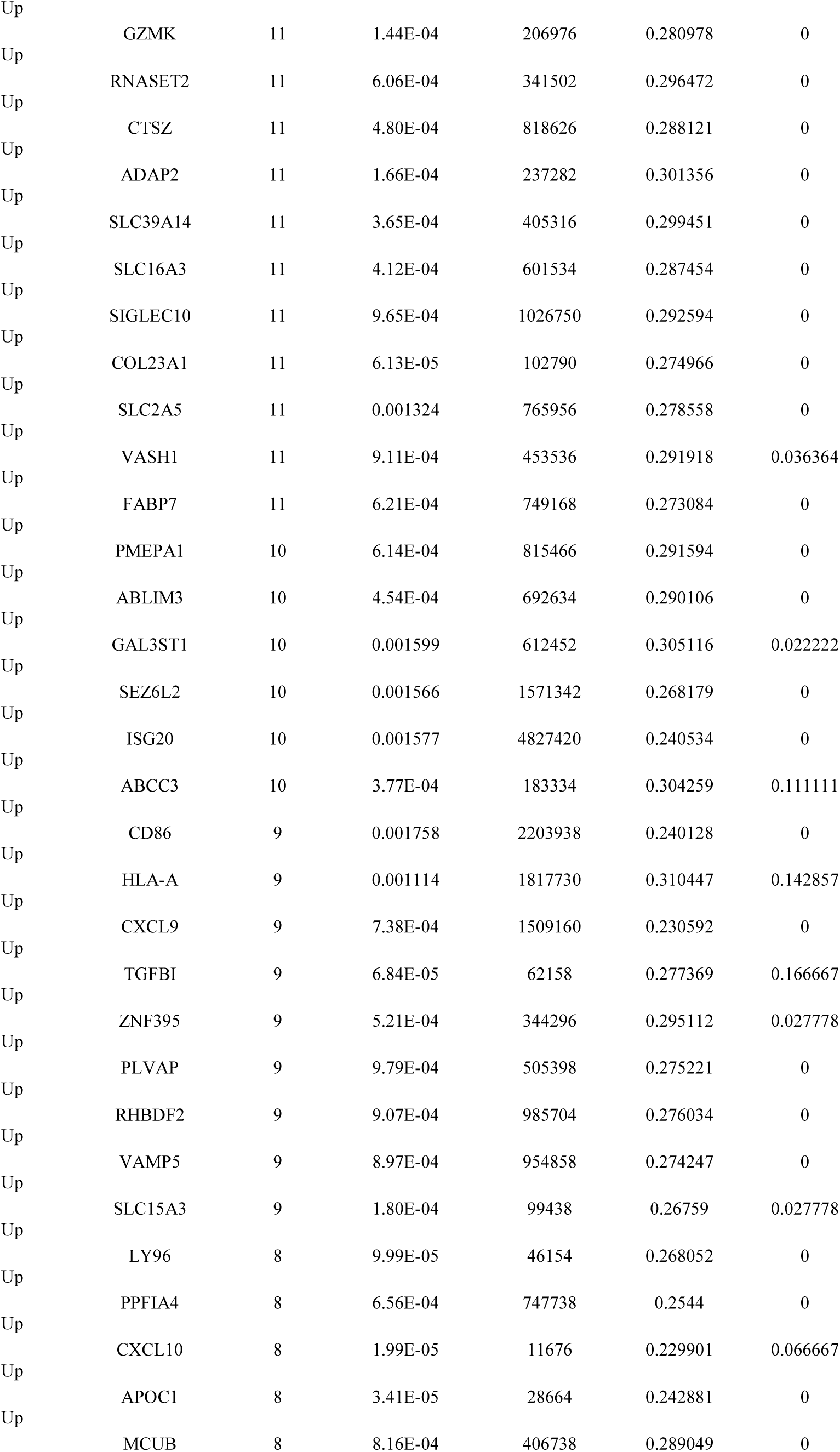
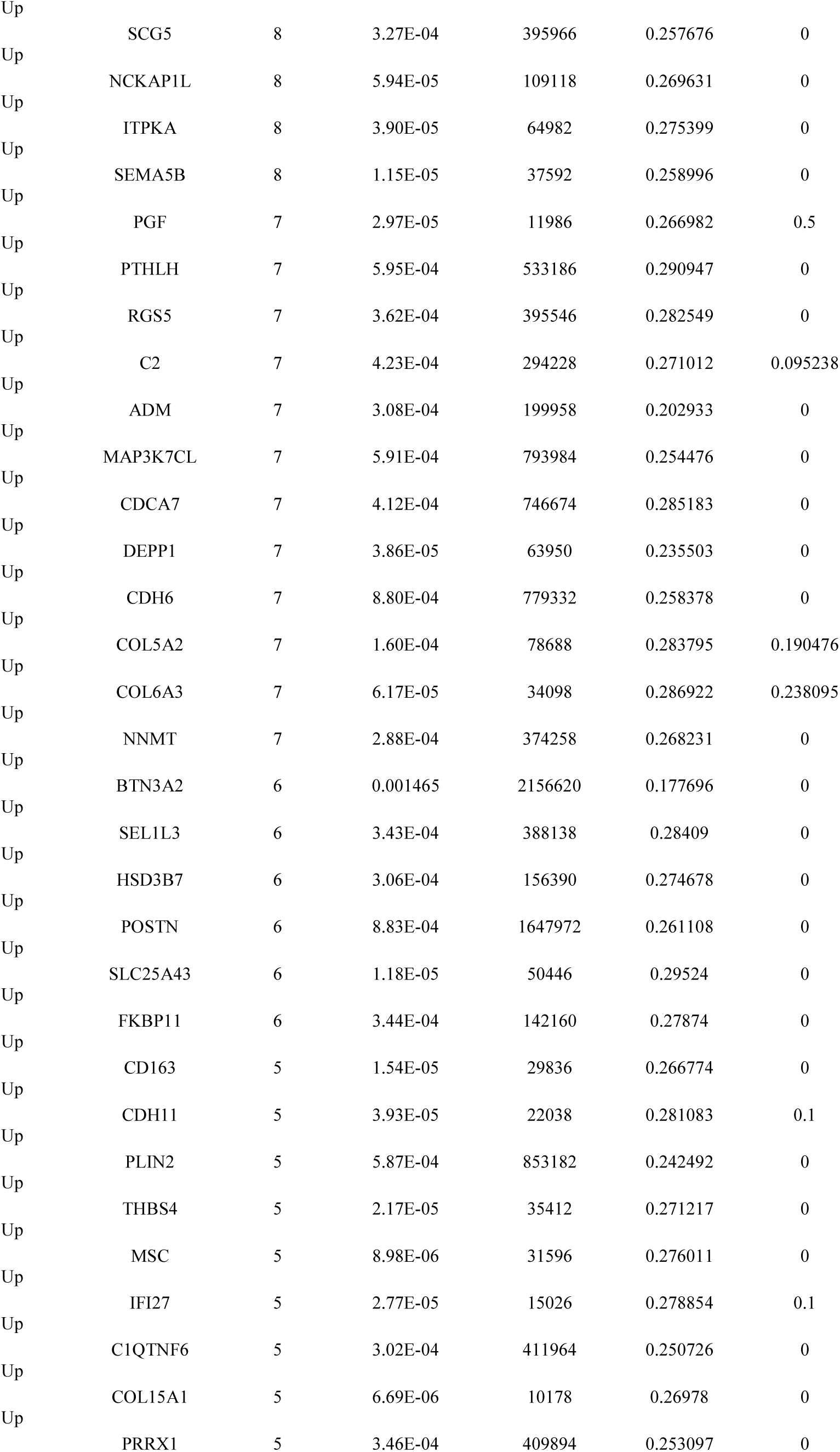
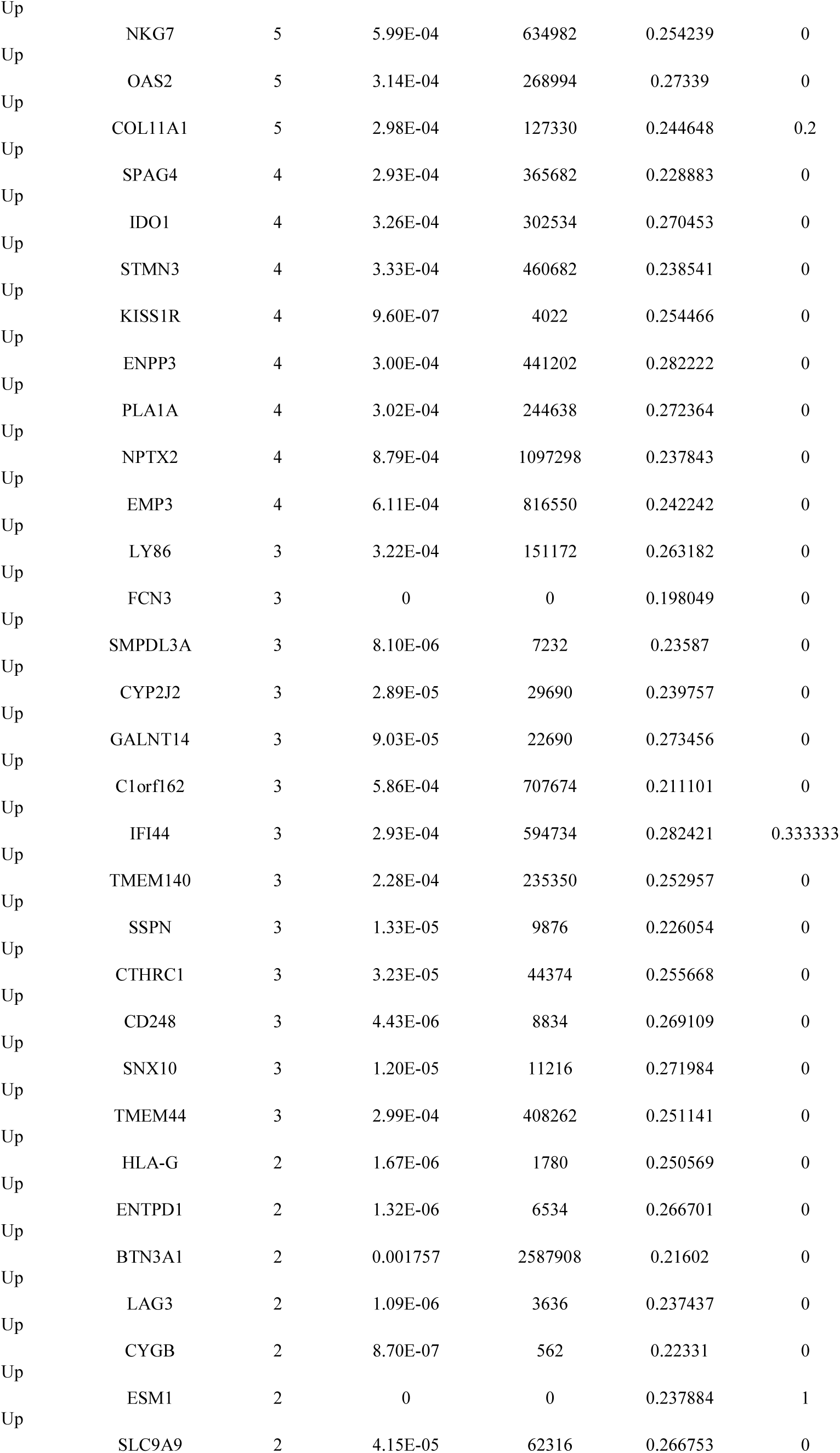
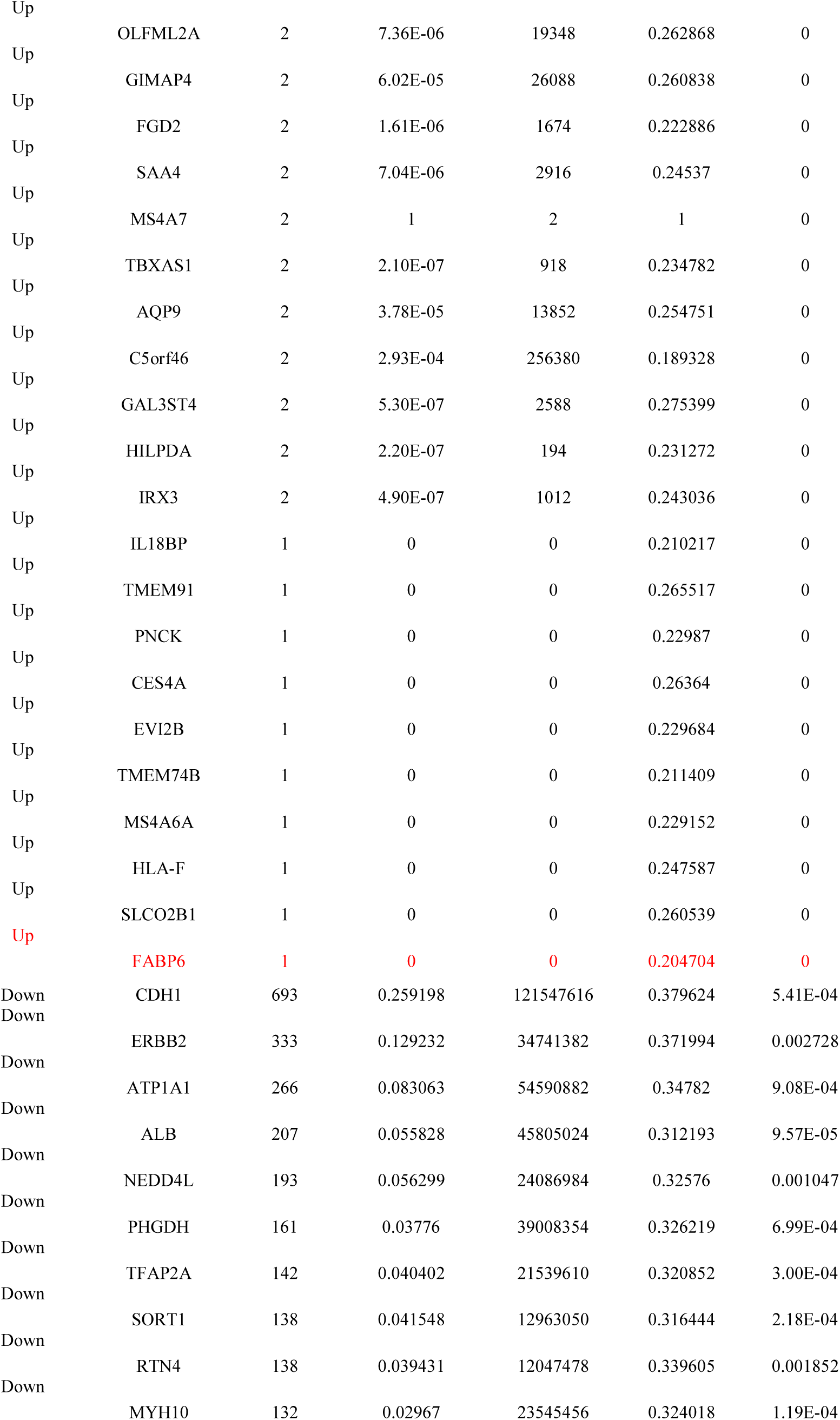
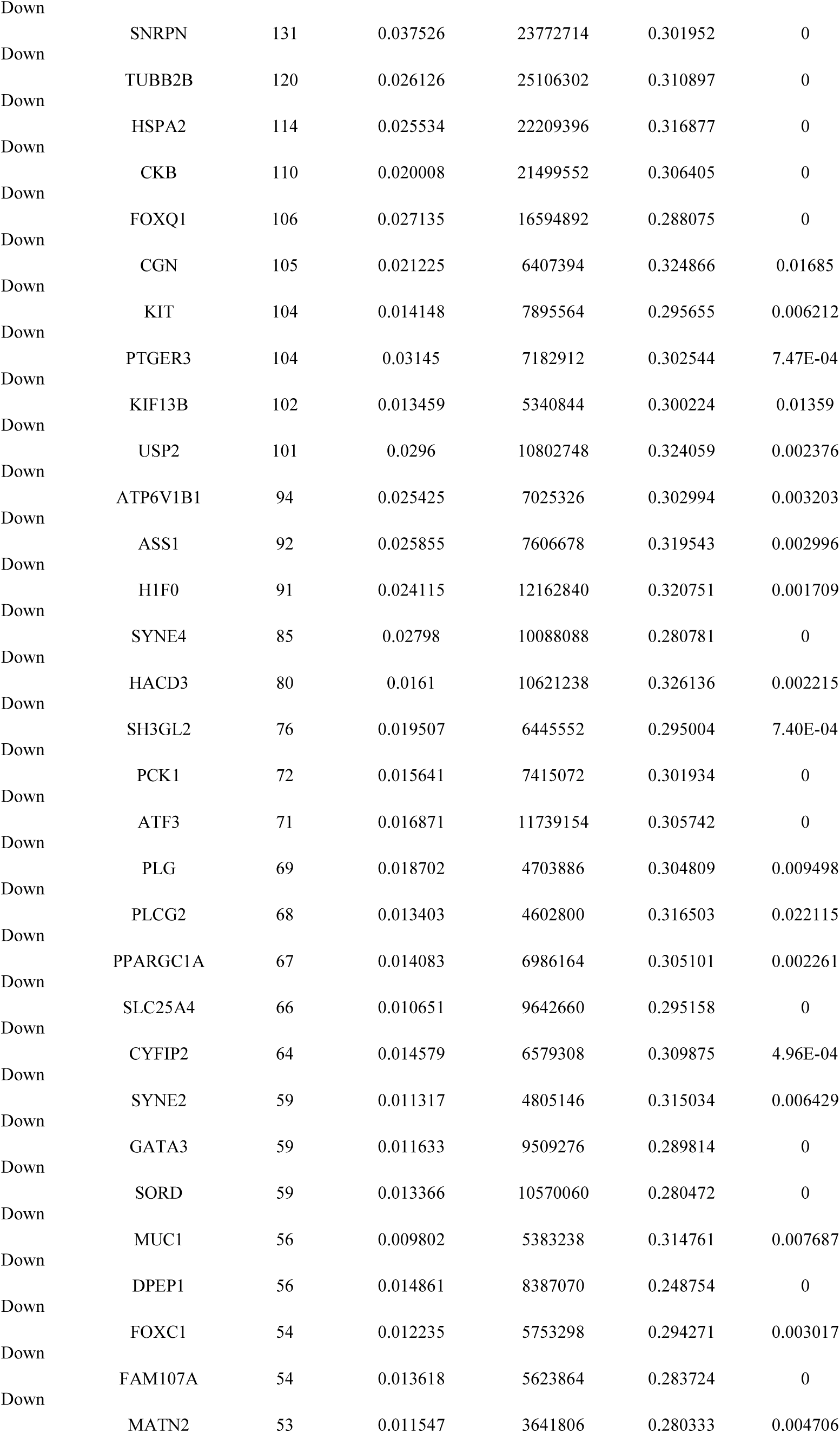
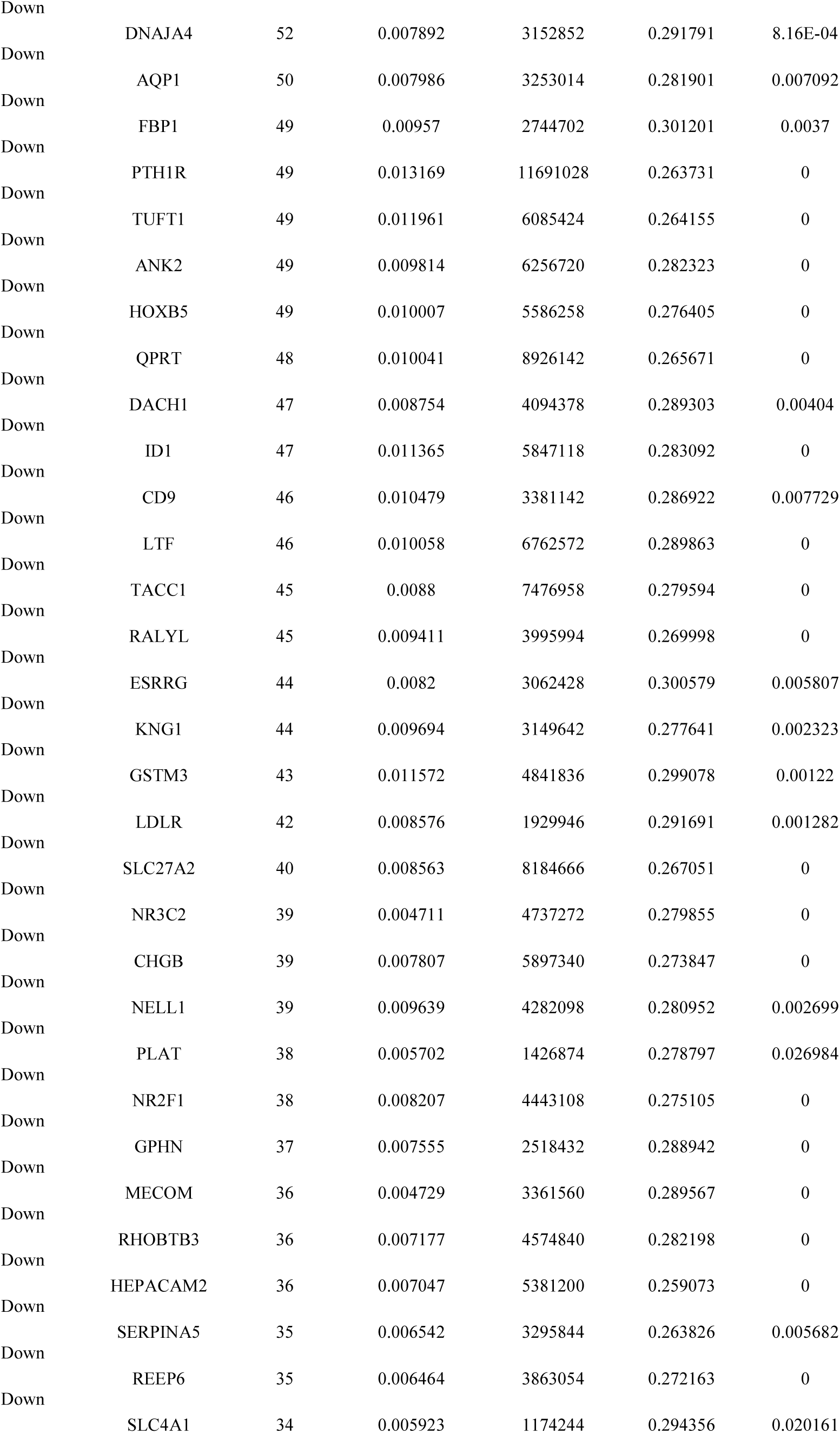
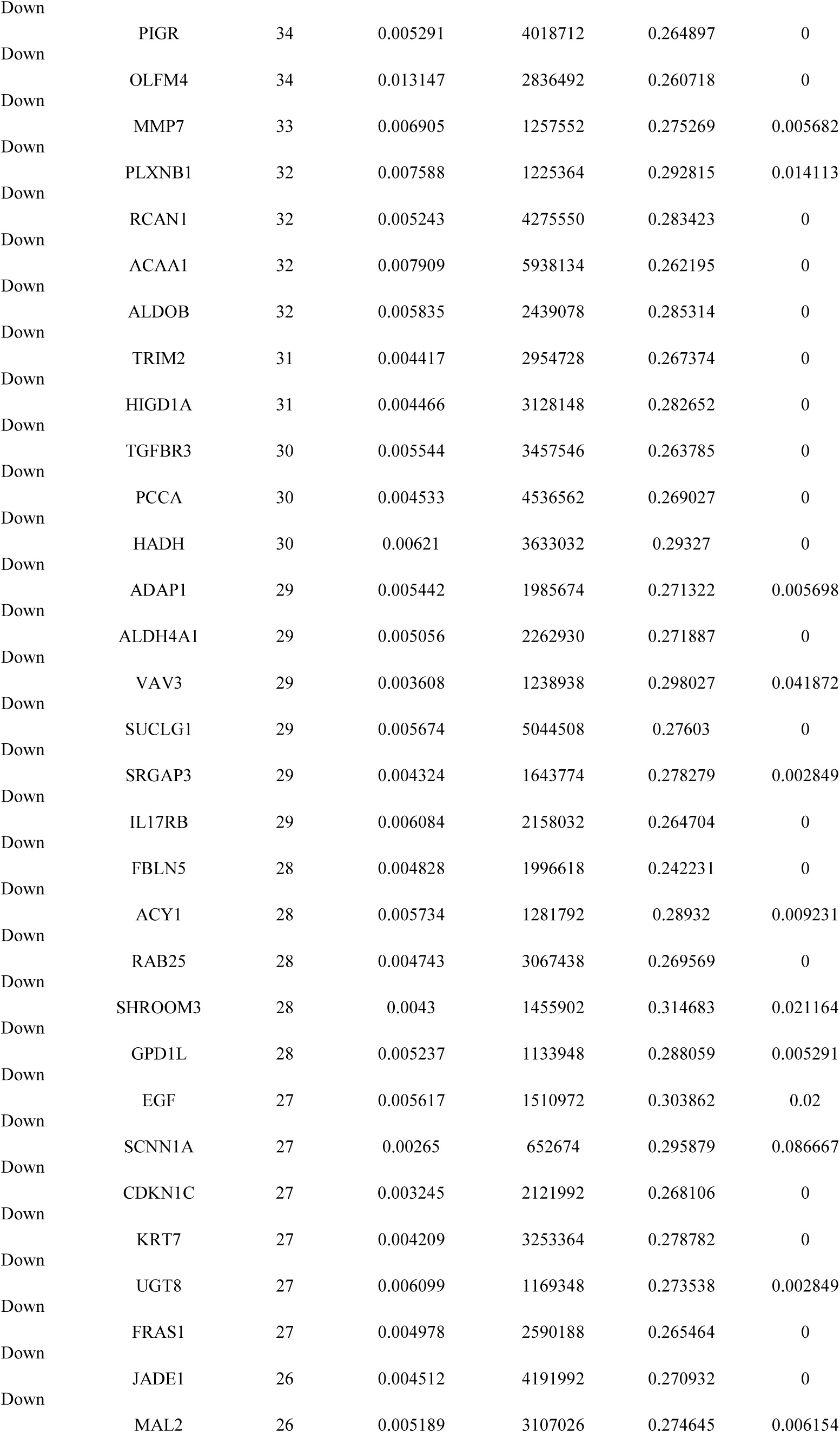
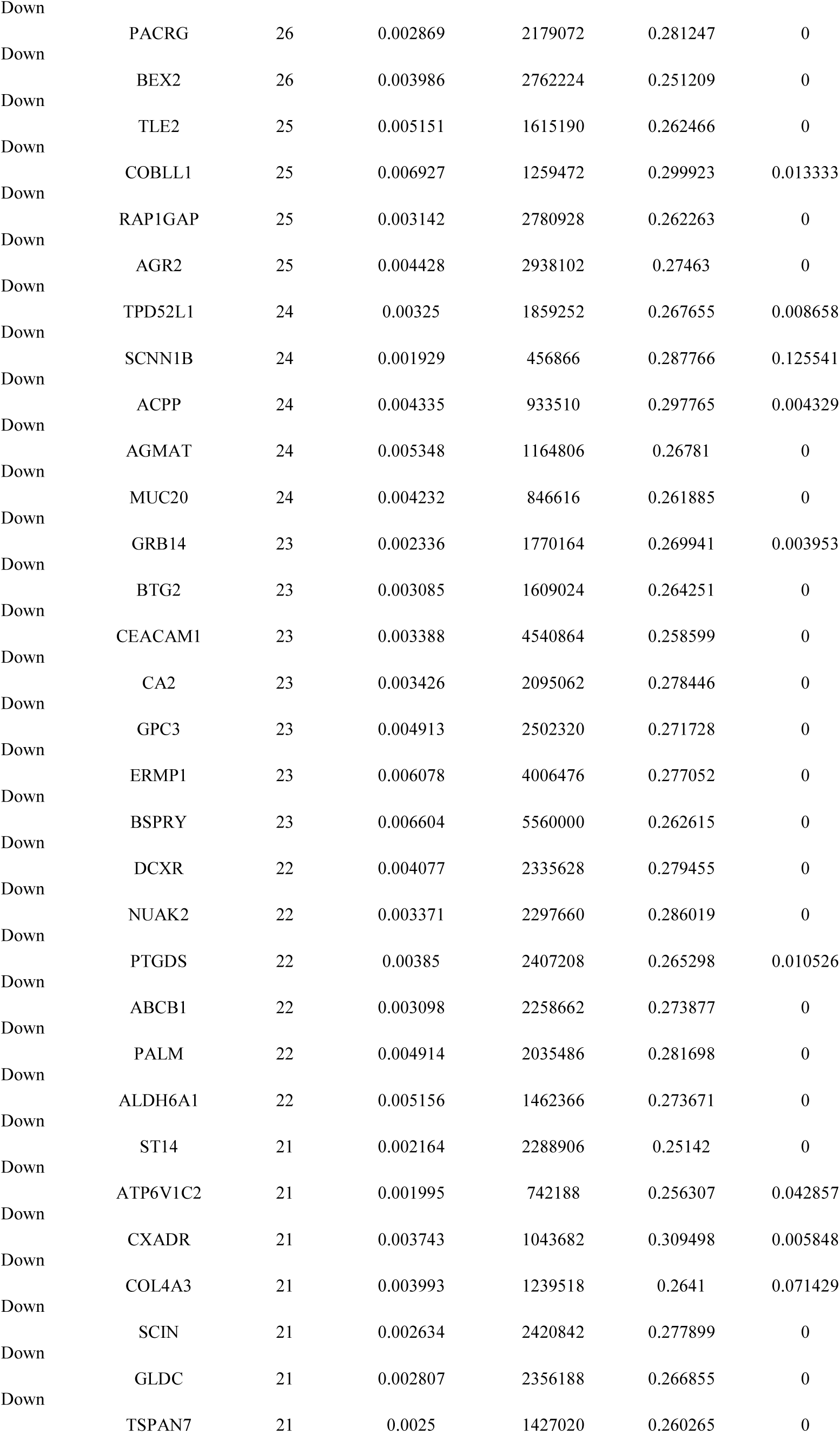
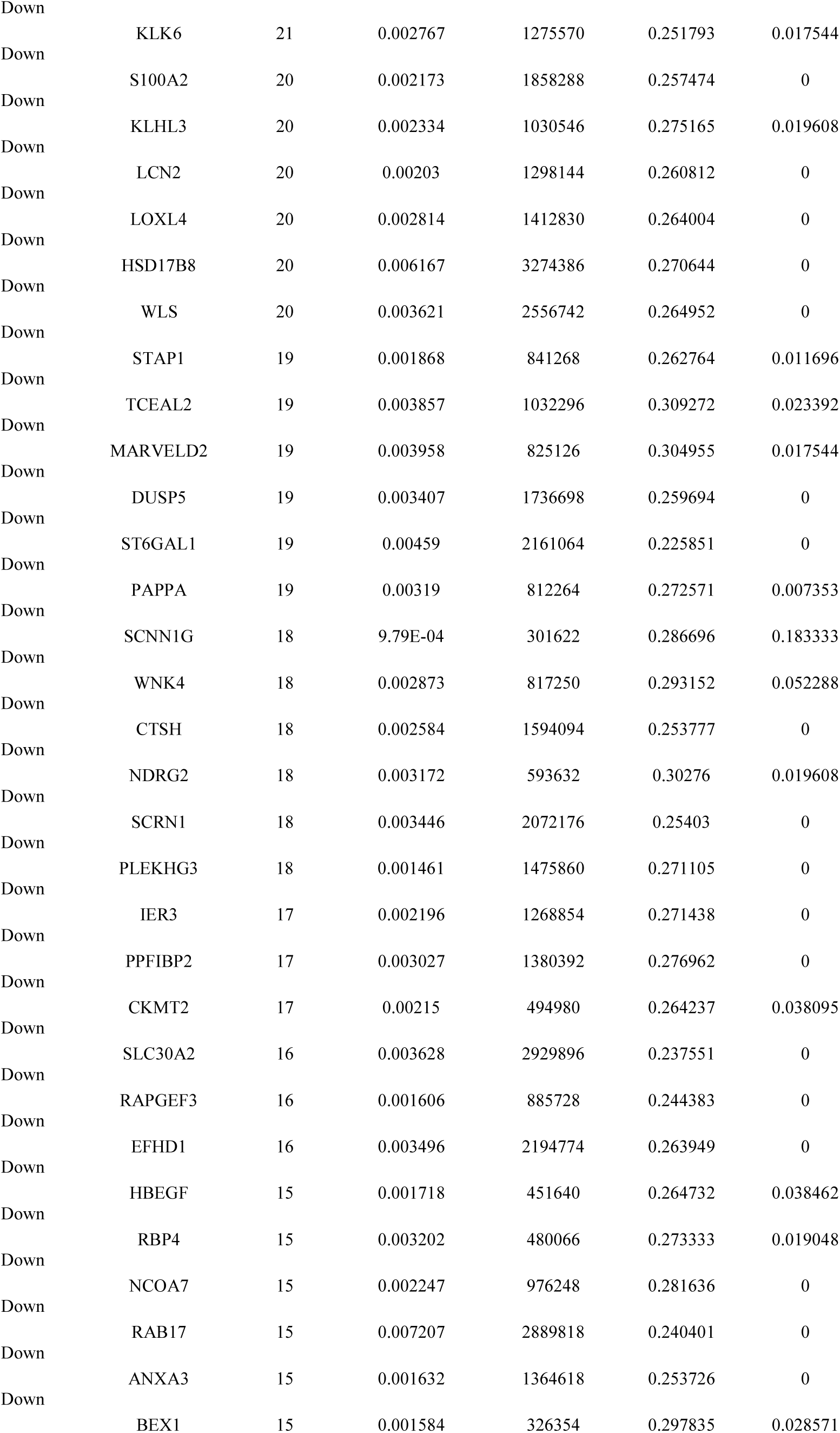
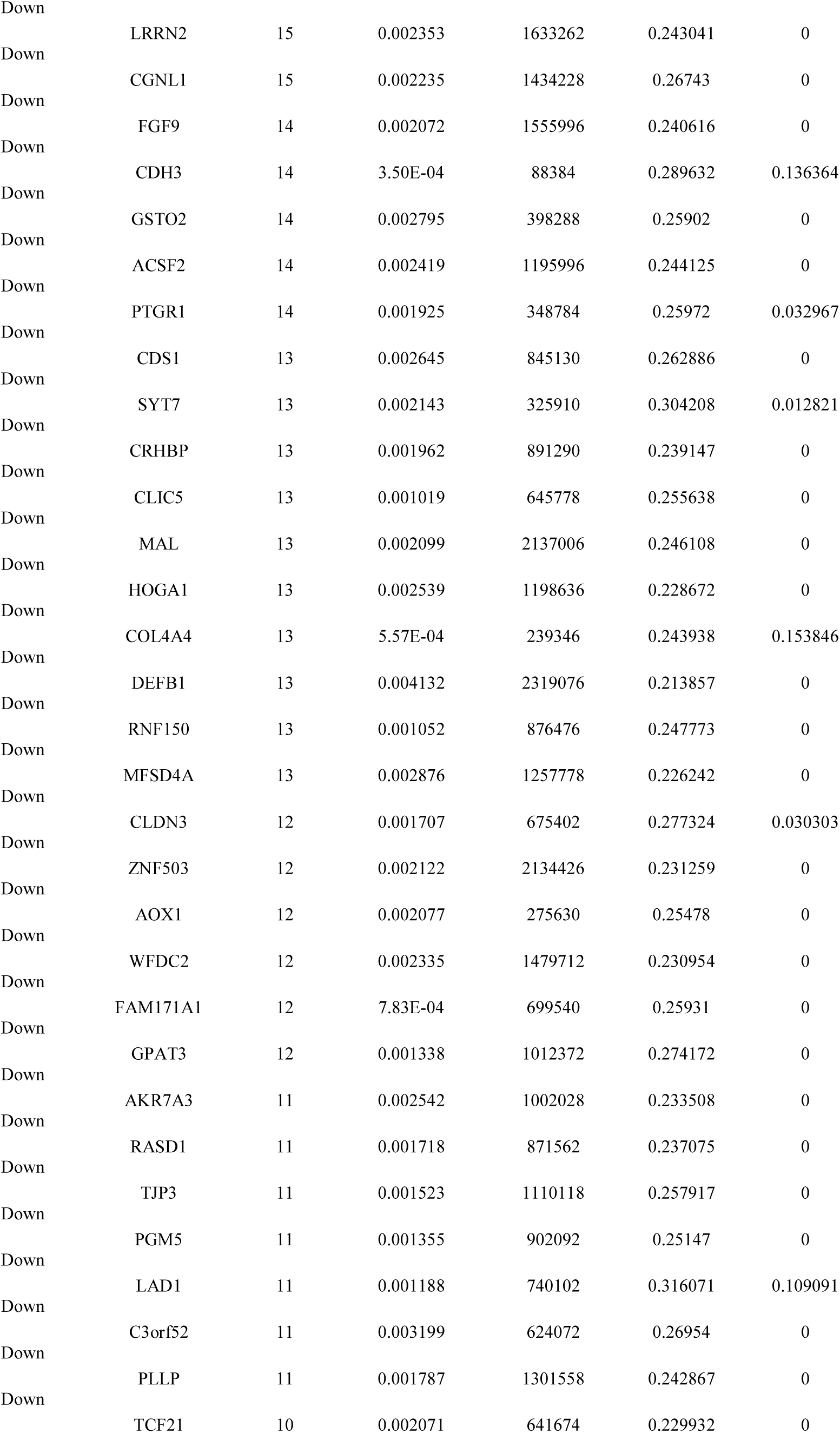
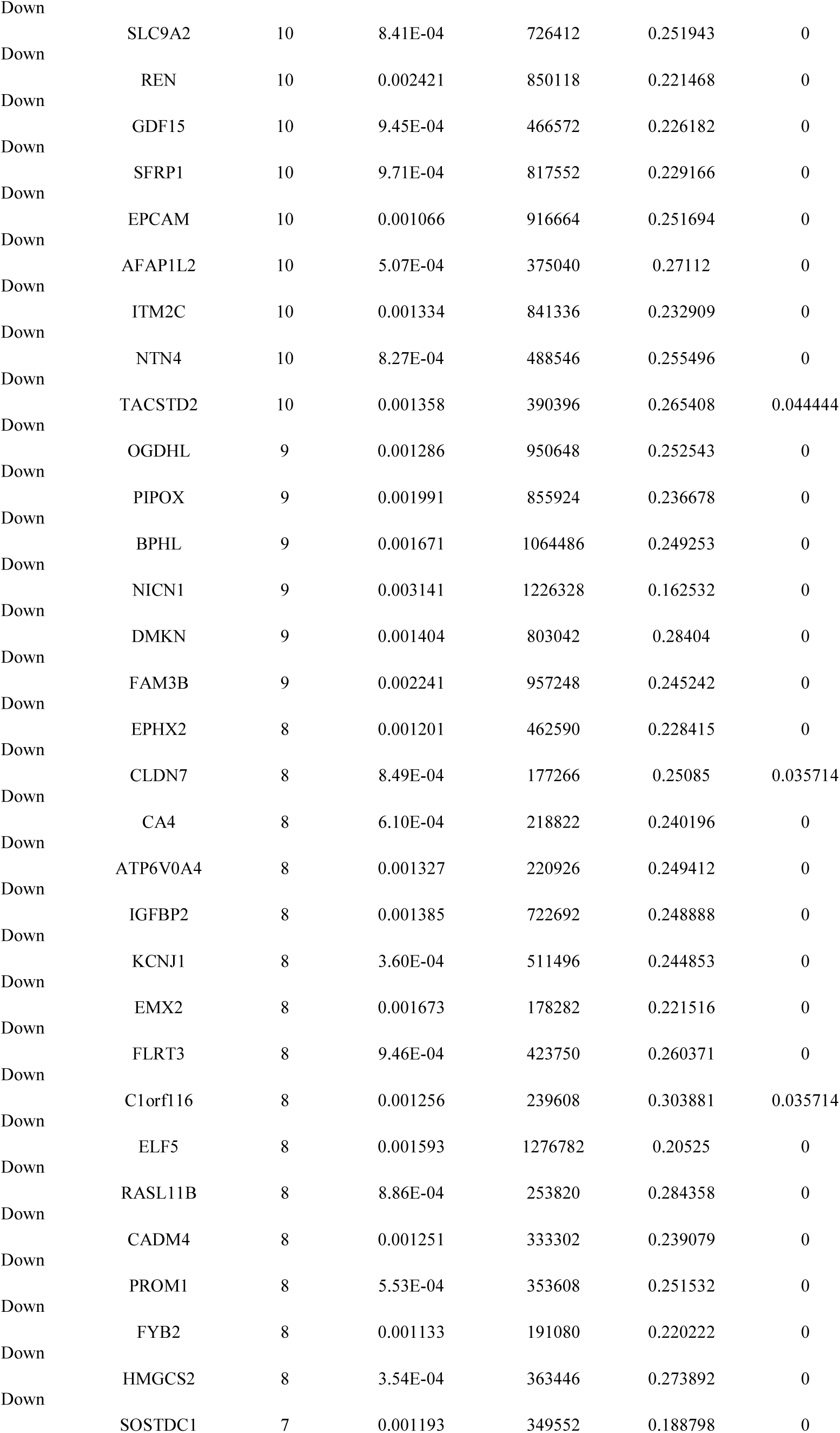
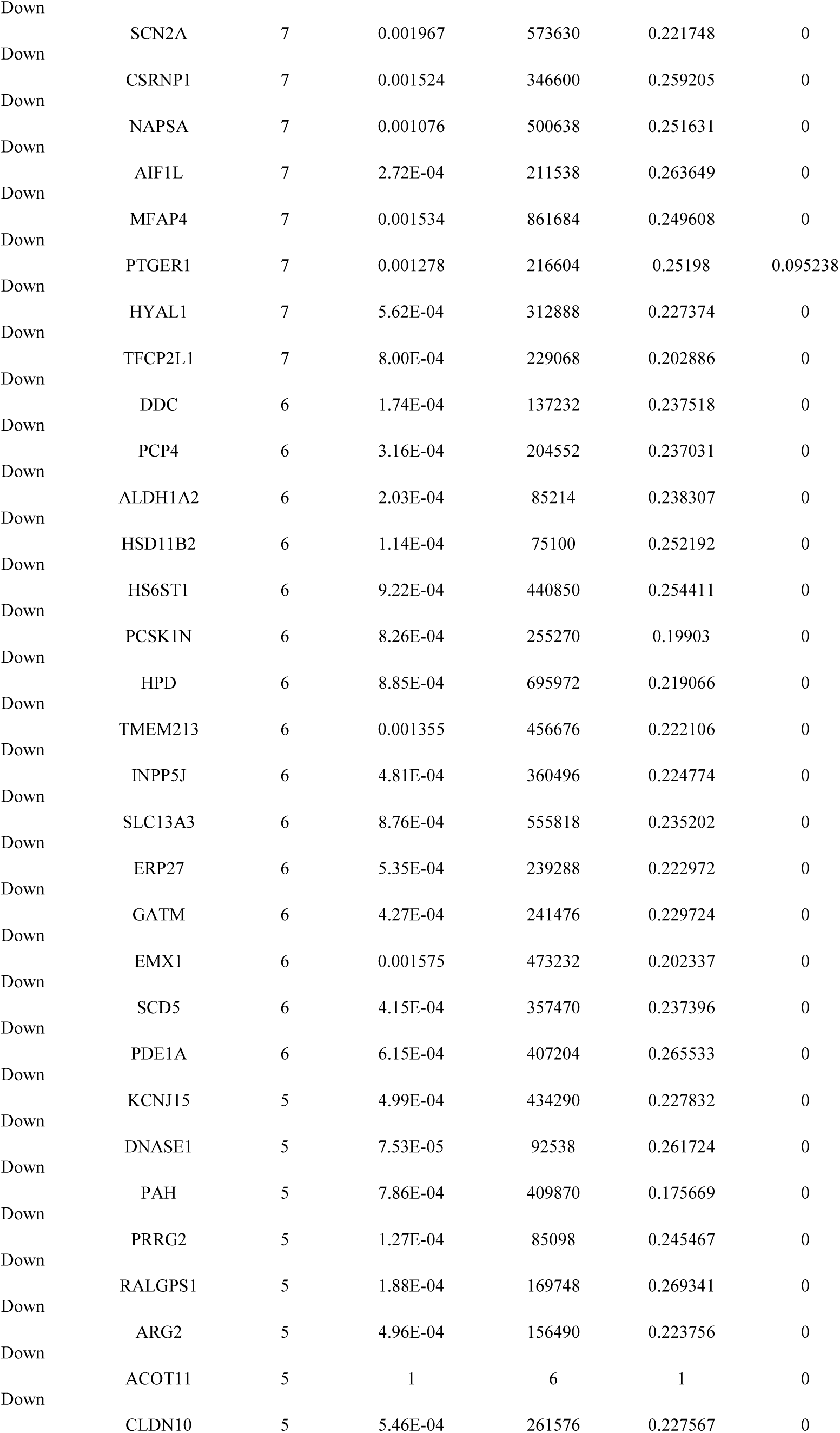
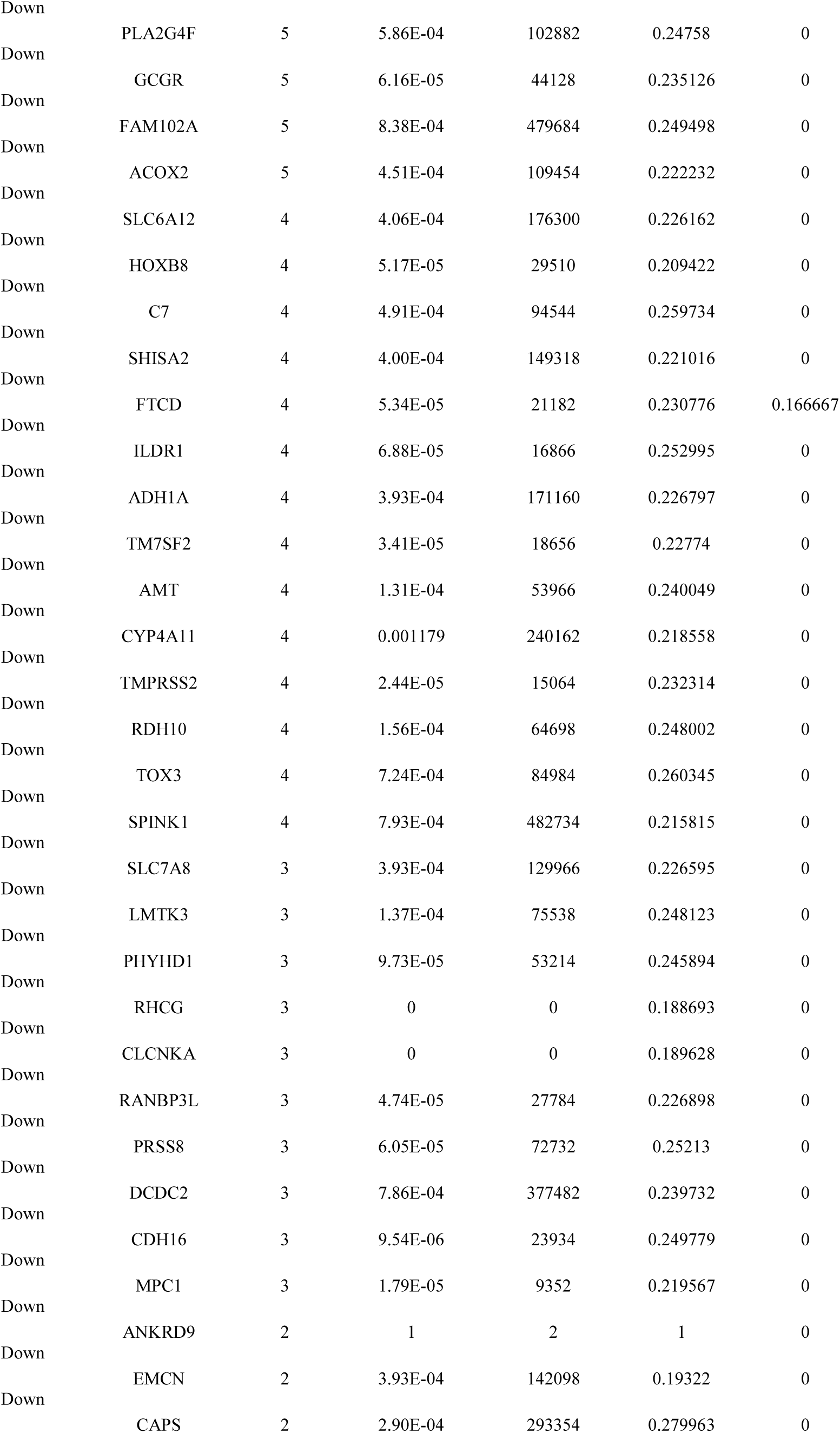
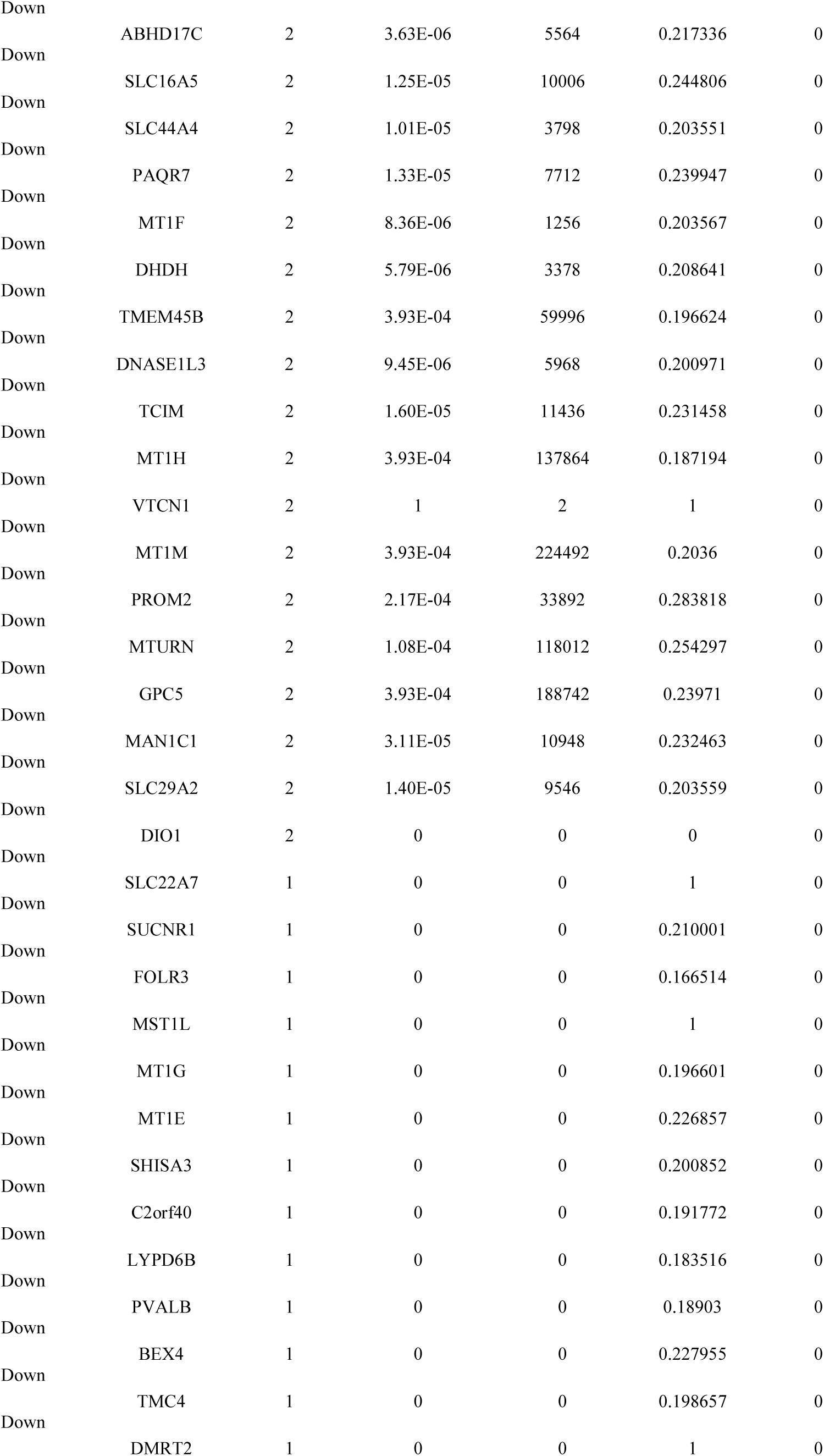
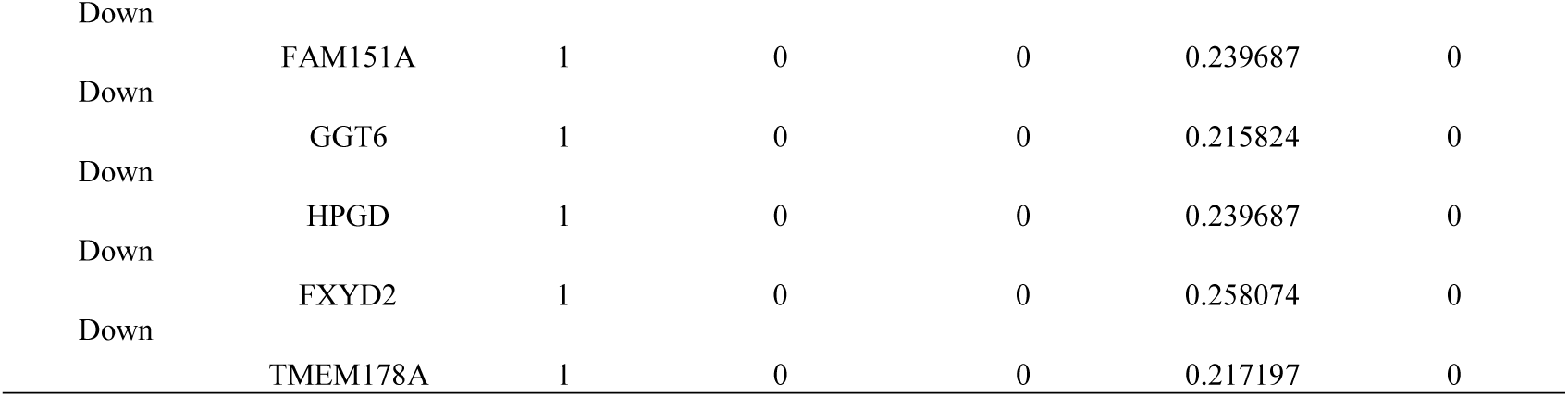
Topology table for up and down regulated genes

### Construction of target genes - TF regulatory network

Many of the TFs were responsible for advancement of CCRCC [Higgins et al 2003]. The current study focused on the five hub genes, and further assessed their TF-target associations. The TF-target regulatory network was based on interactions in the JASPAR database. A target genes - TF regulated network was established, including connections and top five up regulated genes, which are presented in Fig. 13. Specifically, THBS2 regulates 217 TFs (ex; FOXC1), VCAN regulates173 TFs (ex; GATA2), MLKL regulates 102 TFs (ex; YY1), RHBDF2 regulates 100 TFs (ex; FOXL1) and COL11A1 regulates 90 TFs (ex; E2F1) are listed in Table 8. The pathway and GO enrichment analysis reveled that up regulated targate genes were enriched in phagosome, direct p53 effectors, protein-containing complex binding, secretion and extracellular matrix organization. Similarly, target genes - TF regulated network was established, including connections and top five down regulated genes, which are presented in Fig. 14. Specifically, FOXQ1 regulates 175 TFs (ex; GATA2), EFHD1 regulates126 TFs (ex; YY1), NCOA7 regulates86 TFs (ex; FOXL1), PTH1R regulates 84 TFs (ex; USF2) and MUC20 regulates 83 TFs (ex; NFIC) are listed in Table 8. The pathway and GO enrichment analysis reveled that down regulated targate genes were enriched in epithelium development, mitochondrion, signaling receptor binding, signaling by GPCR and signaling events mediated by hepatocyte growth factor receptor (c-Met).

**Fig. 13.**
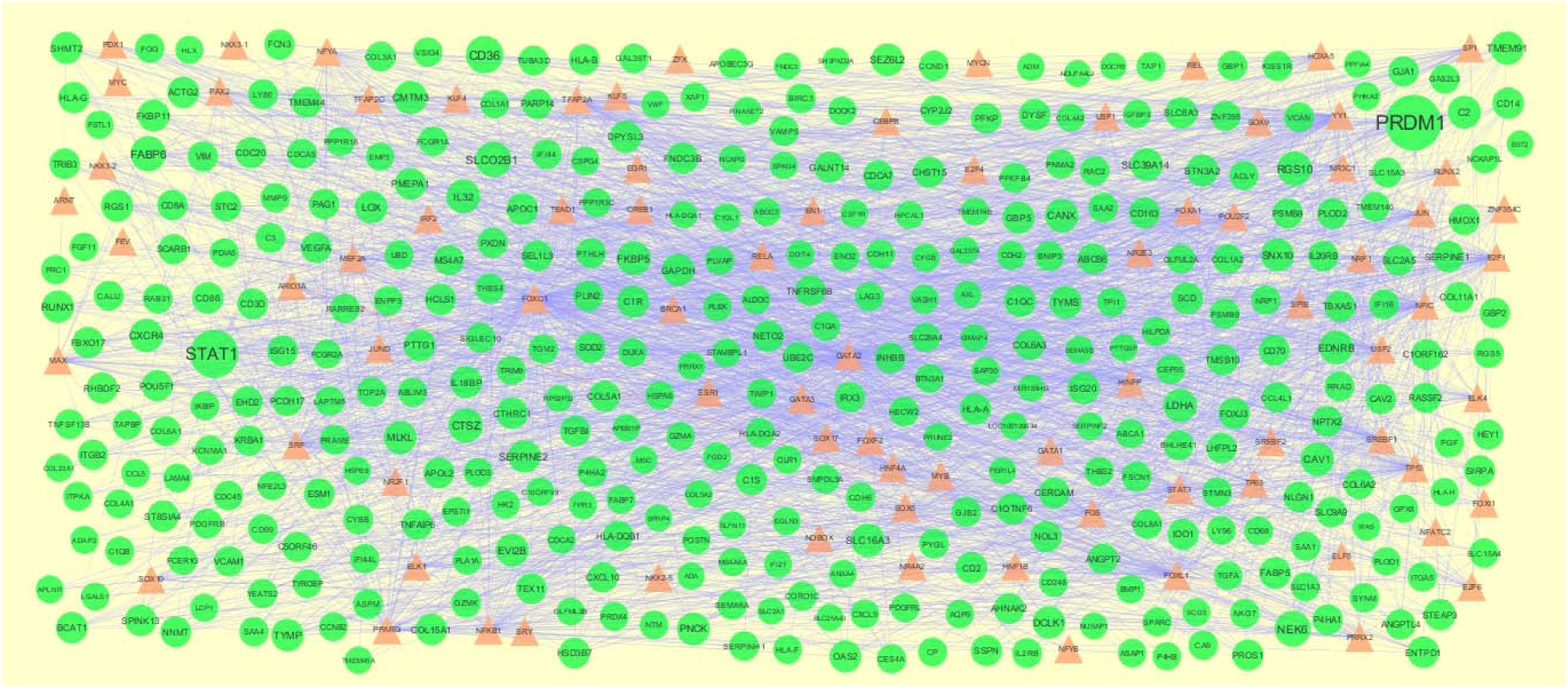
Target gene - TF network of predicted target up regulated genes. (Orange triangle - TFs and green circles-target up regulated genes)

**Fig. 14.**
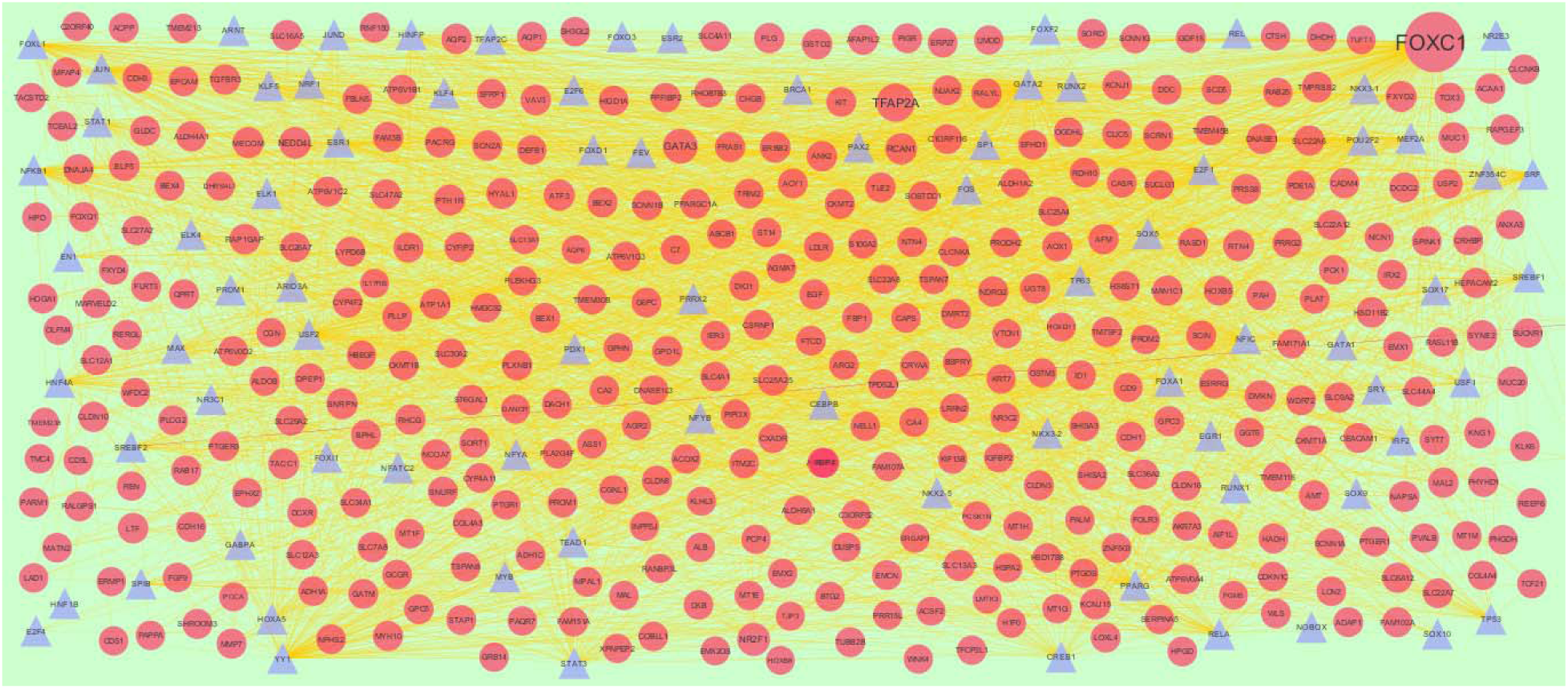
Target gene - TF network of predicted target down regulated genes. (Blue triangle - TFs and red circles-target up regulated genes)

**Table 8.**
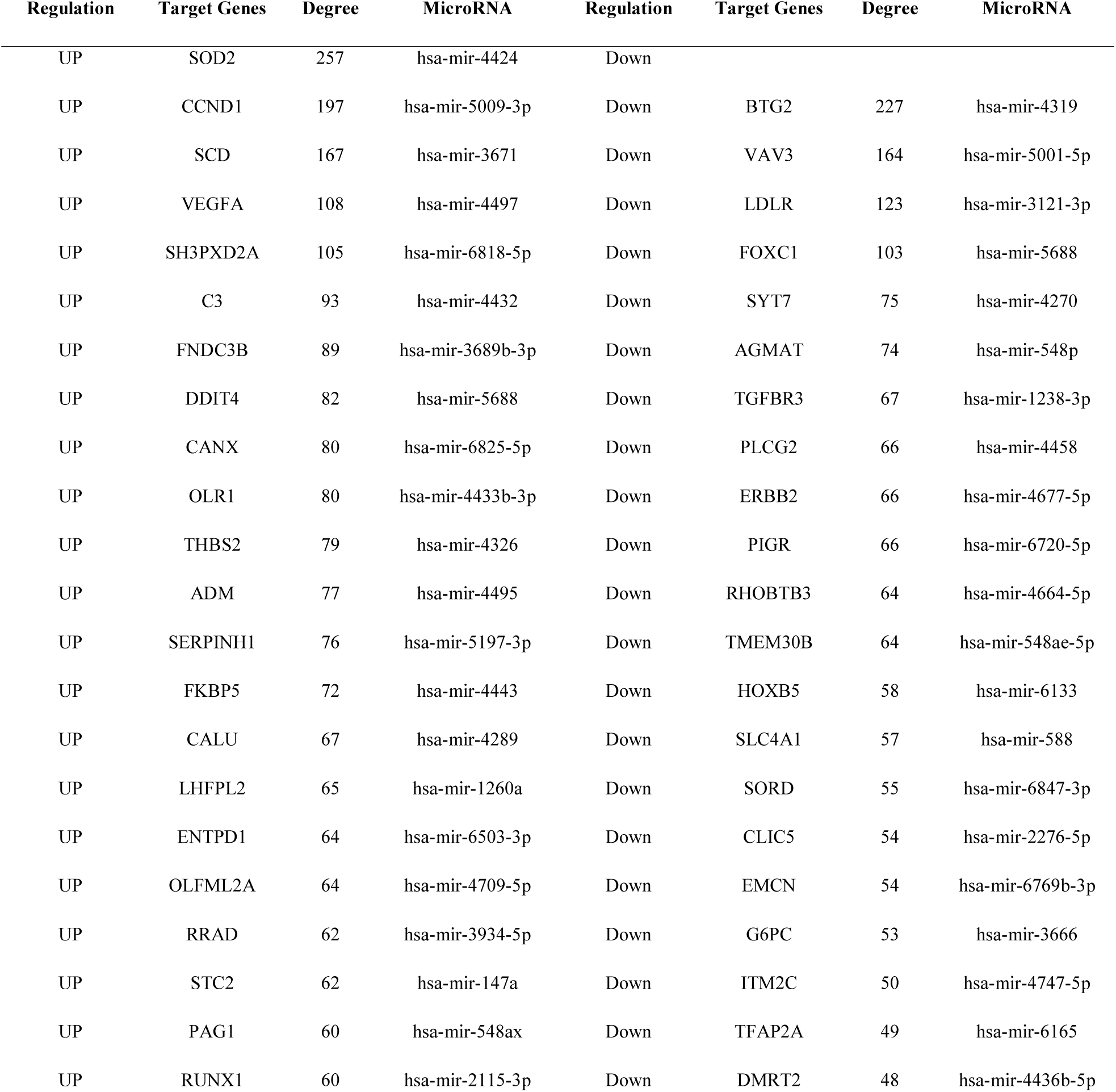
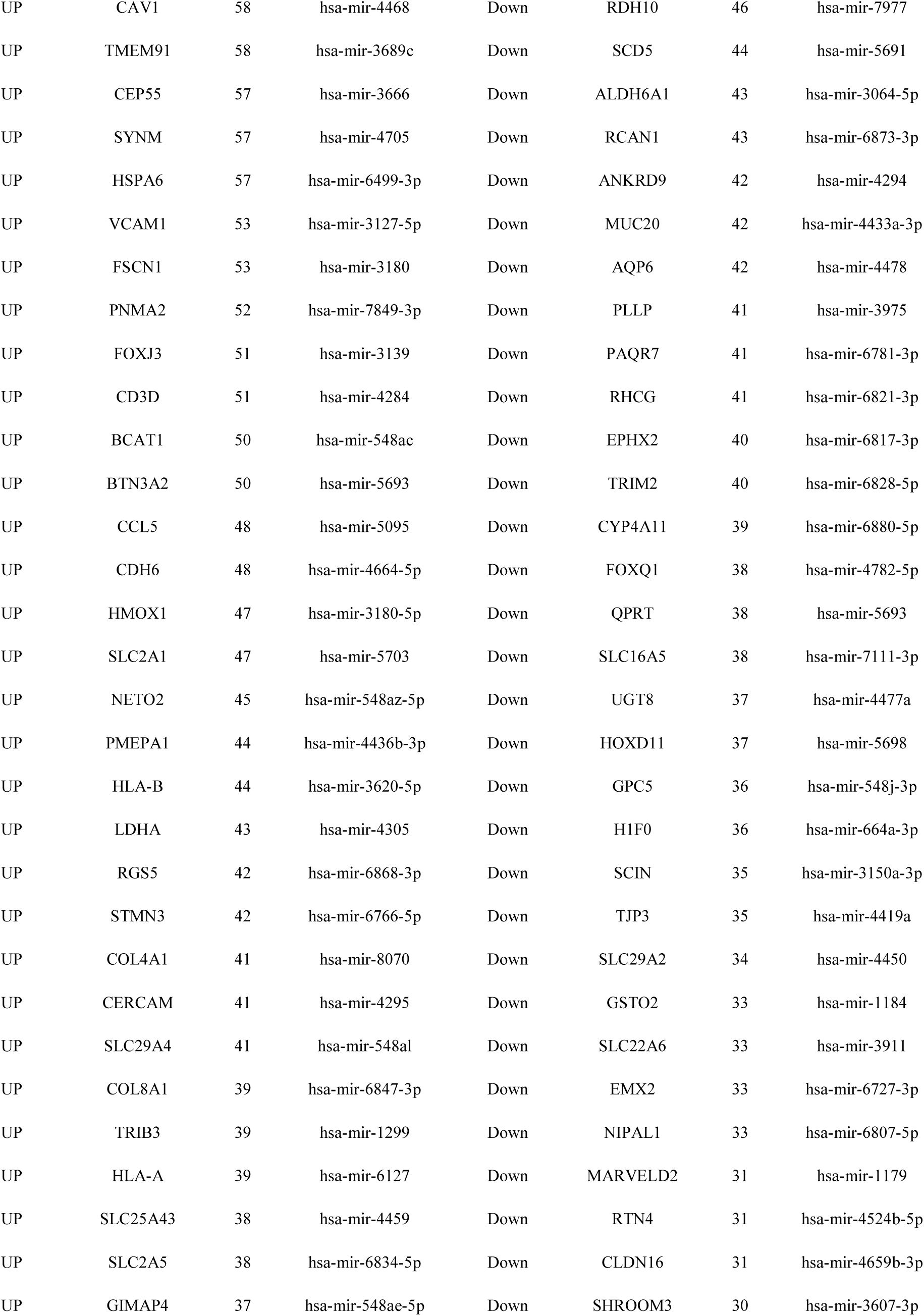
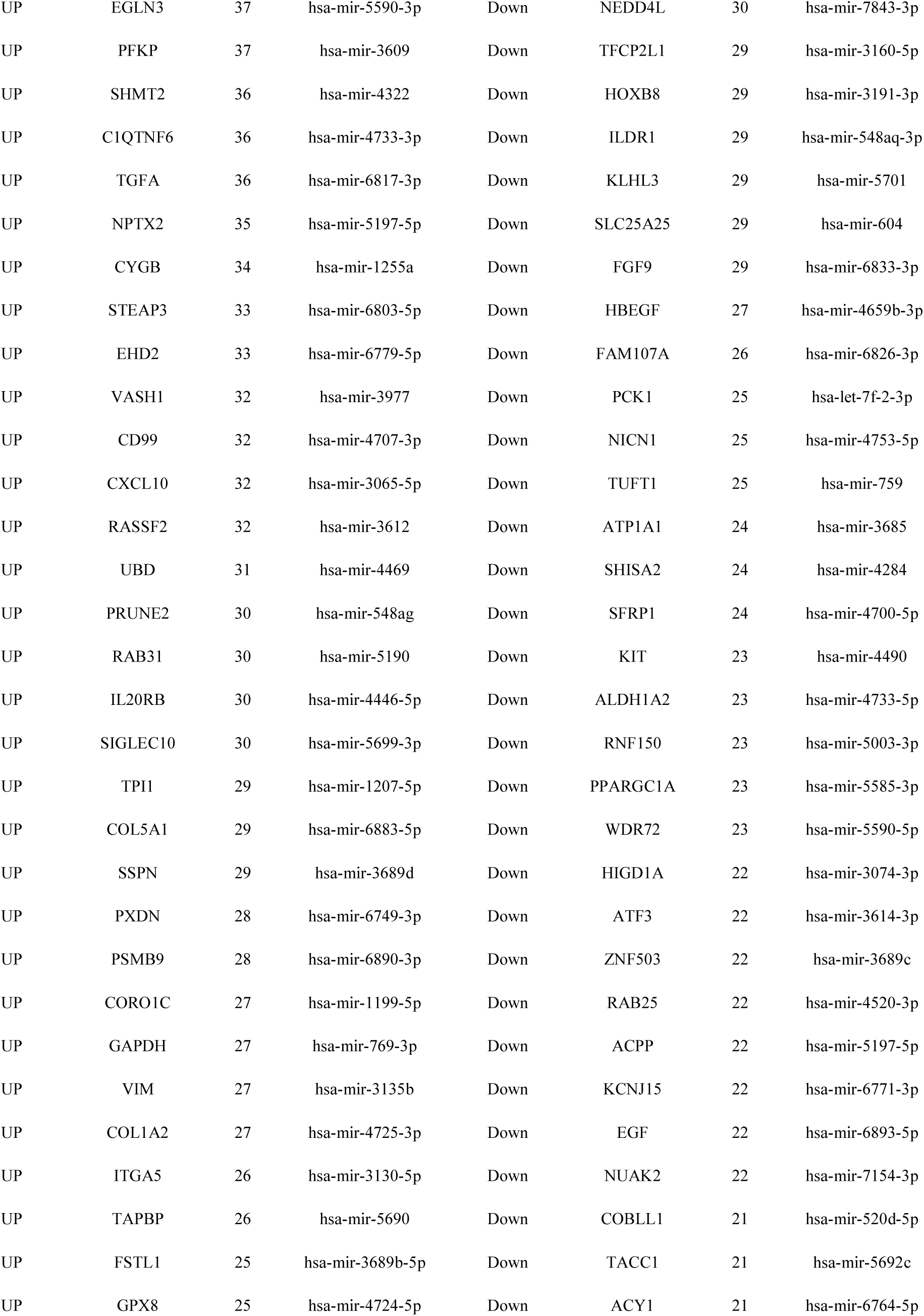
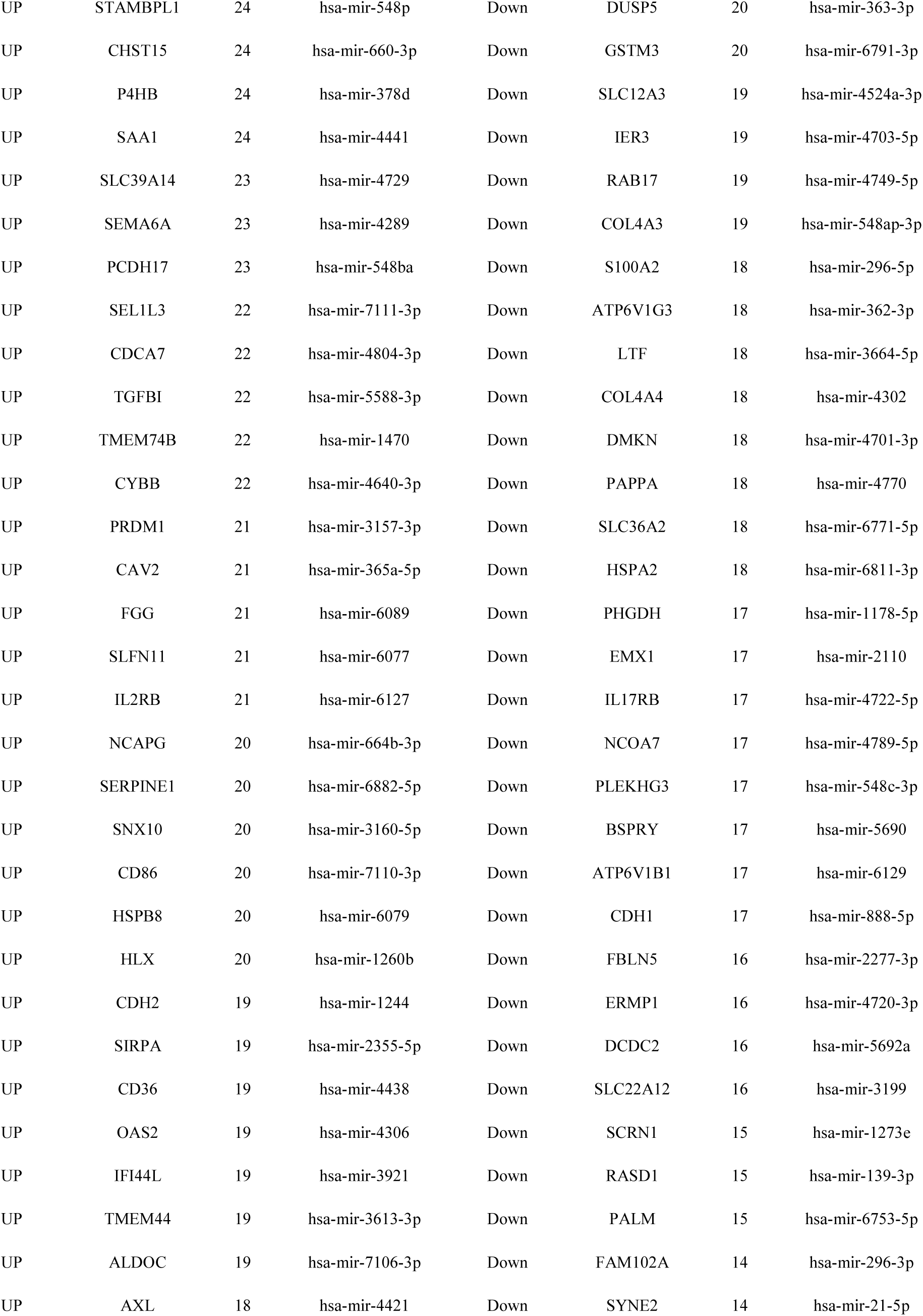
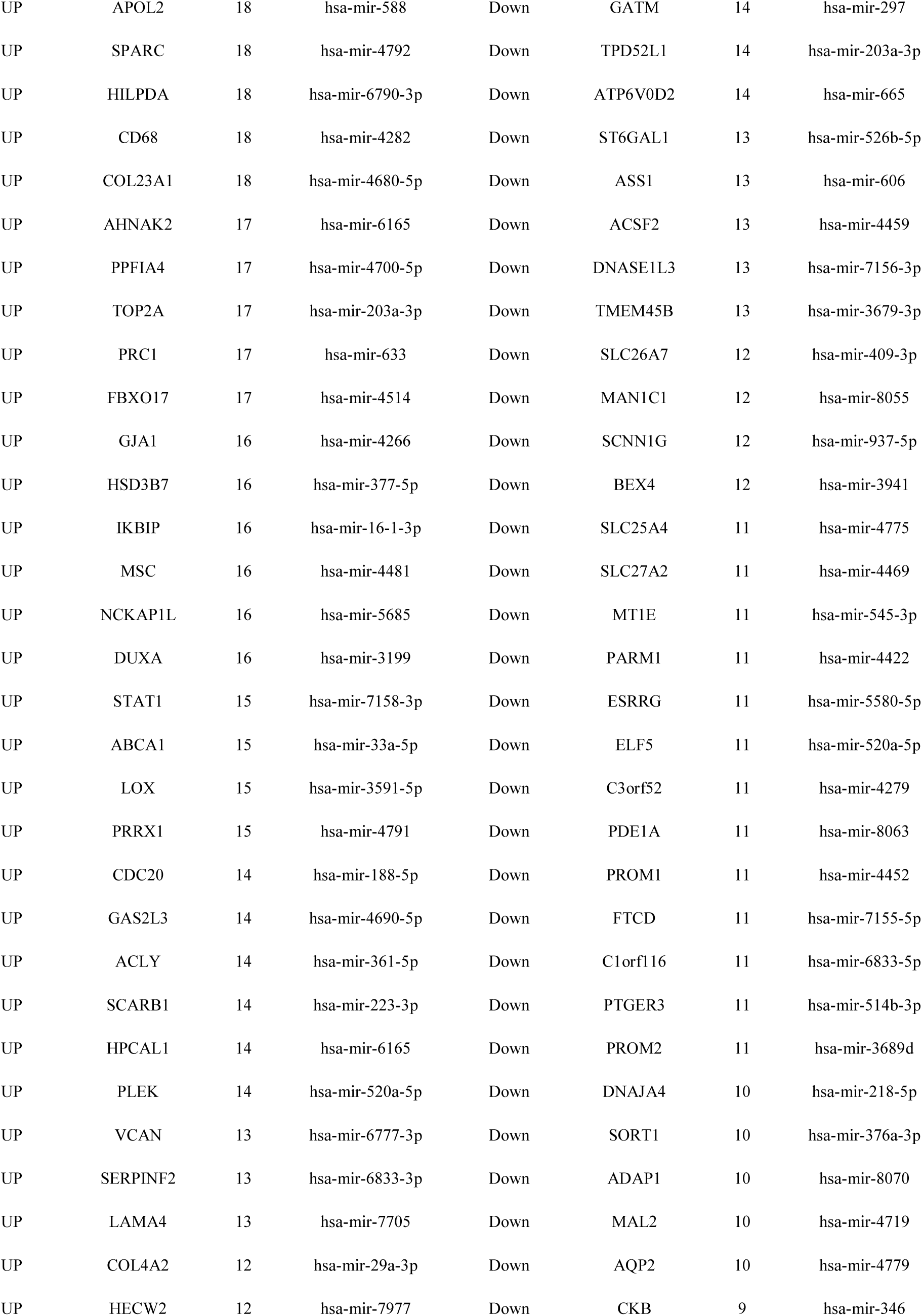
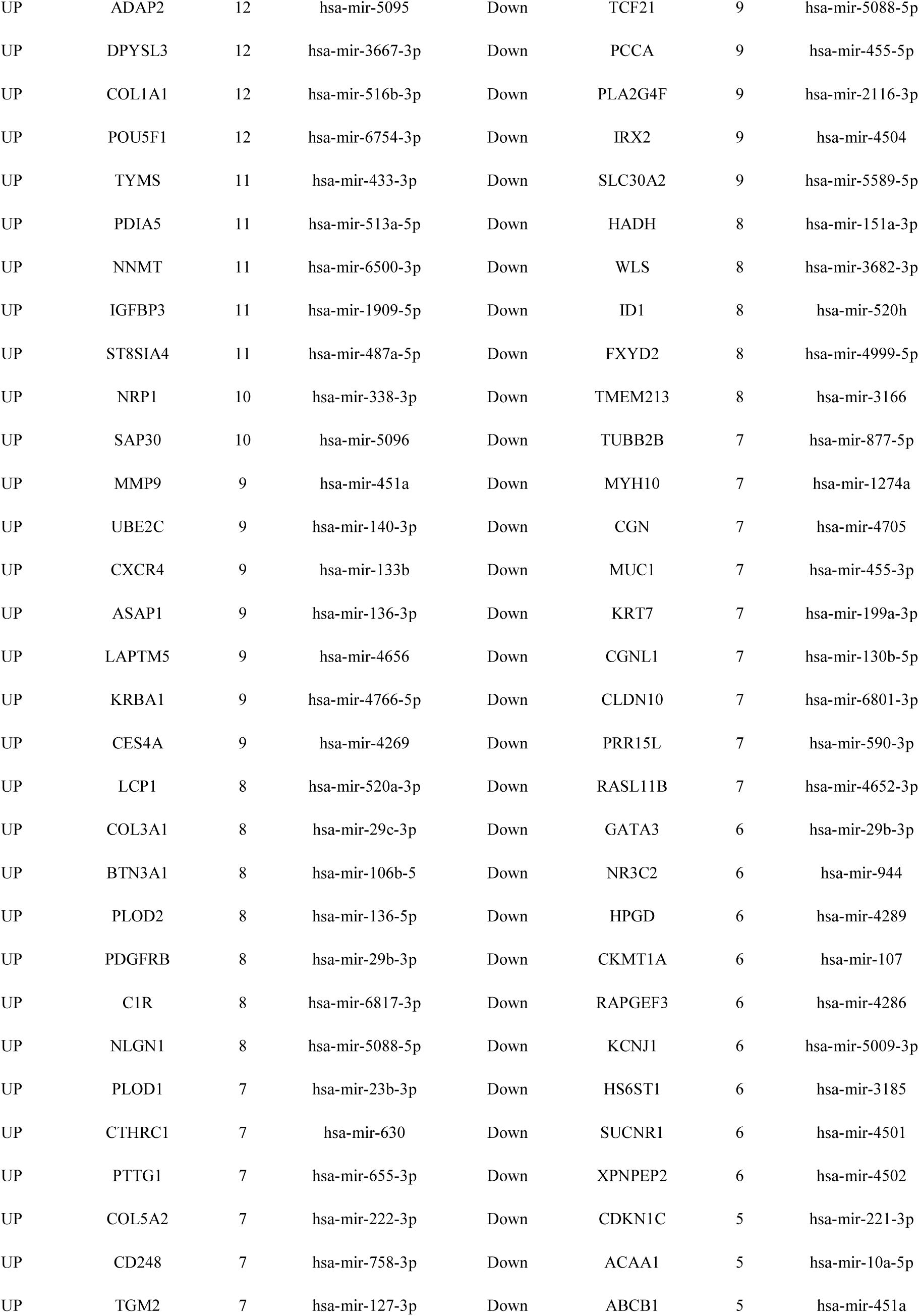
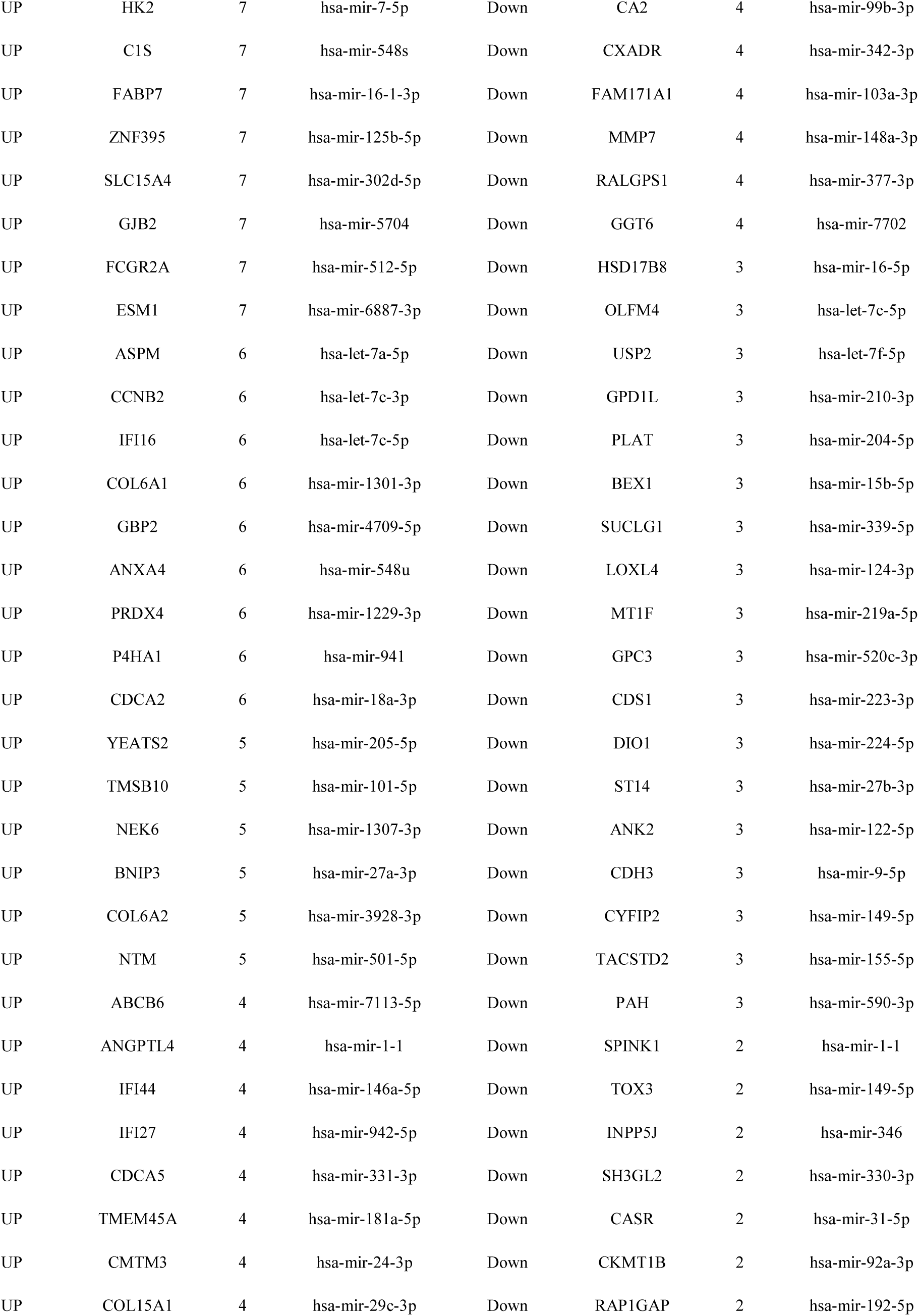
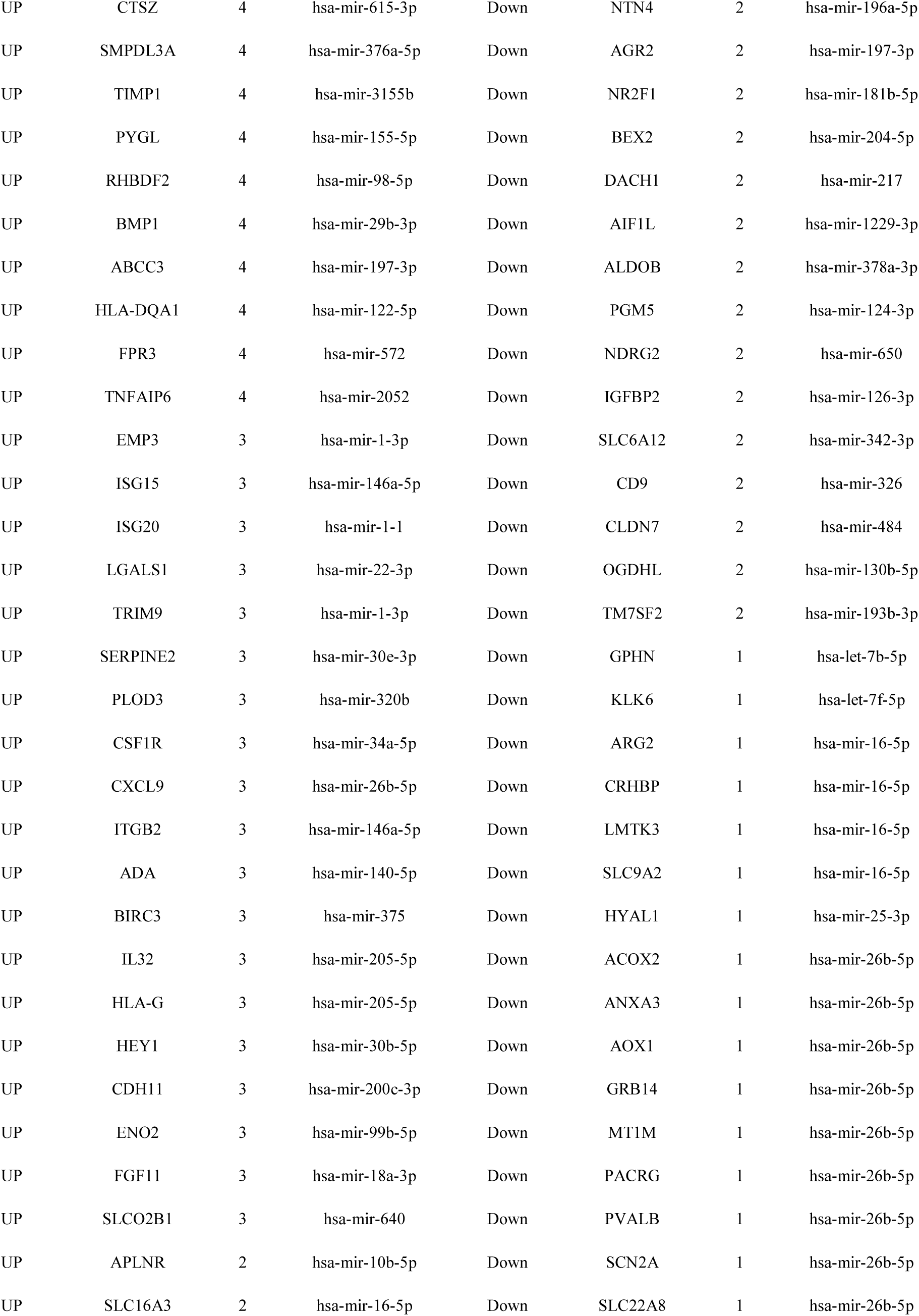
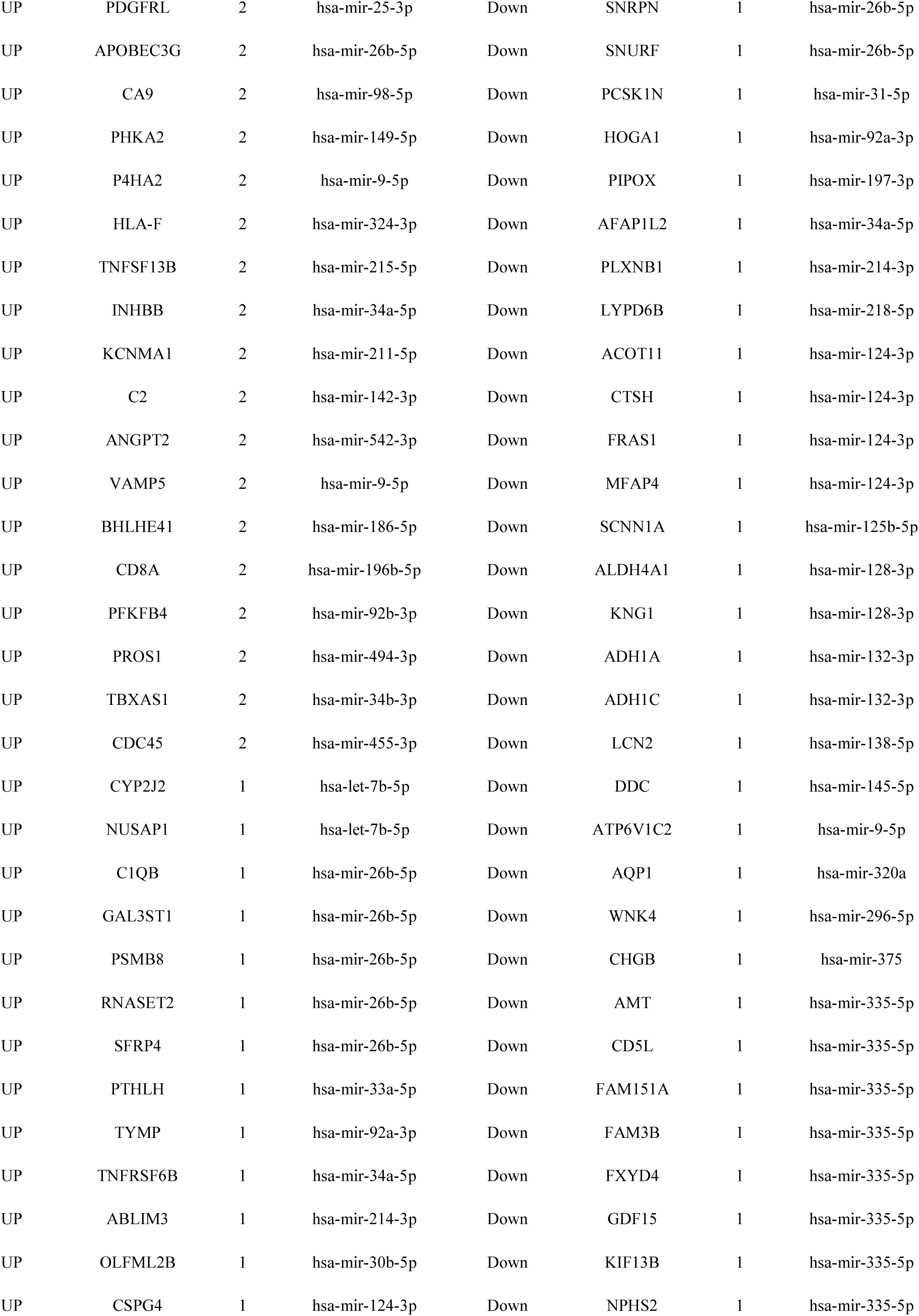
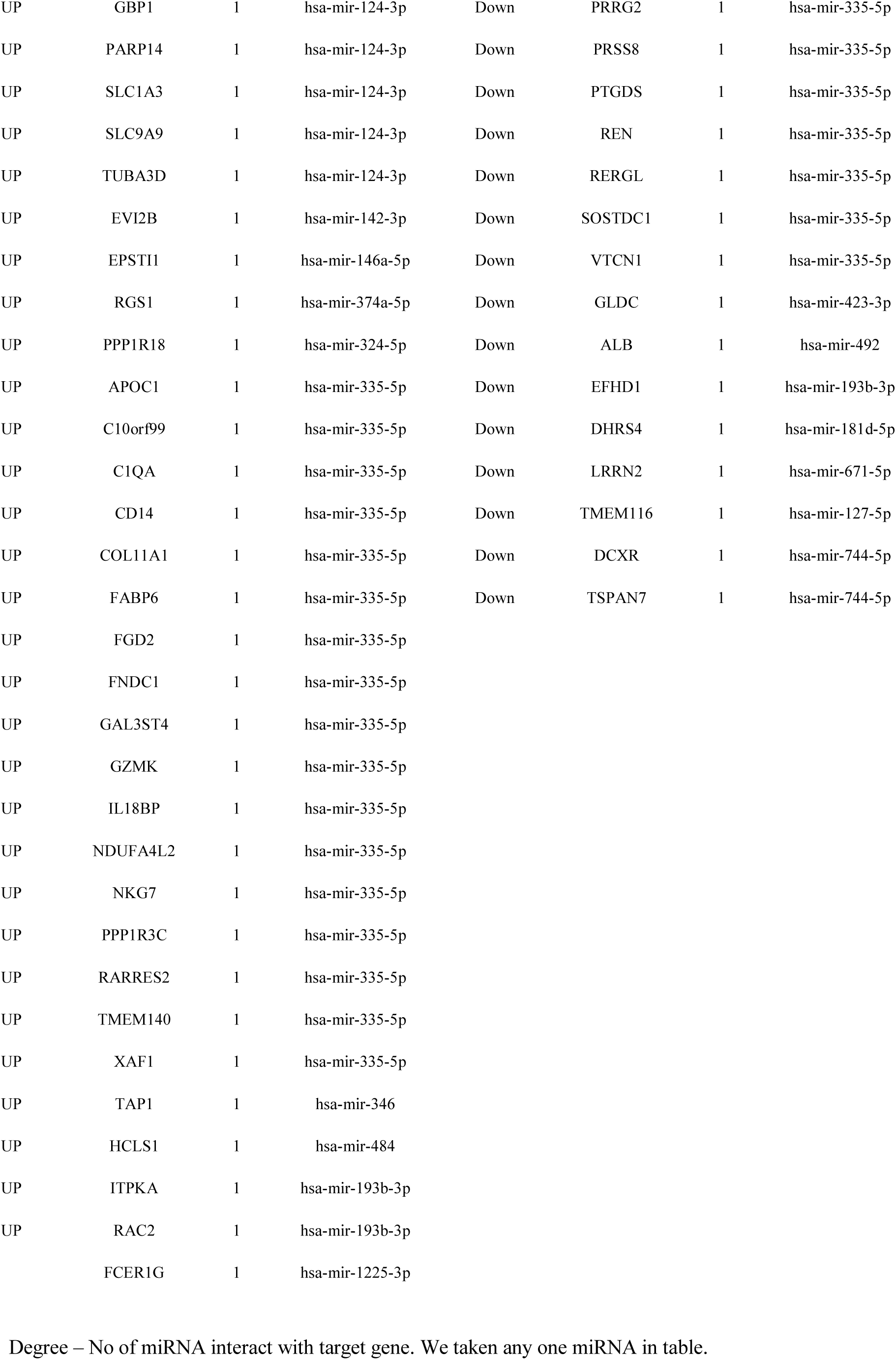
miRNA - target gene interaction table

**Table 9.**
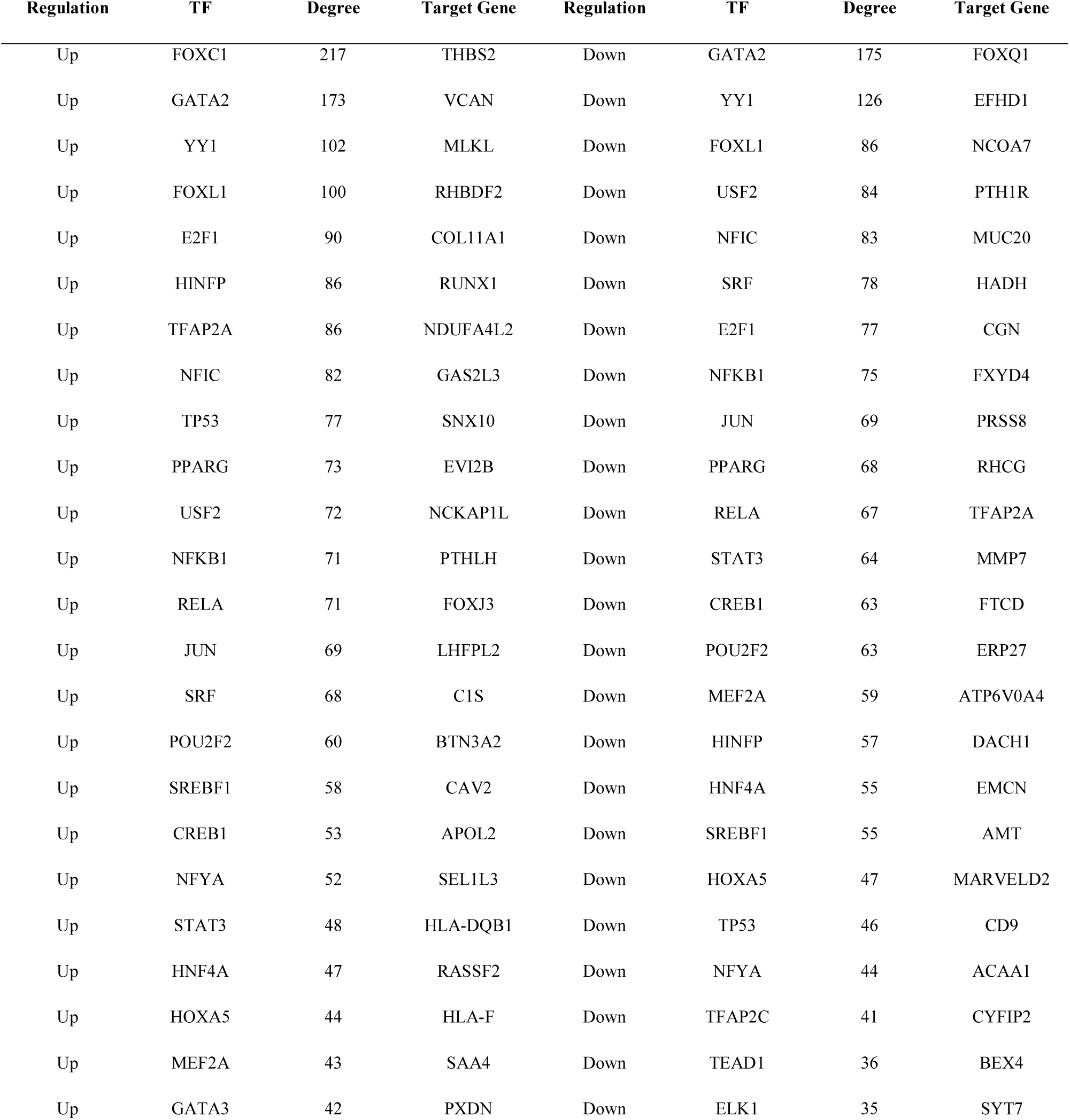
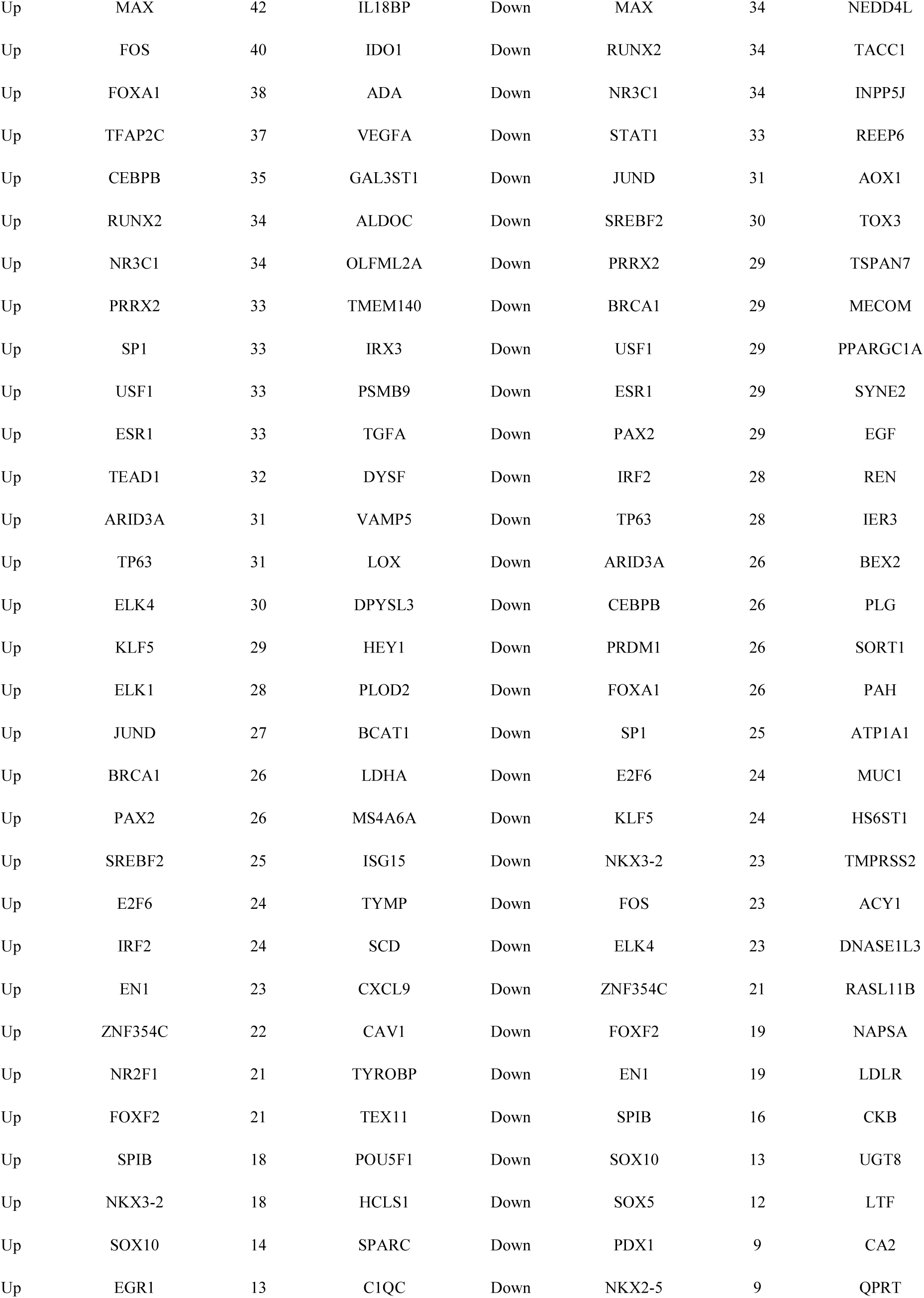
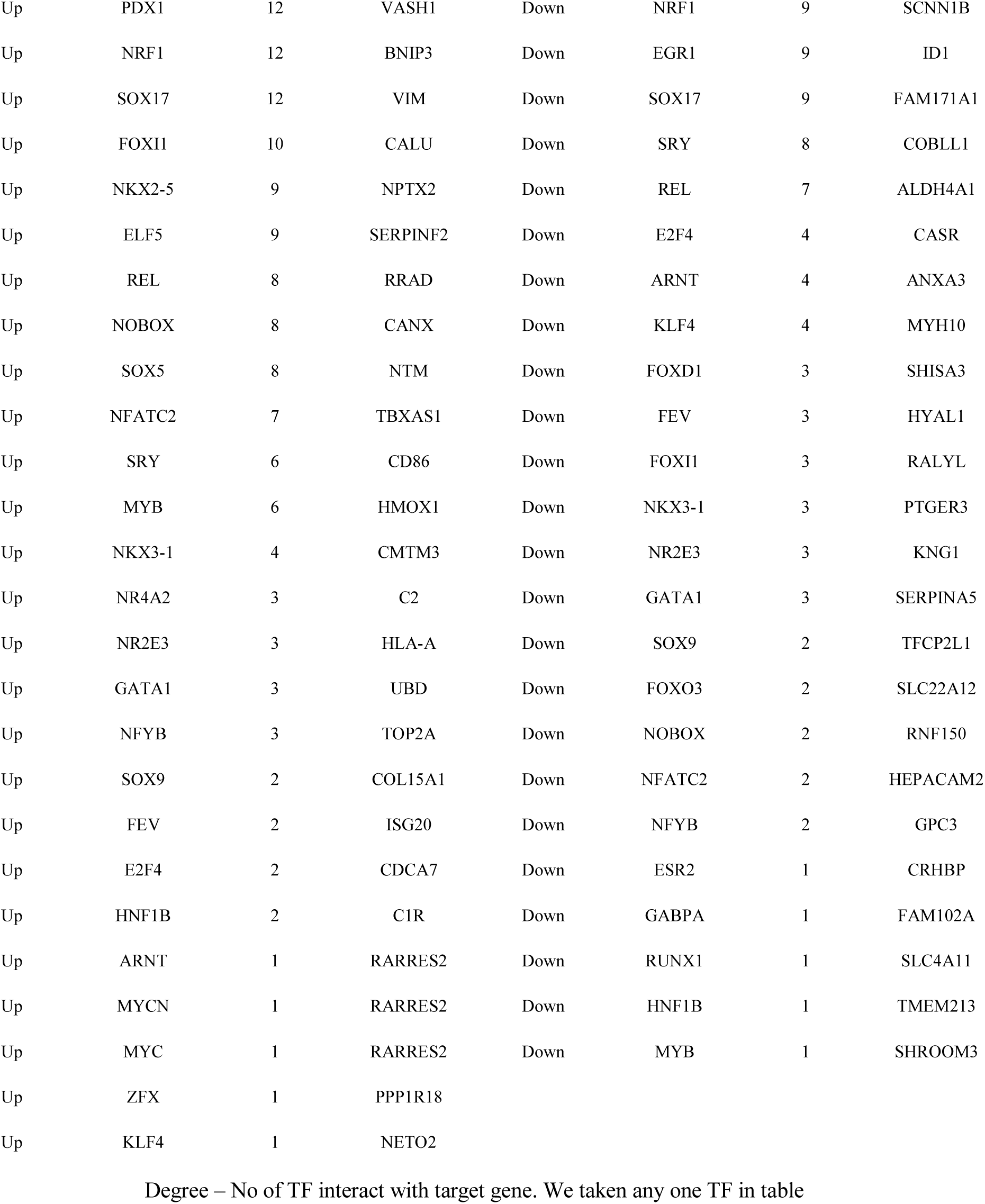
TF - target gene interaction table

### Validation of hub genes

The present study validated the effect of CANX, SHMT2, IFI16, P4HB, CALU, CDH1, ERBB2, NEDD4L, TFAP2A and SORT1 on the prognosis of patients with ccRCC using the UALCAN. According to the median expression level of CANX, SHMT2, IFI16, P4HB, CALU, CDH1, ERBB2, NEDD4L, TFAP2A and SORT1, the patients were divided into a high-expression and a low-expression group. The Kaplan-Meier analysis demonstrated that patients with low CANX expression had shorter survival time than patients with high CANX levels (p = 0.043) (Fig. 15A), high SHMT2 expression had shorter survival time than patients with low SHMT2 levels (p = 0.039) (Fig. 15B), high IFI16 expression had shorter survival time than patients with low IFI16 levels (p < 0.0001) (Fig. 15C), high P4HB expression had shorter survival time than patients with low P4HB levels (p < 0.0001) (Fig. 15D), high CALU expression had shorter survival time than patients with low CALU levels (p = 0.0065) (Fig. 15E), low CDH1 expression had shorter survival time than patients with higher CDH1 levels (p = 0.0037) (Fig. 16A), low ERBB2 expression had shorter survival time than patients with higher ERBB2 levels (p = 6e-04) (Fig. 16B), low NEDD4L expression had shorter survival time than patients with high NEDD4L levels (p = 0.0044) (Fig. 16C), high TFAP2A expression had shorter survival time than patients with low TFAP2A levels (p < 0.0001) (Fig. 16D) and low SORT1 expression had shorter survival time than patients with higher SORT1 levels (p = 0.00099) (Fig. 16E). UALCAN for expression analysis of tumor vs. normal tissue demonstrated that CANX, SHMT2, IFI16, P4HB and CALU were significantly over expressed in ccRCC compared to normal (Fig. 17), while CDH1, ERBB2, NEDD4L, TFAP2A and SORT1 were significantly low expressed in ccRCC compared to normal (Fig. 18). In addition, UALCAN box plots of gene expression by pathological stages based on the TCGA database results shows that CANX, SHMT2, IFI16, P4HB and CALU were more expressed in all stages of ccRCC compared to normal (Fig. 19), while CDH1, ERBB2, NEDD4L, TFAP2A and SORT1 were less expressed in all stages of ccRCC compared to normal (Fig. 20). We used cBioportal tool to identify the specific alterations in hub genes. Percentages of alteration in hub genes such as 8 % CANX (missense mutation, truncating mutation and amplification), 0.3 % SHMT2 (missense mutation), 0.3 % IFI16 (amplification), 0.3% P4HB (amplification), 0.6 % CALU (missense mutation and amplification), 0.3% % CDH1 (missense mutation), 1.1 % ERBB2 (missense mutation, truncating mutation and amplification), 0.6 % NEDD4L (truncating mutation and amplification), 0 % TFAP2A and 0.3 % SORT1 (missense mutation) (Fig.21). In addition, the immunohistochemistry staining obtained from The Human Protein Atlas indicated that the expression of CANX, SHMT2, IFI16, P4HB and CALU were enhanced in ccRCC tissue compared to normal tissue and CDH1, ERBB2, NEDD4L, TFAP2A and SORT1 in ccRCC tissue compared to normal tissue (Fig. 22) and showed consistent conclusions. To further investigate the efficiency of hub genes as potential biomarker of ccRCC, ROC curve between CCRCC patients and controls was performed to analyze the sensitivity and specificity of hub genes in diagnosing ccRCC. The result (Fig. 23) showed that areas under the curve (AUC) of hub genes were CANX (AUC = 0.936), SHMT2 (AUC = 0.915), IFI16 (AUC = 0.932), P4HB (AUC = 0.974), CALU (AUC = 0.944), CDH1 (AUC = 0.880), ERBB2 (AUC = 0.962), NEDD4L (AUC = 0.885), TFAP2A (AUC = 0.799) and SORT1 (AUC = 0.932). The AUC values were more than 0.6, which had great diagnostic value for ccRCC. To verify the reliability of the high-throughput sequencing data, several candidate mRNAs of interest were initially identified for further analysis, including CANX, SHMT2, IFI16, P4HB, CALU, CDH1, ERBB2, NEDD4L, TFAP2A and SORT1. The results demonstrated that CANX, SHMT2, IFI16, P4HB and CALU were up regulated and that CDH1, ERBB2, NEDD4L, TFAP2A and SORT1 were down regulated in ccRCC patients relative to those in the healthy group (Fig. 24). These results demonstrated that the expression patterns of CANX, SHMT2, IFI16, P4HB, CALU, CDH1, ERBB2, NEDD4L, TFAP2A and SORT1 were consistent with those attained by high-throughput sequencing analysis. In the ccRCC, the high expression levels of the CDH1, ERBB2, NEDD4L, TFAP2A and SORT1 all negatively associated with tumor purity (Fig. 25A-E), whereas low expression of CDH1, ERBB2, NEDD4L, TFAP2A and SORT1 were all positively associated with tumor purity (Fig. 25F-J). It can be inferred from this result that all ten hub genes are probably expressed not in the microenvironment, but expressed in the tumor cells.

**Fig. 15.**
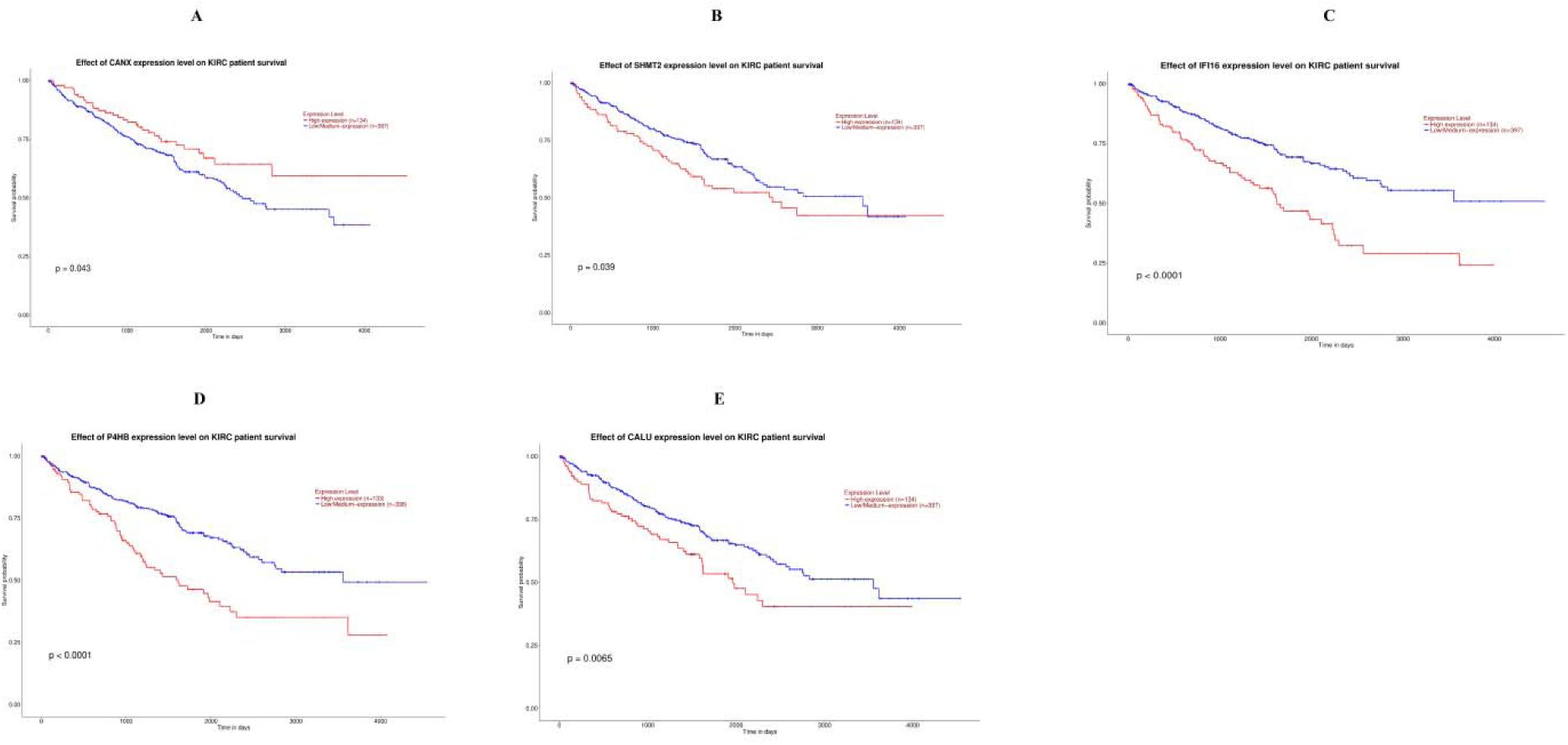
Overall survival analysis of hub genes (up regulated). Overall survival analyses were performed using the UALCAN online platform. A) CANX B) SHMT2 C) IFI16 D) P4HB E) CALU

**Fig. 16.**
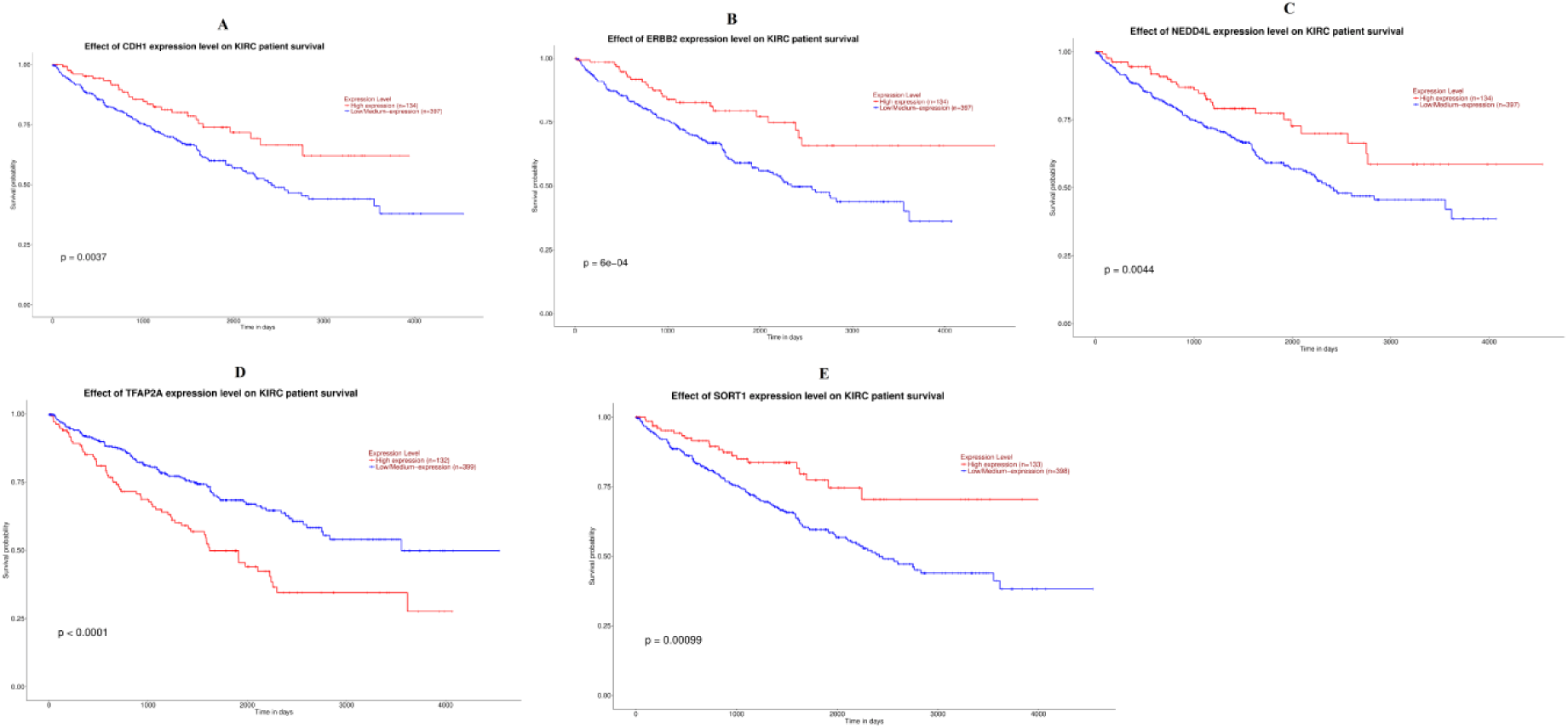
Overall survival analysis of hub genes (down regulated). Overall survival analyses were performed using the UALCAN online platform. A) CDH1 B) ERBB2 C) NEDD4L D) TFAP2A E) SORT1

**Fig. 17.**
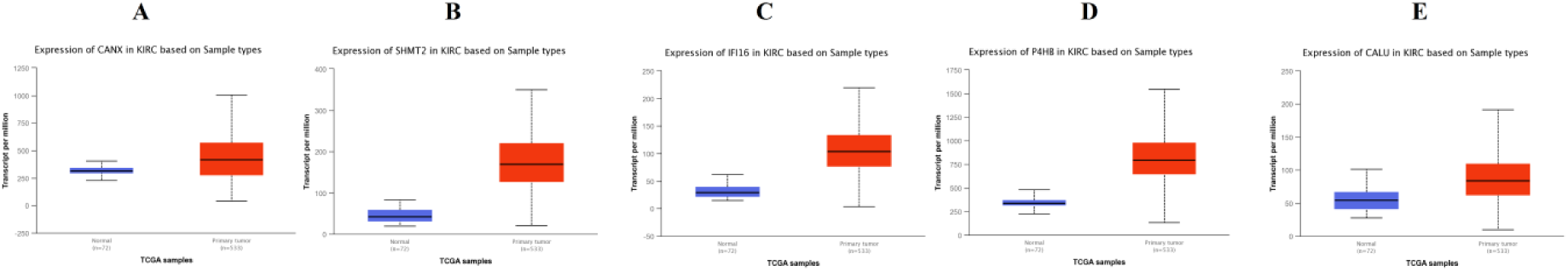
Box plots (expression analysis) hub genes (up regulated genes) were produced using the UALCAN platform. A) CANX B) SHMT2 C) IFI16 D) P4HB E) CALU

**Fig. 18.**
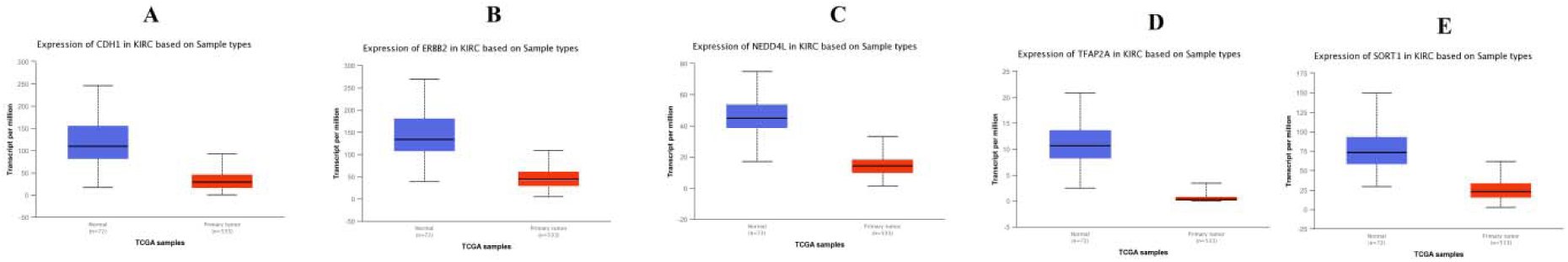
Box plots (expression analysis) hub genes (down regulated genes) were produced using the UALCAN platform. A) CDH1 B) ERBB2 C) NEDD4L D) TFAP2A E) SORT1

**Fig. 19.**
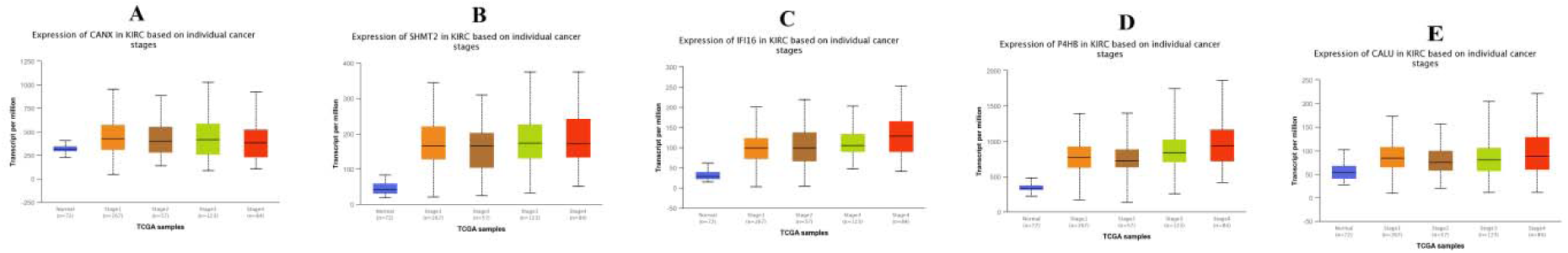
Box plots (stage analysis) of hub genes (up regulated) were produced using the UALCAN platform. A) CANX B) SHMT2 C) IFI16 D) P4HB E) CALU

**Fig. 20.**
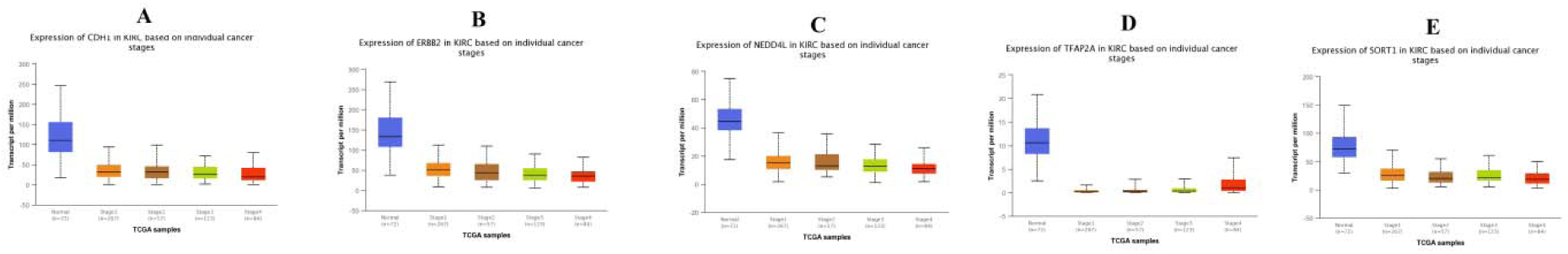
Box plots (stage analysis) of hub genes (down regulated) were produced using the UALCAN platform. A) CDH1 B) ERBB2 C) NEDD4L D) TFAP2A E) SORT1

**Fig. 21.**
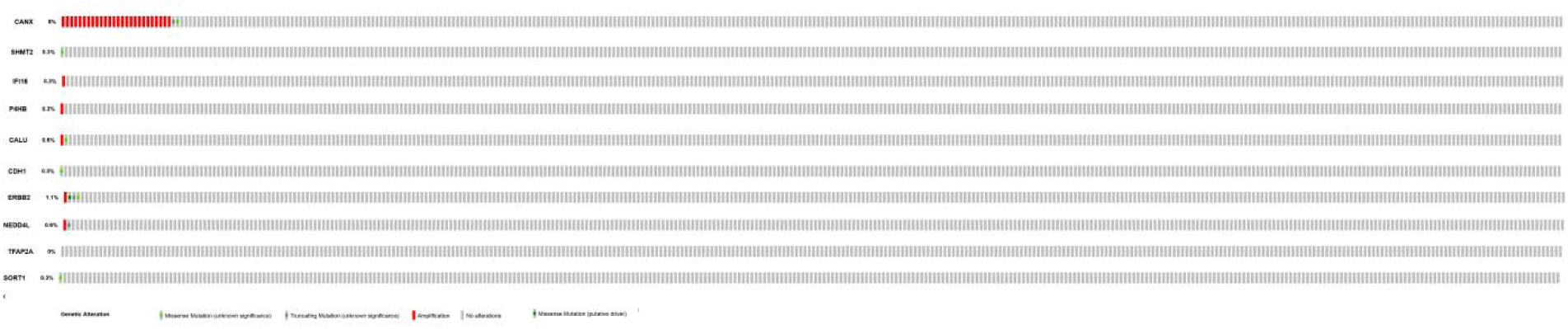
Mutation analyses of hub genes were produced using the CbioPortal online platform

**Fig. 22.**
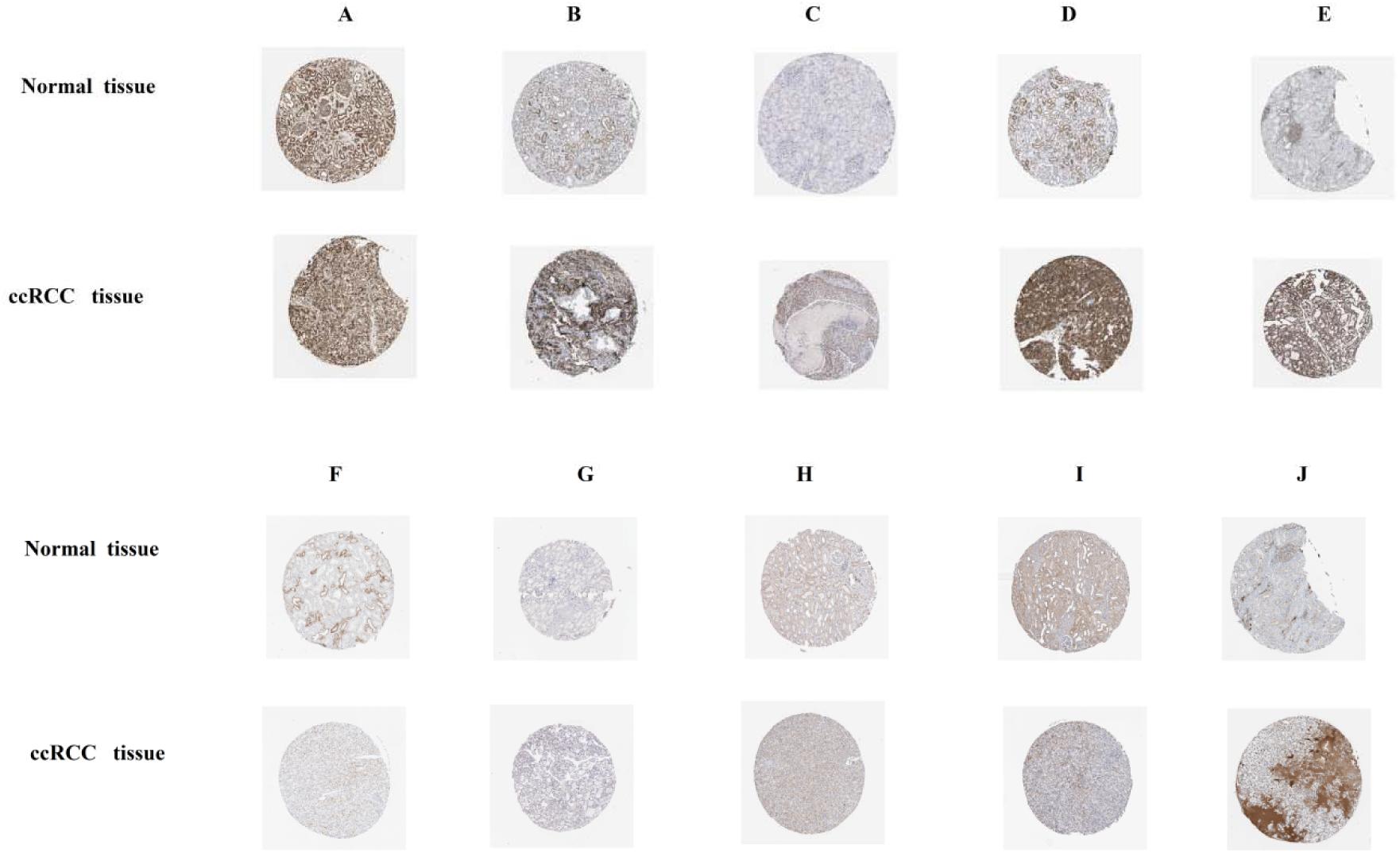
Immuno histochemical analyses of hub genes were produced using the human protein atlas (HPA) online platform. A) CANX B) SHMT2 C) IFI16 D) P4HB E) CALU F) CDH1 G) ERBB2 H) NEDD4L I) TFAP2A J) SORT1

**Fig. 23.**
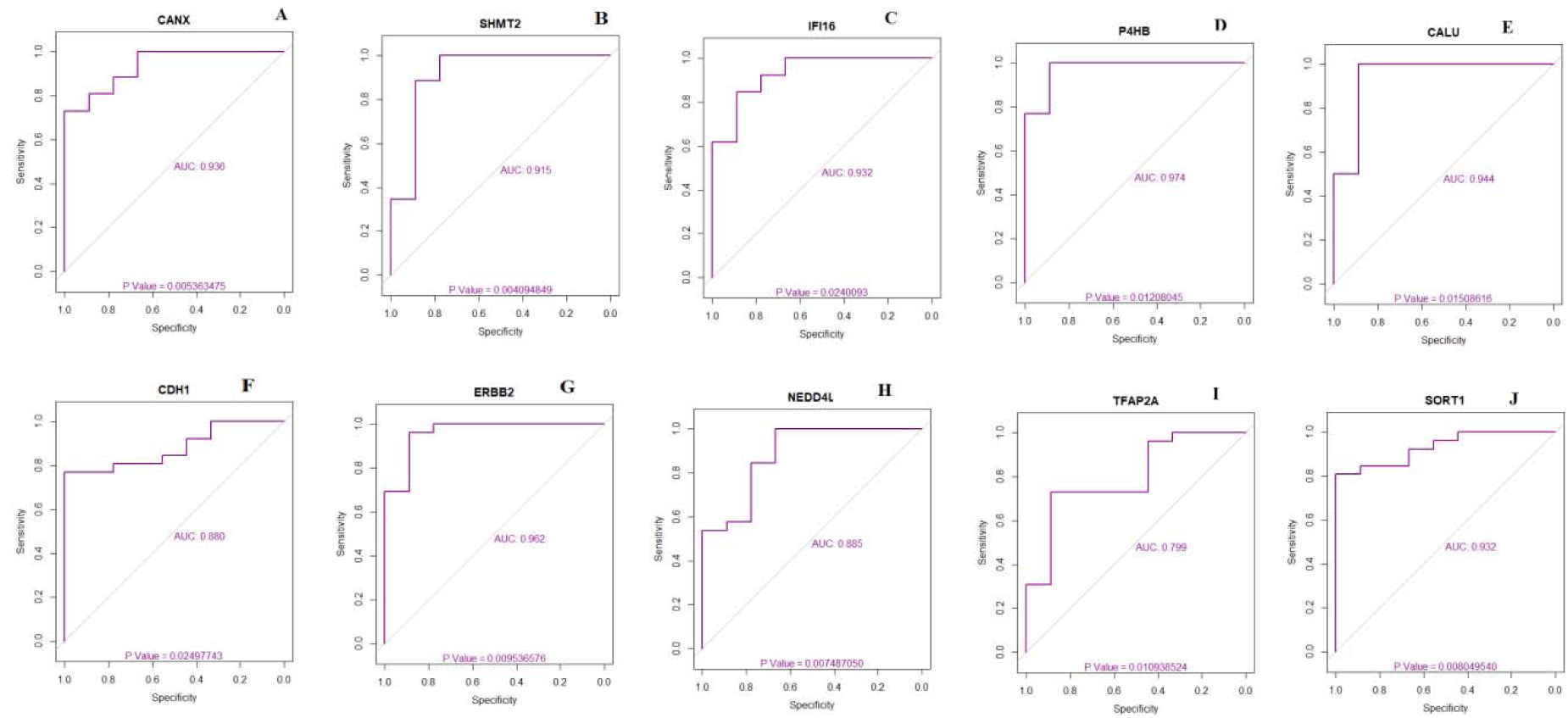
ROC curve validated the sensitivity, specificity of hub genes as a predictive biomarker for GBM prognosis. A) CANX B) SHMT2 C) IFI16 D) P4HB E) CALU F) CDH1 G) ERBB2 H) NEDD4L I) TFAP2A J) SORT1

**Fig. 24.**
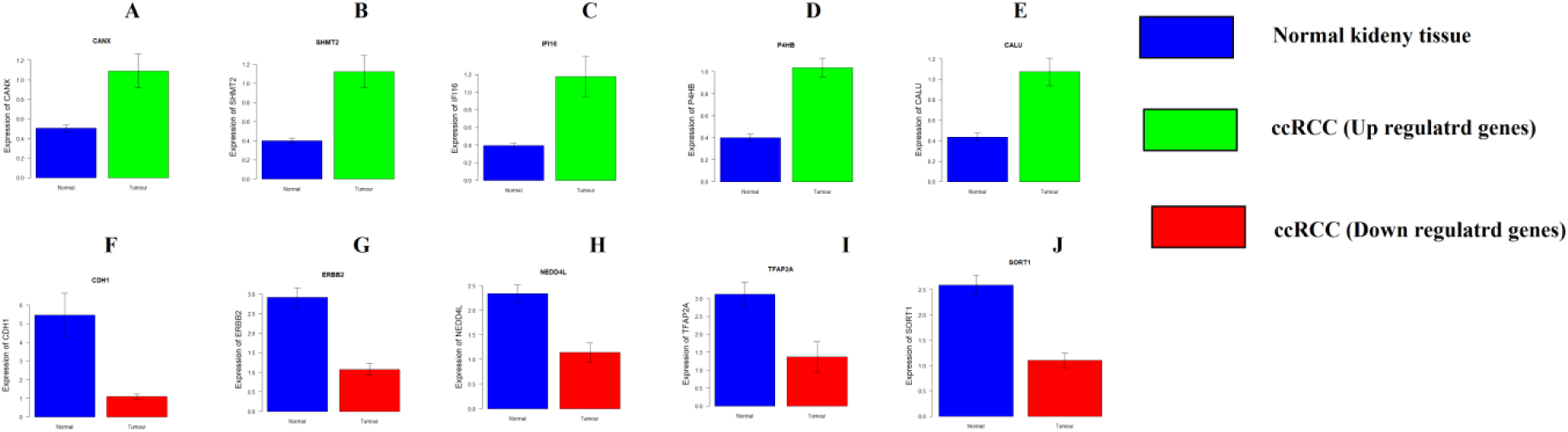
Validation of microarray data by quantitative PCR. A) CANX B) SHMT2 C) IFI16 D) P4HB E) CALU F) CDH1 G) ERBB2 H) NEDD4L I) TFAP2A J) SORT1

**Fig. 25.**
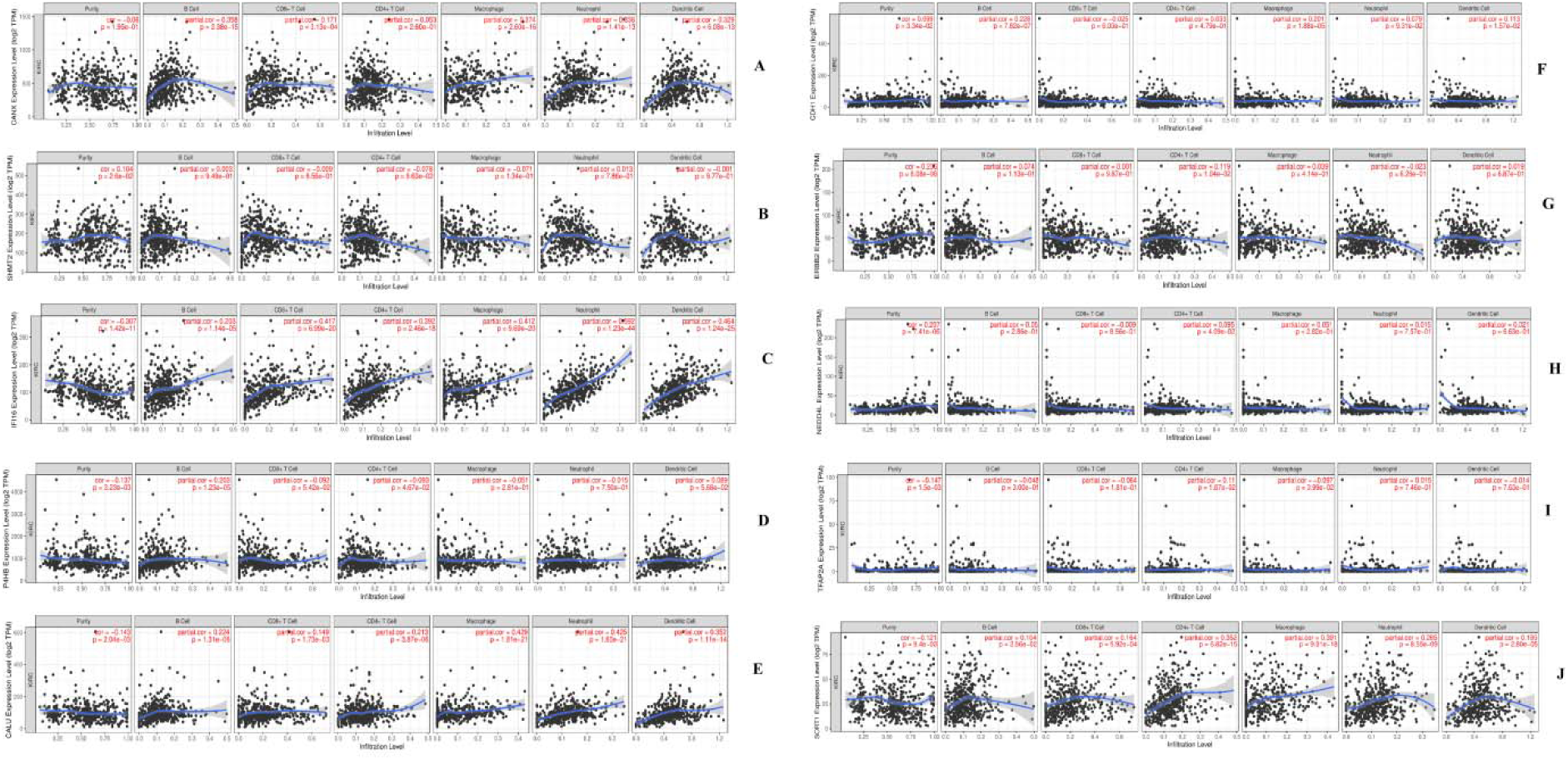
Scatter plot for Immune infiltration of A) ADAM15 B) BATF C) NOTCH3 D) ITGAX E) SDC1 F) RPL4 G) EEF1G H) RPL3 I) RBMX J) ABCC2 in renal cancer

## Discussion

ccRCC is a common malignant neoplasm of the kidney [Gerlinger et al 2014]. The histological type of disease, determined after surgery and chemotherapy, has a key impact on the survival and prognosis of patients with ccRCC. Pathologically determined cancer usually leads to a poor prognosis with metastasis and more difficult surgery or chemotherapy. Furthermore, the 5-year survival rate only reaches 10% [Li et al 2016]. Consequently, a comprehensive understanding of the molecular mechanism behind the occurrence and progression of ccRCC is of essential for management and therapy, as is the progression of new targeted therapies to progress over all survival time. With the popular use of high through output sequencing technologies and microarrays, the expression levels of genes in the human genome are openly incorporated in public databases. Therefore, there is an insistent need for sensitive and new biomarkers of ccRCC and also microarray data can be scrutinized in order to promote the genetic research of ccRCC.

In the current study, based on the original microarray data, we identified 930 DEGs, including 469 up regulated and 461 down regulated genes between metastatic ccRCC tissues and normal renal tissues. Genes such as CKMT1A [Fan et al 2018] and P4HA1 [Zhou et al 2017] were associated with invasion of various cancer cells, but these genes may be liable for invasion of ccRCC cells. Genes such as RAB25 [Liu and Ding, 2014], KCNJ1 [Guo et al 2015], SFRP1 [Ricketts et al 2014], CLDN8 [Osunkoya et al 2009], ATP1A1 [Zhang et al 2017], NDUFA4L2 [Wang et al 2017], SLC16A3 [Fisel et al 2013], ANGPTL4 [Dong et al 2017] and PLOD2 [Kurozumi et al 2016] were responsible for progression of ccRCC.

In pathway enrichment analysis, up regulated genes were enriched in dTMP de novo biosynthesis, phagosome, HIF-1-alpha transcription factor network, extracellular matrix organization, gluconeogenesis, ensemble of genes encoding extracellular matrix and extracellular matrix-associated proteins, integrin signalling pathway and hypertension. Genes such as TYMS (thymidylatesynthetase) [Colavito et al 2009], HLA-A (major histocompatibility complex, class I, A) [Schendel et al 1997], HLA-B (major histocompatibility complex, class I, B) [Ibrahim et al 2003], HLA-G (major histocompatibility complex, class I, G) [Seliger and Schlaf, 2007], TAP1 [Dovhey et al 2000], CD14 [Li et al 2014], SCARB1 [Pośpiech et al 2015], HK2 [Yoshino et al 2013], PLIN2 [Cao et al 2018], ADM (adrenomedullin) [Gao et al 2016], VEGFA (vascular endothelial growth factor A) [Dorević et al 2009], BHLHE41 [Bigot et al 2016], BNIP3 [Huang et al 2014], CXCR4 [D’Alterio et al 2010], CA9 [Bleumer et al 2007], LDHA (lactate dehydrogenase A) [Girgis et al 2014], MMP9 [Liang et al 2012], VCAM1 [Shioi et al 2006], COL3A1 [Su et al 2014], COL6A1 [Wan et al 2015], COL23A1 [Xu et al 2018], VCAN (versican) [Mitsui et al 2017], LAMA4 [Wragg et al 2016], SERPINH1 [Qi et al 2018], TIMP1 [Lu et al 2014], LOX (lysyl oxidase) [Di Stefano et al 2016], PFKP (phosphofructokinase, platelet) [Wang et al 2016], CTHRC1 [Jin et al 2016], ANGPT2 [Gayed et al 2015], POSTN (periostin) [Chuanyu et al 2017], IGFBP3 [Ricketts et al 2009], CXCL10 [Polimeno et al 2013], ESM1 [Leroy et al 2010], LGALS1 [White et al 2017], TGFA (transforming growth factor alpha) [Argilés et al 1992], TGFBI [Lebdai et al 2015], TGM2 [Erdem et al 2014], CAV1 [Butz et al 2015], SOD2 [Gao et al 2013] and GAPDH (glyceraldehyde-3-phosphate dehydrogenase) [Vilà et al 2000] were responsible for progression of ccRCC. SHMT2 was associated with pathogenesis of hepatocellular carcinoma [Woo et al 2016], but expression of this gene may be important for advancement of ccRCC. SNP in genes such as HLA-DQA1 [Kohno et al 2011], HLA-DQB1 [Chan et al 2007], FCGR2A [Gavin et al 2017], THBS4 [Lin et al 2016], C1QA [Azzato et al 2010] and RAC2 [Hertz et al 2016] were responsible for advancement of various cancer, but these polymorphic genes may be liable for progression of ccRCC. Genes such as ITGA5 [Yoo et al 2016], ITGB2 [Liu et al 2018], C3 (complement C3) [Chen et al 2013], THBS2 [Wei et al 2017], CD36 [Ladanyi et al 2018], HMOX1 [Ghosh et al 2016], SERPINE1 [Azimi et al 2017], PLOD1 [Li et al 2017], COL1A1 [Zhang et al 2018], COL1A2 [Ao et al 2018], COL4A1 [Huang et al 2018], COL4A2 [JingSong et al 2017], COL5A1 [Liu et al 2018], COL5A2 [Zeng et al 2018], COL6A3 [Ao et al 2018], COL8A1 [Ma et al 2012], BMP1 [Wu et al 2014], P4HB [Xia et al 2017], ALDOC (aldolase, fructose-bisphosphate C) [Li et al 2016], PGF (placental growth factor) [Zhang et al 2015], SERPINE2 [Wang et al 2015], CXCL9 [Bronger et al 2016], FGF11 [Ye et al 2016], CCL5 [Mi et al 2011], SEMA6A [Lim et al 2017], VWF (von Willebrand factor) [Wang et al 2005], CTSZ (cathepsin Z) [Lines et al 2012], INHBB (inhibin subunit beta B) [Zou et al 2018], FNDC1 [Das and Ogunwobi, 2017], PXDN (peroxidasin) [Zheng and Liang, 2018] and FSTL1 [Yang et al 2017] were important for invasion of various cancer cells, but these genes may be involved in invasion of ccRCC cells. High expression of genes such as CANX (calnexin) [Kobayashi et al 2015], OLR1 [Wang et al 2017], CP (ceruloplasmin) [Balmaña et al 2015], SLC2A1 [Santasusagna et al 2018], FGG (fibrinogen gamma chain) [Zhu et al 2009], ENO2 [Soh et al 2011], C1QTNF6 [Takeuchi et al 2011], TNFSF13B [Qian et al 2014], CSPG4 [Poli et al 2013] and C1QL1 [Celestino et al 2018] were responsible for pathogenesis of various cancer, but over expression of these genes may be linked with progression of ccRCC. Genes such as COL11A1 [Wu et al 2015], PLOD3 [Baek et al 2019] and SFRP4 [Warrier et al 2014] were associated with drug resistance in various cancer, but these genes may be linked with chemo resistance in ccRCC. HLA-DQA2, HLA-F (major histocompatibility complex, class I, F), FCGR1A, CYBB (cytochrome b-245 beta chain), C1R, TUBA3D, EGLN3, COL6A2, COL15A1, SPARC (secreted protein acidic and cysteine rich), P4HA2, COLGALT1, TPI1, SEMA5B, SERPINF2, ANXA4, CCL4L2, FCN3, C1QB, C1QC and TNFAIP6 were identified as novel biomarkers for pathogenesis of ccRCC in these pathways. Similarly, down regulated genes were enriched in 4-hydroxyproline degradation, metabolic pathways, FOXA2 and FOXA3 transcription factor networks, transmembrane transport of small molecules, urea cycle and metabolism of amino groups, platelet amyloid precursor protein pathway, blood coagulation, pathway of urea cycle and metabolism of amino groups and polythiazide pathway. Low expression of genes such as GPHN (gephyrin) [Zhang et al 2019], ASS1 [Qiu et al 2014], INPP5J [Zhu et al 2015], PCK1 [Liu et al 2018] and NEDD4L [Qu et al 2016] were linked with development of various cancer, but less expression of these genes may be liable for progression of ccRCC. Genes such as EPHX2 [Vainio et al 2011], ACY1 [Shi et al 2013] CKB (creatine kinase B) [Li et al 2013], UGT8 [Cao et al 2018], ALDOB (aldolase, fructose-bisphosphate B) [Tian et al 2017], HYAL1 [Tan et al 2011], G6PC [Guo et al 2015], GLDC (glycine decarboxylase) [Berezowska et al 2017], QPRT (quinolinatephosphoribosyltransferase) [Hinsch et al 2009], AQP6 [Ma et al 2016], SLC12A1 [Teng et al 2016], SLC44A4 [Mattie et al 2016], NIPAL1 [Sasahira et al 2018], PLG (plasminogen) [Stillfried et al 2007], COL4A3 [Jiang et al 2013] and KNG1 [Quesada-Calvo et al 2017] were important for progression of various cancer, but these genes may be linked with pathogenesis of ccRCC. Genes such as HMGCS2 [Su et al 2017], PTGDS (prostaglandin D2 synthase) [Shyu et al 2013] and PLAT (plasminogen activator, tissue type) [Strojan et al 1998] were involved in invasion of various cancer cells, but these genes may be responsible for invasion of ccRCC cells. Single nucleotide polymorphism (SNP) in genes such as ADH1A [Cui et al 2018] and ADH1C [Hidaka et al 2015] were linked with advancement of gastric cancer, but these polymorphic genes may be associated with pathogenesis of ccRCC. Genes such as ATP6V1G3 [Shinmura et al 2015], FBP1 [Ning et al 2016], ARG2 [Ochocki et al 2018], GATM (glycine amidinotransferase) [Pflueger et al 2015], DDC (dopa decarboxylase) [Papadopoulos et al 2015], PHGDH (phosphoglycerate dehydrogenase) [Yoshino et al 2017], MAN1C1 [Li et al 2018], ALB (albumin) [Koparal et al 2018], ABCB1 [Diekstra et al 2015], AQP1 [Morrissey et al 2015], SLC47A2 [Gao et al 2019], FXYD2 [Gaut et al 2013] and LCN2 [Perrin et al 2011] were important for pathogenesis of ccRCC. Methylation inactivation of tumor suppressor genes such as ST6GAL1 [Antony et al 2014], OGDHL (oxoglutarate dehydrogenase like) [Khalaj-Kondori et al 2019], ALDH1A2 [Kim et al 2005], SCNN1B [Qian et al 2017] and RHCG (Rh family C glycoprotein) [Ming et al 2018] were liable for advancement of various cancer, but loss of these genes may be important for progression of ccRCC. PRODH2, HOGA1, ALDH4A1, CDS1, ACAA1, GGT6, PIPOX (pipecolic acid and sarcosine oxidase), ACOX2, CKMT1B, CKMT2, HPD (4-hydroxyphenylpyruvate dioxygenase), PLA2G4F, ATP6V1C2, ATP6V0D2, PLCG2, ALDH6A1, AMT (aminomethyltransferase), AOX1, CYP4F2, ATP6V0A4, SORD (sorbitol dehydrogenase), ATP6V1B1, CYP4A11, DHRS4L1, FTCD (formimidoyltransferasecyclodeaminase), SUCLG1, HSD17B8, RDH10, GPAT3, PAH (phenylalanine hydroxylase), TM7SF2, AGMAT (agmatinase), HADH (hydroxyacyl-CoA dehydrogenase), PCCA (propionyl-CoA carboxylase subunit alpha), DCXR (dicarbonyl and L-xylulosereductase), SLC29A2, SLC22A6, SLC22A8, CLCNKA (chloride voltage-gated channel Ka), CLCNKB (chloride voltage-gated channel Kb), SCNN1A, SCNN1G, SLC13A3, AQP2, SLC22A12, SLC4A1, SLC6A12, SLC9A2, SLC12A3, SLC13A1, SLC26A7, SLC34A1, FXYD4, SLC30A2, SLC22A7, SLC36A2, WNK4, SLC7A8 and COL4A4 were identified as novel biomarkers for pathogenesis of ccRCC in these pathways.

In GO enrichment analysis, up regulated genes were enriched in response to cytokine, vesicle organization and signaling receptor binding. Genes such as IL32 [Lee et al 2012], FCER1G [Chen et al 2017], VIM (vimentin) [Yamasaki et al 2012], WAS (Wiskott-Aldrich syndrome) [Liu et al 2015], CSF1R [Soares et al 2009], XAF1 [Kempkensteffen et al 2009], FSCN1 [Zhang et al 2018], AXL (AXL receptor tyrosine kinase) [Zucca et al 2018], CCND1 [Sukov et al 2009], STAT1 [Zhu et al 2012], ACKR3 [Mahmoodi et al 2017], RUNX1 [Xiong et al 2014], CD70 [Ruf et al 2015], ACLY (ATP citrate lyase) [Teng et al 2018], ENPP3 [Thompson et al 2018], DYSF (dysferlin) [Ha et al 2019], CD163 [Ma et al 2018], CAV2 [Yamasaki et al 2013], CMTM3 [Xie et al 2014], PDGFRB (platelet derived growth factor receptor beta) [Shim et al 2015], SLC6A3 [Hansson et al 2017], STC2 [Ma et al 2015], PTHLH (parathyroid hormone like hormone) [Yao et al 2014], NRP1 [Cao et al 2008], EDNRB (endothelin receptor type B) [Wuttig et al 2012], CD2 [Tienari et al 2005] and NETO2 [Oparina et al 2012] were responsible for progression of ccRCC. Methylation inactivation of tumor suppressor genes such as MSC (musculin) [Pirini et al 2017] and TRIM9 [Mishima et al 2015] were important for progression of various cancer, but loss of these genes may be liable for advancement of ccRCC. Elevated expression of genes such as SAA1 [Kim et al 2015], UBD (ubiquitin D) [Zhao et al 2015], PARP14 [Iansante et al 2015], IFI16 [Alimirah et al 2007], ISG15 [Burks et al 2015], PLVAP (plasmalemma vesicle associated protein) [Wang et al 2014], TNFRSF6B [Chen et al 2010], SIRPA (signal regulatory protein alpha) [Nagahara et al 2015], IL18BP [Carbotti et al 2013], PRDX4 [Ummanni et al 2012], APOC1 [Su et al 2018], TAPBP (TAP binding protein) [Shao et al 2013], RAB31 [Sui et al 2015], SAA2 [Kim et al 2015], C10orf99 [Pan et al 2014], LAG3 [Huang et al 2016] and TYMP (thymidine phosphorylase) [Derwinger et al 2013] were associated with development of various cancer, but increase expression of these genes may be involved in progression of ccRCC. Genes such as SNX10 [Cervantes-Anaya et al 2017], LAPTM5 [Chen et al 2017], ABCA1 [Chen et al 2017], RNASET2 [Acquati et al 2011], RARRES2 [Liu-Chittenden et al 2017] and BTN3A1 [Zocchi et al 2017] were liable for pathogenesis of various cancer, but expression of these genes may be linked with development of ccRCC. Genes such as BIRC3 [Mendoza-Rodríguez et al 2017] and BST2 [Kuang et al 2017] were responsible for drug resistance in various cancer, but these genes may be associated with chemo resistance in ccRCC. Genes such as POU5F1 [Cai et al 2016], PSMB8 [Yang et al 2018], GBP1 [Li et al 2015], GBP2 [Zhang et al 2018], DPYSL3 [Yang et al 2018], CORO1C [Lim et al 2017], FABP5 [Wang et al 2016], EHD2 [Kim et al 2017], HSPA6 [Shin et al 2017], SLC2A5 [Weng et al 2018], PROS1 [Che Mat et al 2016], DOCK2 [El Haibi et al 2010], CD68 [Eiró et al 2012], NCKAP1L [Xiong et al 2019] and GJA1 [Busby et al 2018] were liable for invasion of various cancer cells, but these genes may be important for invasion of ccRCC cells. SNP in genes such as IL2RB [Jia et al 2019], CD86 [Azimzadeh et al 2013] and BTN3A2 [Zhu et al 2017] were associate with advancement of various cancer, but these polymorphic genes may be liable for pathogenesis of ccRCC. HLA-H (major histocompatibility complex, class I, H (pseudogene)), IFI27, RHEX (regulator of hemoglobinization and erythroid cell expansion), PSMB9, IL20RB, ISG20, GBP5, NOL3, OAS2, LCP1, HCLS1, TYROBP (TYRO protein tyrosine kinase binding protein), PLEK (pleckstrin), NLGN1, SLC15A4, PYGL (glycogen phosphorylase L), LHFPL2, STEAP3, ADGRE5, LY96, SAA4, HILPDA (hypoxia inducible lipid droplet associated), APOL2, FYB1, CD8A and PRAME (preferentially expressed antigen in melanoma) were identified as novel biomarkers for pathogenesis of ccRCC in these GO categories. Similarly, down regulated genes were enriched in in renal system development, apical part of cell and ion transmembrane transporter activity. Methylation inactivation of tumor suppressor genes such as CDKN1C [Yang et al 2009] and MAL (mal, T cell differentiation protein) [Suzuki et al 2013] were linked with progression of various cancer, but loss of these genes may be liable for development of ccRCC. Genes such as MECOM (MDS1 and EVI1 complex locus) [Hou et al 2016], HPGD (15-hydroxyprostaglandin dehydrogenase) [Fink et al 2014], PROM1 [Qiu et al 2015], CA2 [Luo et al 2017], ACPP (acid phosphatase, prostate) [Chuang et al 2010], MUC20 [Chen et al 2016], IGFBP2 [Zumkeller et al 1993], PTH1R [Liang et al 2012], REEP6 [Park et al 2016], DPEP1 [Tachibana et al 2017], LDLR (low density lipoprotein receptor) [Caruso et al 2002], MAL2 [Li et al 2017], HSPA2 [Zhang et al 2015], CLDN16 [Kuo et al 2010] and SLC4A11 [Qin et al 2017] were important for advancement of various cancer, but expression of these genes may be important for advancement of ccRCC. Genes such as HOXD11 [Sharpe et al 2014], FOXC1 [Han et al 2015], CTSH (cathepsin H) [Jevnikar et al 2013], TACSTD2 [Noorlag et al 2015], IRX2 [Berinstein et al 2012], ERBB2 [Kondratyev et al 2012], CLIC5 [Flores-Téllez et al 2015], ANK2 [Cao et al 2018], ANXA3 [Zhou et al 2017], MPC1 [Zhou et al 2017] and PPARGC1A [Li et al 2017] were involved in the invasion of various cancers cells, but these genes may be associated with invasion of ccRCC. Genes such as GATA3 [Cooper et al 2010], GPC3 [Valsechi et al 2014], TCF21 [Gooskens et al 2015], REN (renin) [Deckers et al 2015], EPCAM (epithelial cell adhesion molecule) [Zimpfer et al 2014], MUC1 [Aubert et al 2009], CEACAM1 [Kammerer et al 2004], CASR (calcium sensing receptor) [Joeckel et al 2014], CD9 [Garner et al 2016], CDH1 [Ricketts et al 2009], CRHBP (corticotropin releasing hormone binding protein) [Tezval et al 2016] and KCNJ15 [Liu et al 2019] were responsible for development of CCRCC. Decrease expression of genes such as RBP4 [Karunanithi et al 2017], EMX2 [Qiu et al 2013], RAB17 [Qi et al 2015], SLC25A25 [Li et al 2016] and PRSS8 [Zhang et al 2016] were associated with advancement of various cancer, but low expression of these genes may be important for pathogenesis of ccRCC. Genes such as TFAP2A [Tang et al 2017] and SLC27A2 [Chen et al 2018] were associated with chemo resistance in various cancer such as gastric cancer and ovarian cancer, but these genes may be linked with drug resistance in ccRCC. SNP in SLC16A5 was linked with advancement of testicular cancer [Drögemölle et al 2017], but this polymorphic gene may be liable for progression of ccRCC. UMOD (uromodulin), FRAS1, KLHL3, NPHS2, PROM2, RAPGEF3, SHROOM3, TJP3, CA4, MARVELD2, SCN2A, SLC25A4, TMC4 and GPD1L were identified as novel biomarkers for pathogenesis of ccRCC in these GO categories.

In PPI network, up regulated genes such as EGLN3, PCLAF (PCNA clamp associated factor), VCAM1, CANX, SHMT2, IFI16 and GAPDH were identified as hub genes showing the highest node degree, betweenness, stress and closeness. Up regulated genes such as MS4A7, FGF11, IKBIP (IKBKB interacting protein), SCD (stearoyl-CoA desaturase) and TRIM9 were identified as hub genes showing the lowest clustering coefficient. High expression of SCD was linked with progression of prostate cancer [Kim et al 2011], but increase expression of this gene may be liable for advancement of ccRCC. PCLAF, MS4A7 and IKBIP were identified as novel biomarkers for pathogenesis of ccRCC. Similarly, down regulated genes such as CDH1, ERBB2, ATP1A1, ALB, NEDD4L, PHGDH and RTN4 were identified as hub genes showing the highest node degree, betweenness, stress and closeness. Down regulated genes such as ACOT11, ANKRD9, VTCN1, SLC22A7 and MST1L were identified as hub genes showing the lowest clustering coefficient. RTN4 was important for pathogensis for ccRCC [Pu et al 2018]. VTCN1 was associated with invasion of bladder cancer cells [Chand et al 2019], but this gene may be responsible for invasion of ccRCC cells. ACOT11, ANKRD9 and MST1L were identified as novel biomarkers for pathogenesis of ccRCC.

In module analysis, up regulated genes such as PCLAF, GAPDH, ENO2, TPI1, VIM, SERPINH1, P4HB, LGALS1, STAT1, CAV1, VEGFA, PDGFRB, COL6A1, SPARC, COL6A2, BMP1, PLOD1, COL1A2, MMP9, TIMP1, COL5A1 and COL1A1 were identified as hub genes showing the highest node degree in all four significant modules. Similarly, down regulated genes such as ERBB2, PLCG2, PLXNB1, KIT, CDH1, RTN4, CGN, SHROOM3, LAD1, NEDD4L, SCNN1A, SCNN1G, SCNN1B, USP2, MATN2, KLK6, COL4A3 and COL4A4 were identified as hub genes showing the highest node degree in all four significant modules. Genes such as KIT (KIT proto-oncogene receptor tyrosine kinase) [Wang and Mills, 2005] and LAD1 [Van Vlodrop et al 2017] were involved in pathogensis of ccRCC. Genes such as PLXNB1 [Worzfeld et al 2012] and USP2 [Qu et al 2015] were assocated with invasion of breast cancer cells, but these genes may be involved in invasion of ccRCC cells. KLK6 was associated with progression of breast cancer [Sidiropoulos et al 2016], but this gene may be involved in pathogensis of ccRCC. PLXNB1, CGN (cingulin) and MATN2 were identified as novel biomarker for pathogenesis of ccRCC.

In target gene - miRNA network, up regulated genes such as SOD2, CCND1, SCD, VEGFA and SH3PXD2A were identified as target genes showing the highest number of integration with miRNAs. Methylation inactivation of tumor supressor SH3PXD2A was liable for advancement of colorectal cancer [Ma et al 2018], but loss of this gene may be responsible for progression of ccRCC. Similarly, down regulated genes such as BTG2, VAV3, LDLR, FOXC1 and SYT7 were identified as target genes showing the highest number of integration with miRNAs. BTG2 was responsible for pathogensis of CCRCC [Struckmann et al 2004]. Genes such as VAV3 [Jiang et al 2017] and SYT7 [Liu et al 2019] were important for invasion of various cancer cells such as breast cancer and lung cancer, but these genes may be associated with invasion of ccRCC cells.

In target gene - TF network, up regulated genes such as THBS2, VCAN, MLKL (mixed lineage kinase domain like pseudokinase), RHBDF2 and COL11A1 were identified as target genes showing the highest number of integration with TFs. RHBDF2 was important for invasion of gastric cancer cells [Ishimoto et al 2017], but this gene may be liable for invasion of ccRCC cells. MLKL was identified as novel biomarker for pathogenesis of ccRCC. Similarly, down regulated genes such as FOXQ1, EFHD1, NCOA7, PTH1R and MUC20 were identified as target genes showing the highest number of integration with TFs. Genes such as FOXQ1 [Liu et al 2017] and NCOA7 [Xie et al 2016] were involved in the invasion of various cancer cells, but these genes may be linked with invasion of CCRCC cells. Methylation inactivation of tumor supressor EFHD1 was associated with development of colorectal cancer [Takane et al 2014], but loss of this gene may be responsible for advancement of ccRCC.

CANX, SHMT2, IFI16, P4HB, CALU, CDH1, ERBB2, NEDD4L, TFAP2A and SORT1 were screened. CALU (calumenin) was asspciated with invasion of lung cancer cells [Nagano et al 2015], but this gene may be involved in the progression of ccRCC. These include high expression of, TFAP2A, SHMT2, IFI16, P4HB and CALU were predicting shorter survival of ccRCC, while low expression of CANX, CDH1, ERBB2, NEDD4L, and SORT1 were predicting longer survival of CCRCC. Low expression of genes such as CDH1, ERBB2, NEDD4L, TFAP2A and SORT1 were associated with advancement of ccRCC, while high expression of genes such as CANX, SHMT2, IFI16, P4HB and, CALU were associated with advancement of ccRCC. All 10 hub genes were showed altered expressed in all four stage of ccRCC. The mutation analysis found that mutations or alterations in all ten hub genes could result in importantely reduced disease-free survival and poorer clinical outcomes in ccRCC. All ten hub genes were validated by ICH analysis and RT-PCR. ROC curve analysis revealed that the combination of these hub genes was a good predictive model for ccRCC. Immune infiltration analysis revealed that hub genes were positively and negatively associated with tumor purity.

In conclusion, a risk assessment tool for assessing the prognosis of patients with ccRCC was developed and validated in the current study. The 10 identified prognostic genes were associated with several cellular processes and signaling pathways and may be recommended as promising prognostic biomarkers or a diagnostic biomarkers or therapeutic targates of ccRCC. The current risk assessment tool may help identify patients with a high risk of ccRCC, and the proposed prognostic and diagnostic genes as well as therapeutic targates may help elucidate the progression of ccRCC.

## Acknowledgement

I thank Sunil Sudarshan, UAB, Urology, 510 20th Street South, Birmingham, Alabama, USA, very much, the author who deposited their microarray dataset, GSE105261, into the public GEO database.

## Ethical approval

This article does not contain any studies with human participants or animals performed by any of the authors.

## Informed consent

No informed consent because this study does not contain human or animals participants.

## Author Contributions

Basavaraj Vastrad was associated with methodology, manuscript preparation, review and editing. Chanabasayya Vastrad was performed software, supervision, formal analysis and validation.

## Availability of data and materials

The datasets supporting the conclusions of this article are available in the GEO (Gene Expression Omnibus) (https://www.ncbi.nlm.nih.gov/geo/) repository. [(GSE105261) (https://www.ncbi.nlm.nih.gov/geo/query/acc.cgi?acc=GSE105261)]

## Consent for publication

Not applicable.

## Author Contributions

B. V. - Writing original draft, and review and editing

C. V. - Software and investigation

I. K. - Supervision and resources

